# Library Preparation and Sequencing Platform Introduce Bias in Metagenomic-Based Characterizations of Microbiomes

**DOI:** 10.1101/592154

**Authors:** Casper S. Poulsen, Claus T. Ekstrøm, Frank M. Aarestrup, Sünje J. Pamp

## Abstract

Metagenomics is increasingly used to describe microbial communities in biological specimens. Ideally, the steps involved in the processing of the biological specimens should not change the microbiome composition in a way that it could lead to false interpretations of inferred microbial community composition. Common steps in sample preparation include sample collection, storage, DNA isolation, library preparation, and DNA sequencing. Here we assess the effect of three library preparation kits and two DNA sequencing platforms. Of the library preparation kits, one involved a polymerase chain reaction (PCR) step (Nextera), and two were PCR-free (NEXTflex and KAPA). We sequenced the libraries on Illumina HiSeq and NextSeq platforms. As example microbiomes, we assessed two pig fecal samples and two sewage samples of which aliquots were stored at different storage conditions (immediate processing and storage at −80°C). All DNA isolations were performed in duplicate, totaling 80 samples excluding controls. We found that both library preparation and sequencing platform had systematic effects on the inferred microbial community composition. The different sequencing platforms introduced more variation than library preparation and freezing the samples. The results highlight that all sample processing steps need to be considered when comparing studies. Standardization of sample processing is key to generate comparable data within a study, and comparisons of differently generated data, such as in a meta-analysis, should be performed cautiously.

**Importance:** Previous research has reported effects of sample storage conditions and DNA isolation procedures on metagenomics-based microbiome composition; however, the effect of library preparation and DNA sequencing in metagenomics has not been thoroughly assessed. Here, we provide evidence that library preparation and sequencing platform introduce systematic biases in the metagenomic-based characterization of microbial communities. These findings suggest that library preparation and sequencing are important parameters to keep consistent when aiming to detect small changes in microbiome community structure. Overall, we recommend that all samples in a microbiome study are processed in the same way to limit unwanted variations that could lead to false conclusions. Furthermore, if we are to obtain a more holistic insight from microbiome data generated around the world, we will need to provide more detailed sample metadata, including information about the different sample processing procedures, together with the DNA sequencing data at the public repositories.

## Introduction

Microbes are omnipresent and inhabit even the most extreme environments on Earth. Metagenomics-based analyses have provided unprecedented insight into these microbial communities. Metagenomics is applied heavily to human microbiomes, as well as animal and environmental microbiomes, and is being implemented to understand disease states (1–4), for diagnostic purposes (5), and surveillance of pathogens and antimicrobial resistance (6–9). The data from such studies are a growing resource that can be utilized in meta-analysis and data mining, revolutionizing medicine, agriculture, and food production (6, 9–12).

Findings from microbiome studies can be difficult to replicate as observed in different meta-analyses of 16S rRNA gene amplicon studies (13–16). Considering the large number of features (including functional and taxonomic) under investigation in metagenomics, it is not surprising that studies do not seem to lack significant results (17). Data dredging is a real concern in metagenomics, which brings to mind the “replication crisis” that has been highlighted in the field of psychology (18, 19). Due to the challenges of replicating results, one must not over-emphasize the results from exploratory research and keep in mind that there is a need to continually validate the robustness and ability to replicate results in microbiome studies (20, 21). With the improvement of genome reference databases and bioinformatics tools, the validation is an ongoing process (22–25).

Technical variation due to sample processing is an important factor that researchers have to minimize to make proper inferences in metagenomics studies. For example, the DNA isolation procedure has been shown to impact on microbiome composition (26–28). The effect of library preparation and sequencing platform has been investigated in metagenomics primarily on human fecal samples. Library preparation was found to affect taxonomic and functional characterization of human fecal samples and *in silico* constructed mock communities (21, 29). In a study by Costea et al. (26), the effect of library preparation was found to be lower compared with DNA isolation and intra- and inter-sample variation in general. The choice of sequencing platform also appears to have an effect on the characterization of microbiomes (30).

The aim of the present study was to assess the effect of library preparation (Nextera, KAPA PCR-free, NEXTflex PCR-free) and sequencing platform (Illumina HiSeq and NextSeq) on the metagenomics-based inference on DNA samples from two different pig feces and two different sewage microbiomes from a previous study (31). We show that library preparation and sequencing platform introduce systematic bias in the inferred microbial community composition for both sample types and that this effect is important when comparing similar samples, such as pig feces, in the present study. This highlights the need for consistent sample processing and demonstration of caution when comparing data from different studies.

## Results

A subset of DNA samples was selected from a large-scale study (31), to assess the effect of library preparation and DNA sequencing on inferred microbiome composition based on metagenomics. The DNA samples originated from two pig fecal samples (Pig feces 1 and Pig feces 2) and two sewage samples (Sewage 1 and Sewage 2). For the present study, we selected DNA aliquots from fecal and sewage samples that originally were processed immediately (time point 0) and were subjected to storage at −80C, respectively, to not only assess whether one can distinguish different samples, but also samples that have the same origin but exhibit differences due to different handling conditions. The DNA aliquots underwent a total of four different strategies for library preparation and DNA sequencing (Figure 1A), namely KAPA PCR-free library preparation + sequencing on a HiSeq (KAPA HiSeq), NEXTflex PCR-free library preparation + sequencing on a HiSeq (NEXTflex HiSeq), NEXTflex PCR-free library preparation + sequencing on a NextSeq (NEXTflex NextSeq), and Nextera library preparation + sequencing on a NextSeq (Nextera NextSeq). The latter sequencing strategy was performed twice (Nextera 1 NextSeq and Nextera 2 NextSeq).

**Fig. 1.**
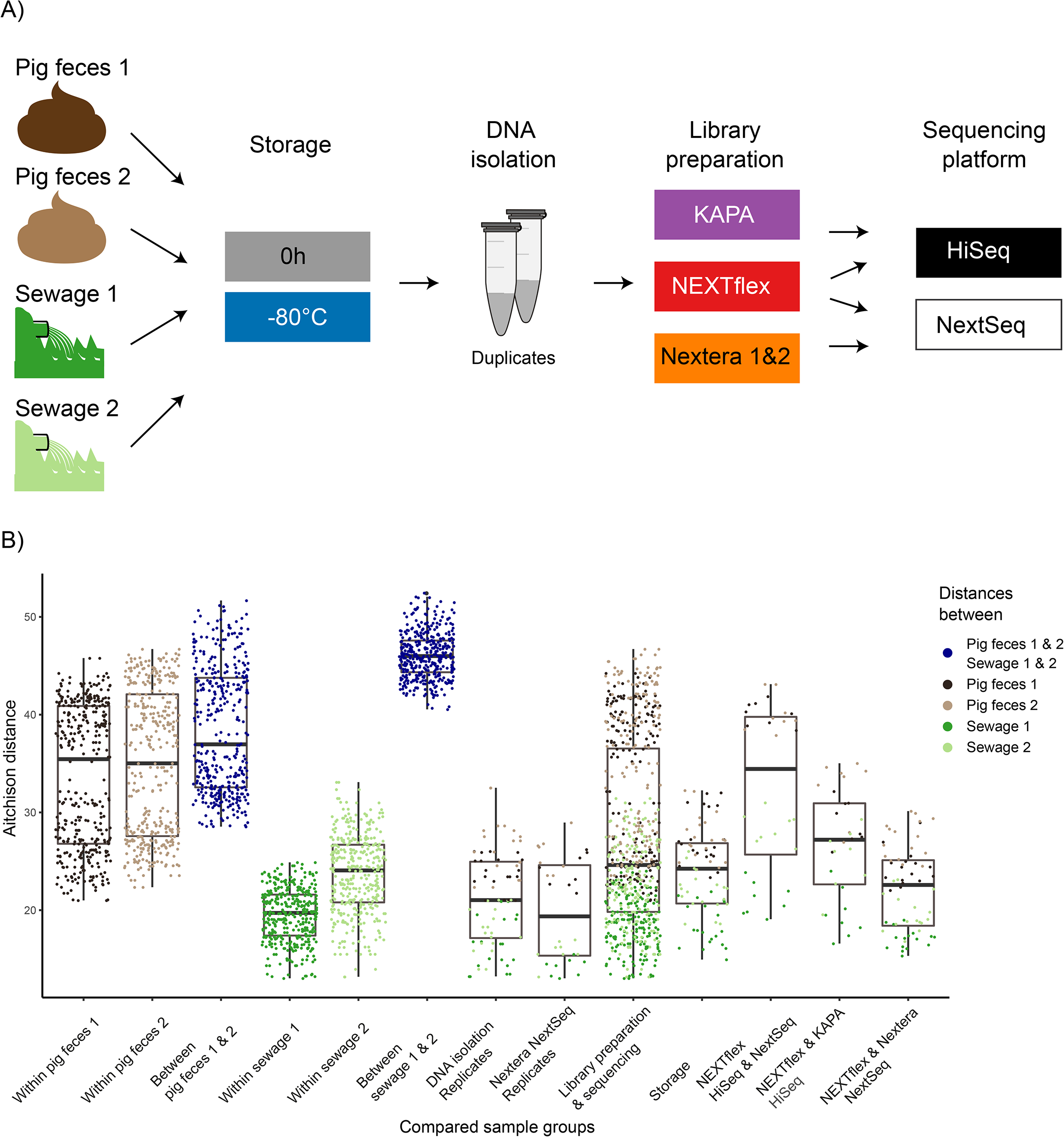
Study design & Comparison between sample groups. A) Two pig feces samples and two sewage samples were processed directly or after storage at −80 °C for 64 hours. The DNA isolation was performed in duplicates, respectively. Library preparation and sequencing was performed in four different combinations: NEXTflex PCR-free library preparation + sequencing on a HiSeq (NEXTflex HiSeq), KAPA PCR-free library preparation + sequencing on a HiSeq (KAPA HiSeq), NEXTflex PCR-free library preparation + sequencing on a NextSeq (NEXTflex NextSeq), and Nextera library preparation + sequencing on a NextSeq (Nextera NextSeq). The latter sequencing strategy was performed twice (Nextera 1 NextSeq, and Nextera 2 NextSeq). The setup resulted in a total of 80 metagenomes plus five negative controls (i.e. DNA extraction controls). B) The boxplots displays pairwise Aitchison distances between different groupings of samples. Within the different groups, dots representing the distances were colored according to which sample the comparison was made in. Blue dots represent a distance between two different samples.

### Quality control of sequencing output

The number of raw reads from the different library preparations and sequencing platforms were similar with about a factor two difference when comparing the medians. The highest number of reads was obtained from the NEXTflex HiSeq run (median: 12.1, range: 6.3 – 30.8 million reads) and the lowest from the NEXTflex NextSeq run (median: 7.6, range: 2.7 – 9.4 million reads) (Table S1). The outputs from the KAPA HiSeq run (median: 9.4, range: 7.8 – 17.4 million reads) and the Nextera NextSeq runs (median: 10.2, range: 6.5 – 16.5 million reads) were about the same. More reads were obtained from the pig fecal samples compared with the sewage, but a larger proportion of the sewage reads mapped to the reference databases. The microbial community of the sewage samples exhibited a higher Simpson diversity than the pig feces (Table S1). The number of mapped reads were higher for the sewage samples, and many of the samples had reached a plateau as observed when drawing a rarefaction curve (Figure S1). Similar results were obtained when comparing the mean of the percent of unmapped reads of the same sample across the different library preparation and sequencing platform runs (Pig feces 1: 87.4-88.4, Pig feces 2: 89.7-90.5, Sewage 1: 70.1-74.1, Sewage 2: 54.2-59.3) (Table S1).

### Sample processing impacts on inferred microbiome structure

The pairwise Aitchison distances were calculated between all the samples and visualized using PCA (Figure S2A). The sample type explained the greatest variance, and pig feces and sewage samples were clearly separated on the first axis. A clear separation of the two sewage samples was observed on the second axis, while the two pig fecal samples clustered together. Ordination of the pig feces and sewage samples separately revealed that it was possible to differentiate the two pig fecal samples (Figure S2B). However, there were also two clusters within each pig fecal sample. A clear separation of the two sewage samples was still observed (Figure S2C). Also in a boxplot visualization, library preparation, sequencing platform, and storage condition did not hamper the ability to differentiate between the two sewage samples (Figure 1B). However, we observed an overlap between pig feces 1 and 2 comparisons, relative to comparing within the two samples representing the effect of the different sample processing parameters. Nevertheless, the median suggested there is a difference between pig feces 1 and 2 (Figure 1B). In general, larger distances were calculated for the comparisons of sample processing parameters in pig fecal samples compared with sewage. The shortest distances were observed when comparing the DNA isolation replicates and the replicates of the Nextera NextSeq runs, respectively. The distances between samples that differed in library preparation and sequencing platform were greater, compared with samples that differed in storage conditions (i.e. whether they were processed directly or after freezing at −80 °C for 64 hours). The sequencing platform appeared to be a major contributor of variation when comparing the samples that were prepared with NEXTflex and sequenced on both, an Illumina HiSeq and an Illumina NextSeq (Figure 1B). Whereas, using two different preparation kits, i.e. NEXTflex and KAPA, sequenced on an Illumina HiSeq introduced a relatively lower variation. The differences were observed to be even lower when samples were prepared with the two library preparation kits NEXTflex and Nextera and sequenced on an Illumina NextSeq (Figure 1 B).

To investigate the effect of sample processing further, principal component analyses (PCAs) were performed for the individual samples (Pig feces 1, Pig feces 2, Sewage 1 and Sewage 2). Similar patterns were observed in all samples, indicating that there was a systematic effect from sequencing platform, library preparation, and storage condition (Figure 2). The samples clustered primarily according to sequencing platform and library preparation along the x-axis that represents most of the variation. On the y-axis samples clustered according to storage condition. In general, the DNA isolation replicates were similar, as well as the two Nextera NextSeq runs (Figure 2). All the parameters had a significant effect based on permutational multivariate analysis of variance (PERMANOVA), except for storage when comparing all of the samples and in pig feces 2 (Table 1). The percent variation in pig feces attributed to sample (Pig feces 1 and Pig feces 2) (21.1%) and sequencing platform (19.1%) were at similar levels, further emphasizing the importance of sample processing when comparing communities that are more similar to each other (Table 1).

**Fig. 2.**
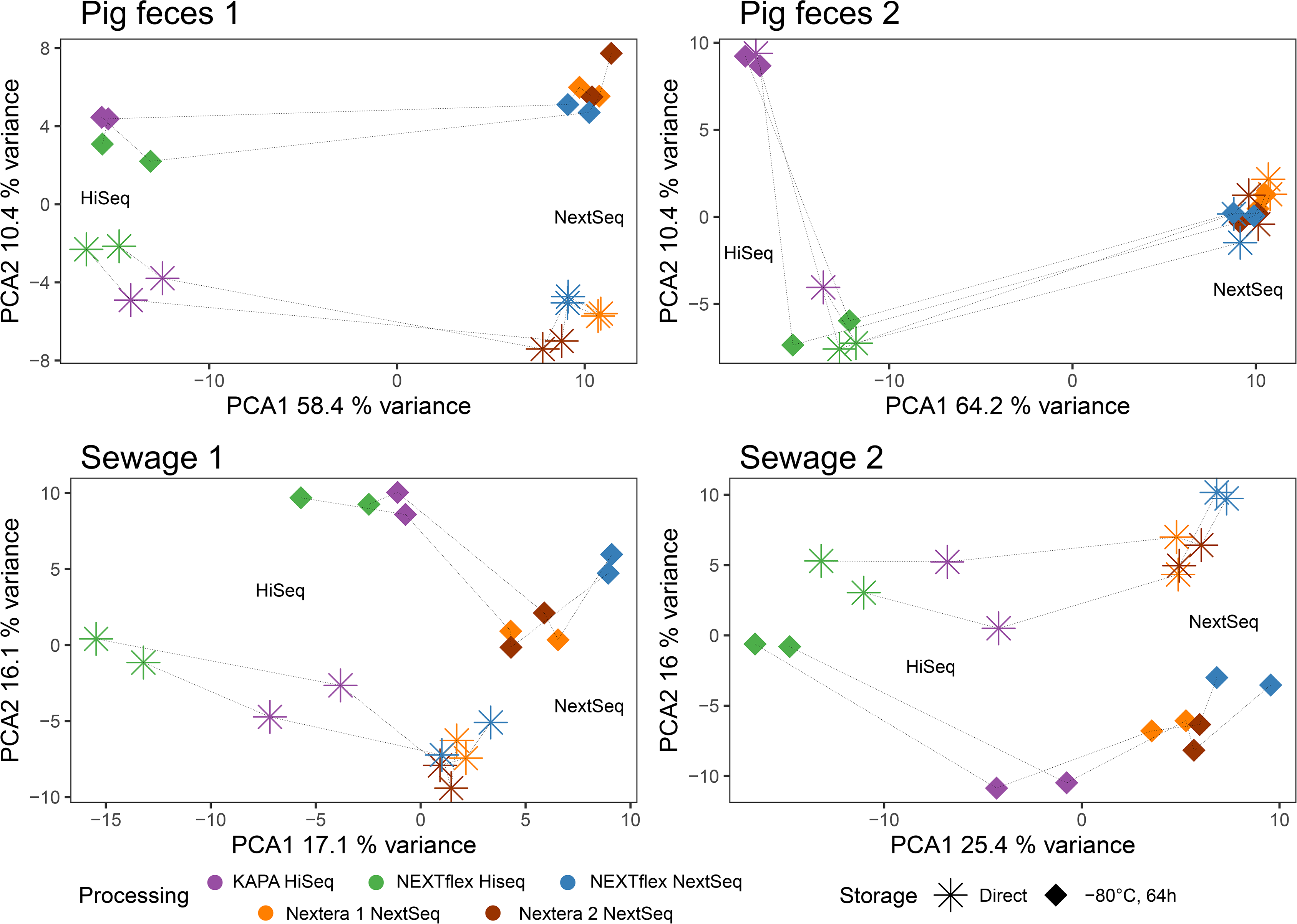
Principal component analysis (PCA) subset to the different sample matrices. Euclidean distances were calculated after performing centered log-ratio transformation (CLR) of the count data (Aitchison distances). Variance explained by the two first axes are included in their labels. The same DNA samples processed differently are connected with dotted lines.

**Table 1:** Effect of sample origin (Pig feces 1, Pig feces 2, Sewage 1 and Sewage 2) and different parameters in sample processing (Library preparation, DNA sequencing). Statistical tests were performed by multiple permutations partitioning sum of squares (PERMANOVA). The *p-value*, as well as the percent of variation explained by the parameters, is reported, testing different sample sets (all, pig feces, sewage, Pig feces 1, Pig feces 2, Sewage 1 and Sewage 2).

| Samples included | Sample<br>*p-value* (%<sup>a</sup>) | Storage<br>*p-value* (%<sup>a</sup>) | Library preparation<br>*p-value* (%<sup>a</sup>) | Sequencing platform<br>*p-value* (%<sup>a</sup>) |
| --- | --- | --- | --- | --- |
| All | <10<sup>-5</sup> (81.9) | 6.4×10<sup>-2</sup> (0.5) | 4.2×10<sup>-2</sup> (1.0) | 3.0×10<sup>-4</sup> (1.8) |
| Pig feces | <10<sup>-5</sup> (21.1) | 3.8×10<sup>-3</sup> (3.3) | 5.7×10<sup>-4</sup> (6.2) | <10<sup>-5</sup> (19.1) |
| Sewage | <10<sup>-5</sup> (61.7) | 2.5×10<sup>-2</sup> (2.9) | 3.0×10<sup>-2</sup> (4.1) | 4.4×10<sup>-3</sup> (4.5) |
| Pig feces 1 | NA<sup>b</sup> | 2.8×10<sup>-3</sup> (9.7) | 2.8×10<sup>-2</sup> (8.9) | <10<sup>-5</sup> (26.2) |
| Pig feces 2 | NA | 0.17 (2.7) | 5.4×10<sup>-3</sup> (12.3) | <10<sup>-5</sup> (25.3) |
| Sewage 1 | NA | <10<sup>-5</sup> (15.1) | 3.6×10<sup>-4</sup> (14.4) | <10<sup>-5</sup> (12.8) |
| Sewage 2 | NA | <10<sup>-5</sup> (14.0) | 6.0×10<sup>-5</sup> (17.8) | <10<sup>-5</sup> (19.6) |

<sup>a</sup>Proportion of the variation explained in the PERMANOVA
<sup>b</sup>Not Applicable, no *p-value* obtained when variable subset to a single sample (Pig feces 1, Pig feces 2, Sewage 1 and Sewage 2).

### Sample processing impacts on inferred microbial abundances

To investigate the effect of library preparation and sequencing platform on the abundance of specific microorganisms, an overview of the 30 most abundant genera was visualized in heatmaps (Figure 3). For the pig samples, the aliquots appeared to cluster mainly based on the sequencing platform (NextSeq vs. HiSeq) (Figure 3A). In contrast, it was possible to distinguish the two sewage samples, who clustered according to sample origin (Sewage 1 vs. Sewage 2) (Figure 3B). A clustering of samples was also observed to a certain degree for both pig feces and sewage according to storage condition and library preparation.

**Fig. 3.**
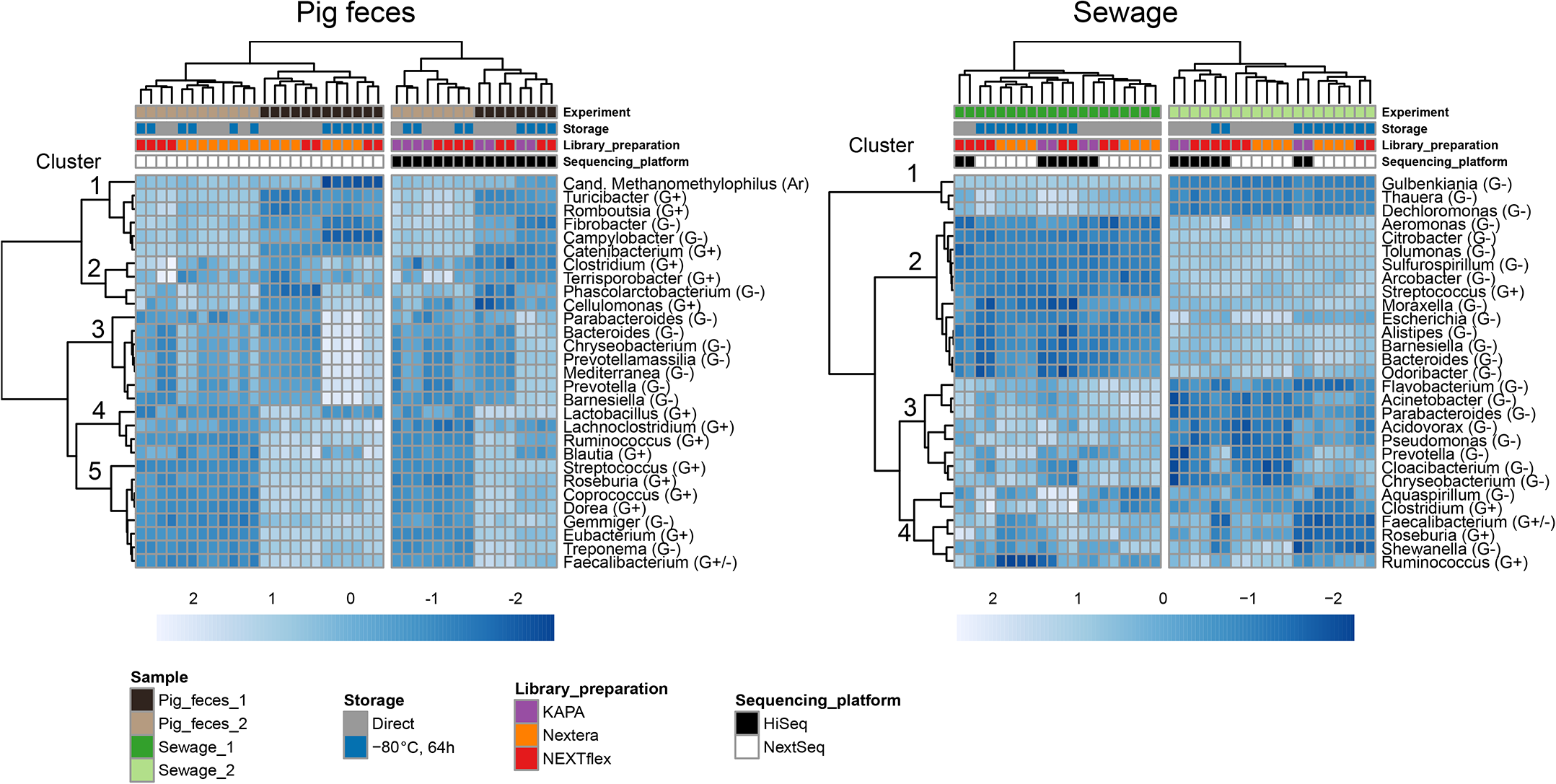
Heatmaps of pig feces and sewage samples separately with the 30 most abundant genera. Complete-linkage clustering was performed to create dendograms for both genera and samples. Spearman correlation was used to cluster the genera and Aitchison distances were used to cluster the samples. Genera abundance depicted in the cells were CLR transformed counts standardized to zero mean and unit variance. Grouping of organisms were included in genera names according to cell wall structure based on Gram positive staining (G+), Gram negative staining (G-) or belonging to Archaea (Ar). (A) Heatmap of all pig feces samples, where the first branching was according to sequencing platform. The third cluster of genera exclusively contained Gram–negatives. (B) Heatmap of all sewage samples. The fourth cluster mainly consisted of Gram–positives. A few Gram–positives were also present in the other clusters.

The pig feces contained both Gram-negative and Gram-positive bacteria, and cluster 3 exclusively consisted of Gram-negatives. There were a few Gram-negatives in the other clusters, indicating that sample processing shifts the abundance profiles for specific groups of organisms, in this case, it appeared to be associated with cell wall structure (Figure 3A). A similar pattern was observed for sewage that mainly consisted of Gram-negatives. The majority of Gram-positives were part of cluster 4, including *Clostridium*, *Faecalibacterium*, *Roseburia* and *Ruminococcus*. However, this cluster also contained Gram-negative genera (Figure 3B).

One explanation for the community differences observed by sample processing could be a possible contamination during the library preparation and sequencing steps. To elucidate this, sparse partial-least-squares discriminant analysis (sPLS-DA) was performed, assessing which genera best characterize the library preparation and sequencing platform processing methods. Components 1, 2 and 3 were included in the model containing 5, 50 and 20 different genera, respectively (Figure S3). The majority of microorganisms were abundant organisms observed across all of the sample processing methods. However, a few were clear indicators of contamination during library preparation and sequencing and were mainly present in a single processing method. This included *Methylobacterium* in the KAPA HiSeq run and *Cutibacterium* in the second Nextera NextSeq run, bacteria that previously have been associated with kit contamination (32). A heatmap of the 30 most abundant genera in the blank controls additionally revealed a high abundance of *Ralstonia* in the Nextera NextSeq runs that where performed with the same kit reagents (Figure S4). Overall, the organisms associated with contamination were limited. The separation of the samples according to the different processing parameters therefore appeared to be real changes to the relative abundances between organisms inherently present in the microbiomes and not due to contamination.

A constrained ordination analysis, also subset according to whether samples were processed directly or after freezing, was performed to assess whether groups of organisms at a taxonomically higher level were associated with a specific library preparation and sequencing method. In the pig feces, Proteobacteria seemed associated with the HiSeq runs (Figure S5). However, this was not observed for sewage. For sewage, Archaea were associated with the HiSeq runs, but also Eukaryotes consisting of Fungi and *Cryptosporidium* seemed associated with the HiSeq runs in sewage 1 (Figure S5). Overall, it was difficult to observe a pattern when assessing this grouping of genera, highlighting that it might be difficult to generalize the effect of sample processing in different sample types and different samples of the same type.

## Discussion

With the increasing amount of metagenomic data in public repositories, meta-analysis and pooling of data from different studies are exciting new opportunities to gain further insight into the microbial world (10–12, 24, 33). Data generation is usually not performed with a standard procedure across studies, and sample processing is an important factor to be aware of when trying to make inferences in these cross-study investigations (21, 26). In the present study, both library preparation and sequencing platform had a significant effect on explaining the variance in the data (Table 1). That these parameters affect the community description has also been observed previously (21, 29, 30). In the study by Costea et al. (26), DNA isolation had the largest effect compared with other technical variations. In the present study, DNA isolation was performed centrally by the same person, while library preparation and sequencing was performed in-house or at external providers, but not in any of the cases by the same person, possibly increasing variation due to DNA shipping and handling in this specific step. When performing a validation study assessing the technical variation of sample processing, the large number of methodologies and variations thereof make it impossible to test all parameters. It is likely that selecting methods that are based on different principles and for specific purposes yield results that highlight the importance of this specific step. Bowers et al. (29) investigated community changes using different amounts of input DNA, and observed that this modification had a significant effect on community description. This effect can increase bias associated with library preparation and sequencing platform in other studies were starting material is of variable quality. In the present study, investigation of sequencing platforms were limited to the NextSeq and HiSeq, which are both Illumina platforms resembling each other in technology, and which were selected due to their popularity in metagenomics with low cost relative to output (34). However, the platforms have been reported to exhibit differences in index hopping (35). In the present study, a large effect was attributed to the sequencing platform and that was also observed when using the same library preparation kit (NEXTflex PCR-Free) (Figure 1B). The library preparation included two methods that required pre-fragmented DNA that was prepared PCR-free (KAPA and NEXTflex). It was decided to include the Illumina Nextera library preparation as well, to compare with a technique that does not resemble the others in having enzymatic fragmentation, and which involved a PCR step that is commonly applied when not enough DNA is available to prepare DNA for sequencing PCR-free. However, the two Nextera runs were relatively similar compared with the NEXTflex run when sequenced on the NextSeq (Fig. 2). The present study was not a full factorial experiment and this should be emphasized when comparing the effect sizes of specific processing parameters.

One explanation for the differences observed between the processing runs can be contamination bias. When designing a metagenomics study, it is to some extent possible to remove kit contaminations or carry-over between sequencing runs from the data *in-silico*, if for instance, blank controls are included or by rotating indexing primers between adjacent runs, respectively (36). In the present study, comparing the sPLS-DA results with the blank controls rarely identified the same genera, indicating that the genera reported to explain the specific sample processing the most were not due to contamination during DNA extraction. The general variation associated with redoing the library preparation and sequencing was low when comparing the two Nextera sequencing runs (Figures 1 and 2). The differences observed are therefore most likely due to true variation associated with the sample processing. Furthermore, it was possible to detect that these patterns were systematic in the different samples (Figure 2), and that this could be partly explained with some crude features such as distinguishing between Gram-negative and Gram-positive bacteria or at a higher taxonomic classification (Figures 3 and S5). The grouping of genera into Gram-negative and Gram-positive might be confounders of an underlying explanation that could be associated with DNA characteristics such as guanine-cytosine percent (GC%) or other specific DNA patterns. Another possibility is that DNA fragmentation during sampling, storage and DNA isolation provided DNA of different quality for specific organism groups. A shift in community structure is then reflected in the selection of different fragment sizes during the library preparation and sequencing. Practical limitations were also an issue when designing the study. To reduce the bias associated with DNA extraction, the QIAamp Fast DNA Stool Mini Kits were all ordered together ensuring that kits were from the same manufacturing batch. Another possible bias might arise from DNA samples that were frozen in between processing them for sequencing. However, only small changes were observed between the two Nextera NextSeq runs.

The Aitchison distances obtained from comparing within the two pig fecal samples separately relative to within the two sewage samples also revealed that storage, library preparation and sequencing platform has a larger effect in pig feces (Figure 1B). Since the distances between the two pig fecal samples were smaller relative to the distances between the two sewage samples, it was difficult to discern the two pig fecal samples when samples were processed differently (Figure 3). It is concerning that the variation due to sample processing might hamper the ability to differentiate between two different pig fecal samples, and this might obstruct the ability to draw meaningful conclusions when technical variations cannot be distinguished from true changes. These results should on the other hand not be overstated; the two pig fecal samples were obtained from an in-bred race raised under very similar conditions including feeding, even though they were obtained from two different healthy pigs at two different farms, the two communities are relatively similar. The finding highlights that the importance of technical variation depends on the differences that one is trying to detect (16). The technical variation did not hamper the ability to differentiate between the two sewage samples.

We show that library preparation and sequencing platform introduce systematic bias in the metagenomic-based characterization of microbial communities. These findings suggest that library preparation and sequencing are important parameters to keep consistent when aiming to detect small changes in community structure. In the present study, the bias was somewhat dependent on sample type, highlighting the importance of assessing the effect of sample processing in the specific sample type under investigation.

## Materials and Methods

### Sample processing

A subset of 85 DNA samples was selected from a large-scale study examining the effect samples storage conditions on inferred microbiome composition (31). The DNA samples originated from two pig fecal samples (Pig feces 1 and Pig feces 2) and two sewage samples (Sewage 1 and Sewage 2). The two pig fecal samples were collected on different occasions from different conventional pig production farms near the laboratory. The two sewage samples were collected at a local wastewater treatment facility on different occasions. DNA isolation was performed in duplicate with a modified QIAamp Fast DNA Stool Mini Kit (Qiagen) protocol, including an initial bead beating step (MoBio garnet beads) (27) (Figure 1B). A DNA extraction (blank) control was included at each time of DNA isolation (5 controls as part of the present study). For a detailed list of all samples included in this study, see Table S1 (column H, ‘Other study (Library preparation)’) in Poulsen *et al.* 2021 (31). The concentration of DNA samples was measured with the Qubit dsDNA High Sensitivity (HS) assay kit on a Qubit 2.0 fluorometer (Invitrogen, Carlsbad, CA) before storing the DNA at −20°C.

### Library preparation and sequencing

Library preparation and sequencing were performed in the order described below and the DNA was frozen between the sequencing runs:

#### KAPA PCR-free on a HiSeq (KAPA HiSeq)

DNA was shipped for sequencing to an external provider (Admera Health, New Jersey, USA). The DNA (500 ng) was fragmented mechanically (Covaris E220 evolution, aimed insert size=350bp) using ultrasonication. The KAPA library preparation was run PCR-free according to the manufacturer’s recommendations (KAPA Hyper Prep Kit KR0961 – v6.17). Sequencing was performed on an Illumina HiSeq 4000 (2×150 cycles, paired end).

#### NEXTflex PCR-free on a HiSeq (NEXTflex HiSeq)

DNA was shipped for sequencing to an external provider (Oklahoma Medical Research Foundation, Oklahoma, USA). The DNA (500 ng) was fragmented mechanically (Covaris E220 evolution, aimed insert size=350bp) using ultrasonication. The NEXTflex library preparation was run PCR-free according to the manufacturer’s recommendations (Bioo Scientific NEXTflex PCR Free DNA Sequencing Kit – 5142-01). Sequencing was performed on an Illumina HiSeq 4000 (2×150 cycles, paired end).

#### NEXTflex PCR-free on a NextSeq (NEXTflex NextSeq)

The DNA (500 ng) was fragmented with mechanical fragmentation (Covaris E210, aimed insert size=350bp, duty cvd=10%, Intensity=5, cycle burst=200, treatment time=240 sec.) using ultrasonication. The NEXTflex library preparation was run PCR-free with Nextflex barcodes (NEXTflex-96 DNA barcodes) according to the manufacturer’s recommendations (Bioo Scientific NEXTflex PCR-Free DNA Sequencing Kit – 5142-01). Sequencing was performed in-house on an Illumina NextSeq 500 (mid output v2, 2×150 cycles, paired end).

#### Nextera 1 and 2 on a NextSeq (Nextera 1 NextSeq, Nextera 2 NextSeq)

The Nextera XT library preparation was performed twice. The Nextera XT protocol was carried out according to the manufacturer’s recommendations (Nextera XT DNA Library Prep Kit 15031942v02). This included a tagmentation step that fragments the DNA (1 ng) and ligates adaptors, and a PCR step amplifying DNA and adding indexing primers. Library cleanup was performed with AMPure XP beads and normalized before sequencing was performed in-house on an Illumina NextSeq 500 (mid output v2, 2×150 cycles, paired end). The bioanalyzer results revealed that the aimed insert size of 350 bp was larger than expected (File S6).

### Bioinformatics and statistical analysis

Pre-processing of raw reads included trimming (Phred quality score = 20) and removal of reads shorter than 50bp (BBduk2) (37). Mapping was performed with a Burrows-Wheeler aligner (BWA-mem) as implemented in MGmapper (22). Mapping was performed in the default “best mode” to 11 databases; first filtering against the human database then extracting the number of raw reads mapping to the genomes of bacteria, fungi, archaea, viruses, and Cryptosporidium. A read count correction was implemented to adjust large hit counts to specific contigs as implemented in Hendriksen et al. (9). All counts in the count table were divided by two to account for reads mapping as proper pairs, and then aggregating to genus level. The processed count table, metadata and feature data are available (File S7), and the raw reads are deposited in the European Nucleotide Archive (ENA) (Project acc.: PRJEB31650).

All statistical analyses adhered to the compositional data analysis framework and were performed in R version 3.5.2 (38–40). Alpha diversity was calculated based on the raw count table estimating richness (Chao1), evenness (Pielou’s) and diversity (Simpson) using the diversity function in vegan. Initial filtering of the count matrix was performed by removing all genera below an average count of 5. The estimation of zeroes was performed using simple multiplicative replacement (41). Centered log-ratio transformation (CLR) was used in: principal components analysis (PCA), heatmaps to perform complete-linkage clustering analysis of the samples, boxplots to calculate pairwise Euclidean distances between samples (Aitchison distance), and permutational multivariate analysis of variance (PERMANOVA, with 99999 permutations assessing the marginal effects of the terms), sparse partial-least-squares discriminant analysis (sPLS-DA, with 5 fold cross validation repeated 10 times), and constrained ordination with redundancy analysis (rda) (38, 40, 42, 43). Spearman correlation on CLR transformed data were used to cluster the genera visualized in the heatmap, but the genera abundance depicted in the cells were standardized to zero mean and unit variance after CLR transformation for comparability. Analyses performed are included (File S8) and the code is available from https://github.com/csapou/LibraryPreparationandSequencingPlatform.

## Data availability

The raw reads supporting the conclusions of this article are available at the European Nucleotide Archive (ENA) (Project acc.: PRJEB31650). The processed count data, metadata and feature data are accessible in the supplemental material as File S7, and the statistical analysis performed are provided in the supplemental material in File S8. The statistical analysis is furthermore available at https://github.com/csapou/LibraryPreparationandSequencingPlatform.

## Declarations

### Ethics approval and consent to participate

Not applicable

### Consent for publication

Not applicable

### Authors’ contributions

FA acquired the funding for the project. CP, FA and SP conceived and designed the experiments. CP performed the experiments, performed the literature review and wrote the paper. CP and CE analysed the data. FA, CE and SP revised the manuscript. All authors read and approved the manuscript.

## Acknowledgments

The authors wish to thank Marie Jensen, Berith Knudsen and Carsten Bidstrup for their help with the sampling, and Jacob Jensen and Marlene Dalgaard for technical assistance during in-house library preparation and sequencing. The authors appreciate helpful discussions with Anna Ingham on data analysis. Rolf Kaas is acknowledged for help with the upload of raw sequencing reads to ENA, and Jeffrey Skiby for language editing.

## Supplemental Material

**Table S1. Sequencing quality control and alpha-diversity overview.**

**Figure S1.**
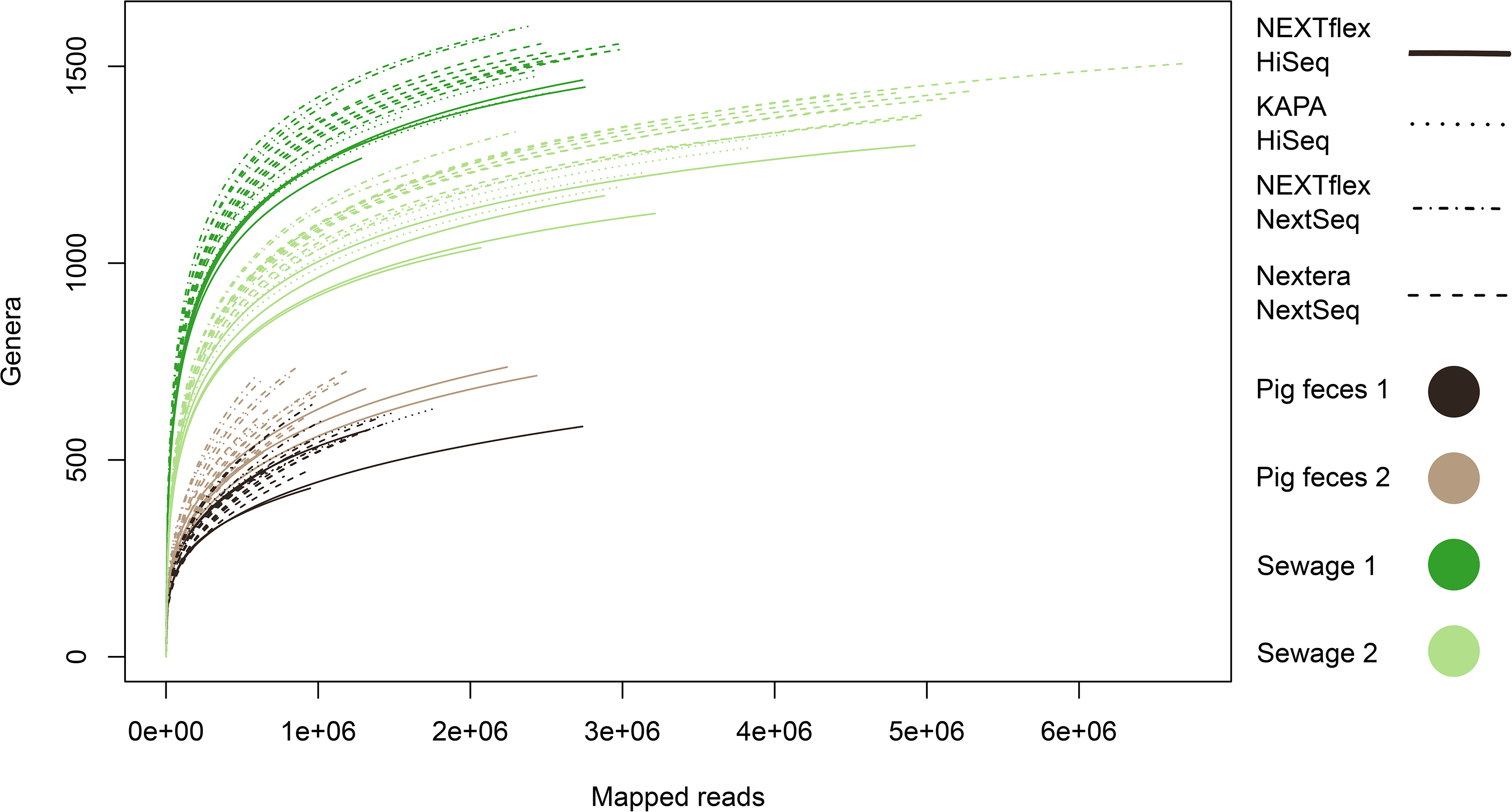
Rarefaction curves.

**Figure S2.**
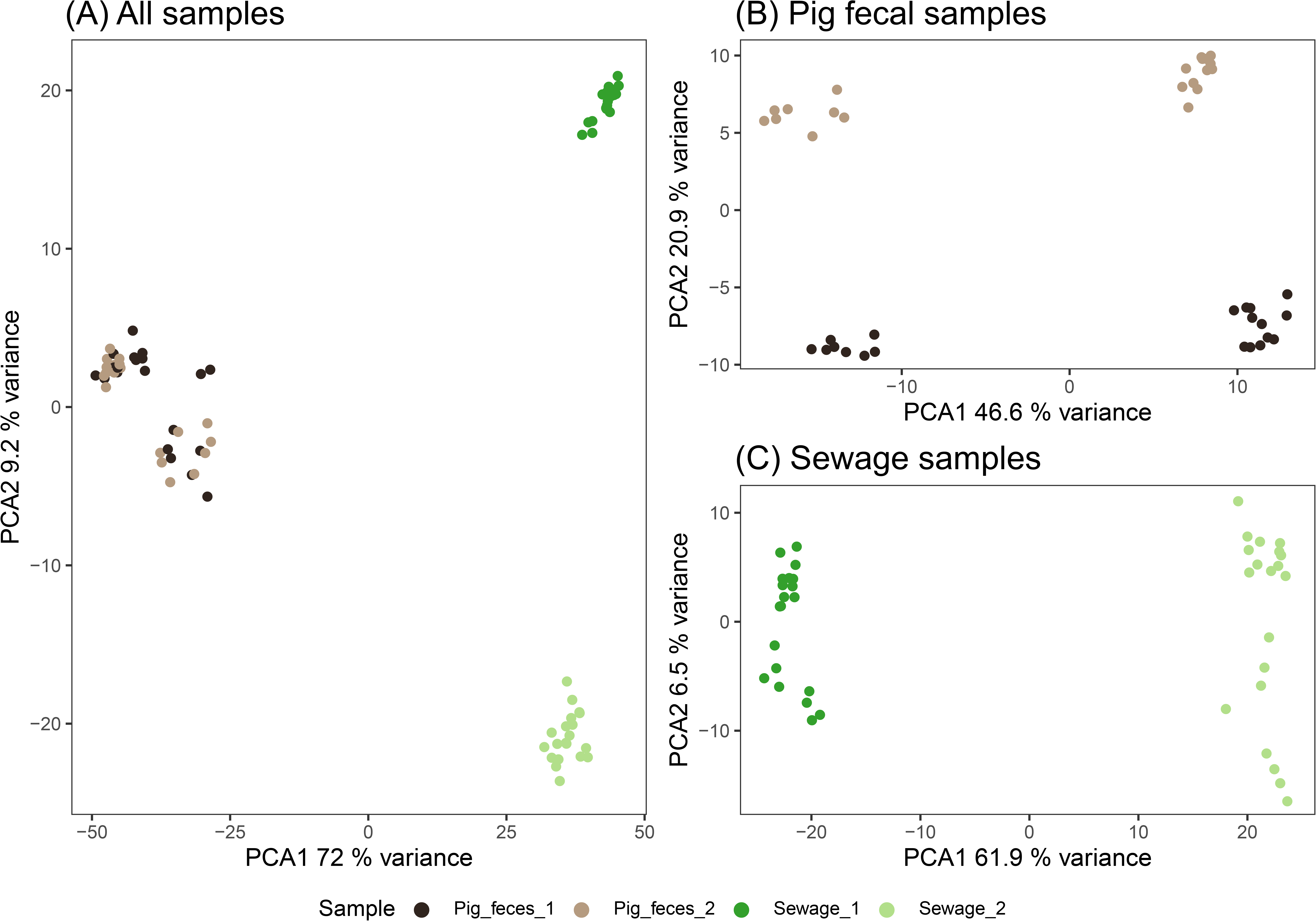
Prinicipal component analysis (PCA) for all samples and subsetted to pig feces and sewage. Variance explained by the two first axes are included in their labels. (A) PCA were generated for all samples forming three clusters with pig feces together and sewage 1 and sewage 2 seperately, (B) PCA were generated for all pig feces samples separation were now observed between pig feces 1 and pig feces 2, and (C) PCA were generated for all sewage samples that were still easy to discriminate.

**Figure S3.**
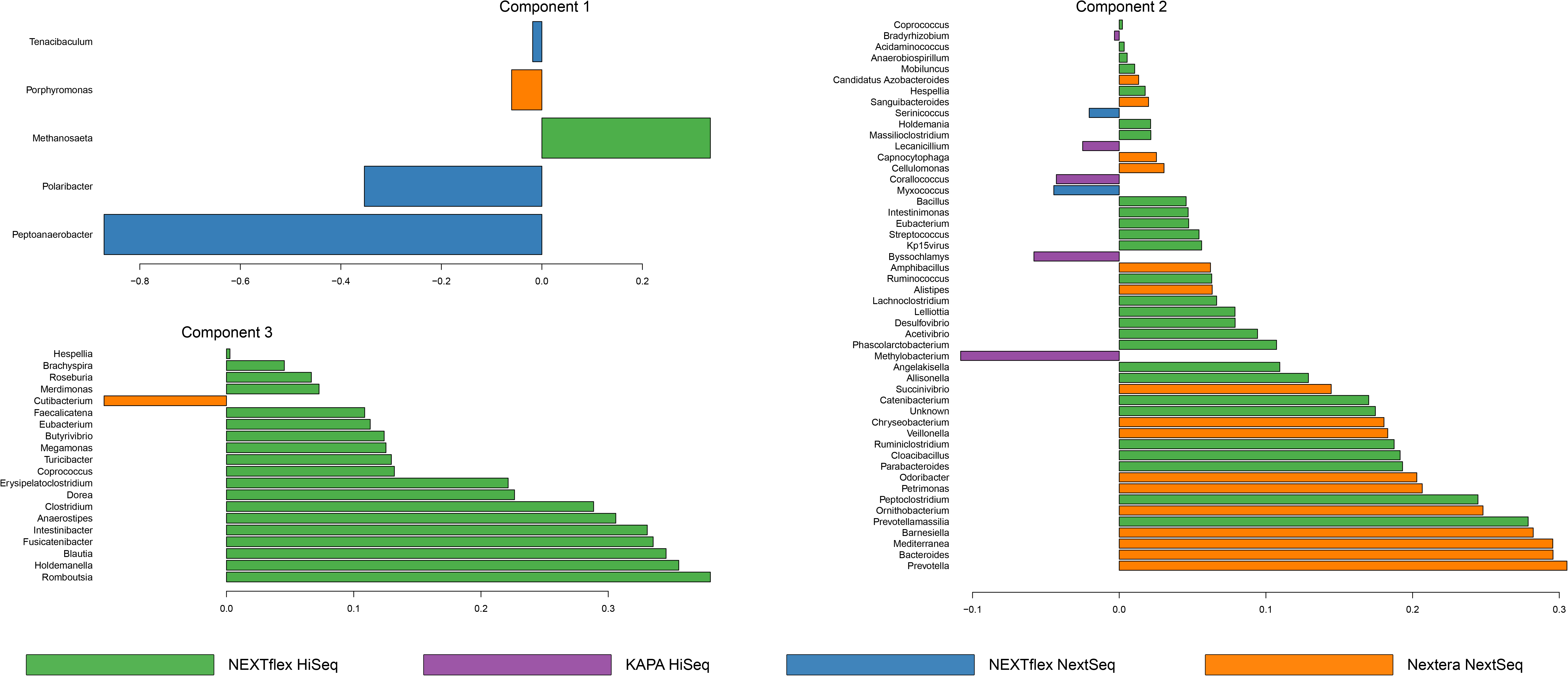
Sparse partial least square discriminant analysis (sPLS-DA). The sPLS-DA were run with a unique identifier for a specific DNA sample that were then processed differently in the generation of libraries and sequencing to select the most discriminative genera in explaining this aspect of sample processing.

**Figure S4.**
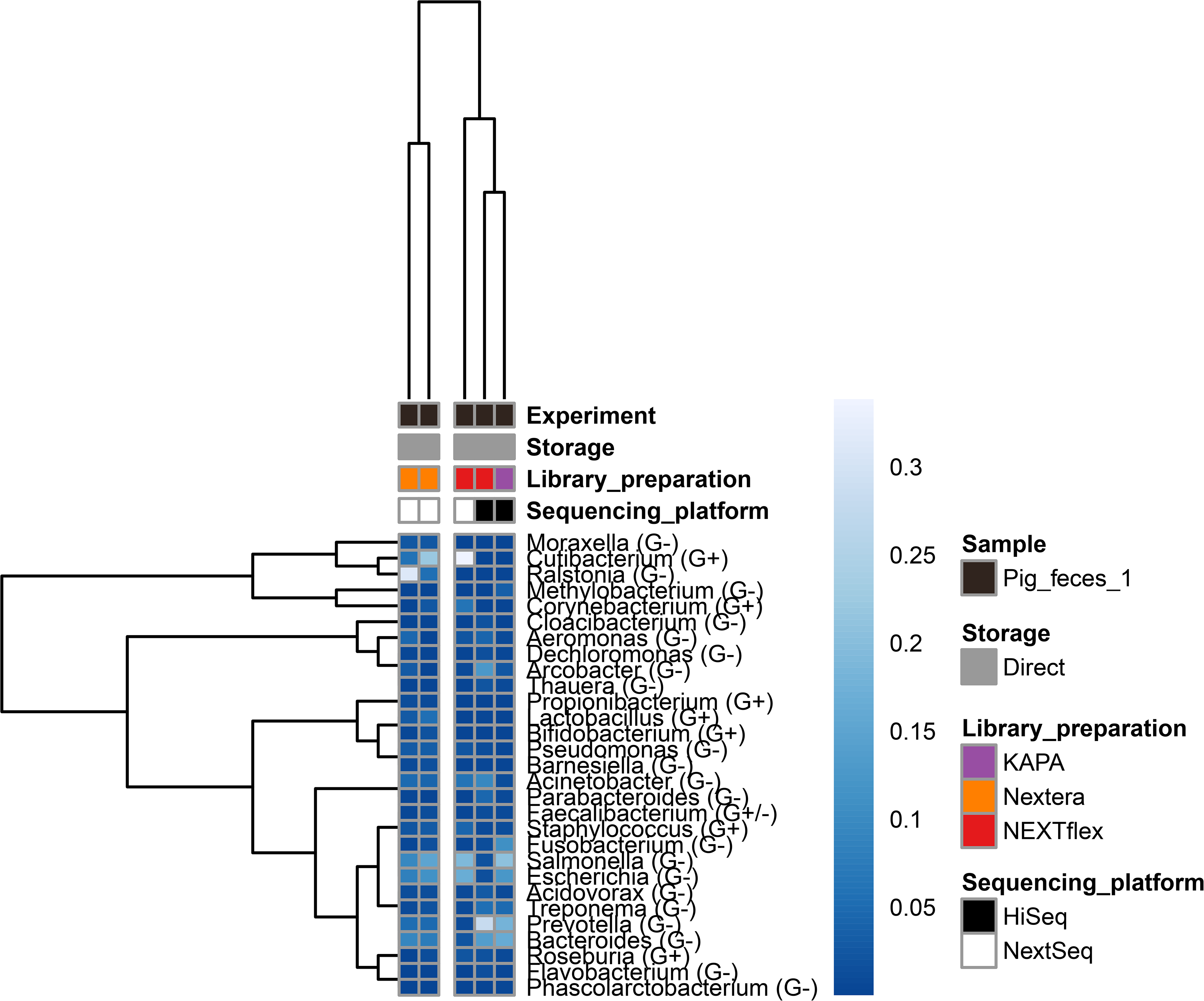
Heatmap of negative controls with the thirty most abundant genera. Complete-linkage clustering was performed to create dendograms for both genera and samples. Spearman correlation was used to cluster the genera and Aitchison distances were used to cluster the samples. Genera abundance depicted in the cells were CLR transformed counts standardized to zero mean and unit variance. Grouping of organisms were included in genera names according to cell wall structure based on Gram-positive staining (G+), Gram-negative staining (G-) or belonging to Archaea (Ar).

**Figure S5.**
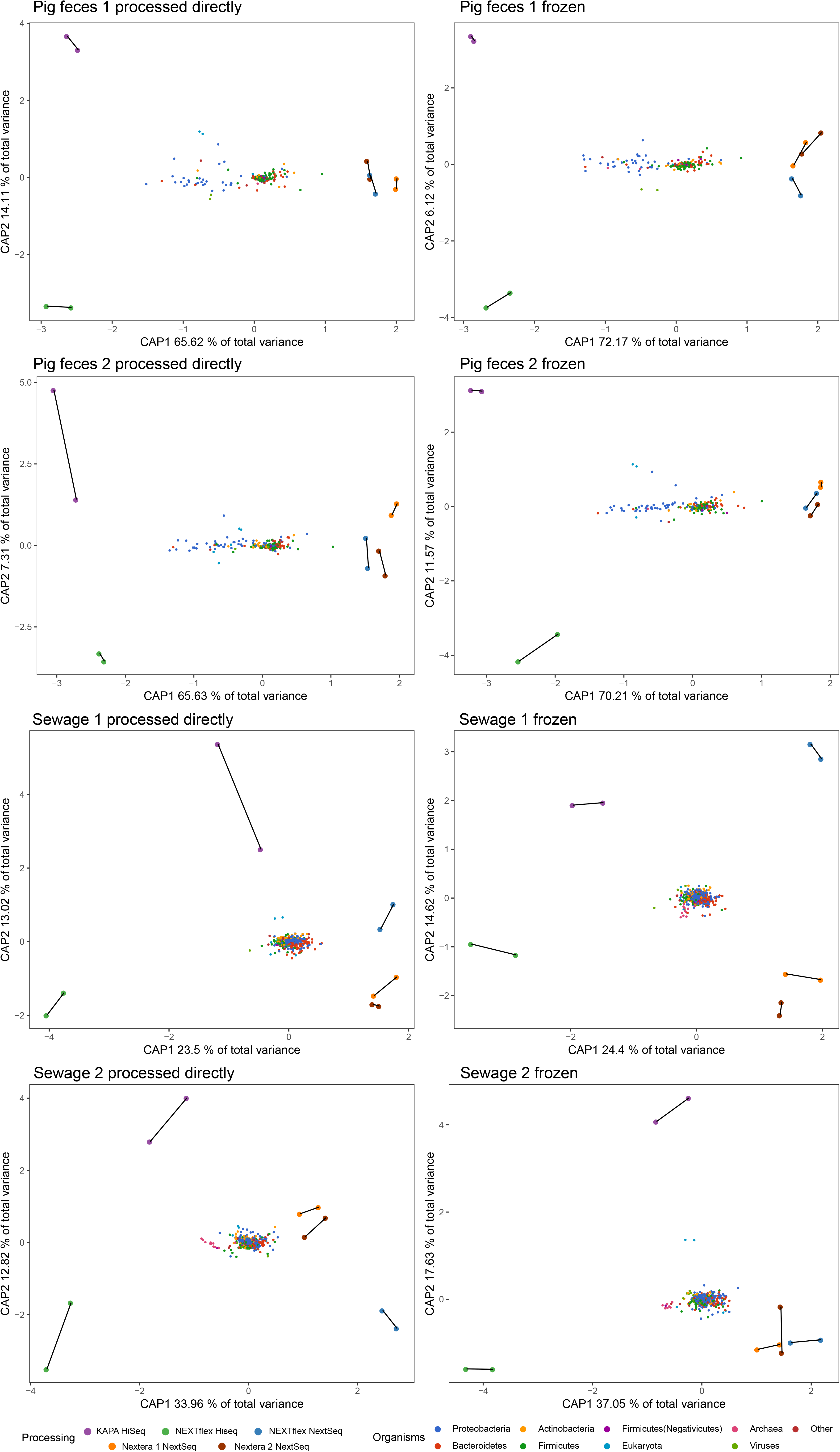
Redundancy analysis (rda) subsetted to sample matrix and if samples were frozen or processed directly. Taxonomic patterns were investigated by plotting genera coloured according to different taxonomic groups. DNA isolation replicates were connected with lines.

**File S6. Bioanalyzer electropherograms.**

**File S7. Count table, metadata and feature data used in the analysis.**

**File S8. R markdown file containing the R analysis.**

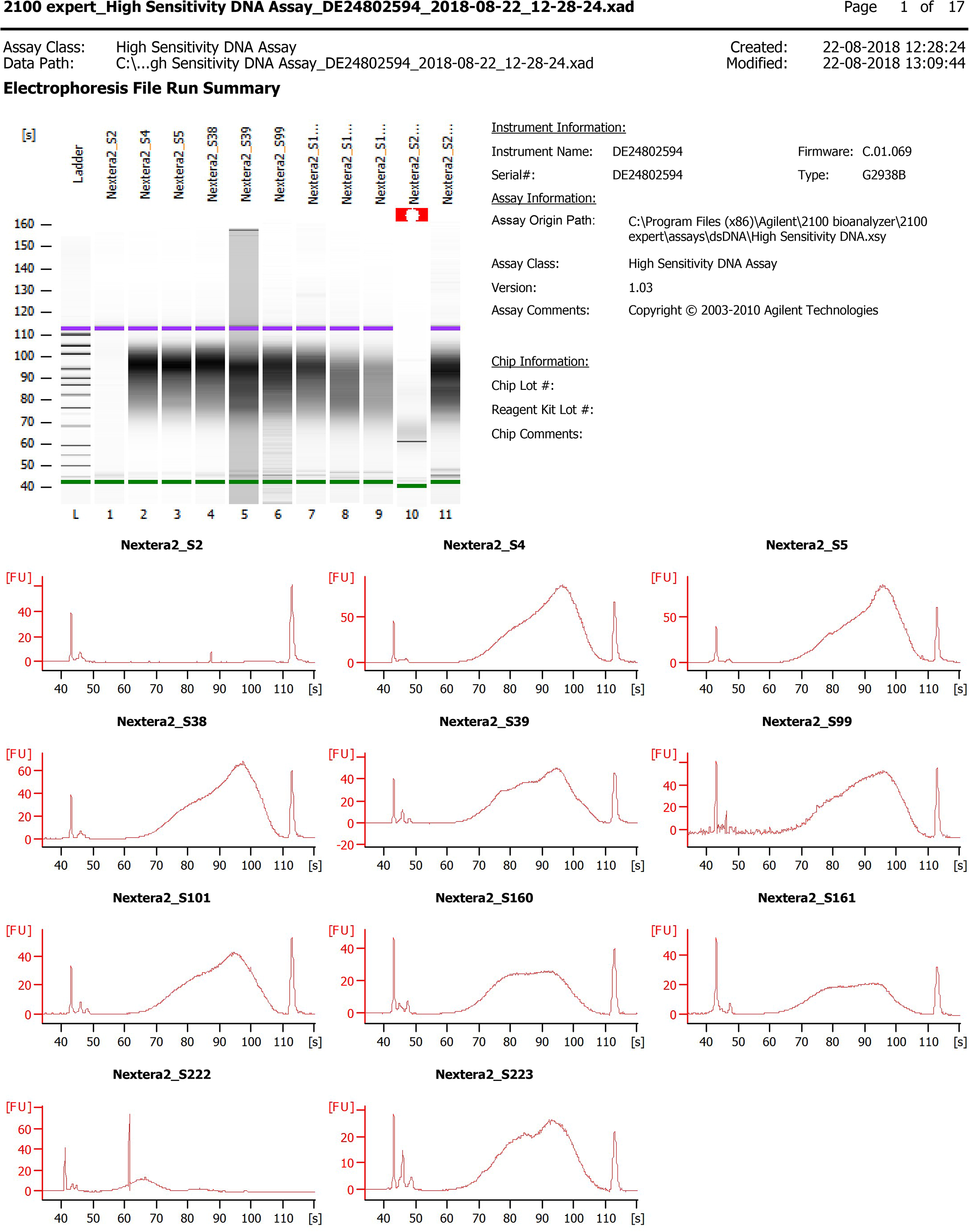
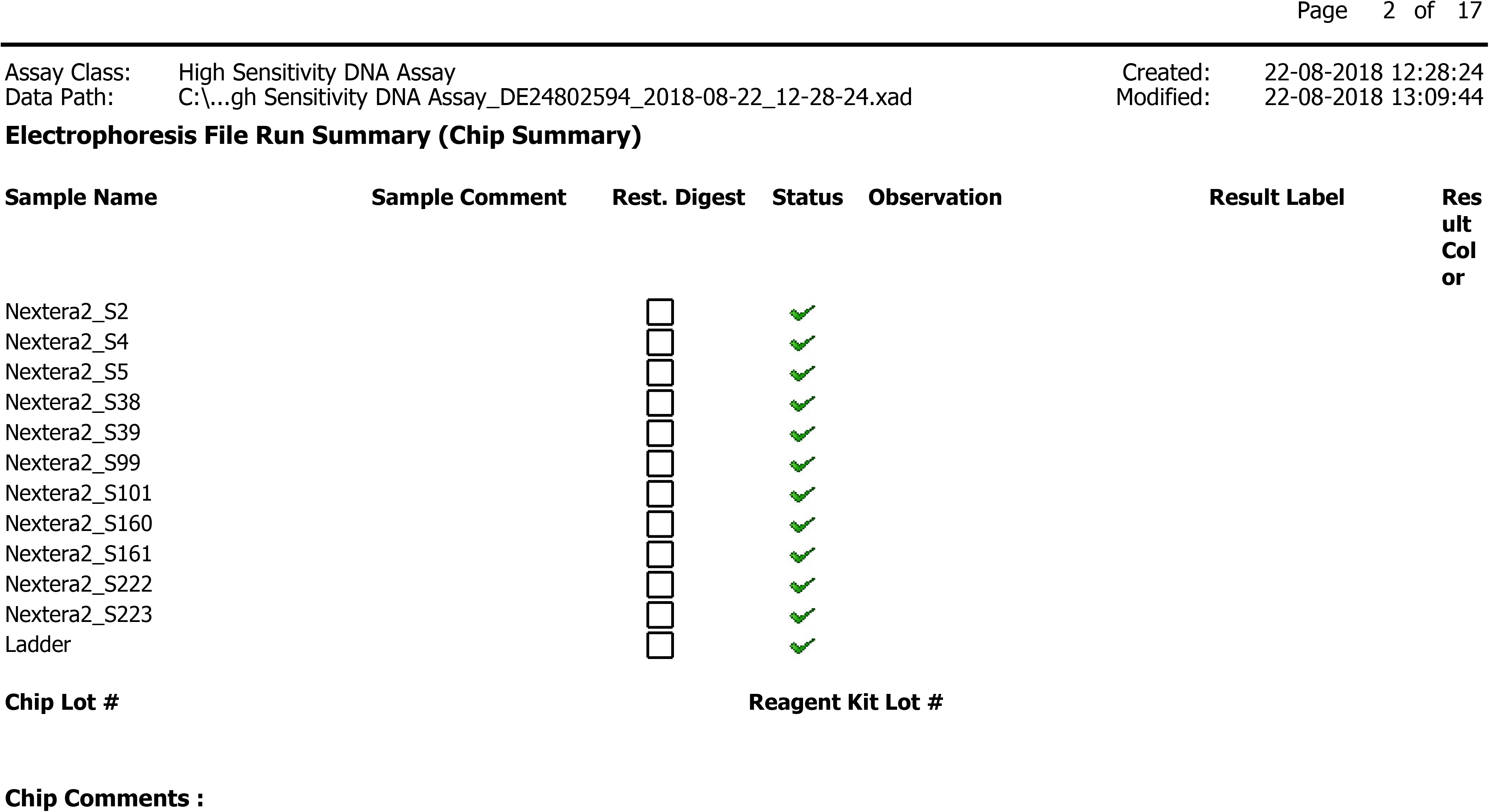
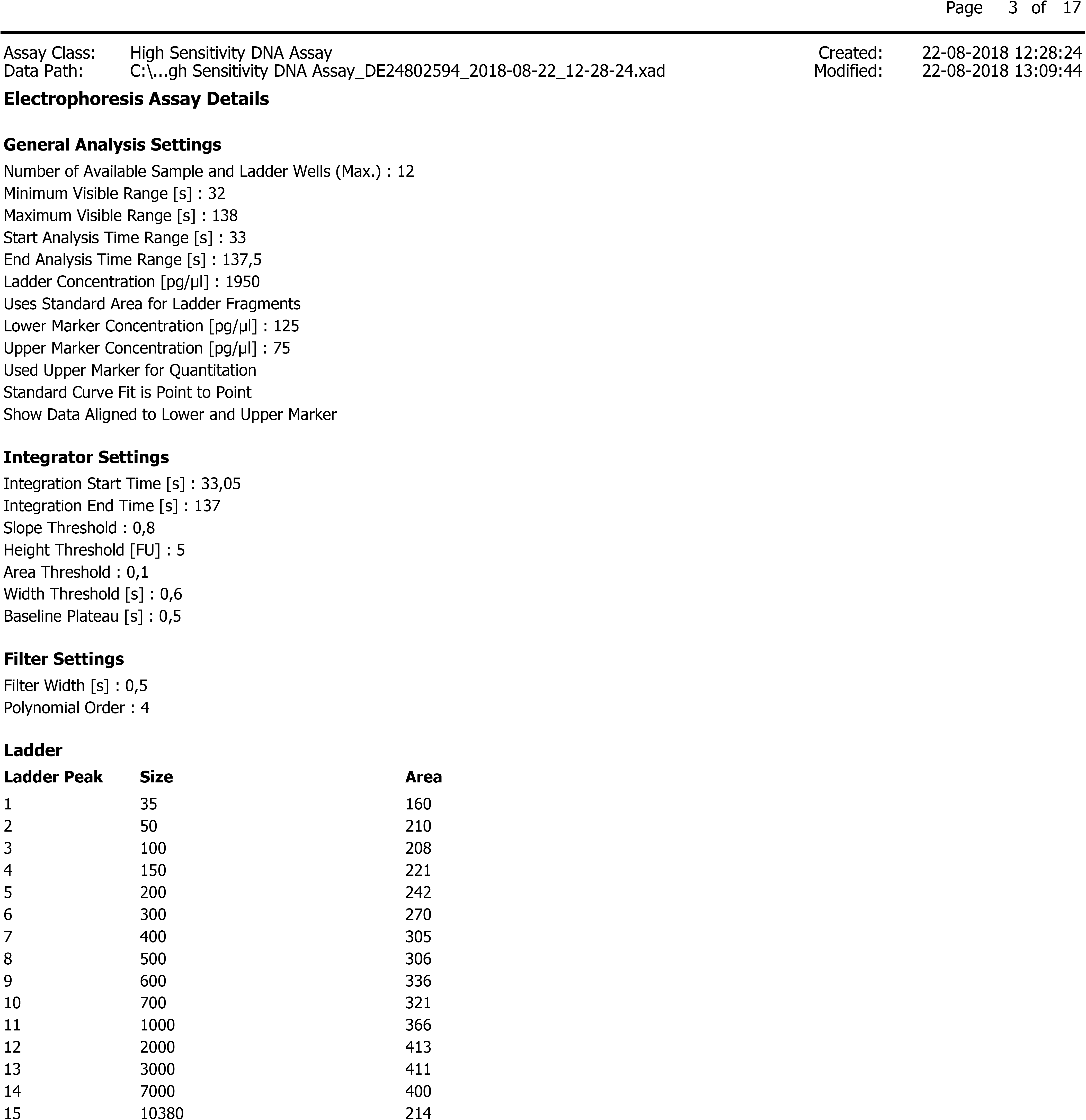
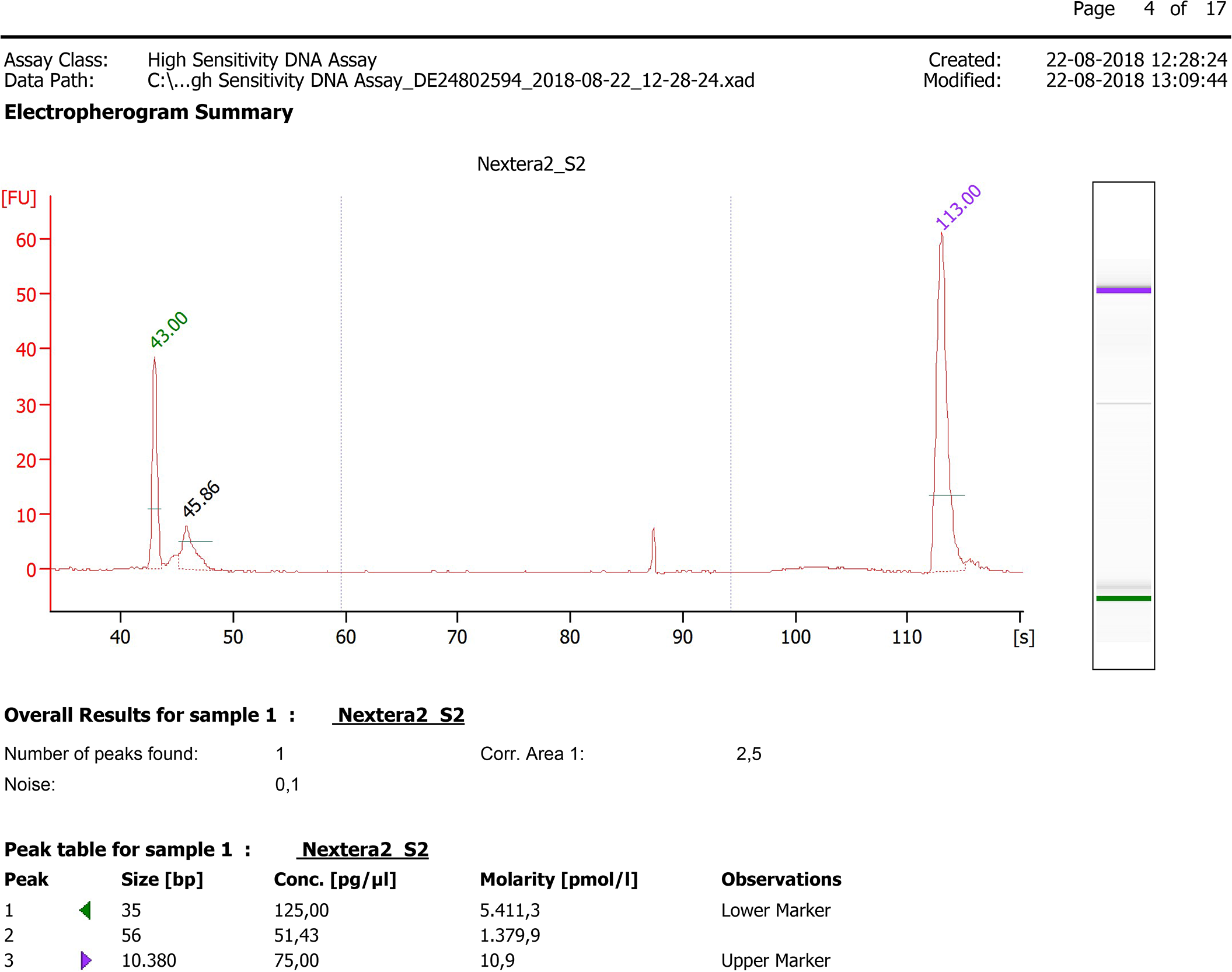
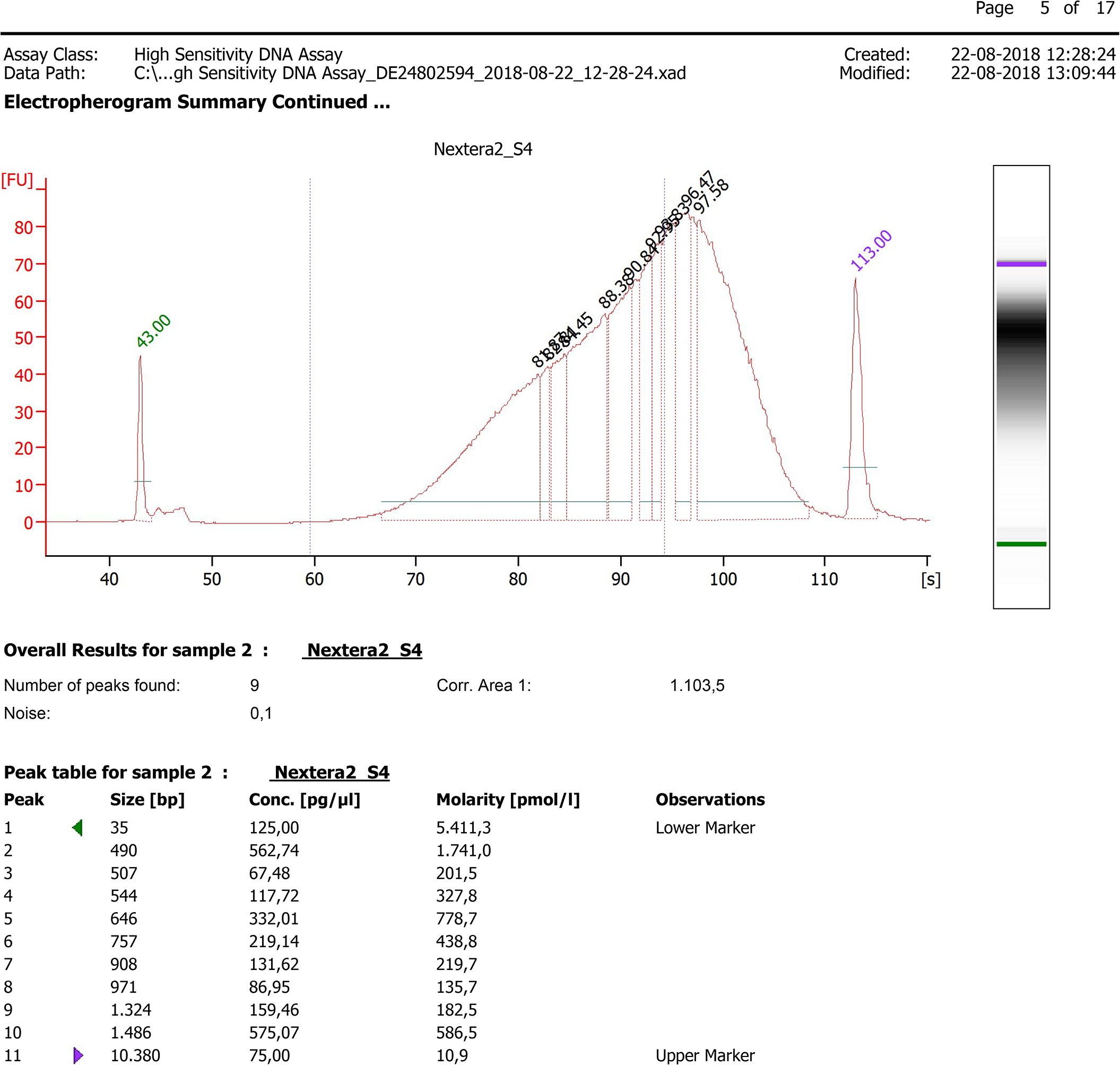
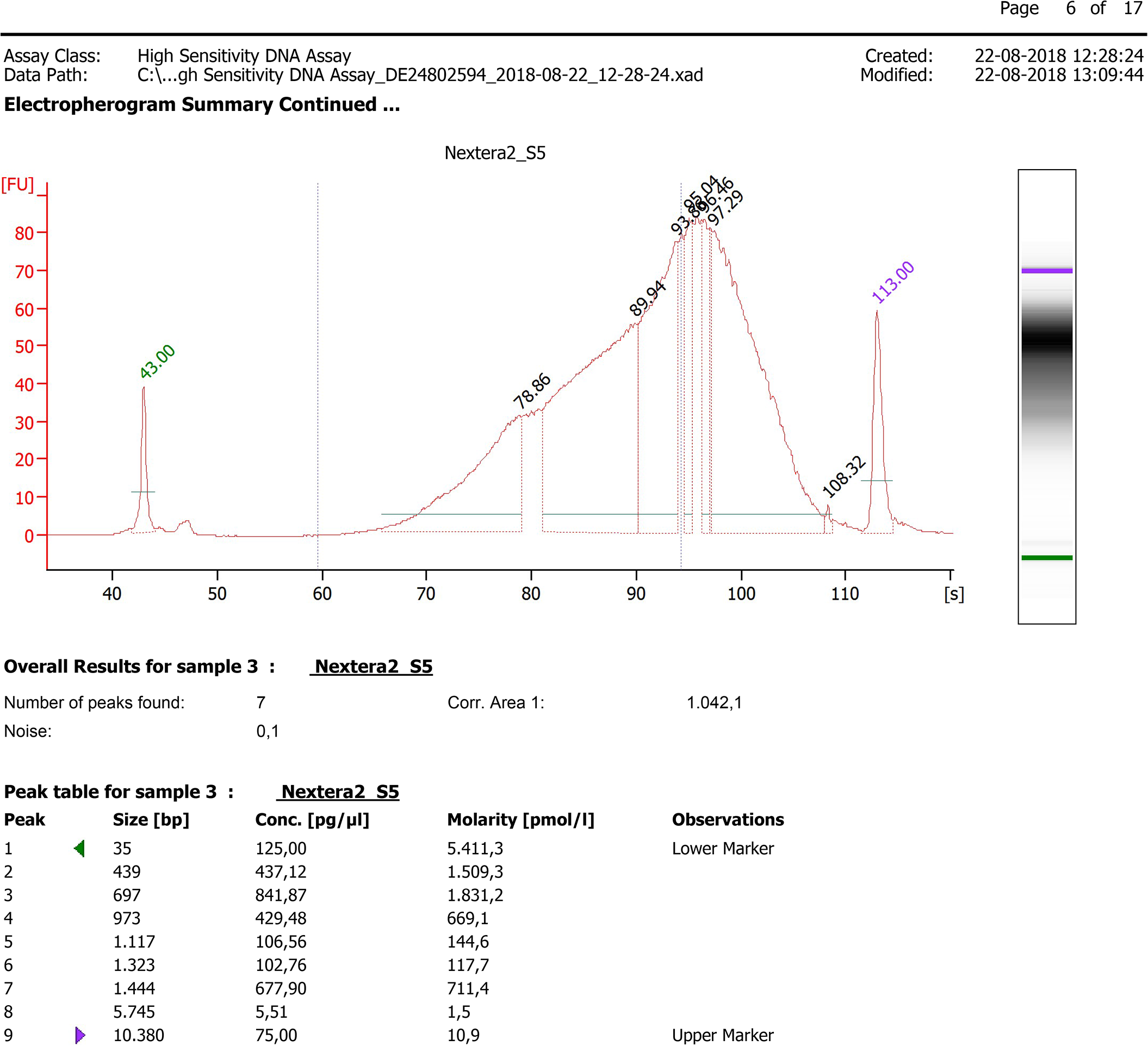
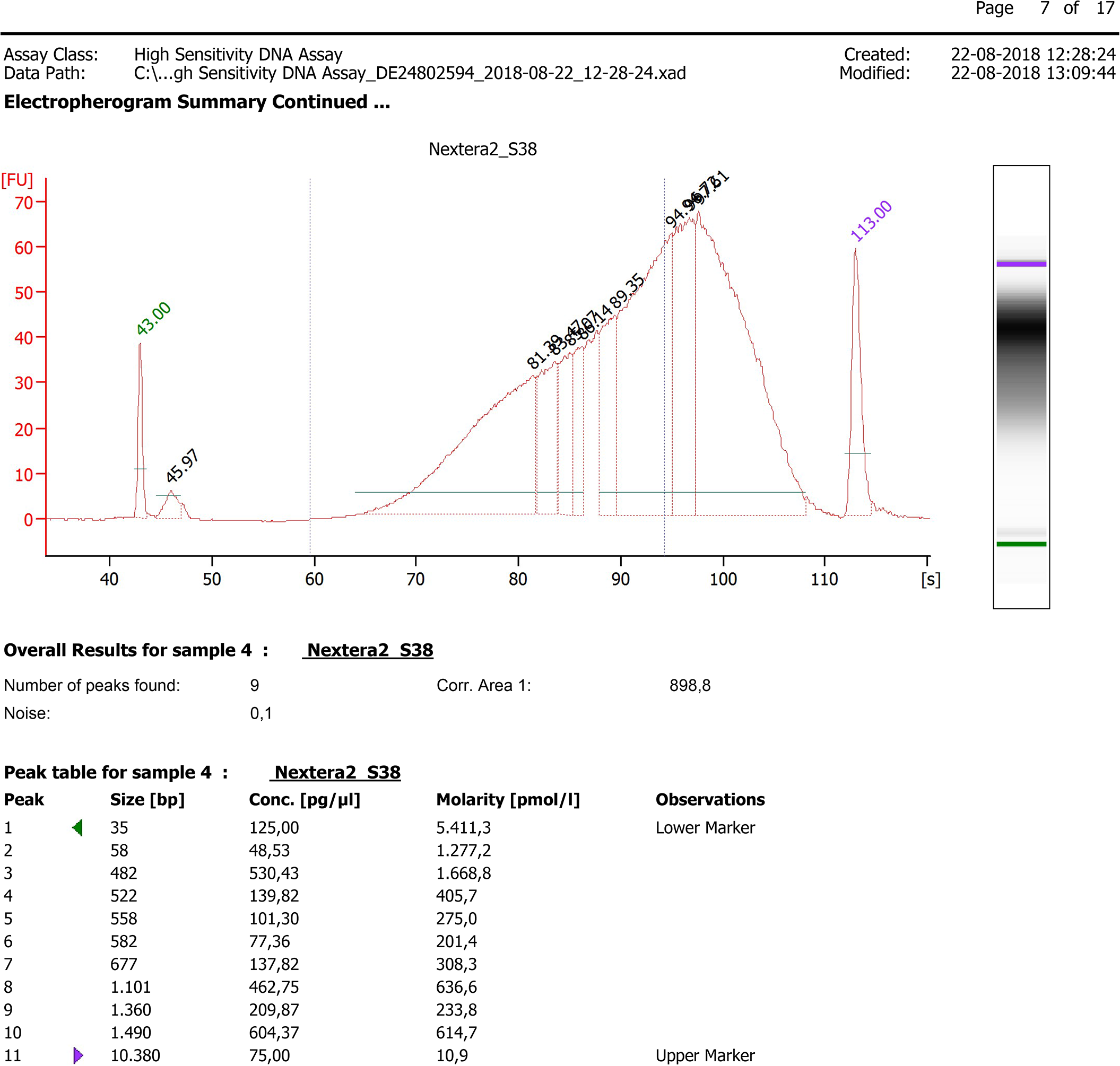
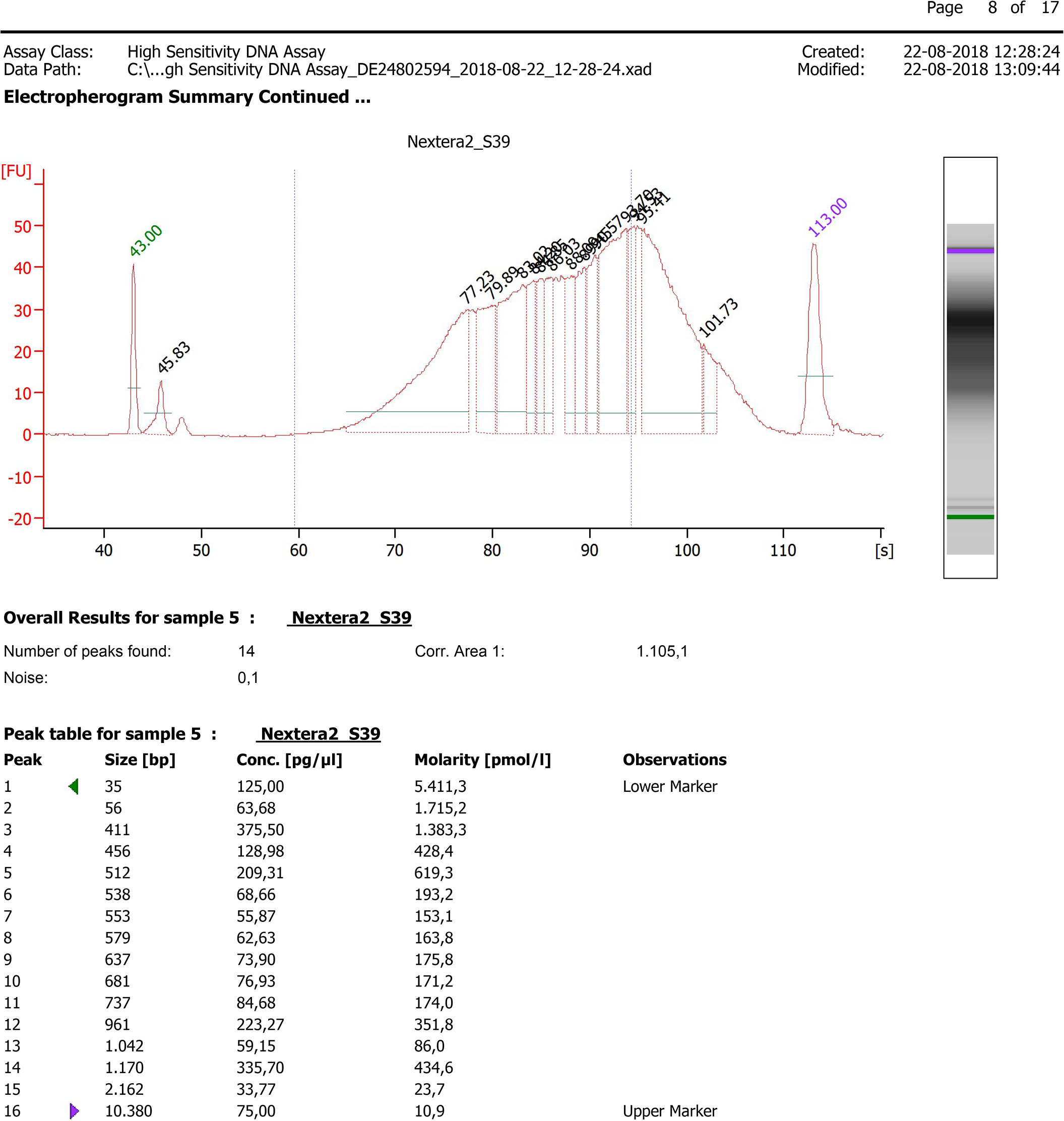
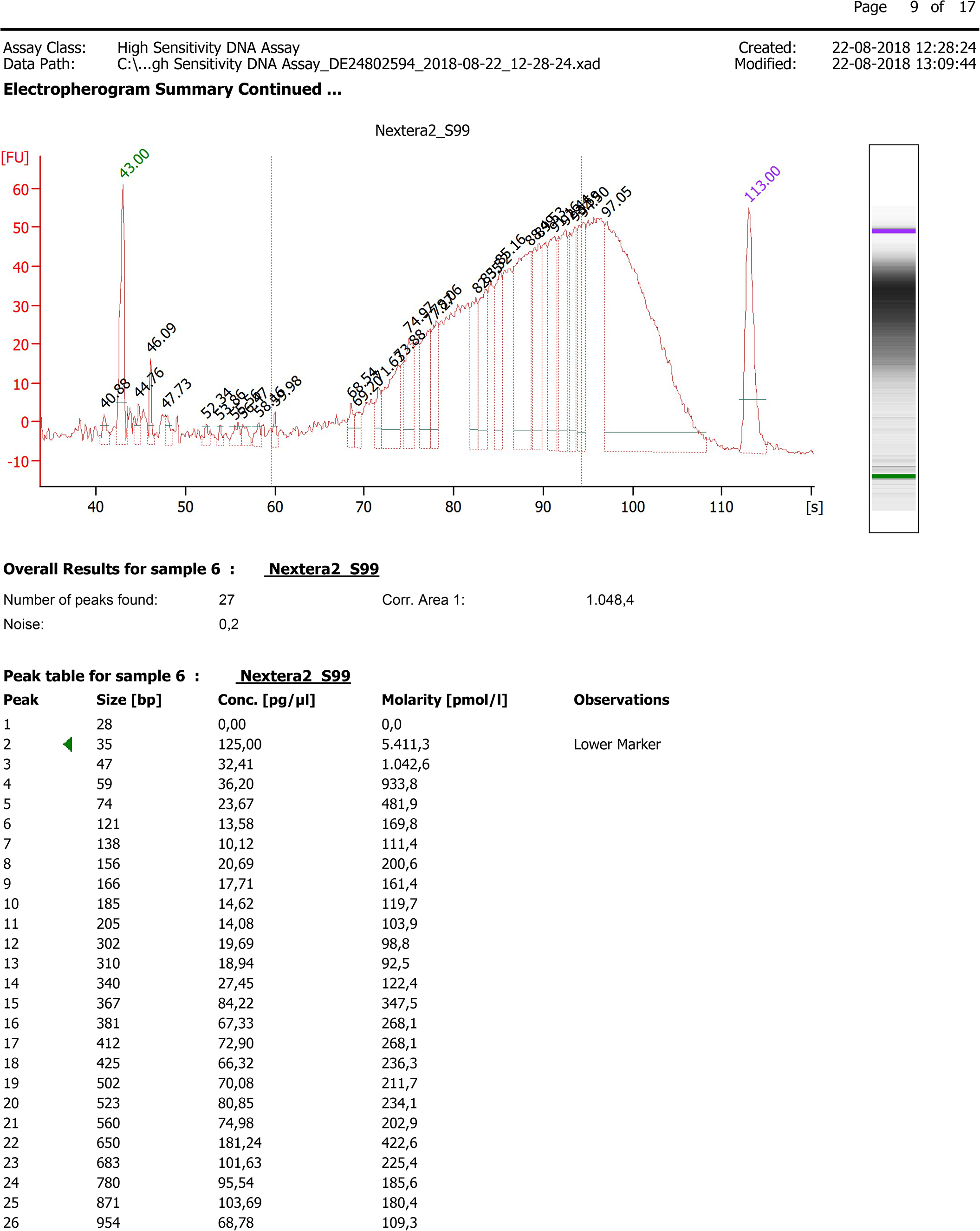
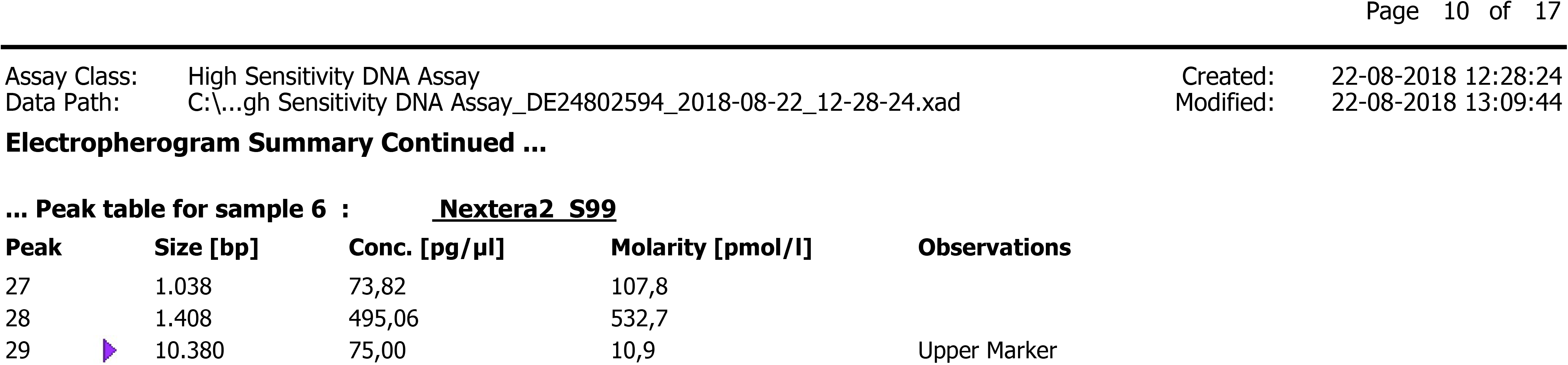
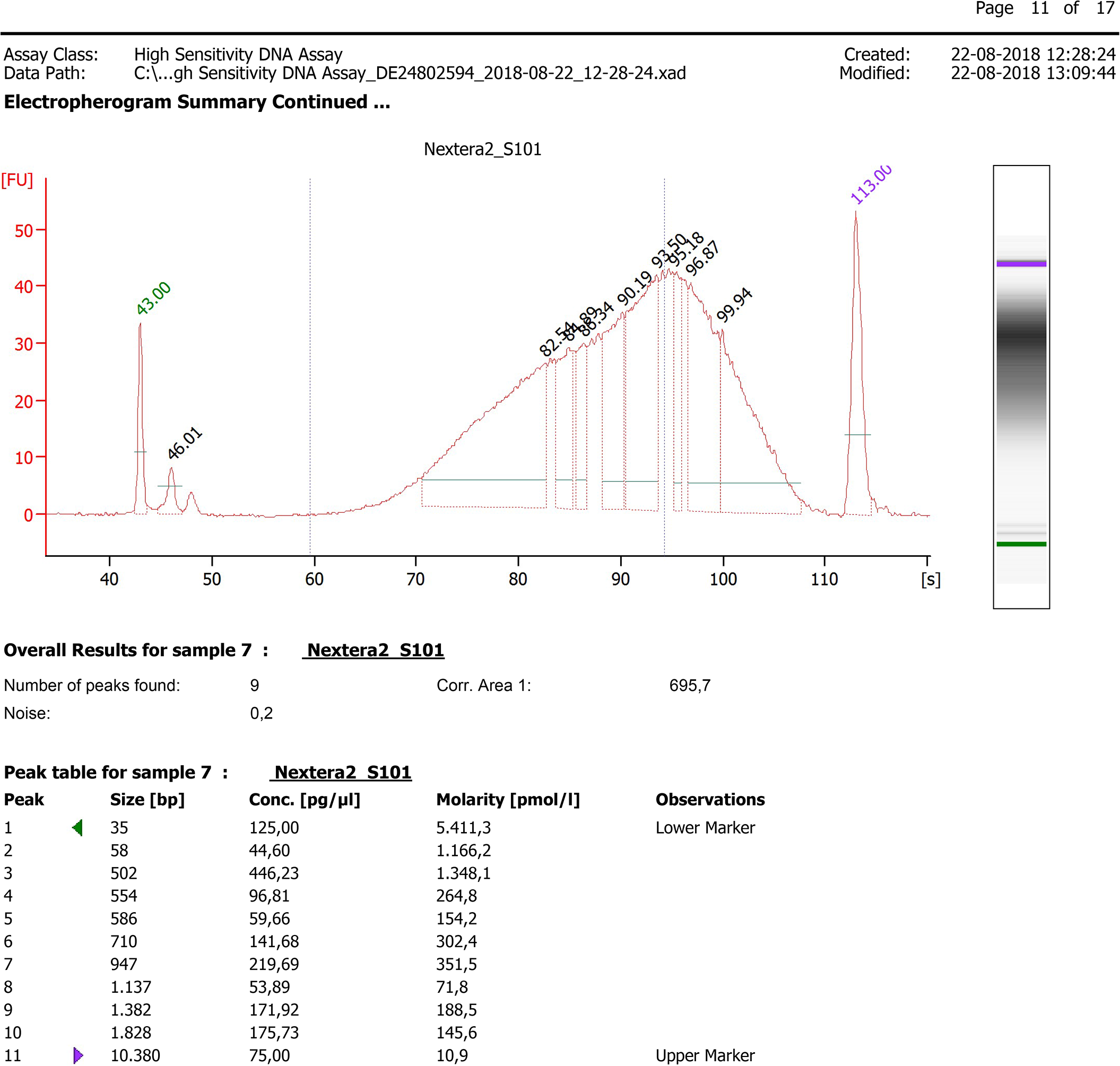
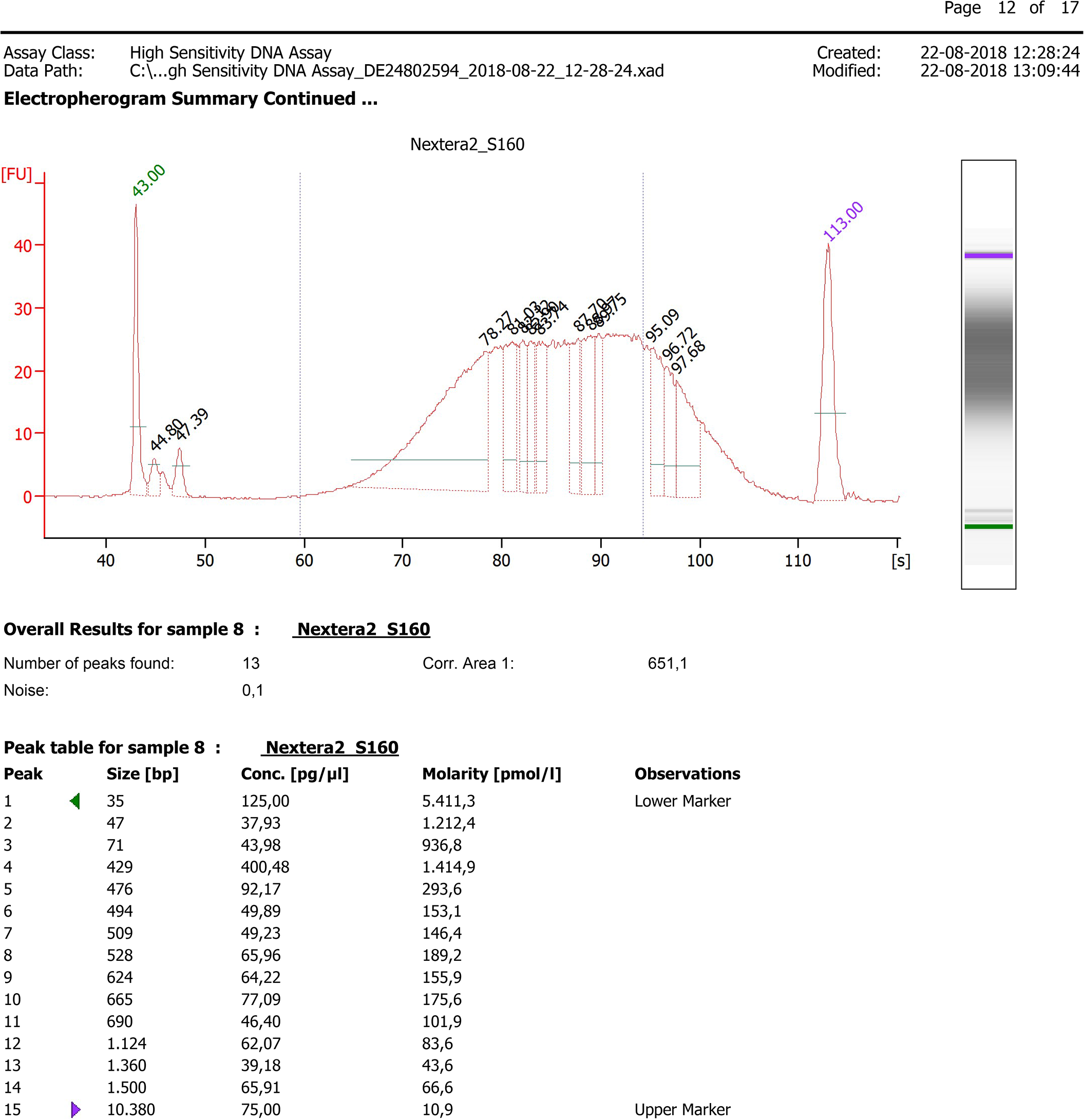
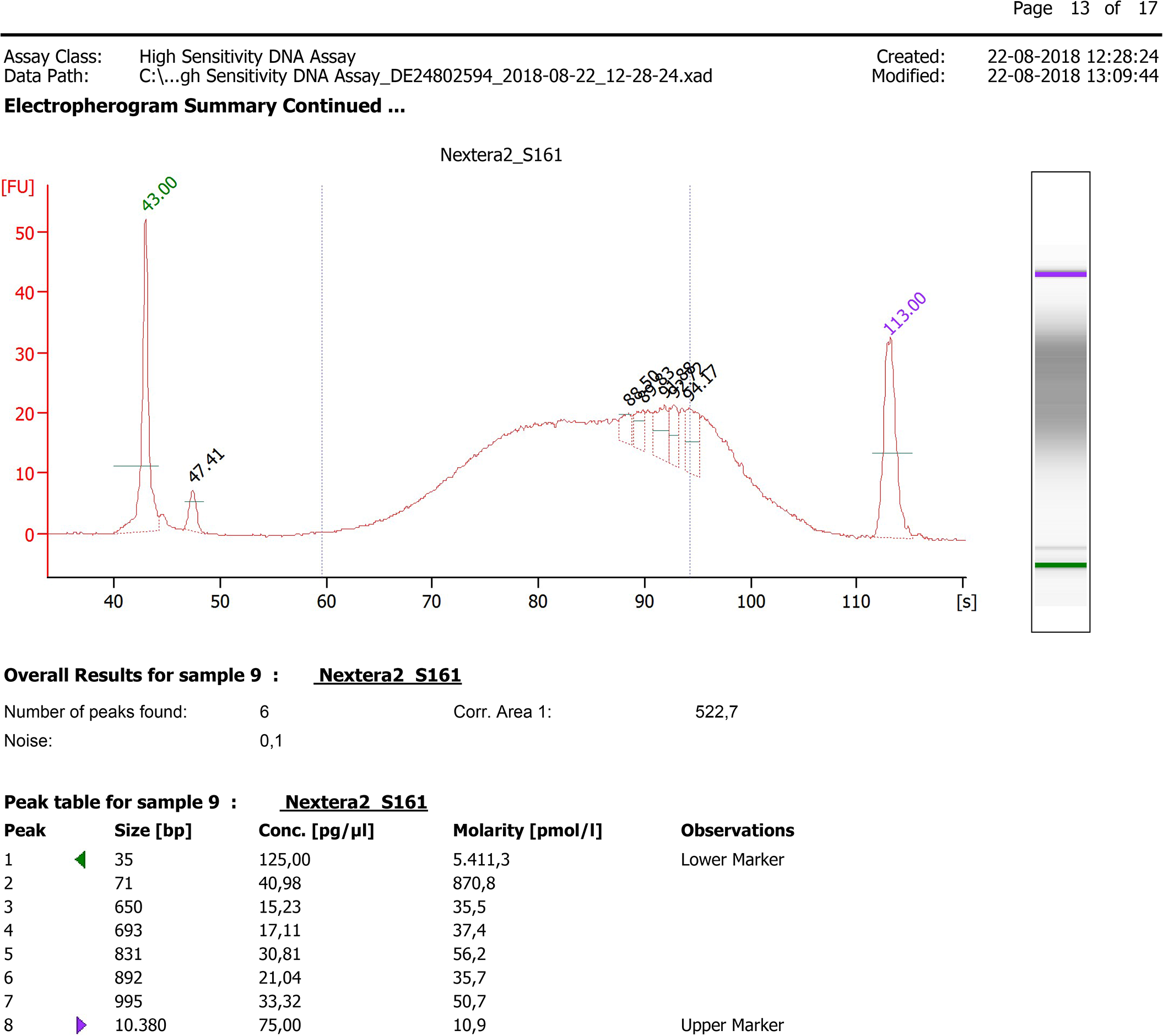
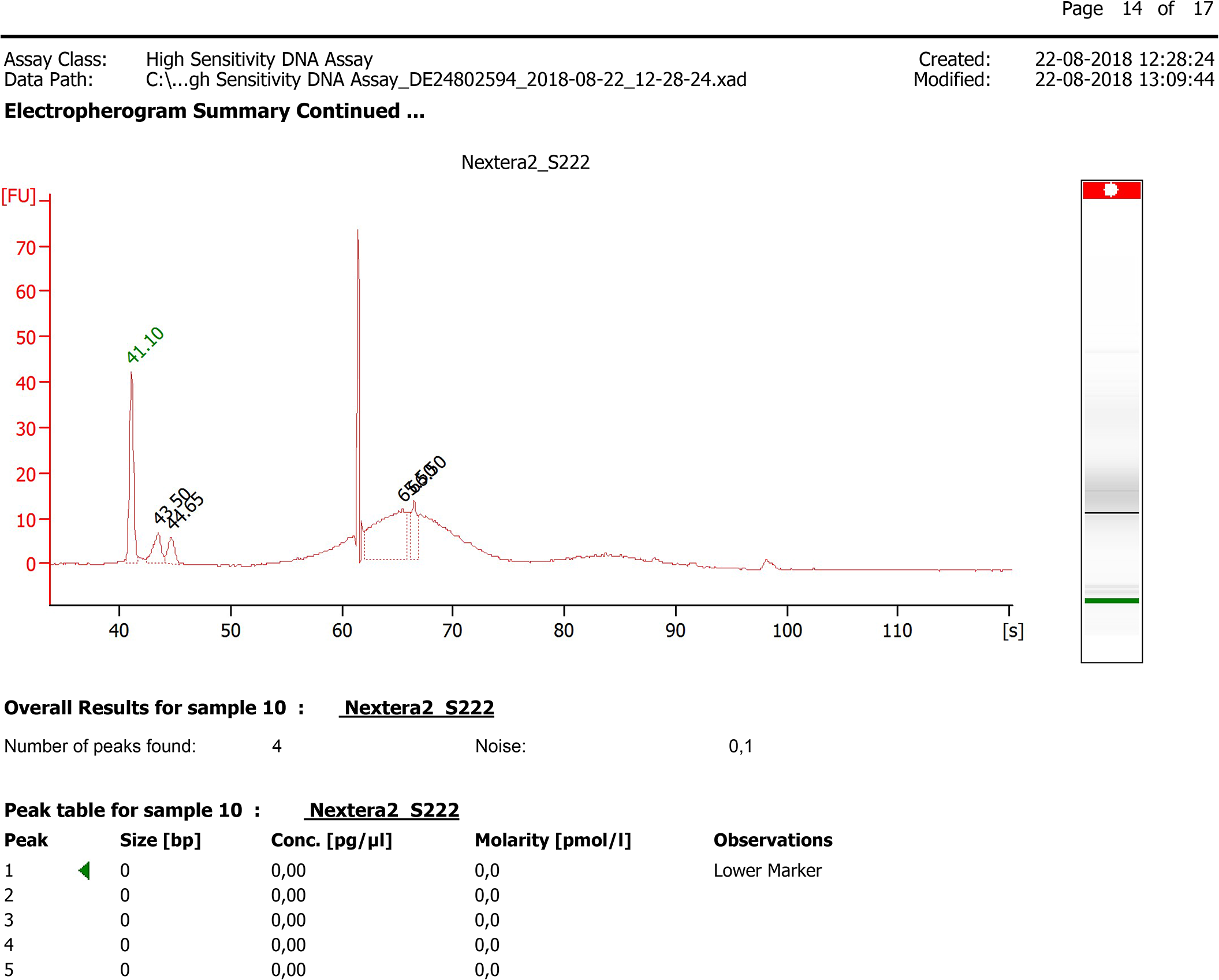
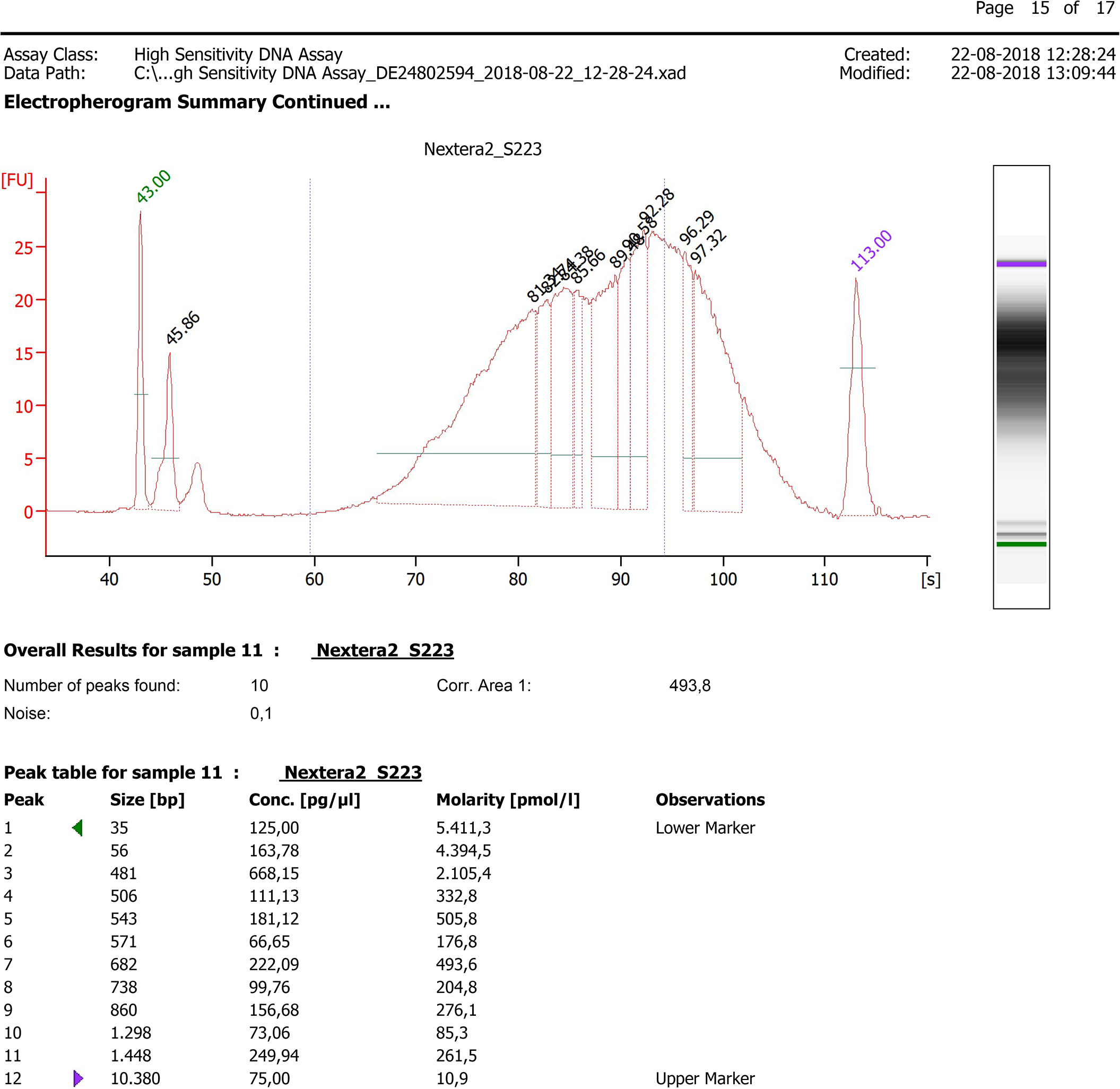
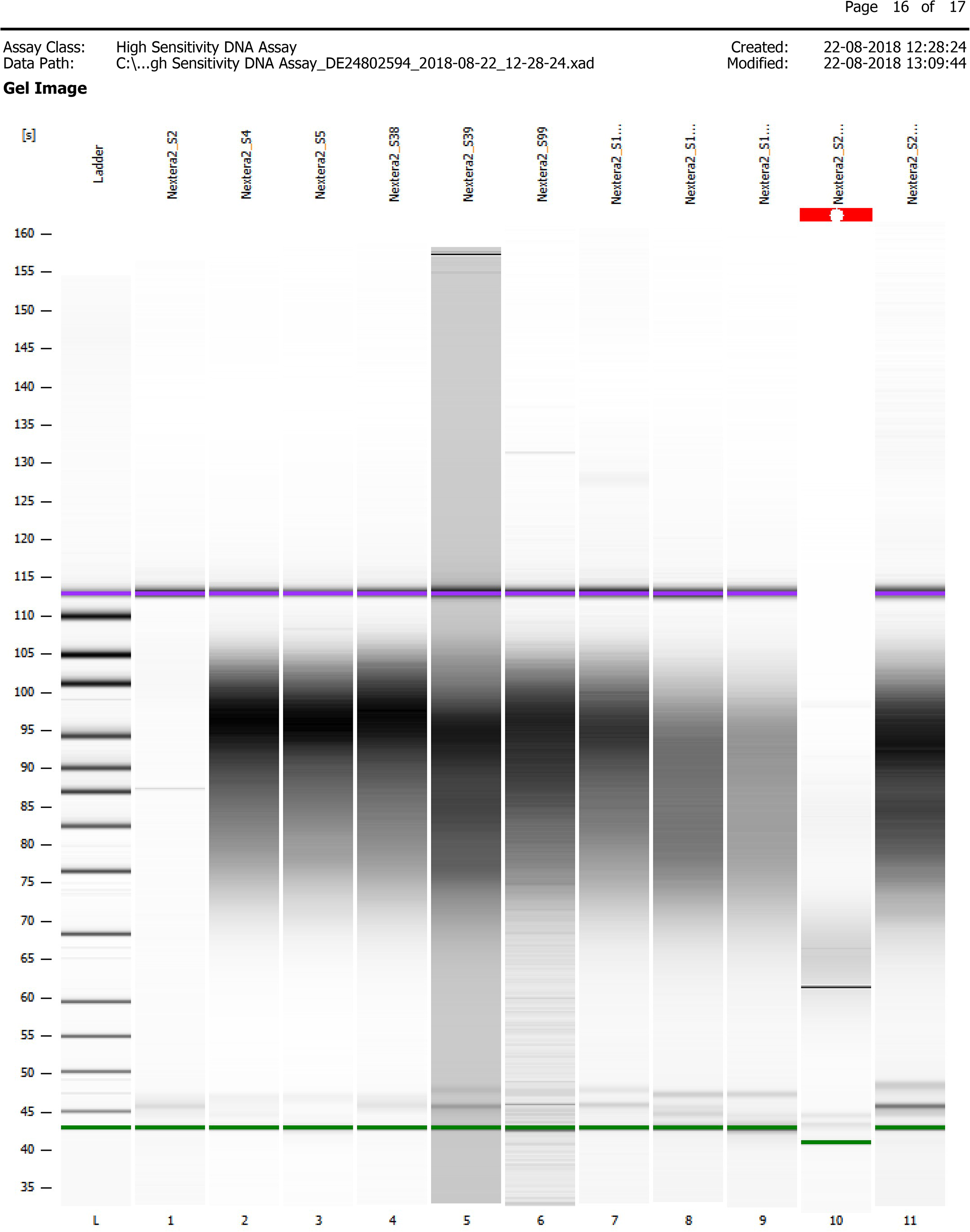
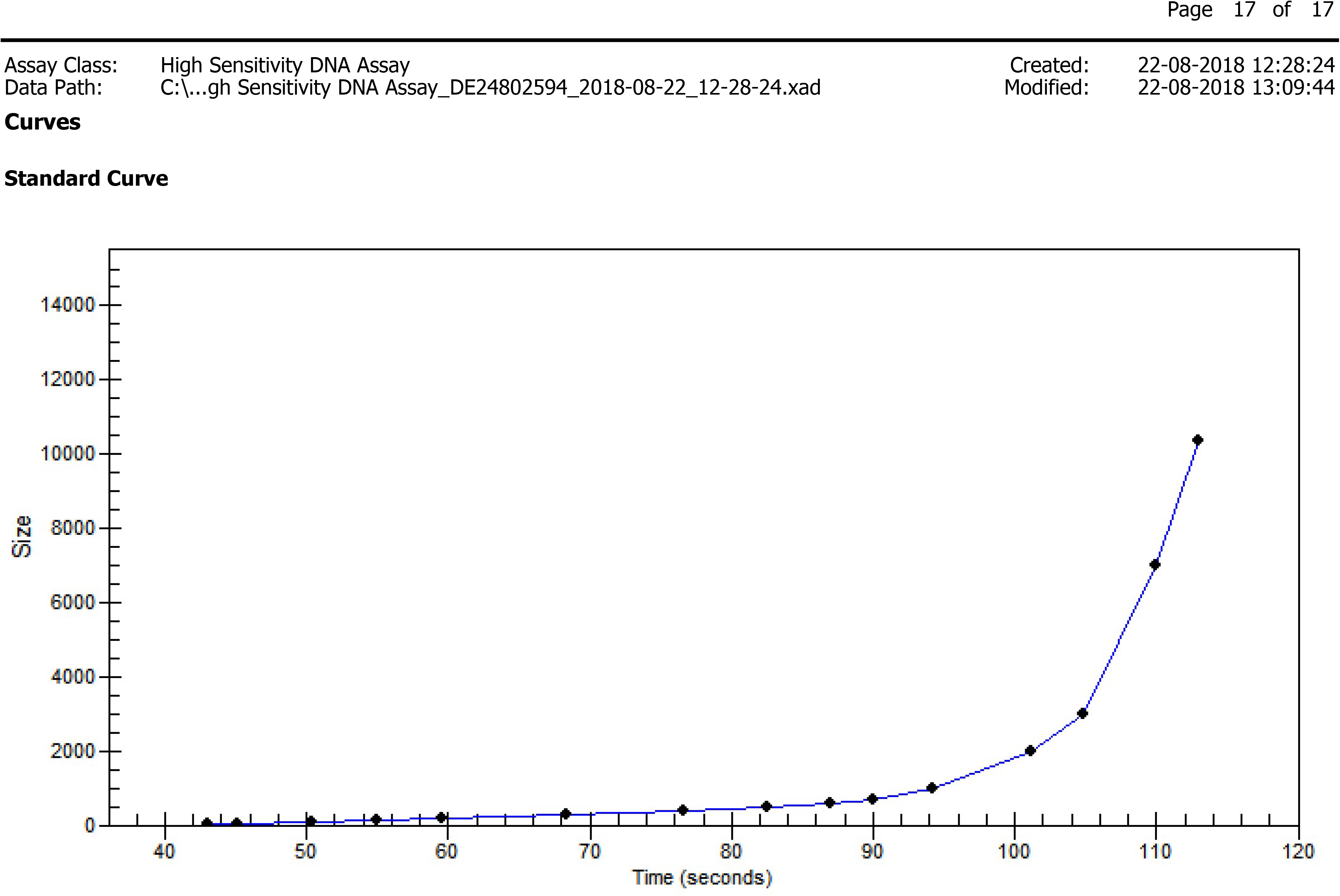

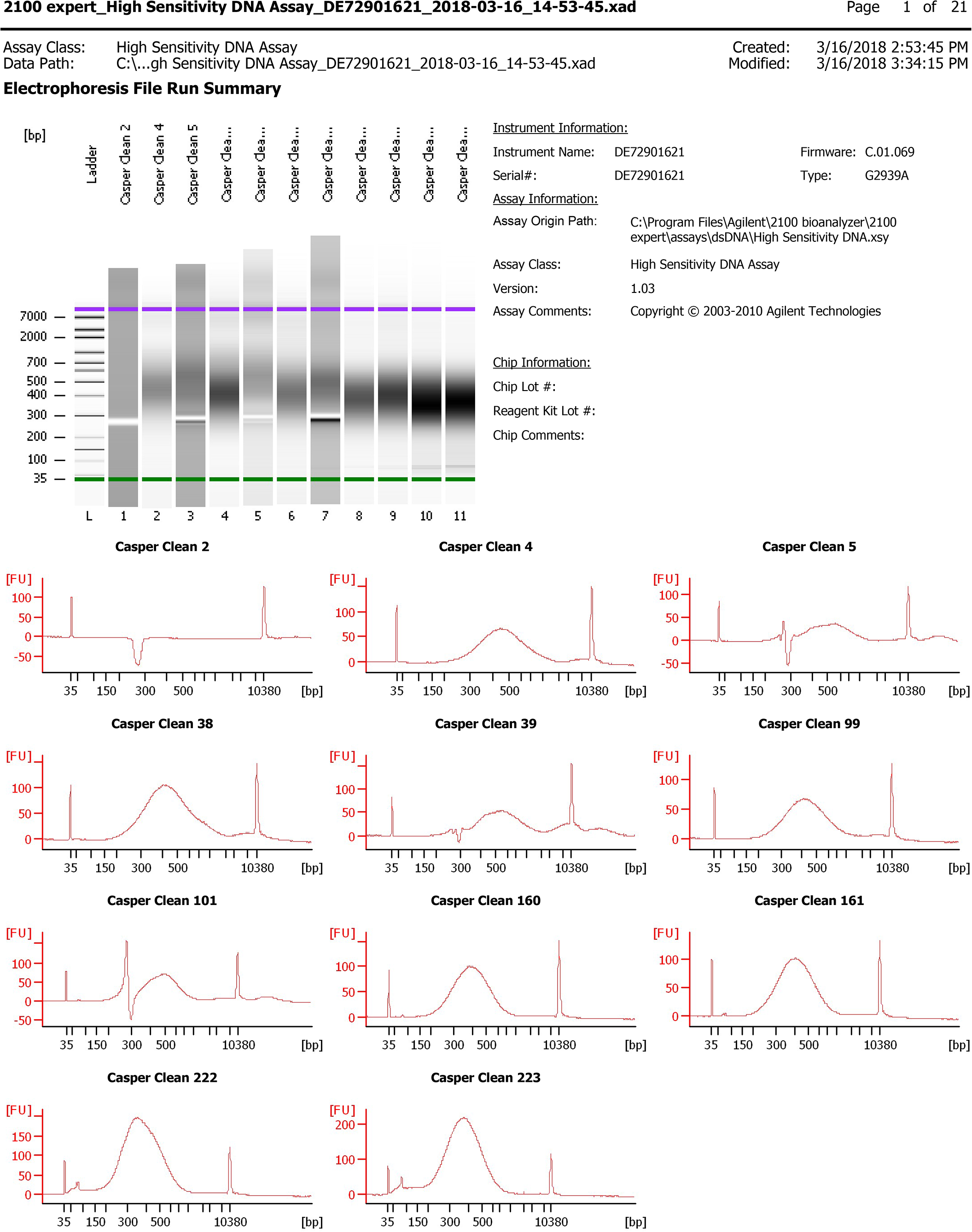
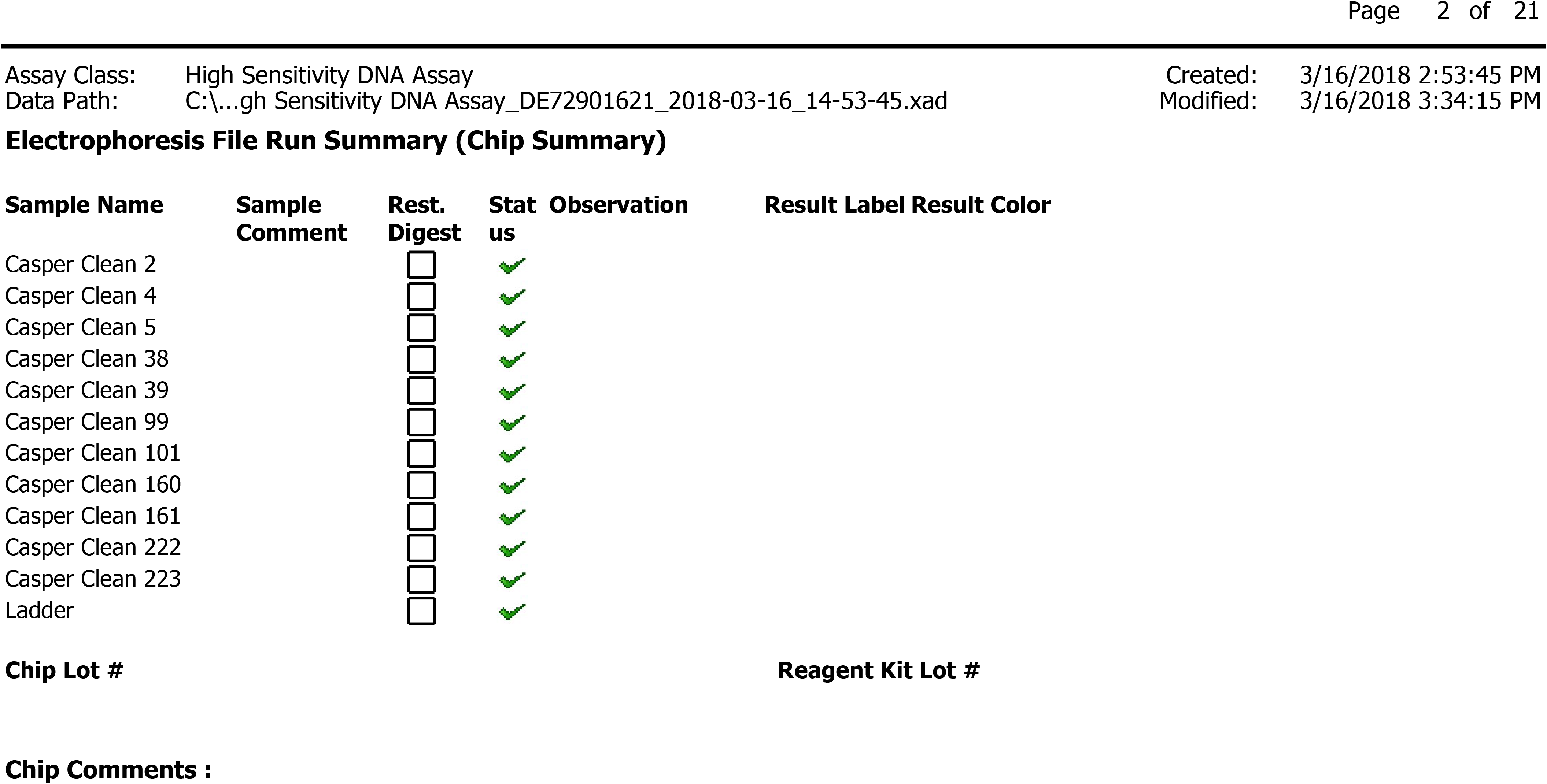
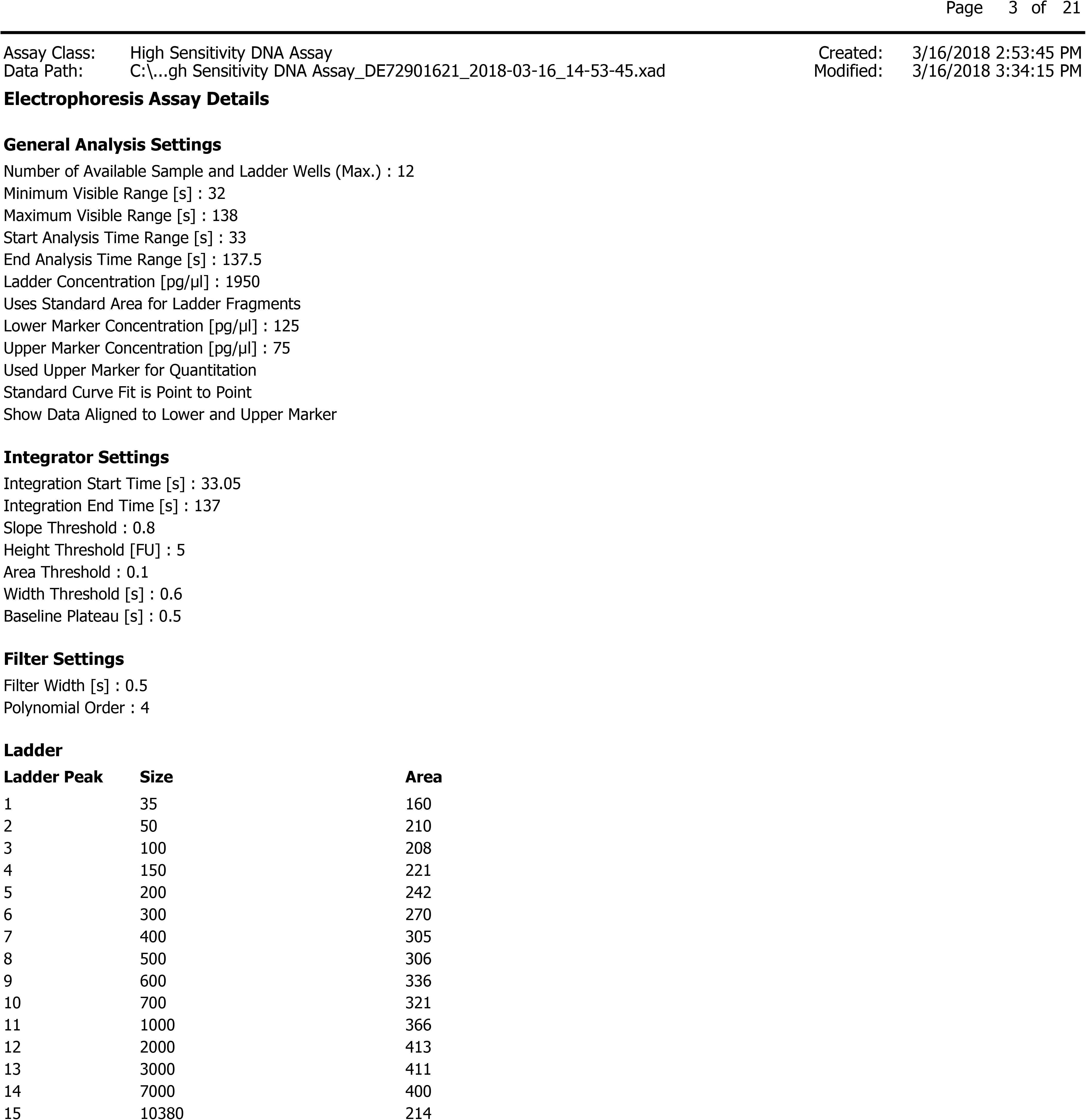
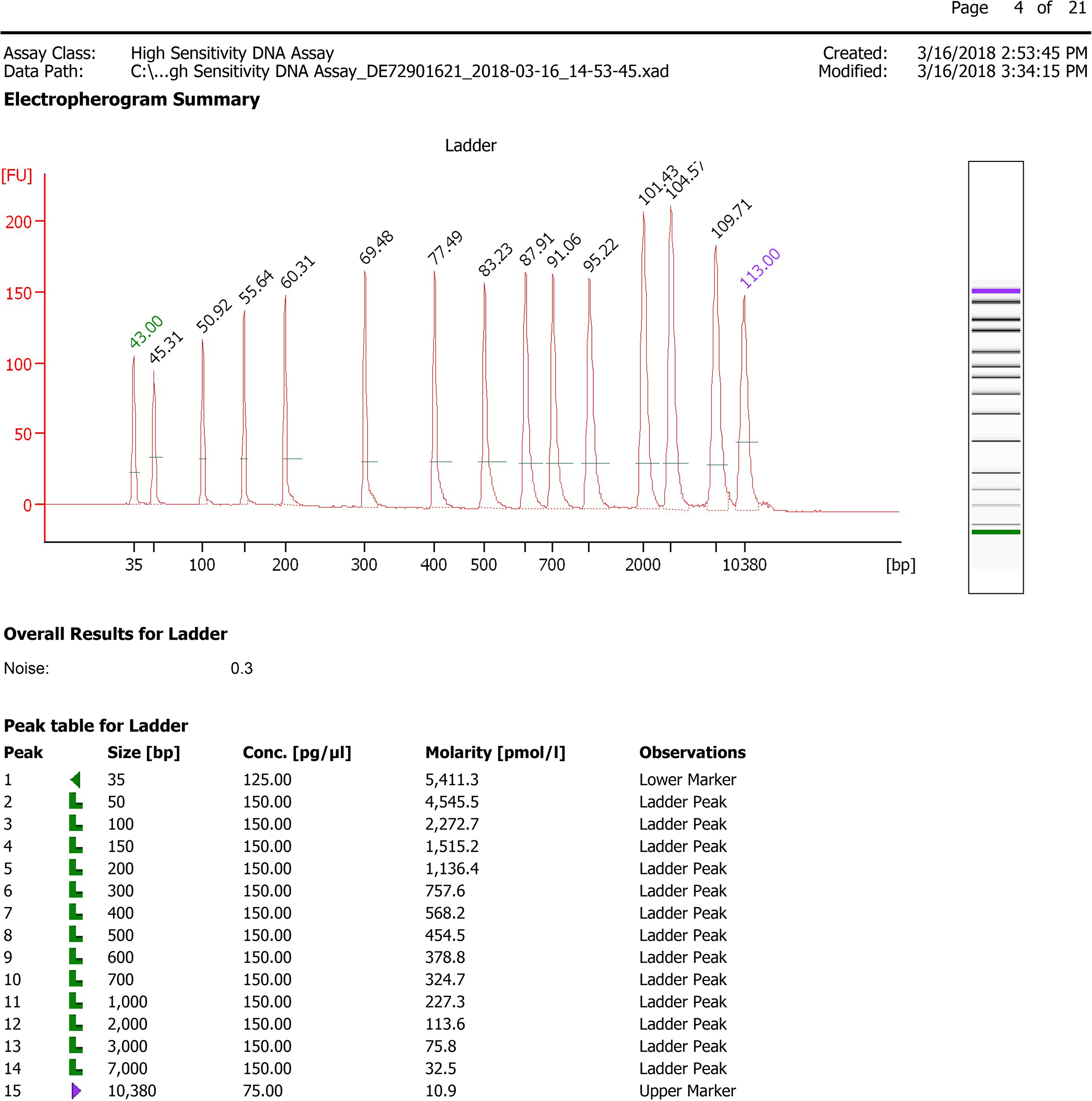
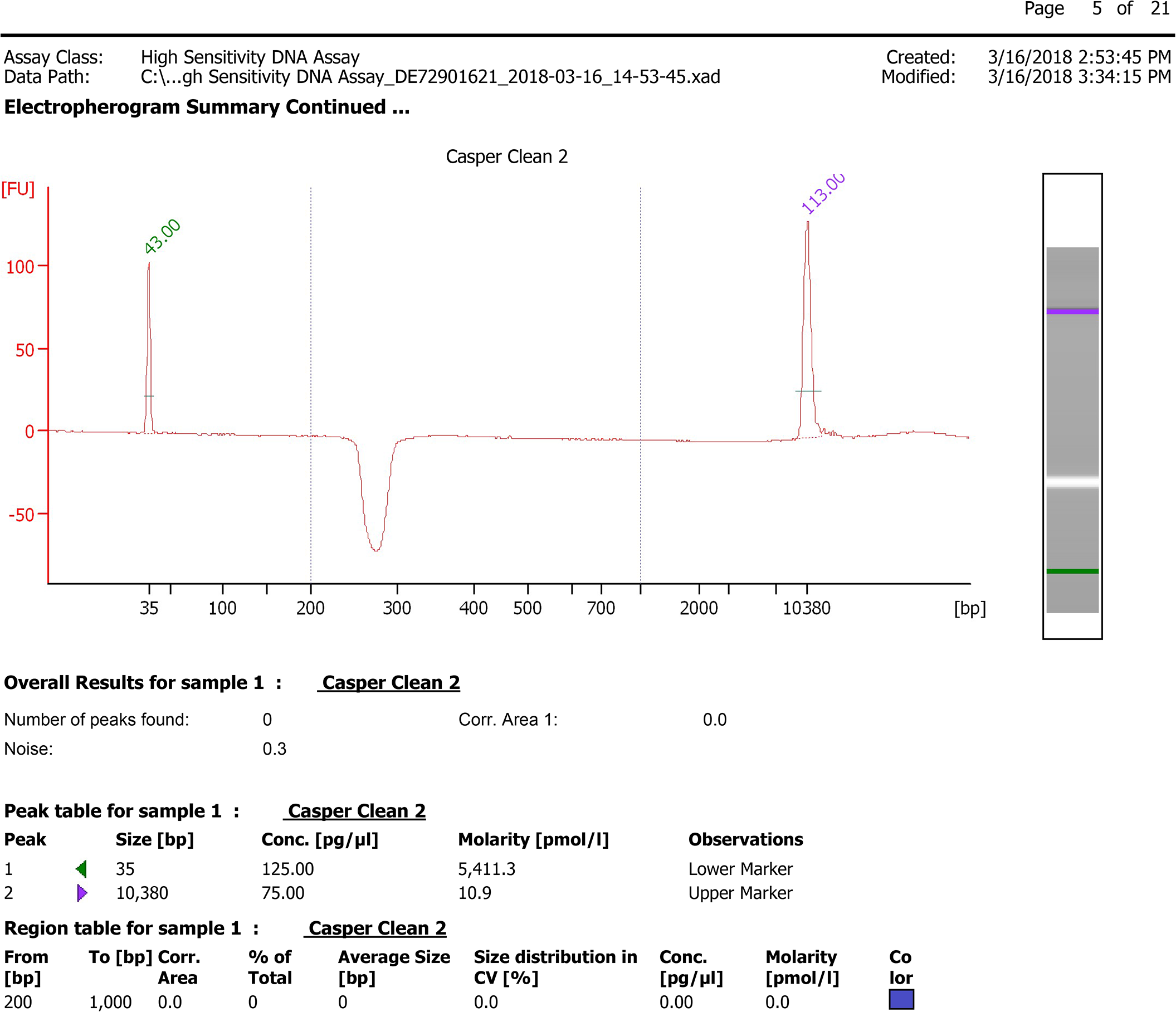
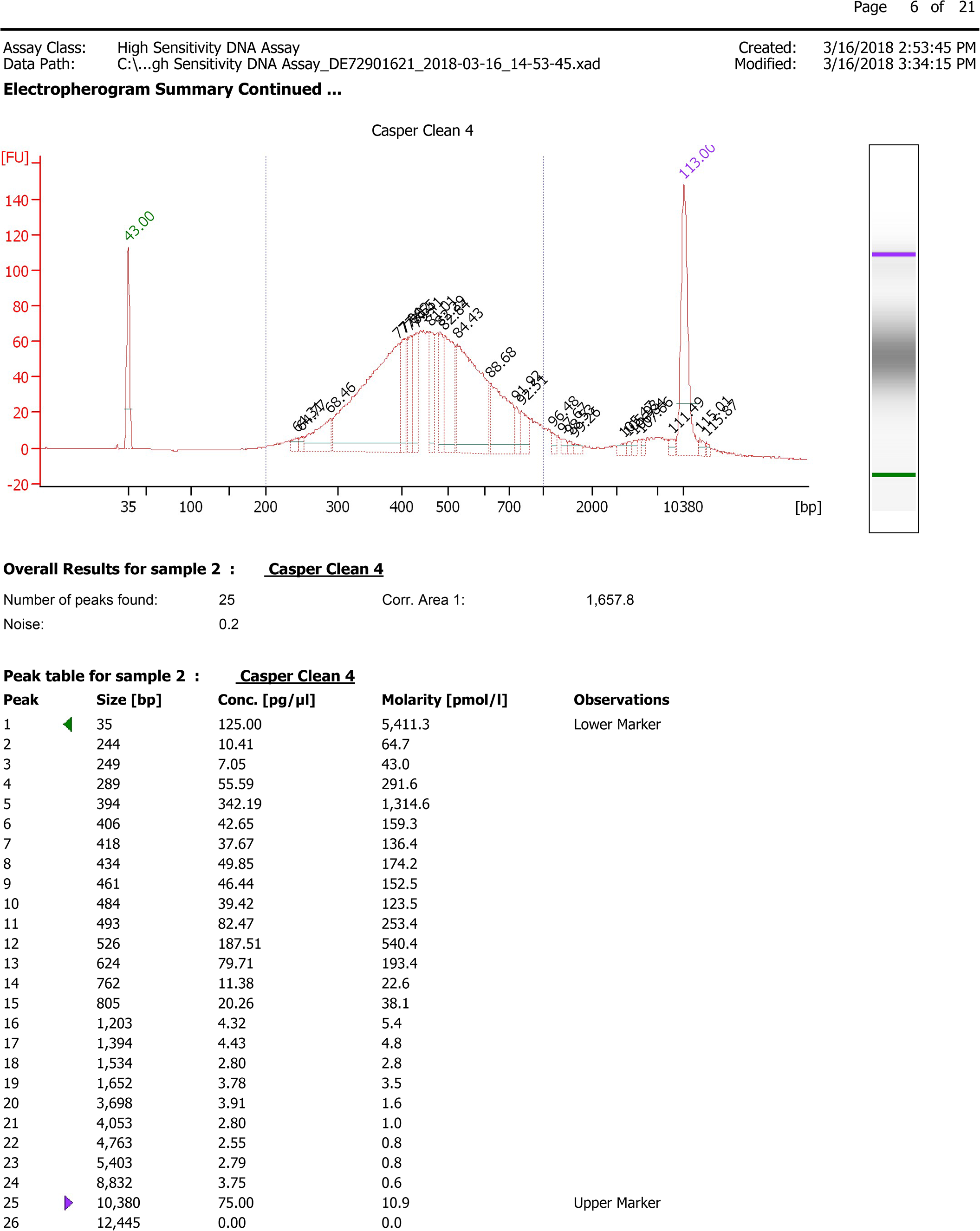
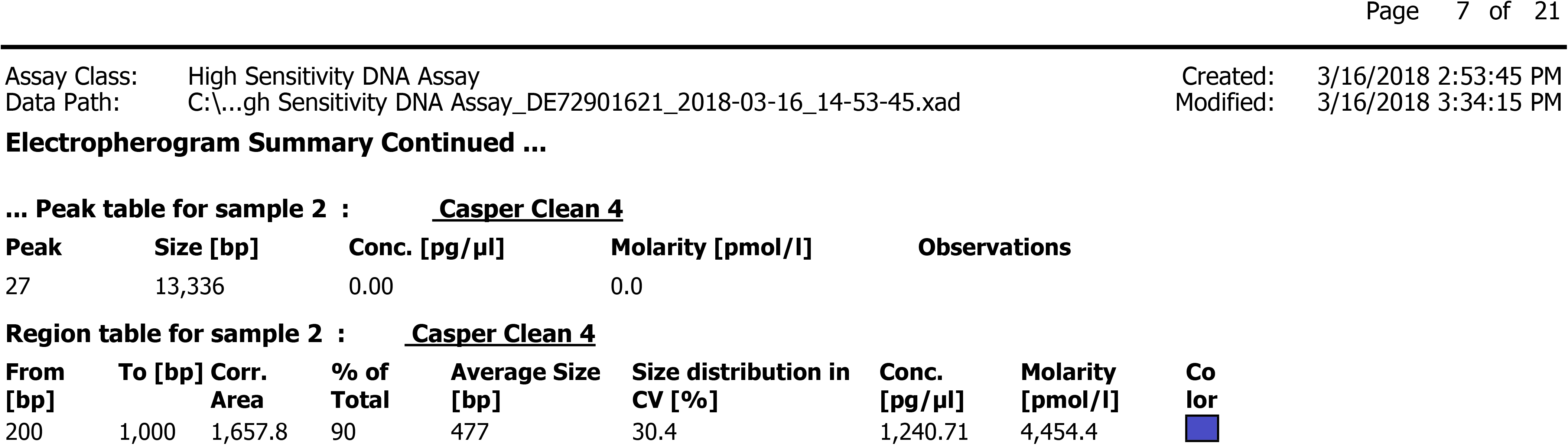
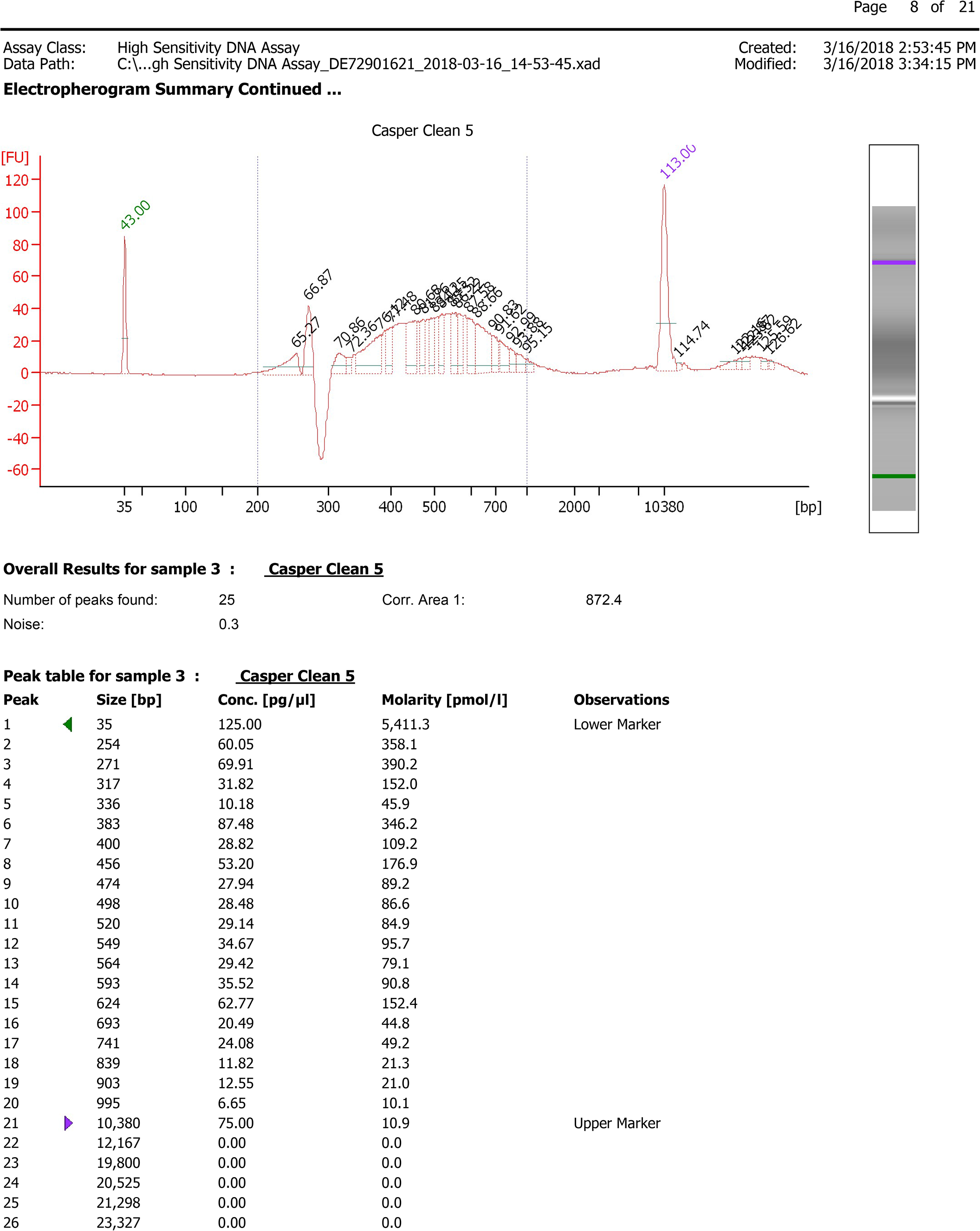
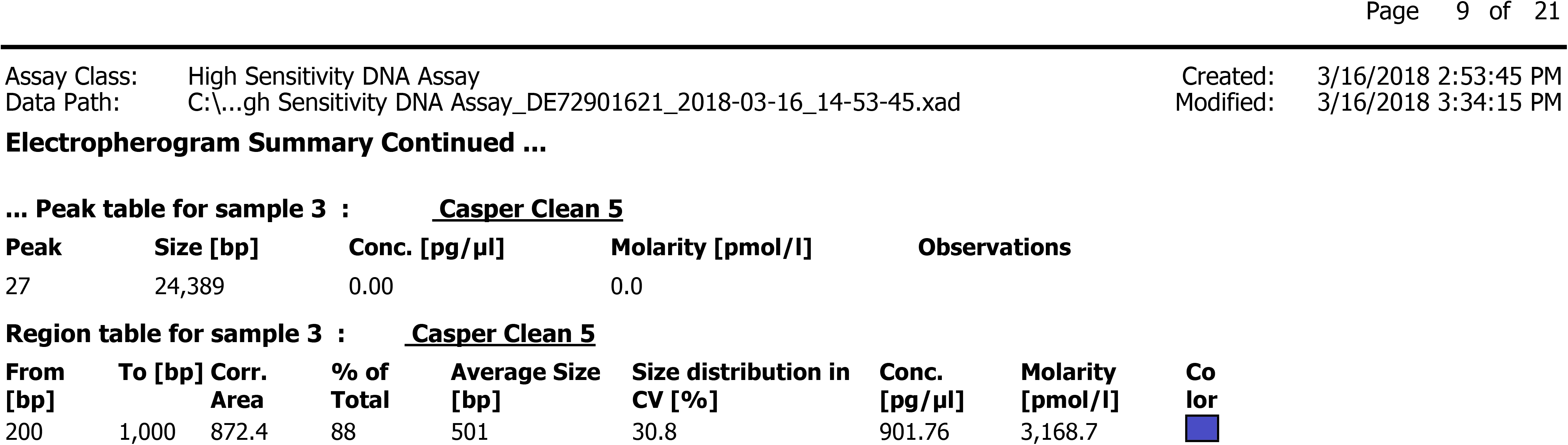
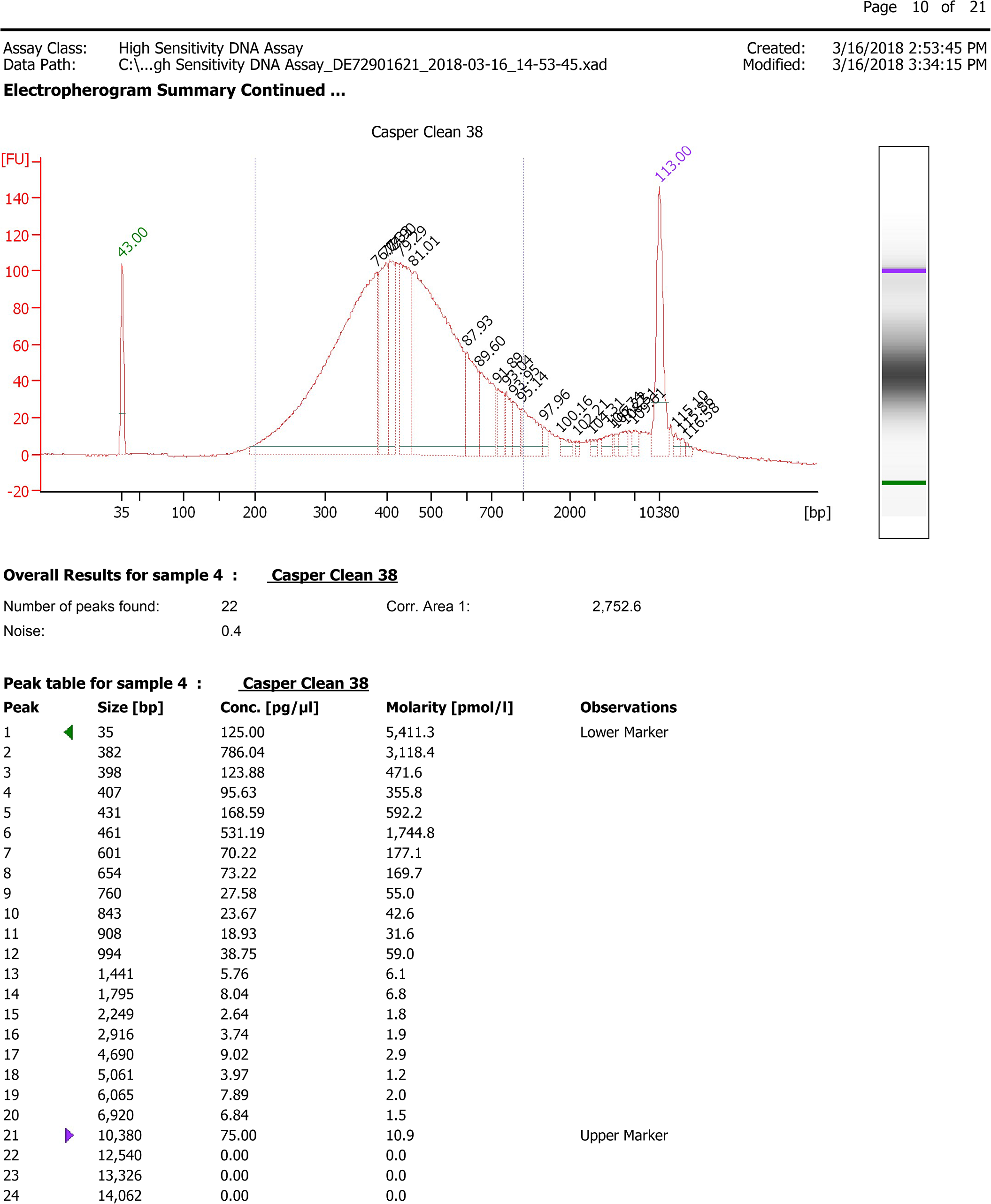
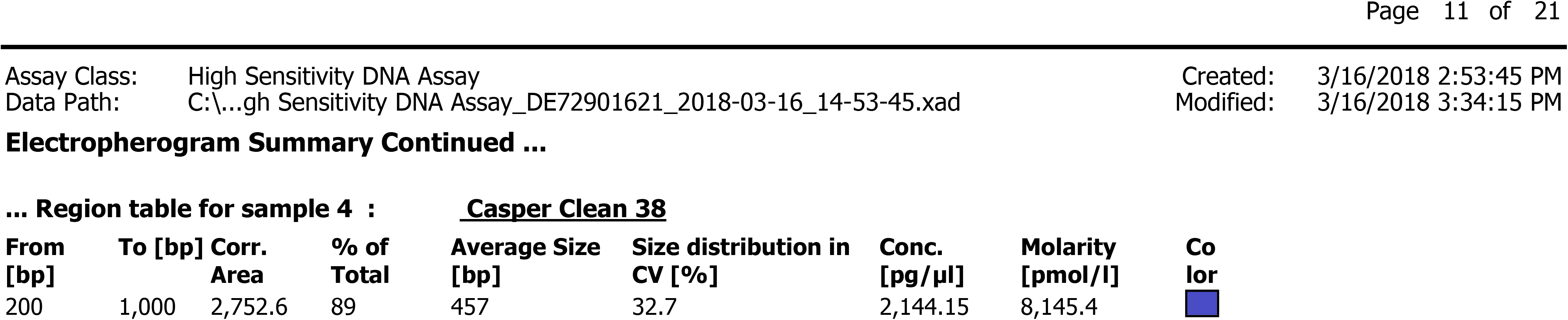
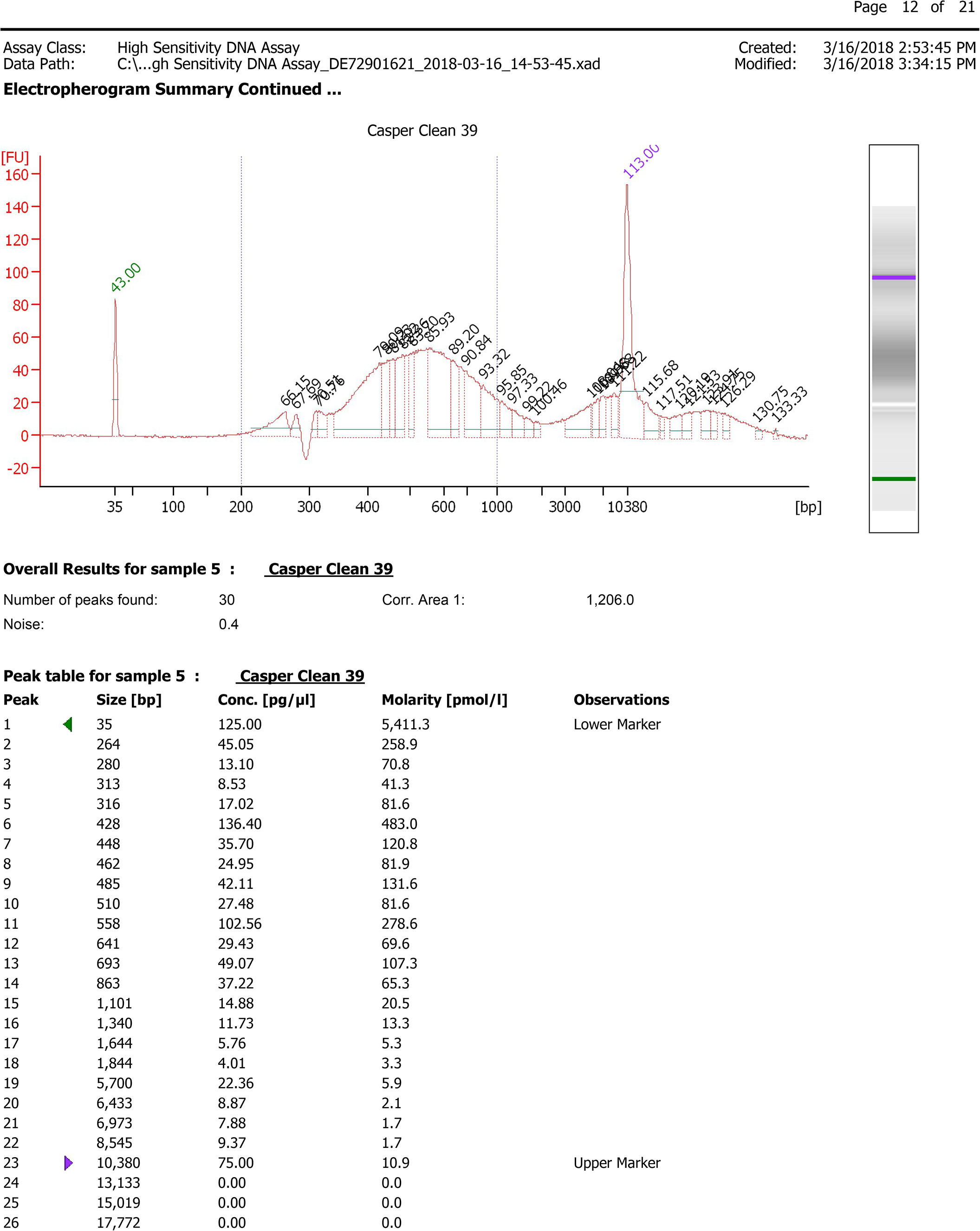
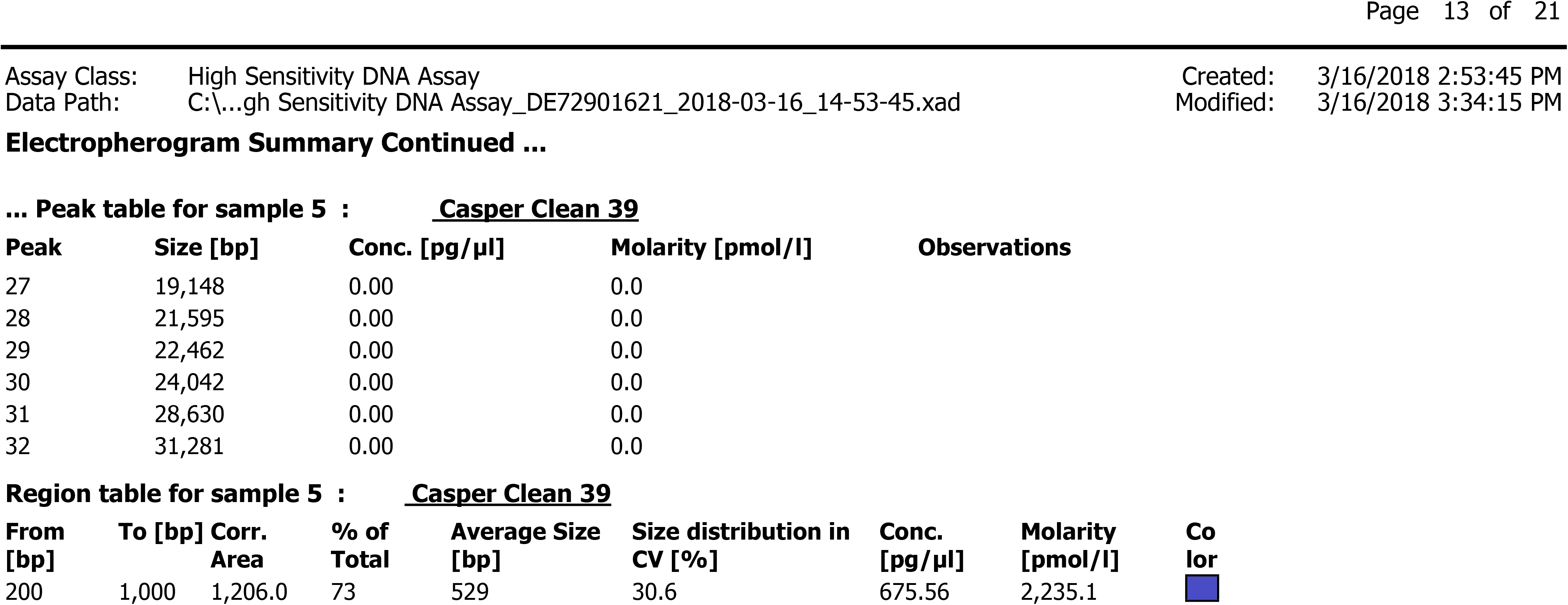
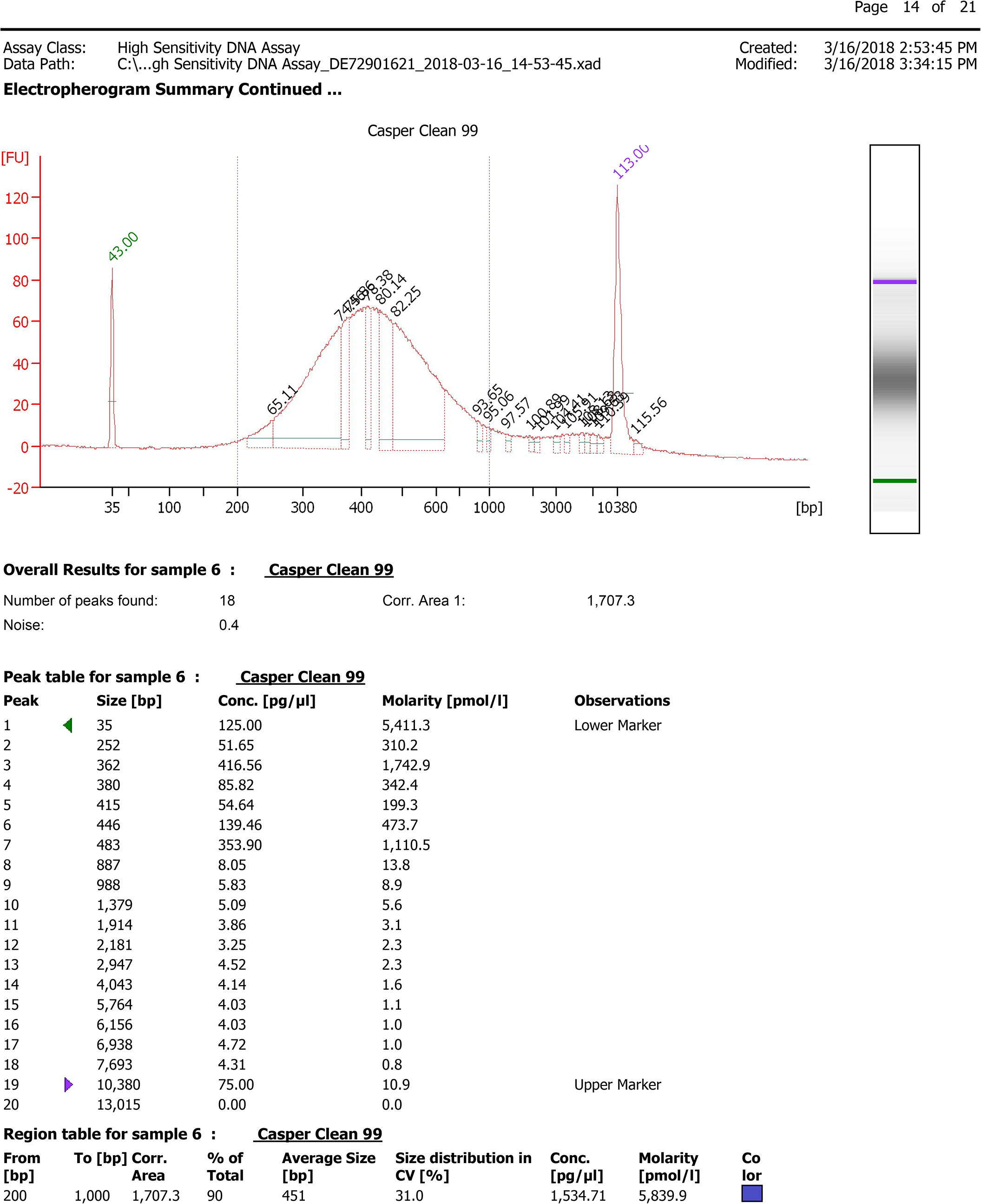
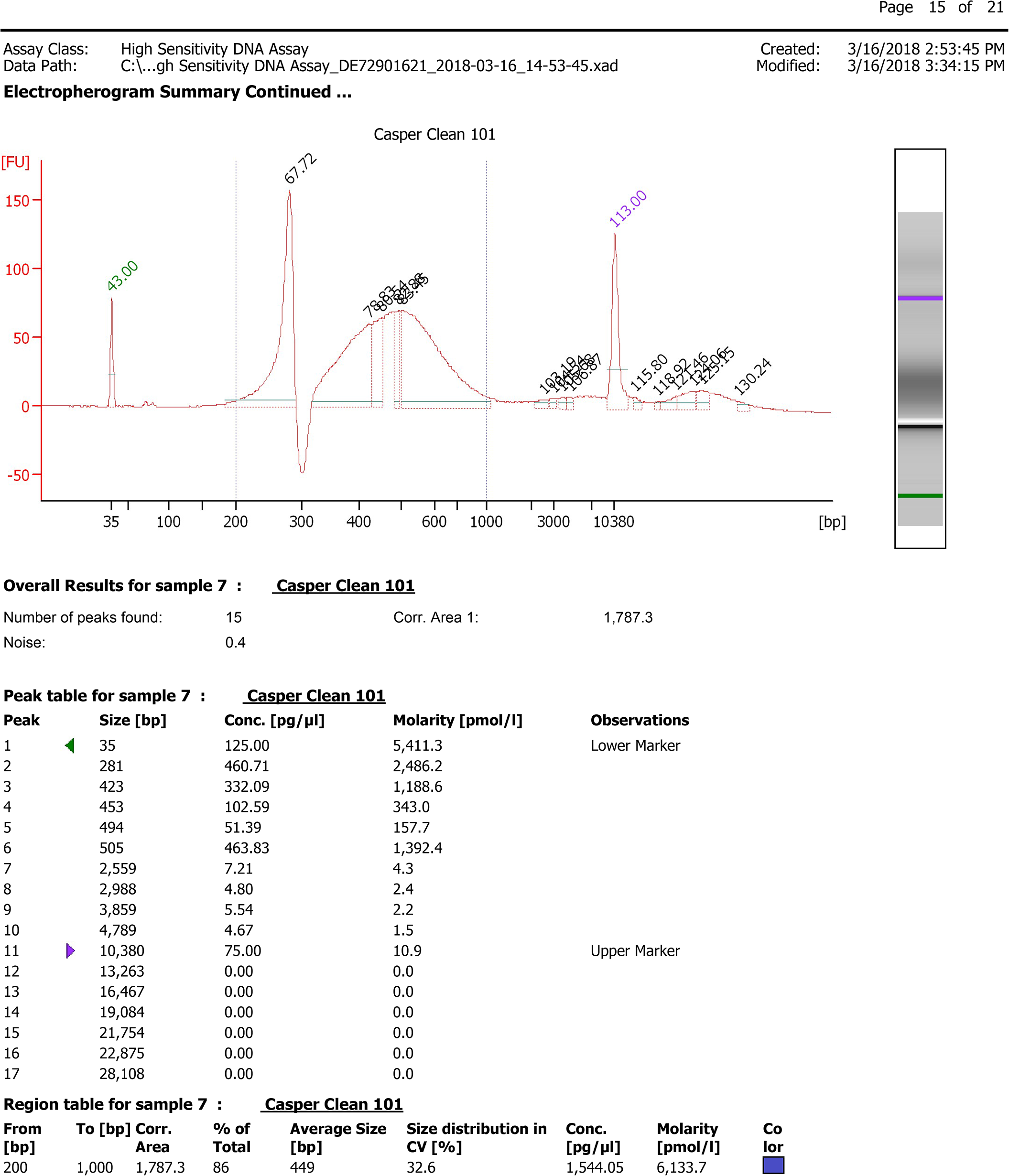
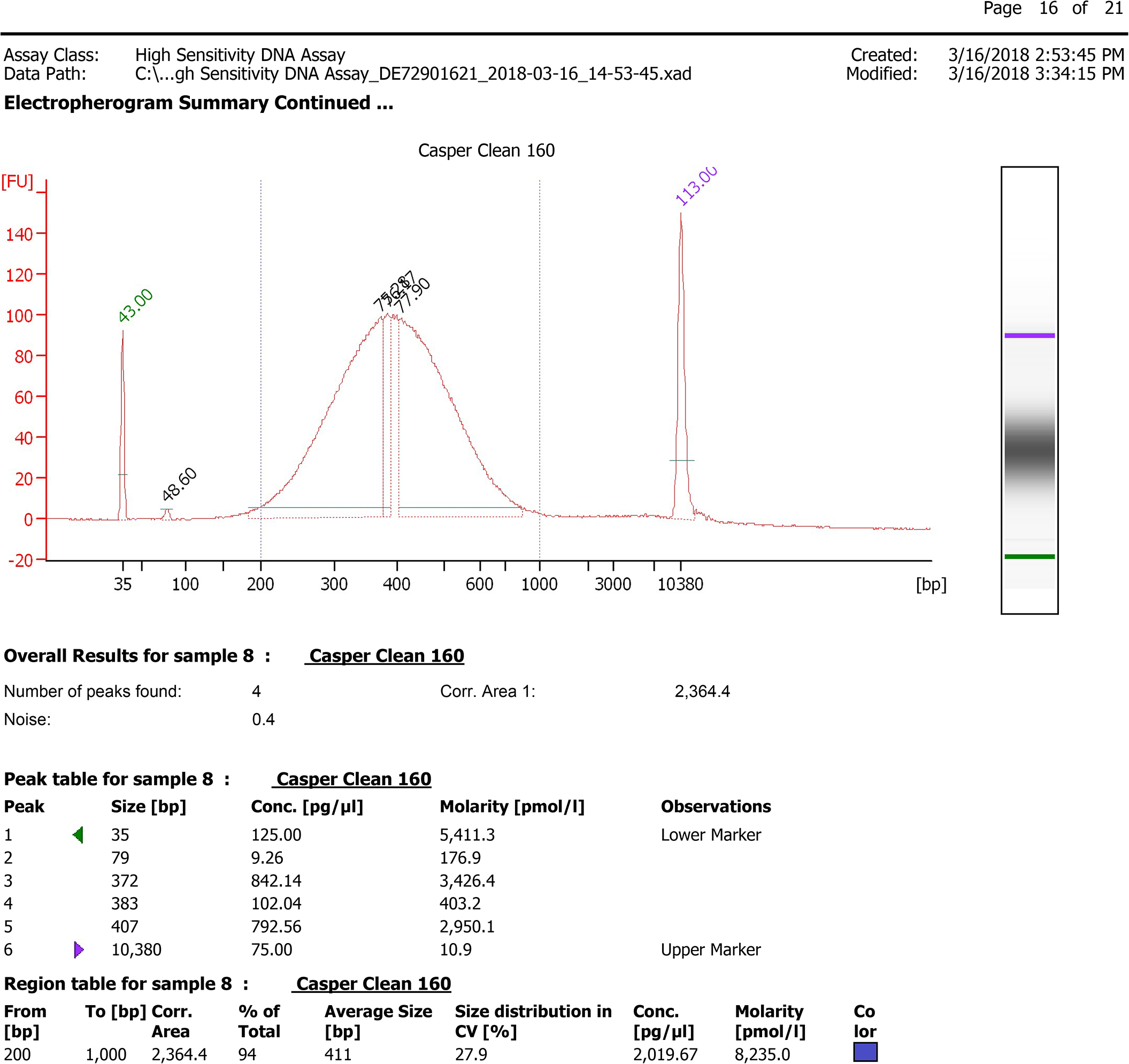
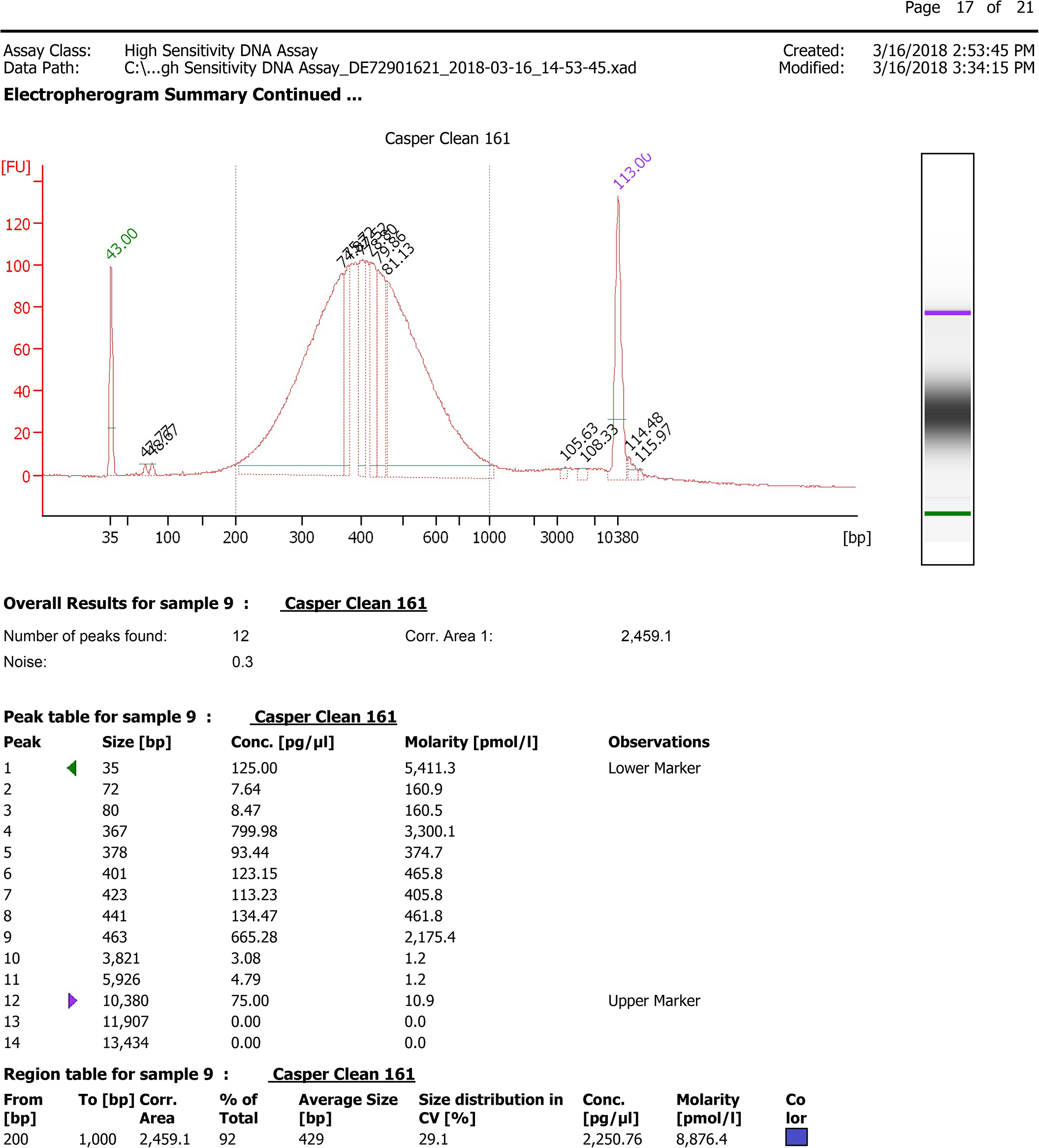
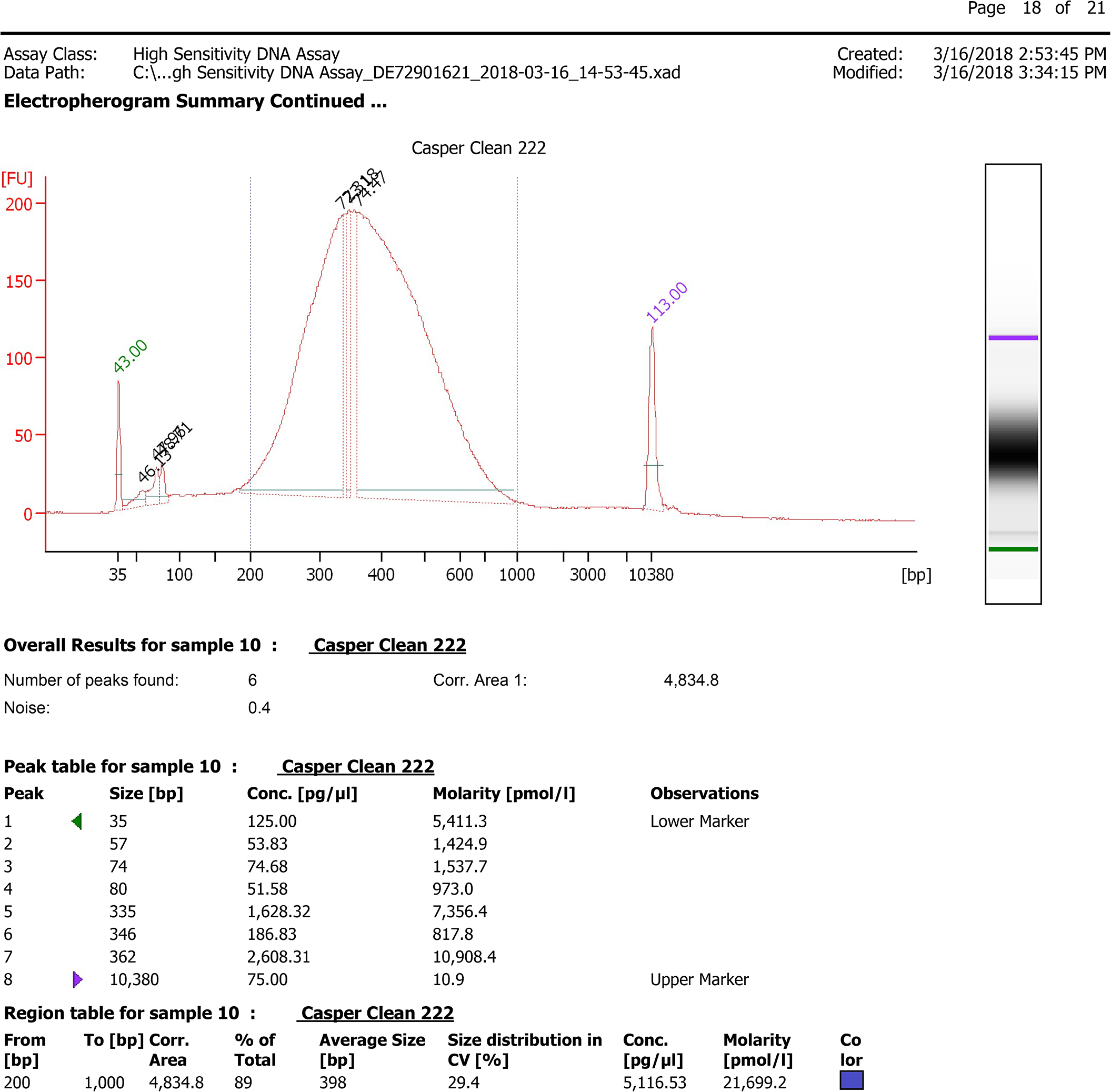
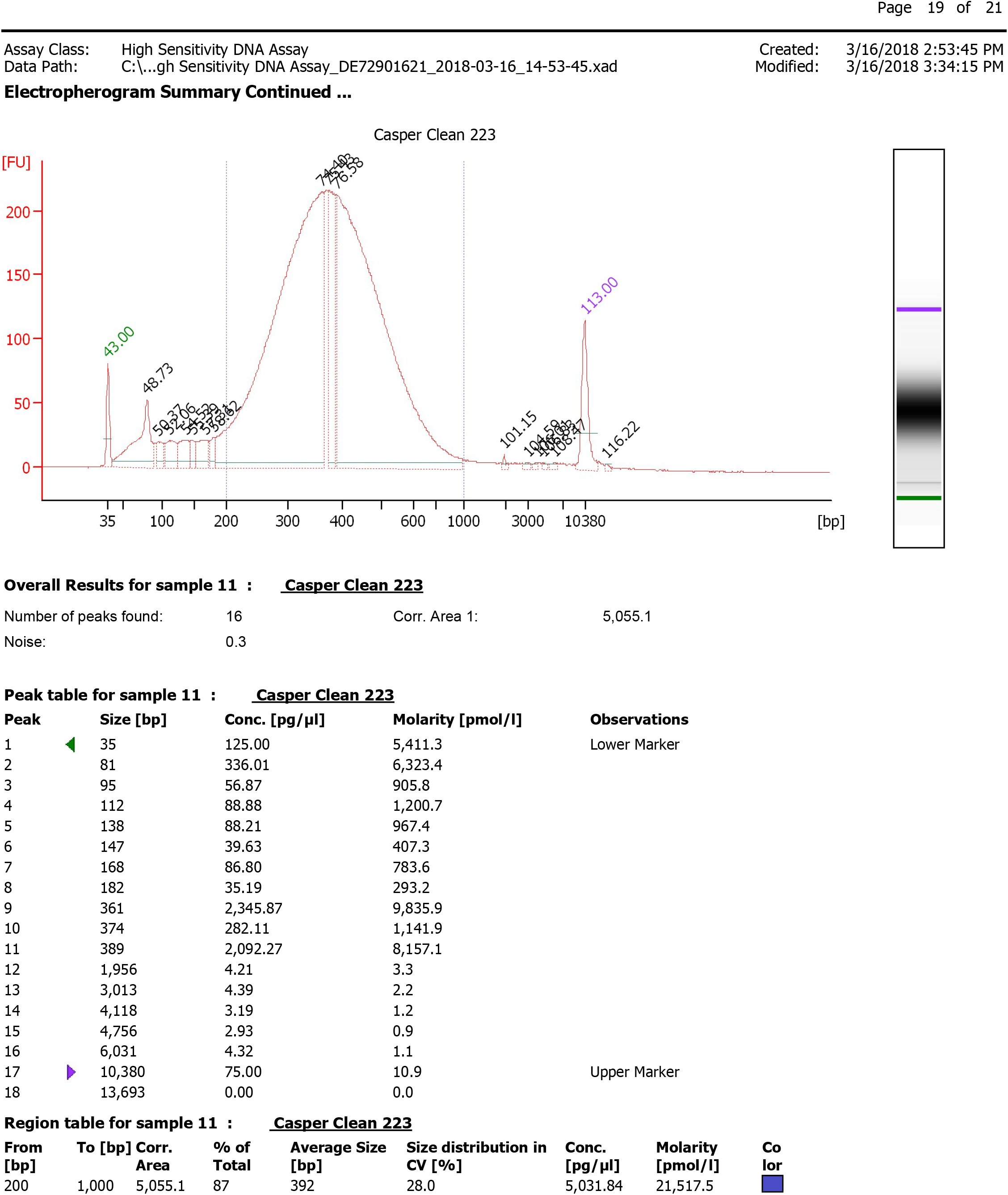
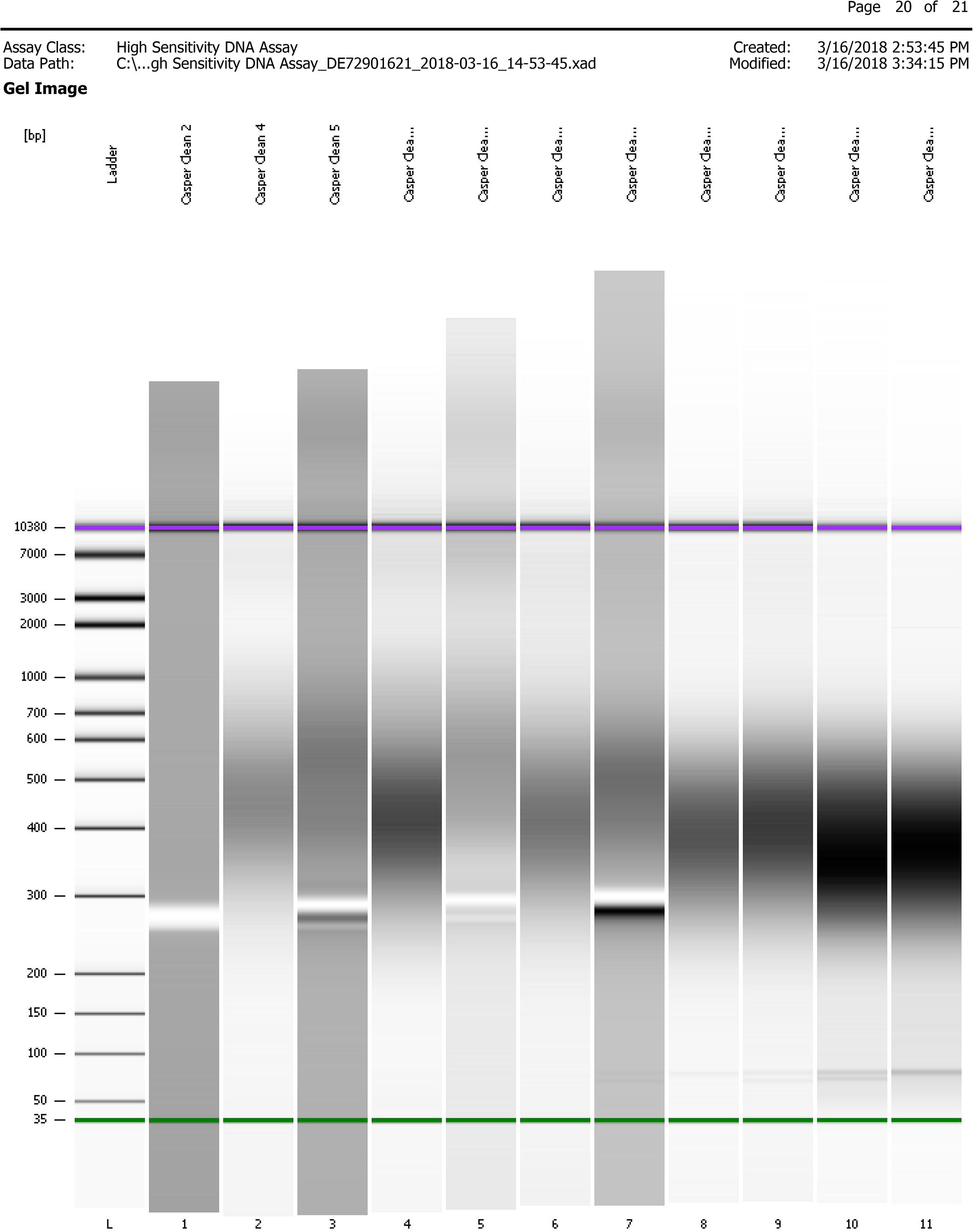
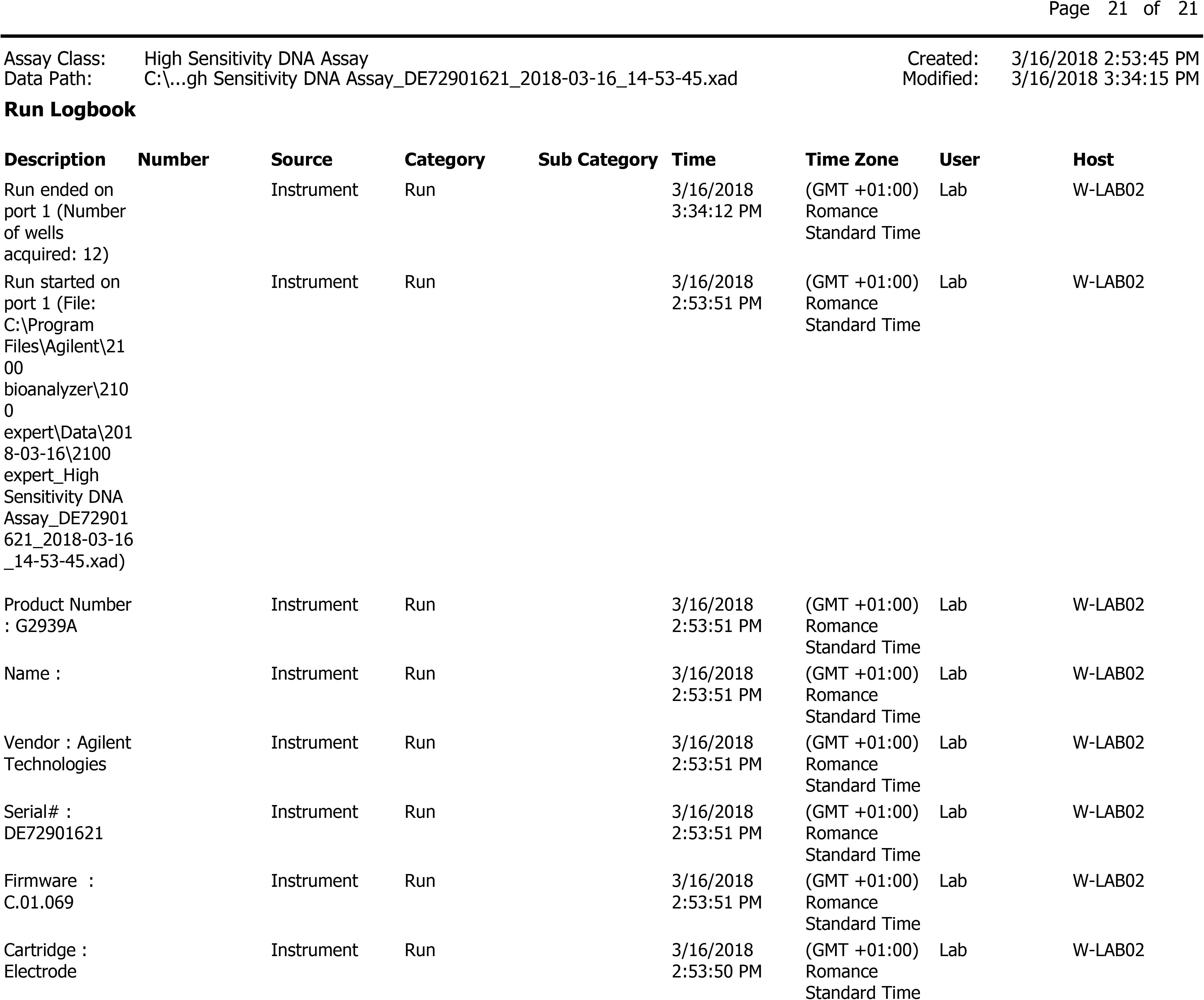

## Analysis: Experiments on the effect of sequencing platform and library preparation

### Introduction

The purpose of the following code is to assess the effect of library preparation using KAPA (PCR free), NEXTflex (PCR free), and Nextera (PCR step), and Illumina sequencing platforms (HiSeq and NextSeq) on two different microbial communities. Raw data is available online (Include the ENA reference). Samples have been quality processed and mapped with MGmapper extracting read counts mapping to the different genera.

### Explanation of samples naming in the taxonomy table

DTU: Performed at the Technical university of Denmark followed by the year of sampling. LPSX: Part of the study on library preparation and sequencing platform study. HX: Part of the handling experiment study included here to represent one library preparation sequencing platform method. P1 & P2: Pig feces 1 and 2. S1 and S2: Sewage 1 and 2. KA: Kapa library preparation. NF: NEXTflex library preparation. NX: Nextera library preparation (Can both be NX1 and NX2 representing the same process to investigate variance associated with redoing library preparation and sequencing). HI: HiSeq. NS: NextSeq. 0h: Direct processing after sample collection, samples were not stored. 64h_80C: After sample collection alliquouts of the same sample as the one processed directly were stored at −80?C for 64 h. a, b, c: Indicating storage replicates, only duplicates where included in this study. MG_XXX: Internal metagenomics sample number.

### Metadata

Contains information on how the samples were processed, sequencing performance, and info on mapping to different databases

**Feature** Contains taxonomic feature information

###Analysis notes All analysis adhere to the compositional data analysis framework by including an isometric log ratio transformation (ILR) to calculate Euclidean distances used to perform prinicipal component analysis (PCA), Heatmap sample clustering and boxplots. A centered log ratio transformation (CLR) were used when keeping genera information was important in sparse partial least square discriminant analysis (sPLS-DA) and redundancy analysis (rda). Important factors to be aware of in this analysis is that raw data is used, filtered according to an average count of 5, and zeroes are estimated with simple multiplicative replacement.

**Colors** In study design and heatmaps KAPA is purple (#984ea3), Nextera is orange (#ff7f00), and NEXTflex is red (#e41a1c). When performing sPLS-DA, PCA, rda the NEXTflex is run either with NextSeq is blue (#377eb8) or HiSeq is green (#4daf4a). Finally in PCA and rda Nextera is divided in 1 and 2, where 1 remains orange and 2 is brown (#993300)

…

#### Packages

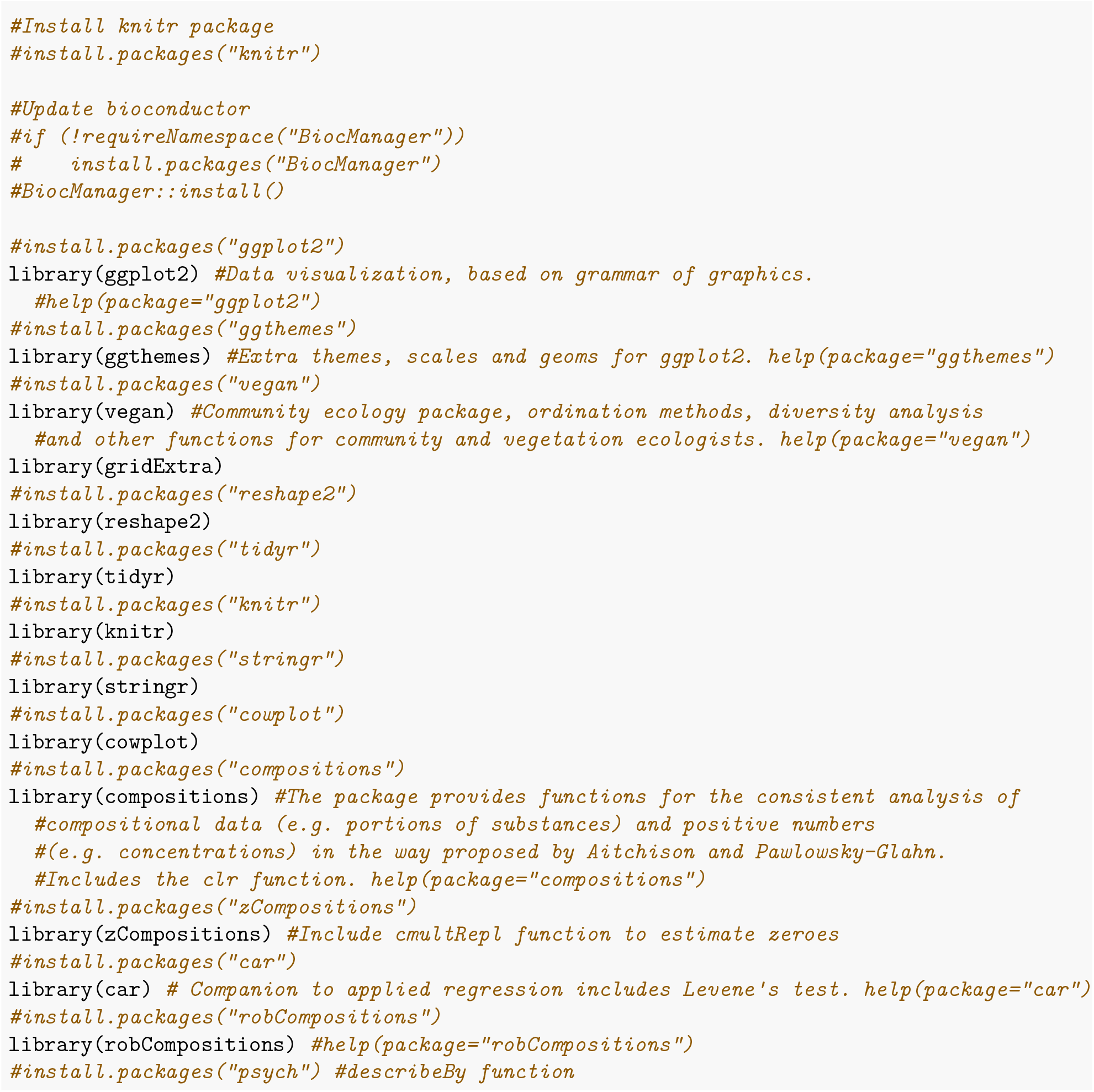
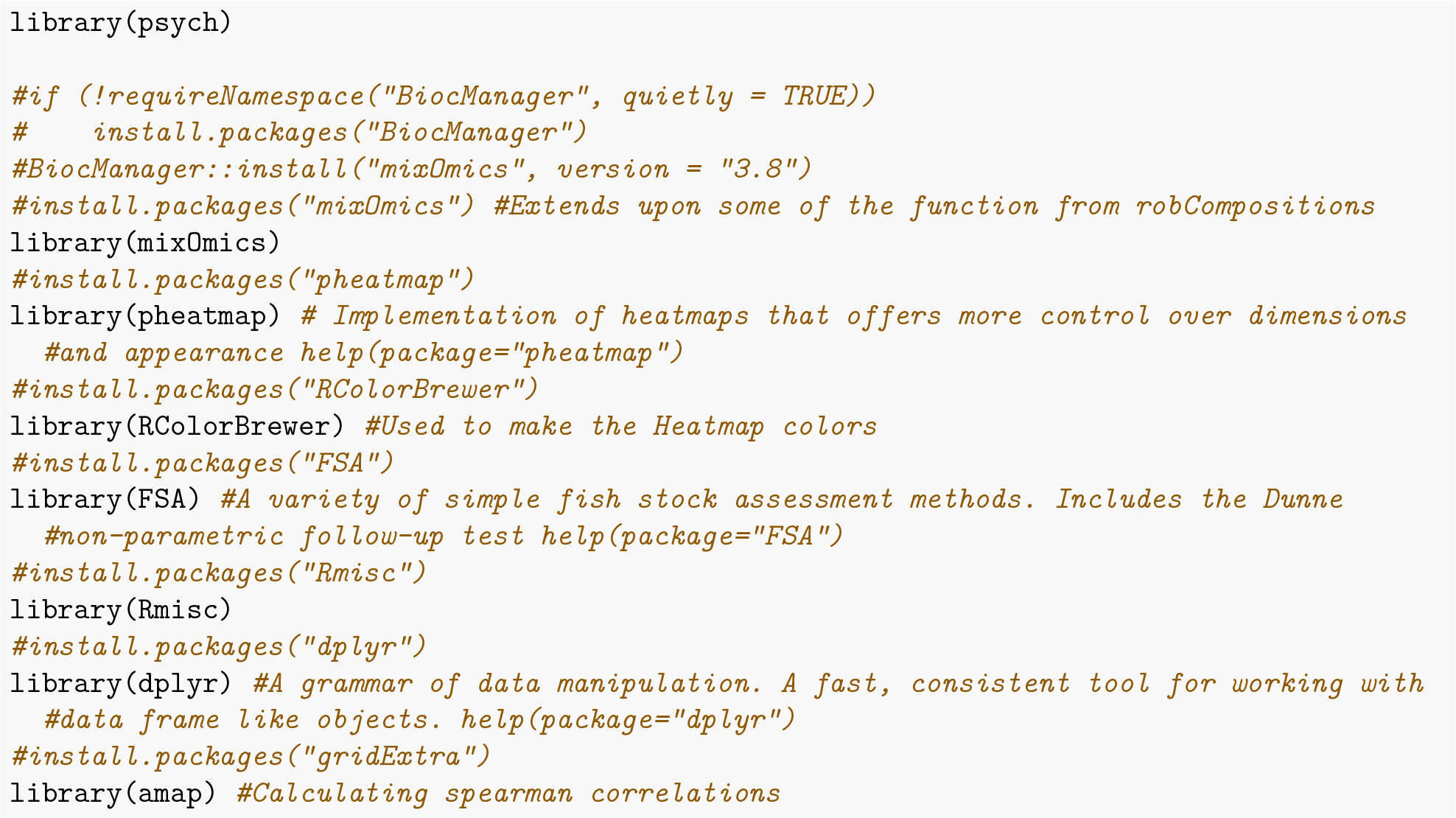

### Read in data

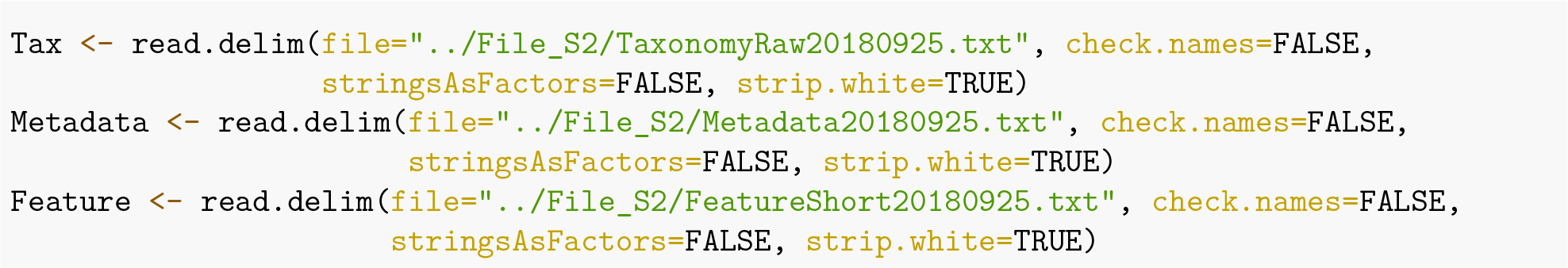

### Change naming

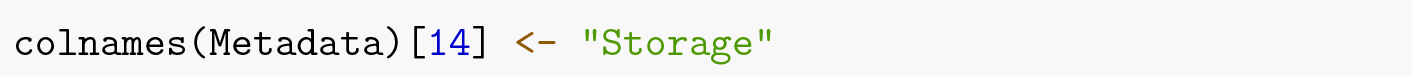

### Analysis

Chunks can be run by themselves or consecutively

#### Quality control of different sequencing specific parameters

Include QC on sequencing output and alpha-diversity summarized in S1_Table and rarefaction curves S2_Fig

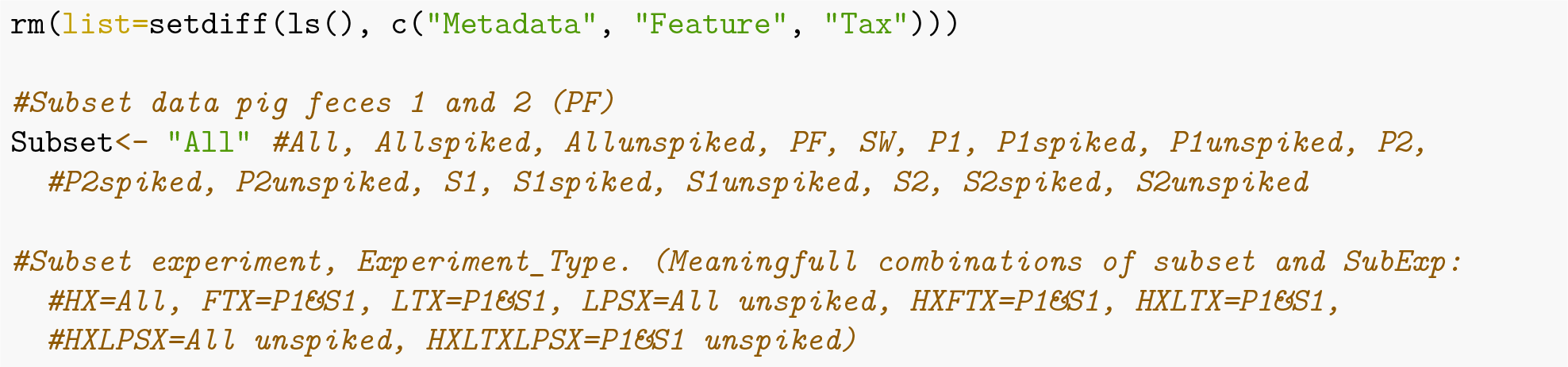
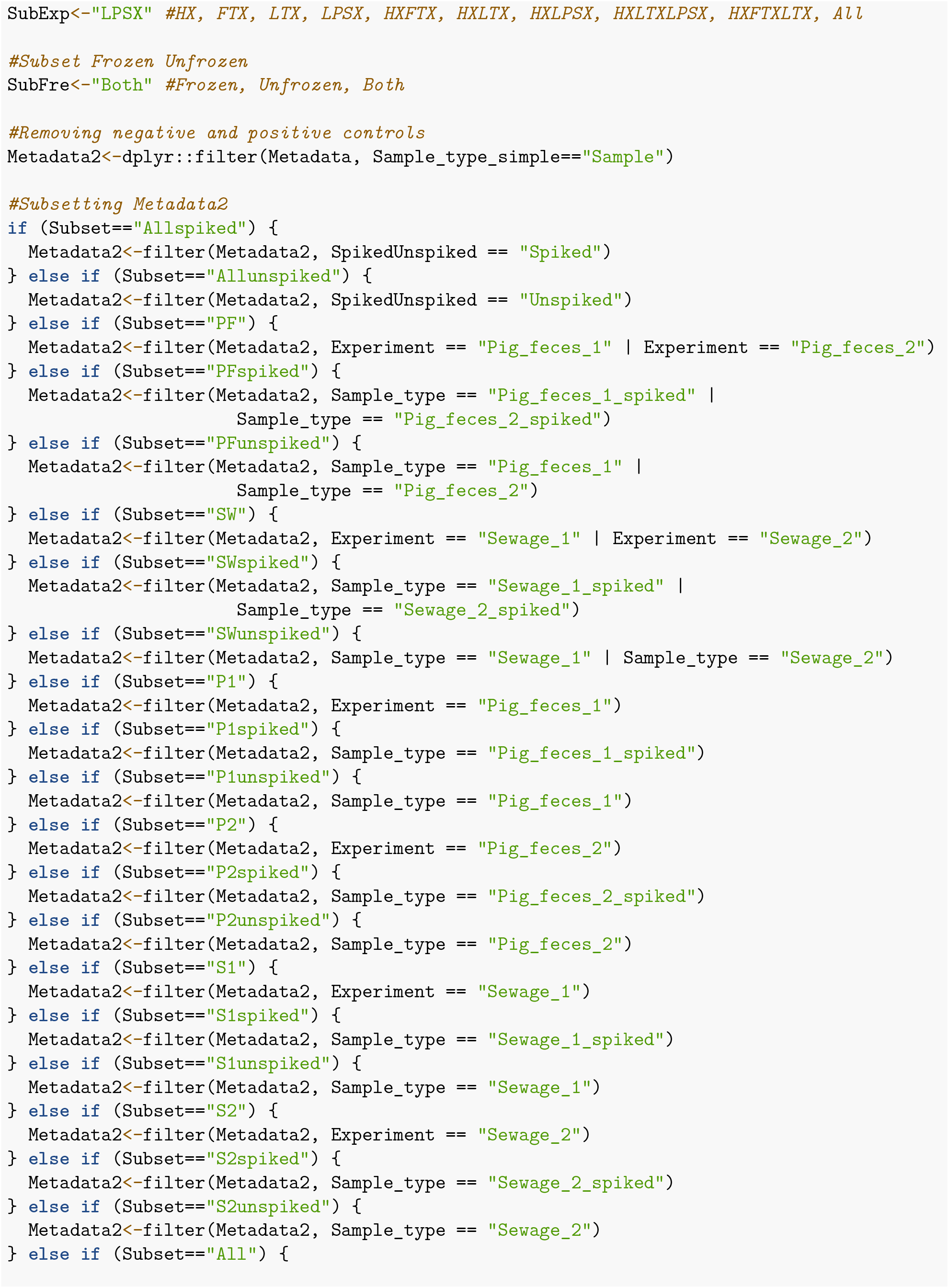
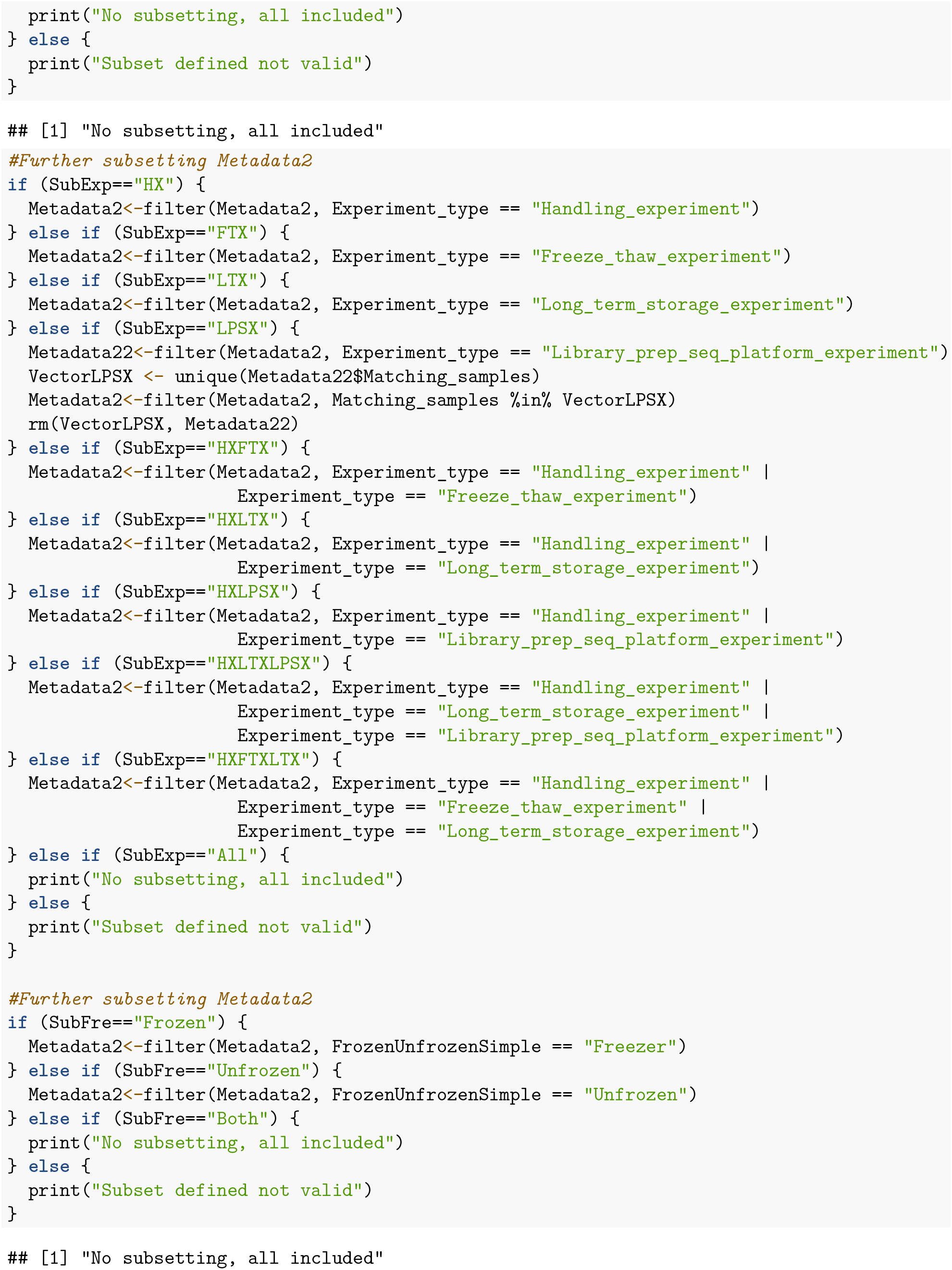
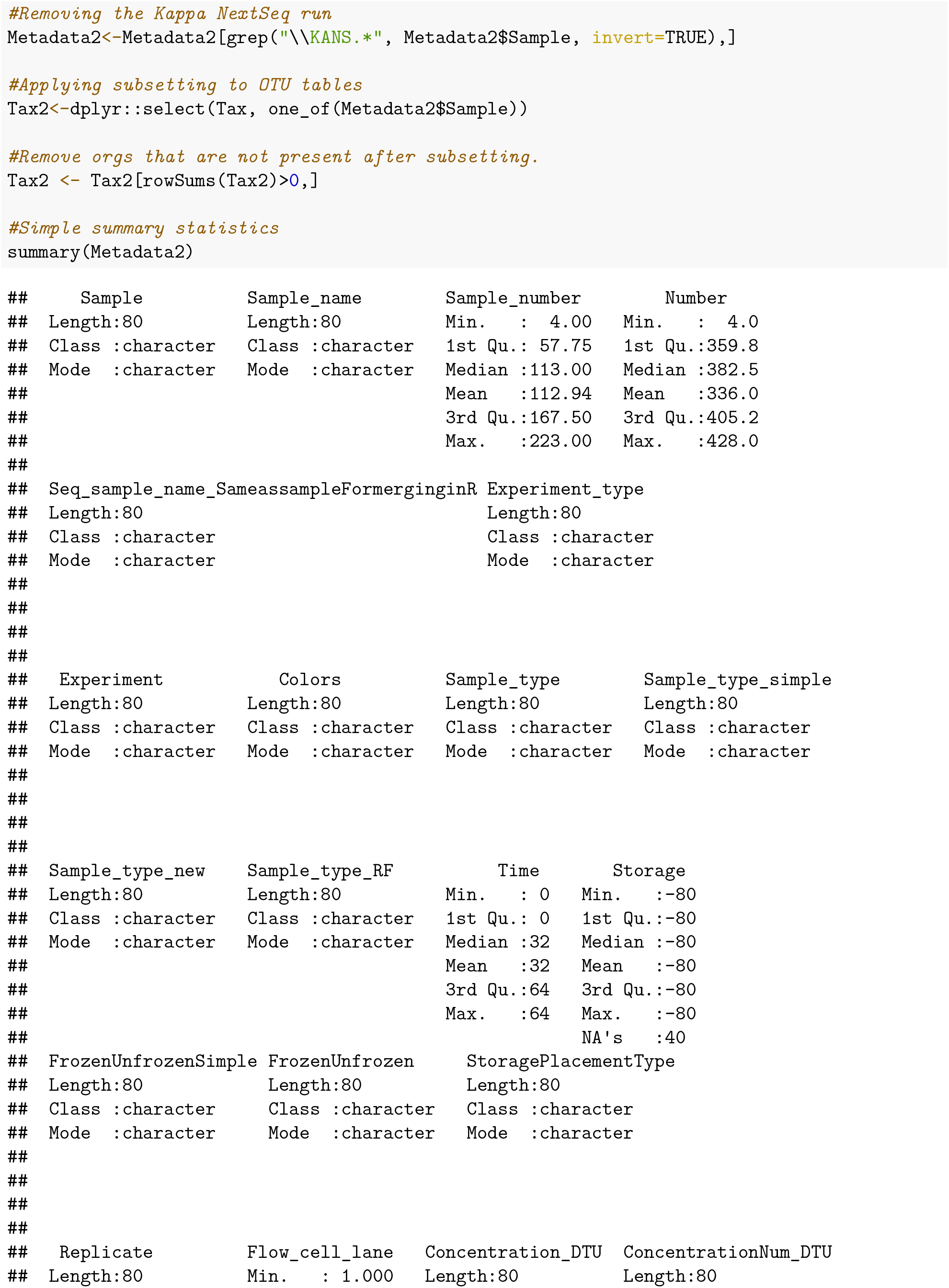
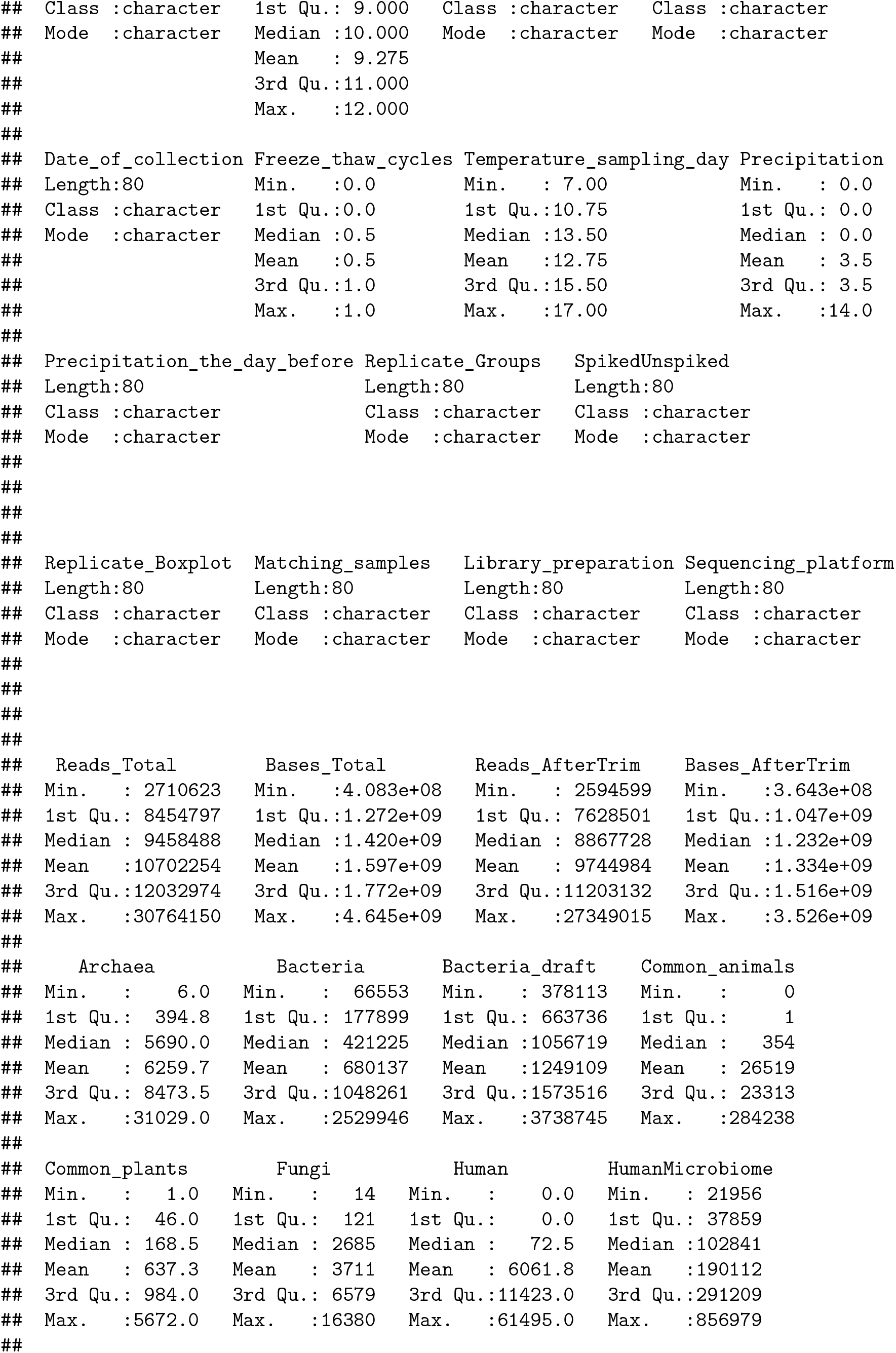
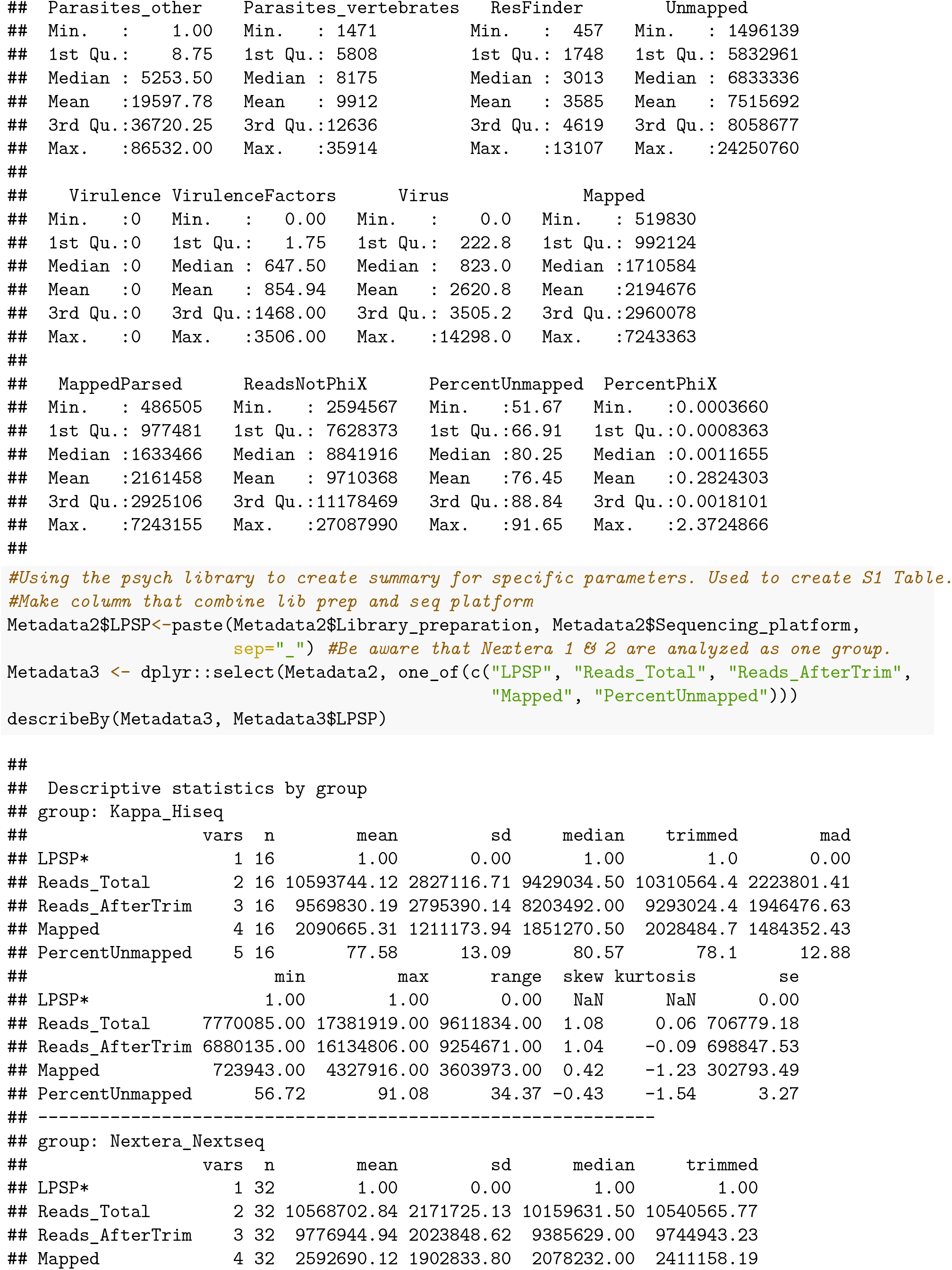
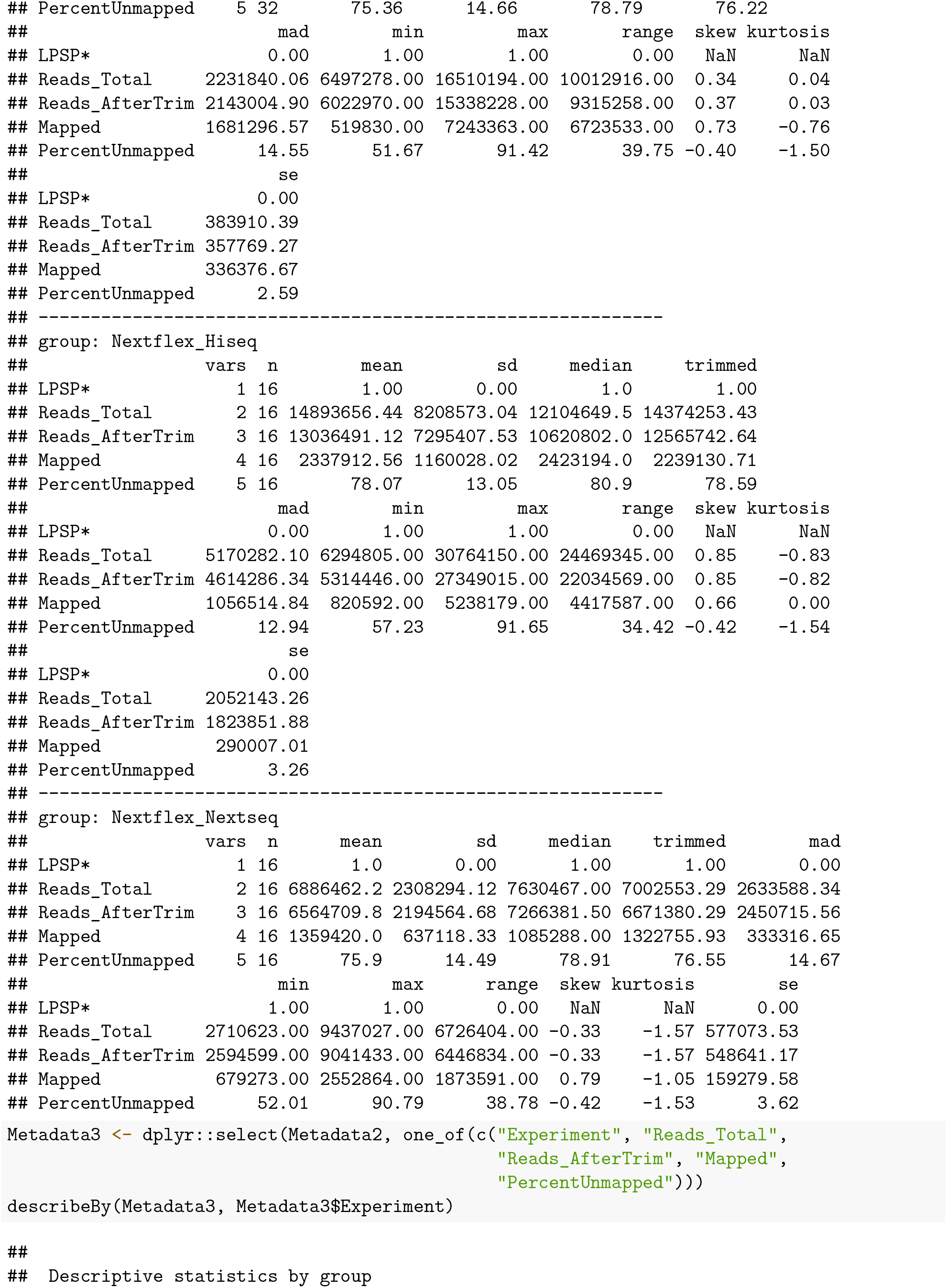
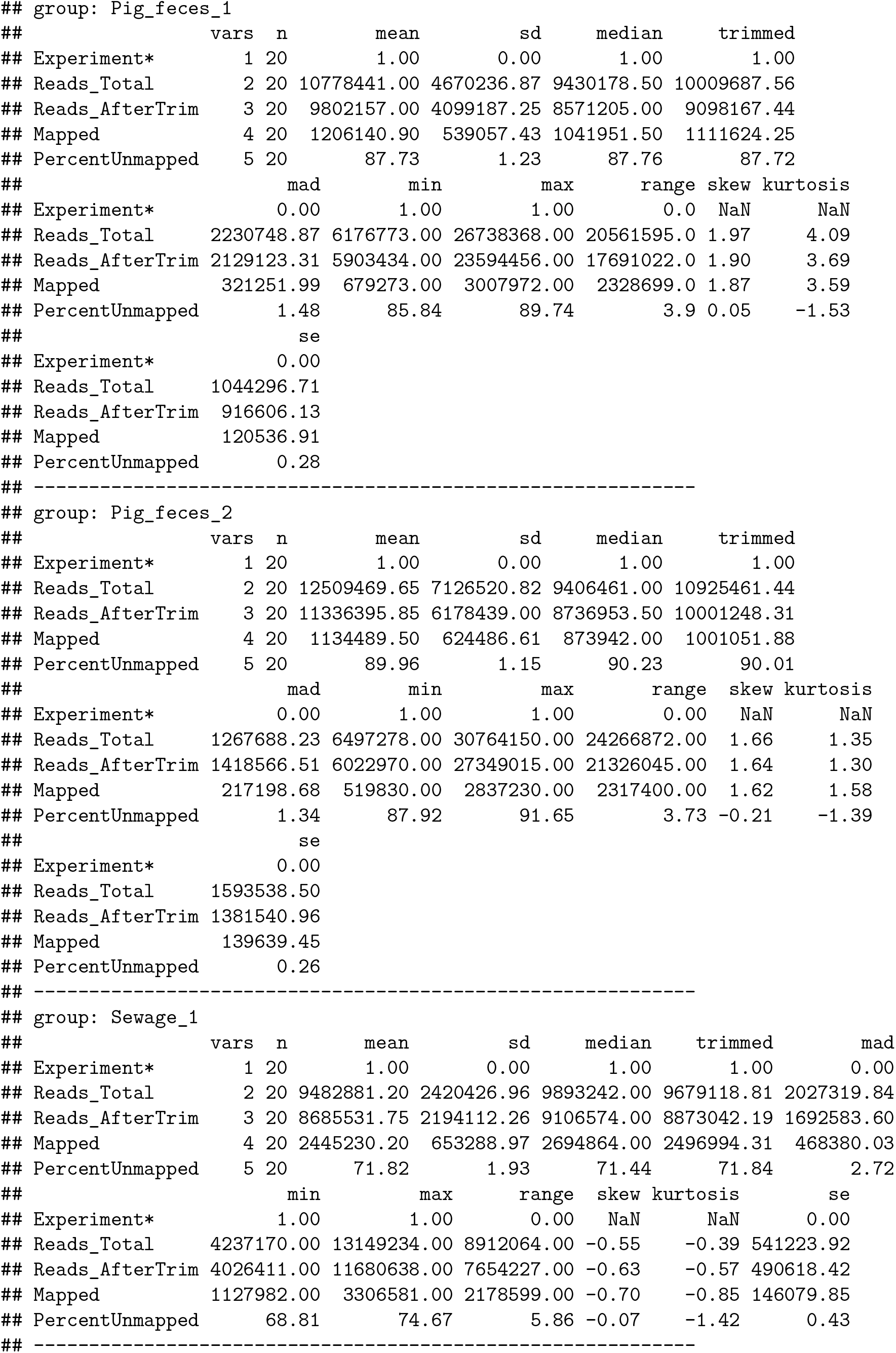
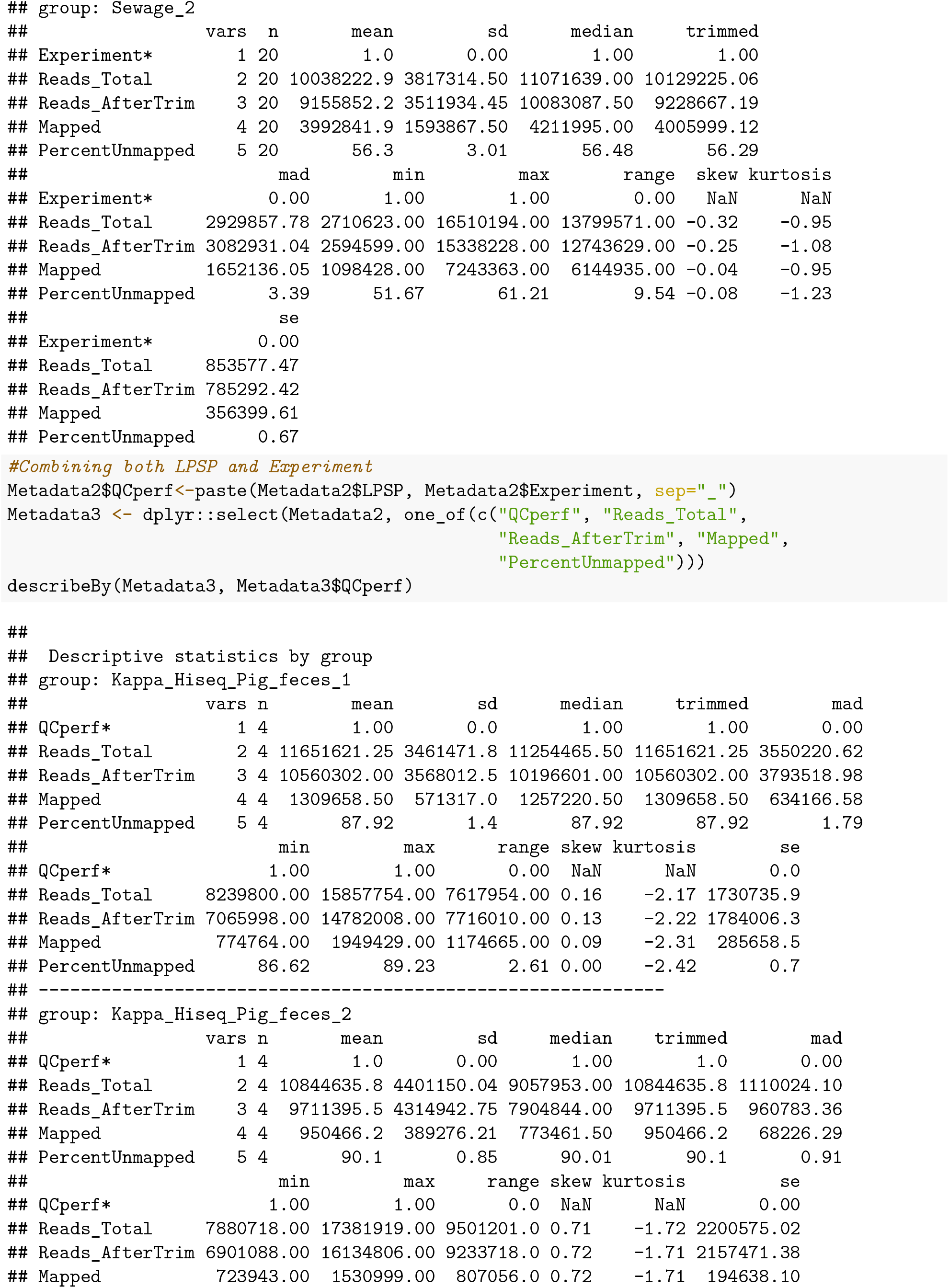
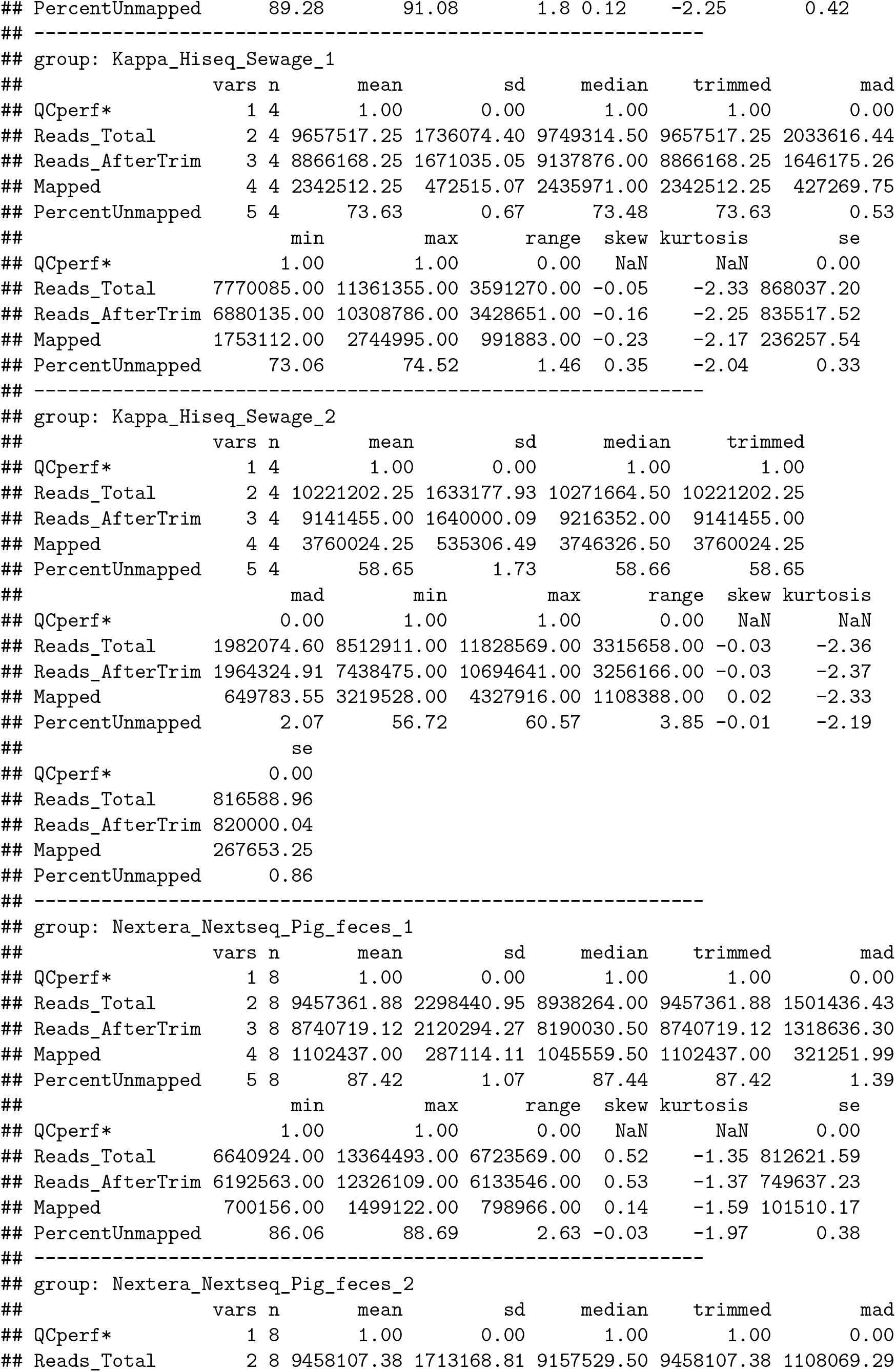
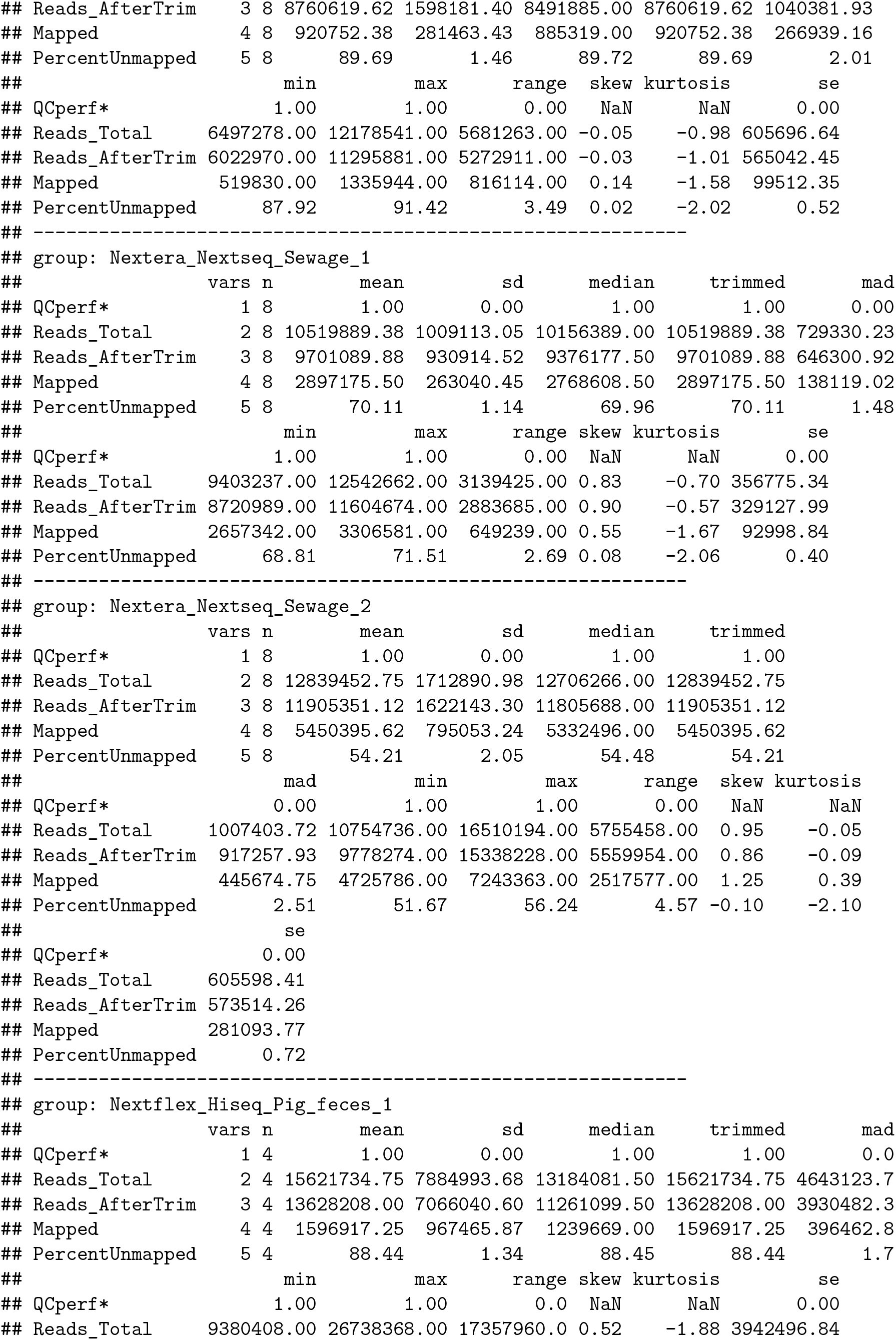
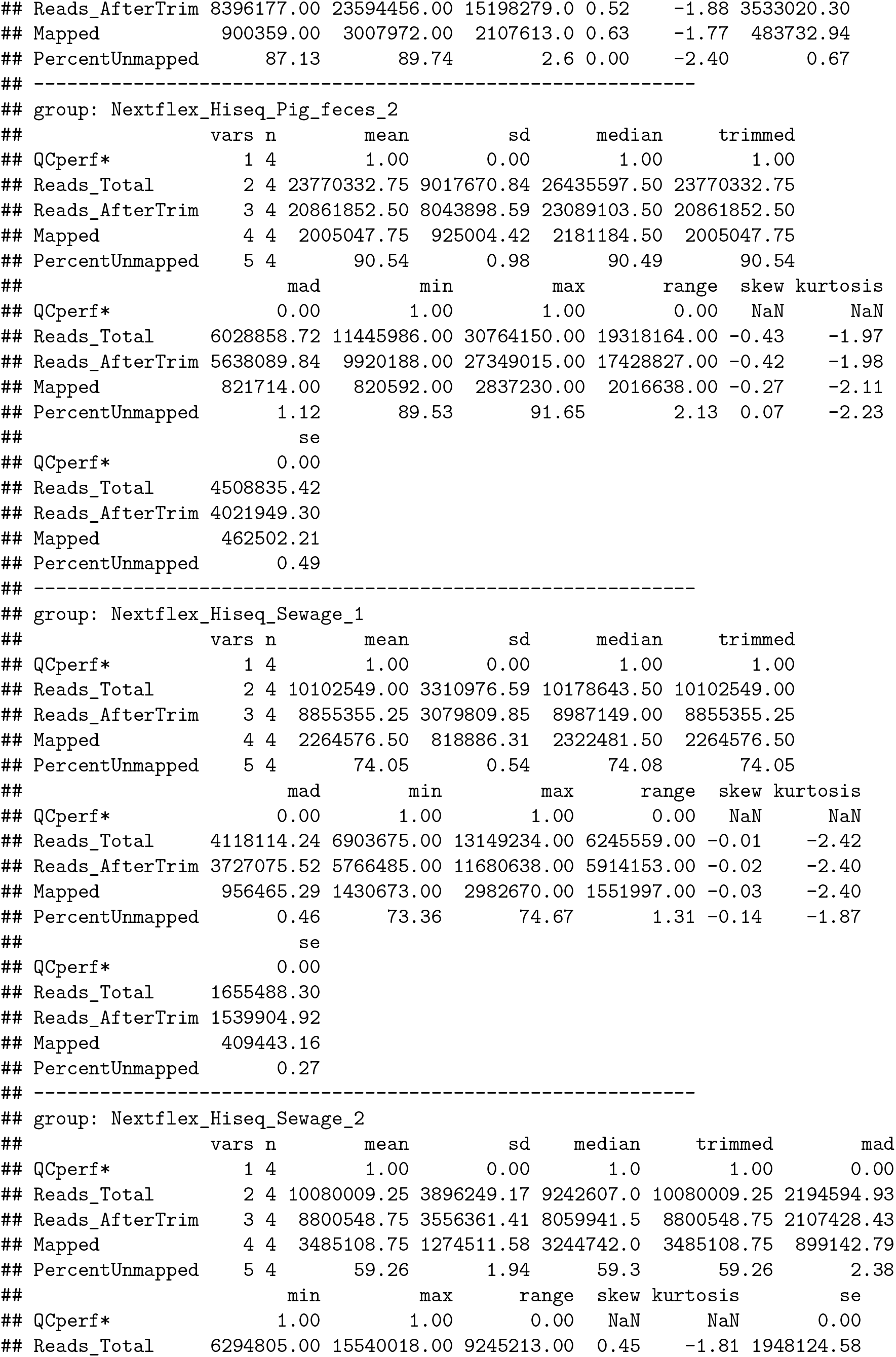
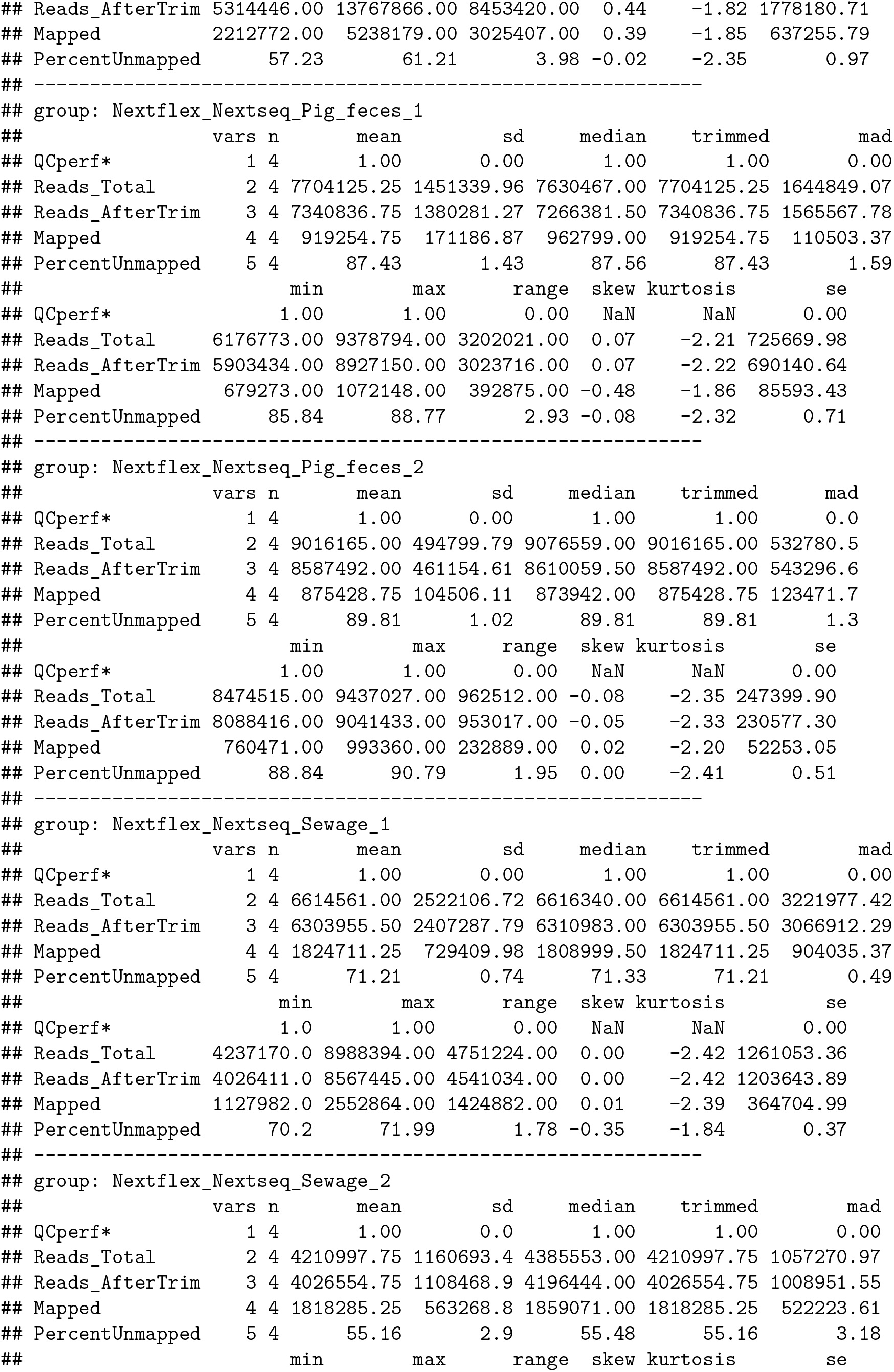
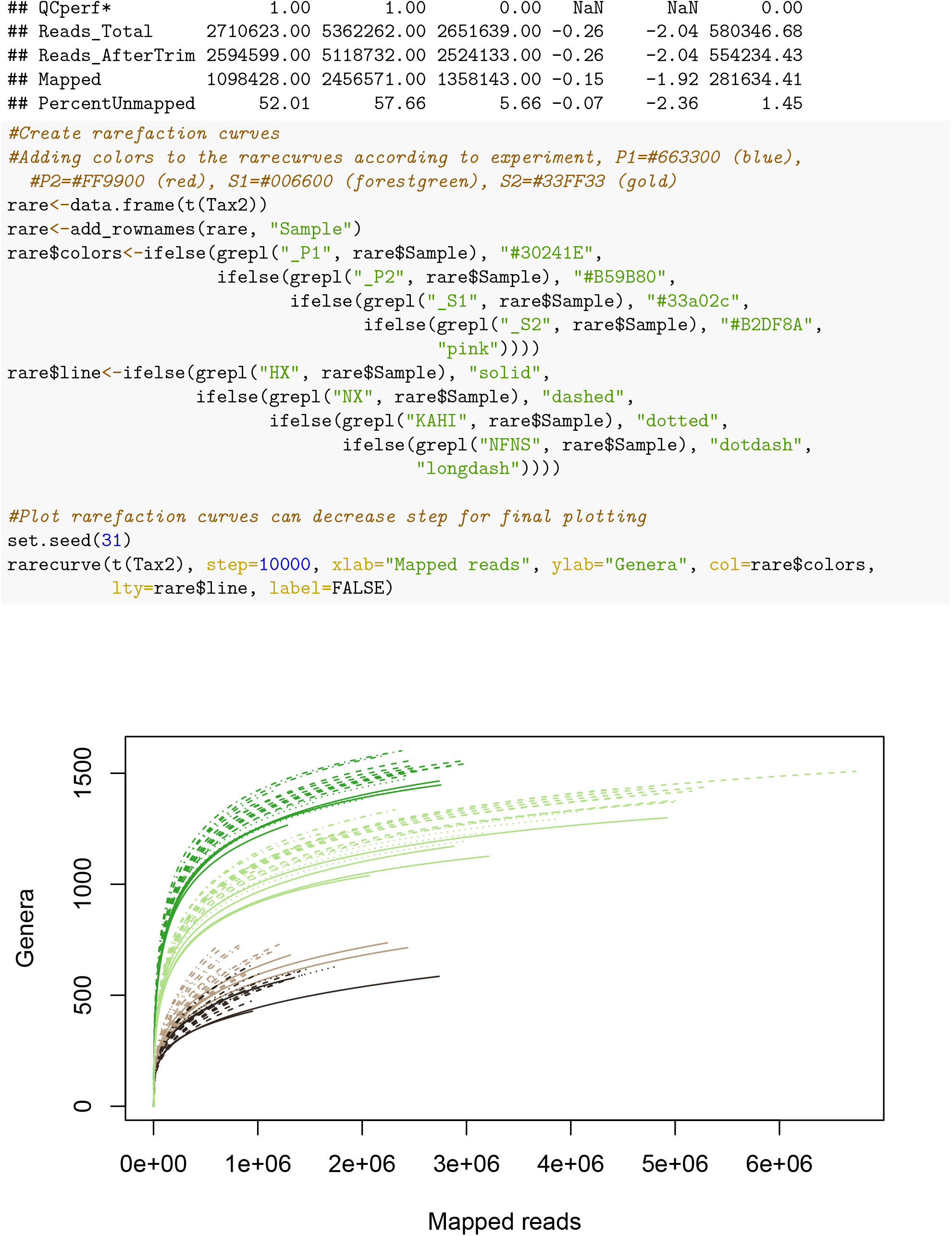
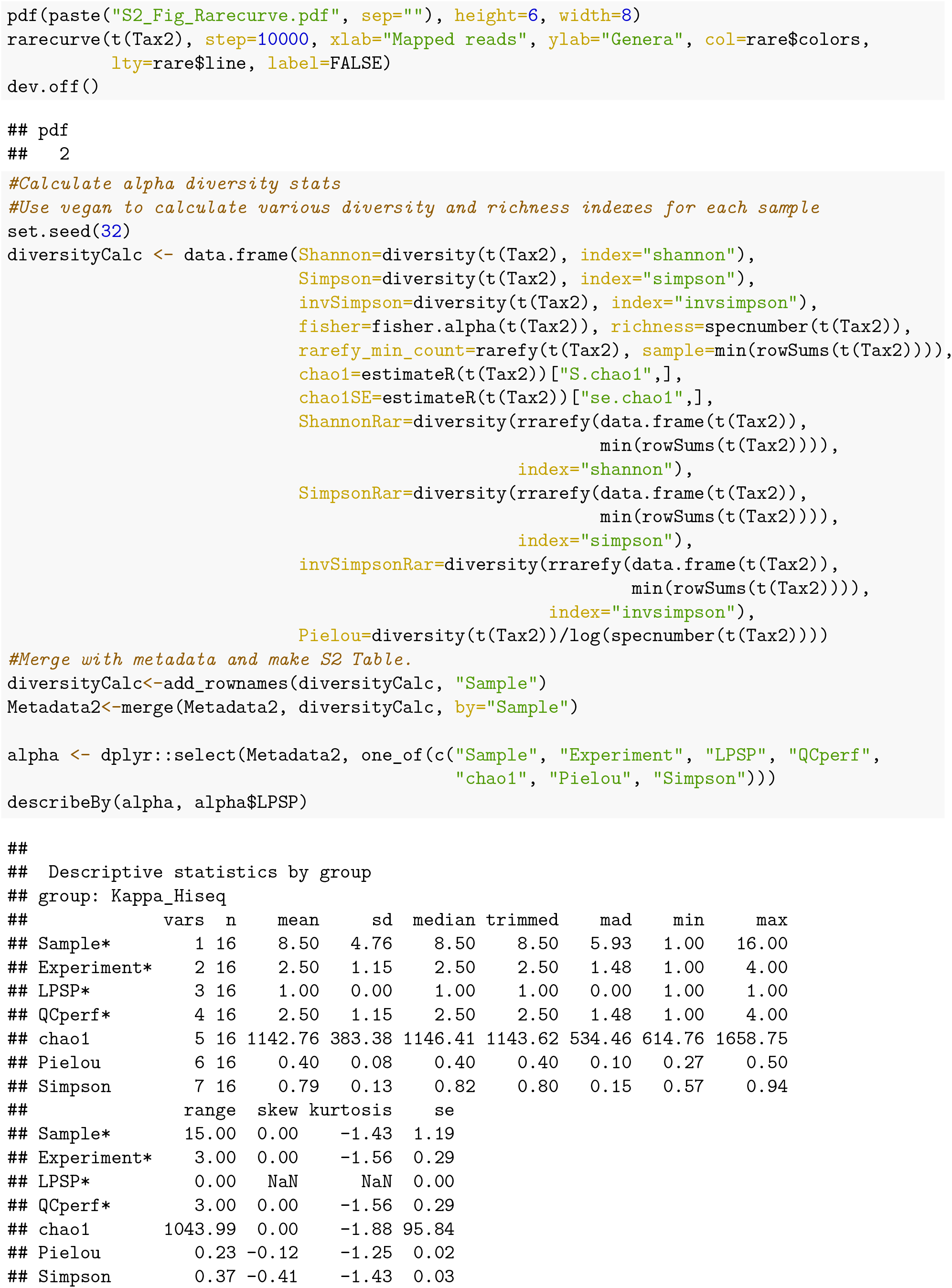
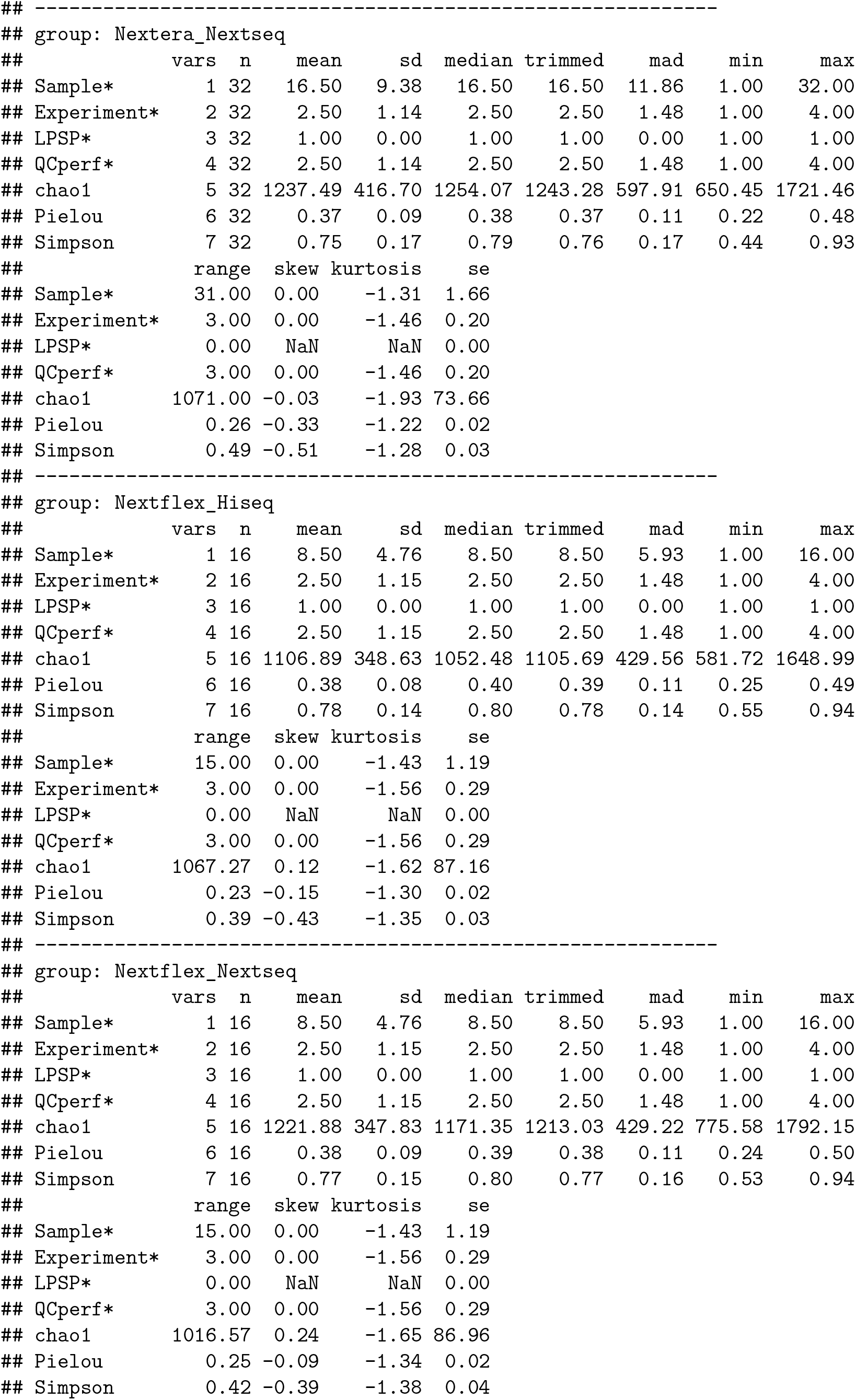
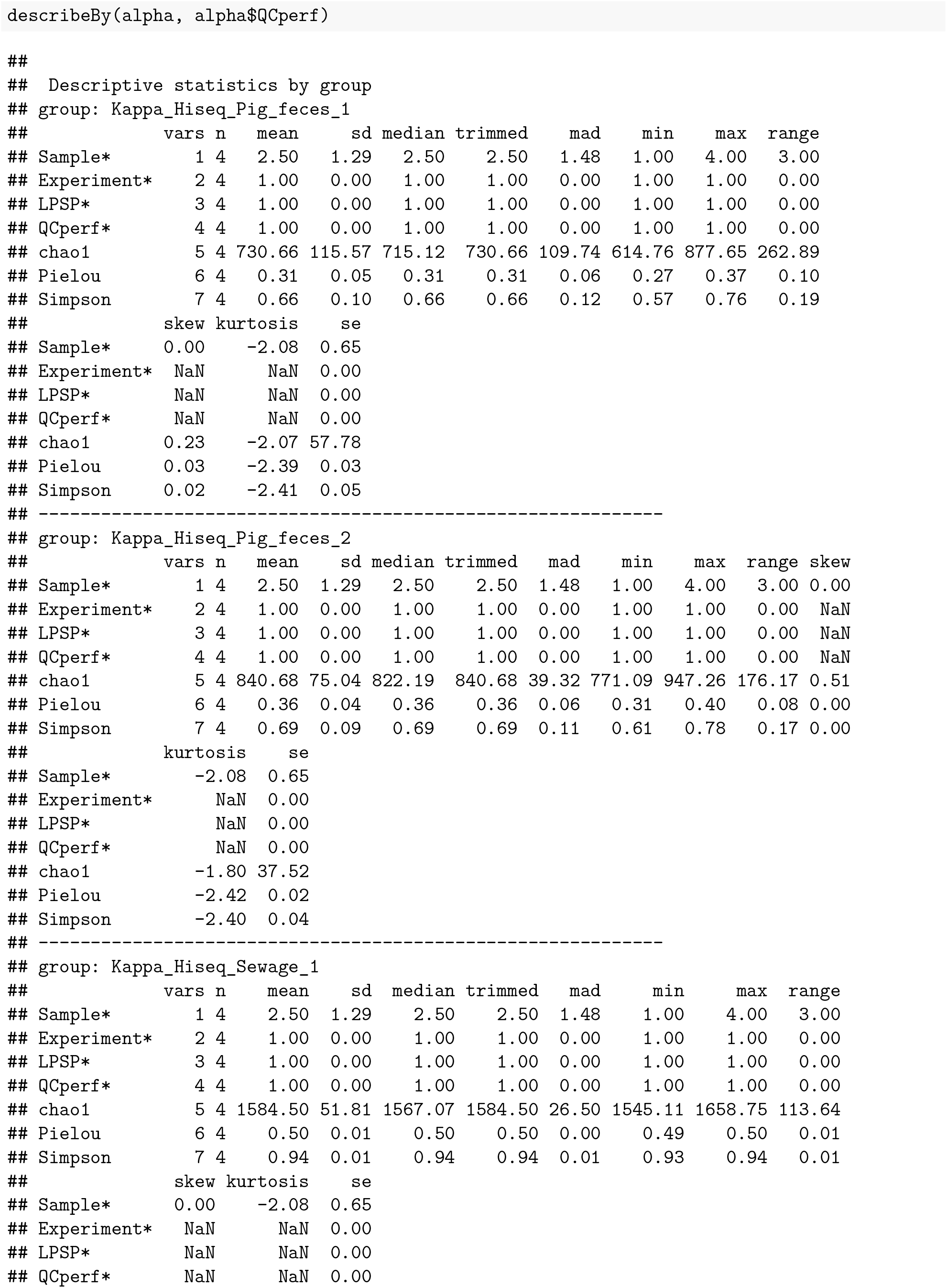
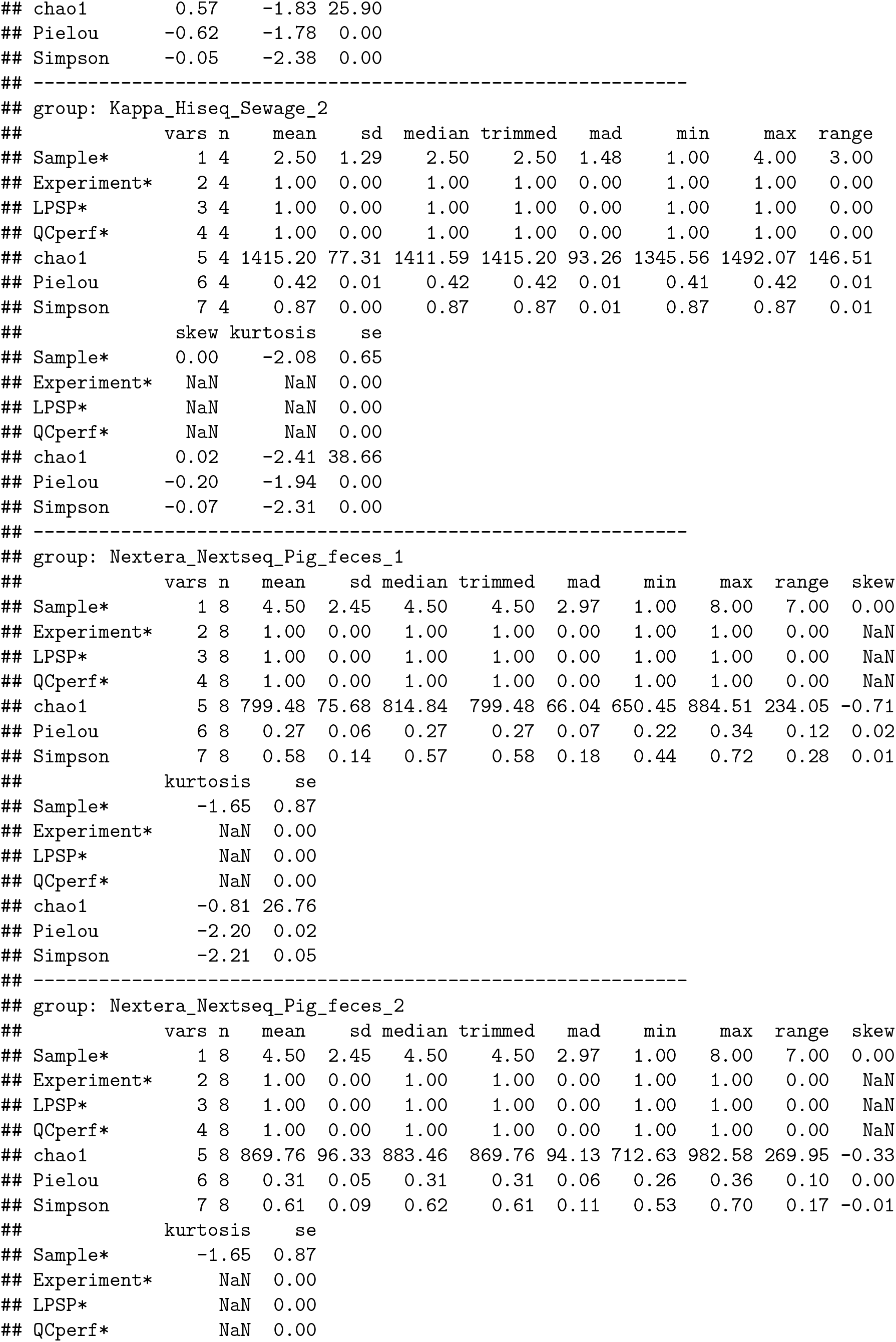
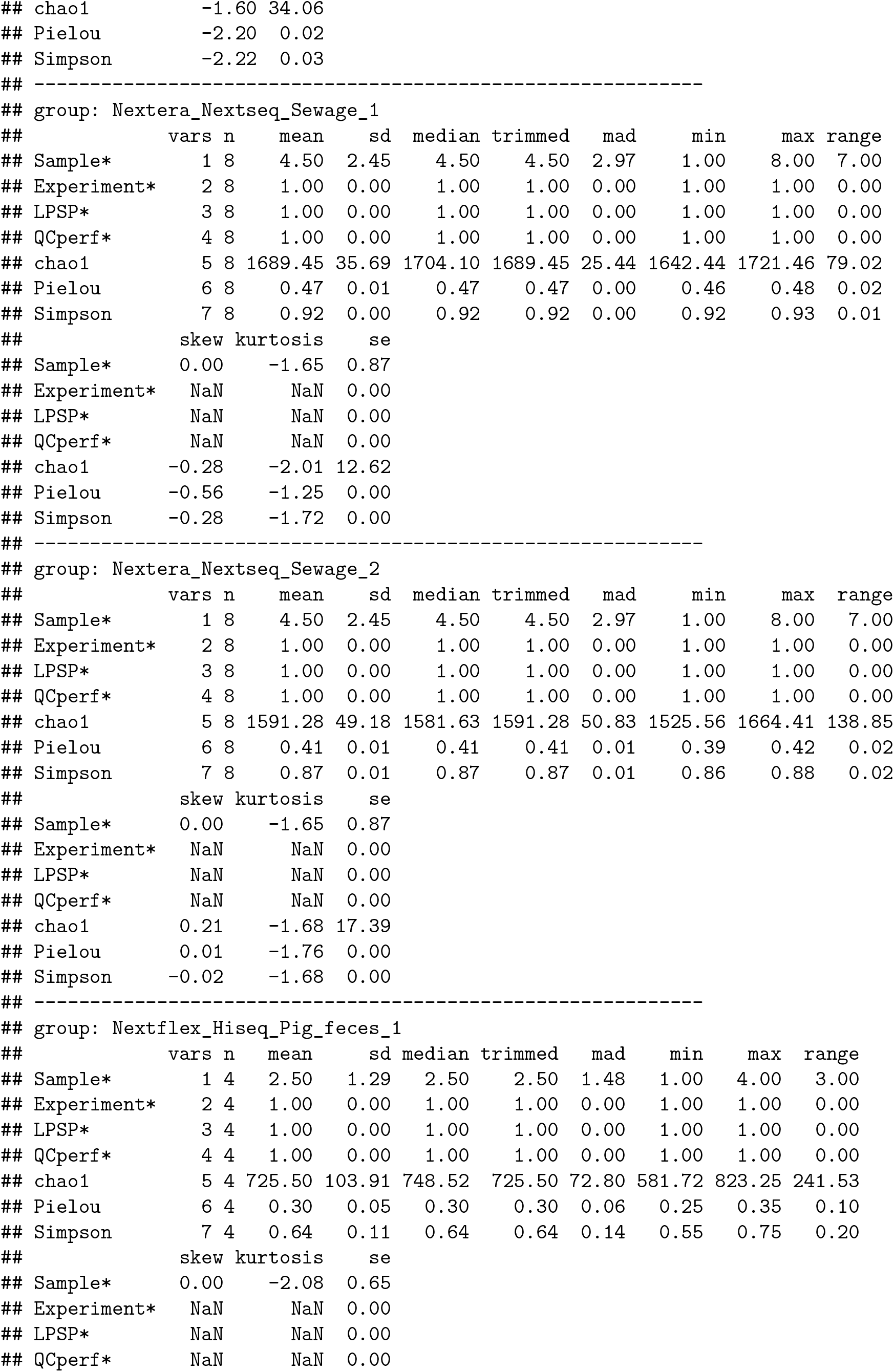
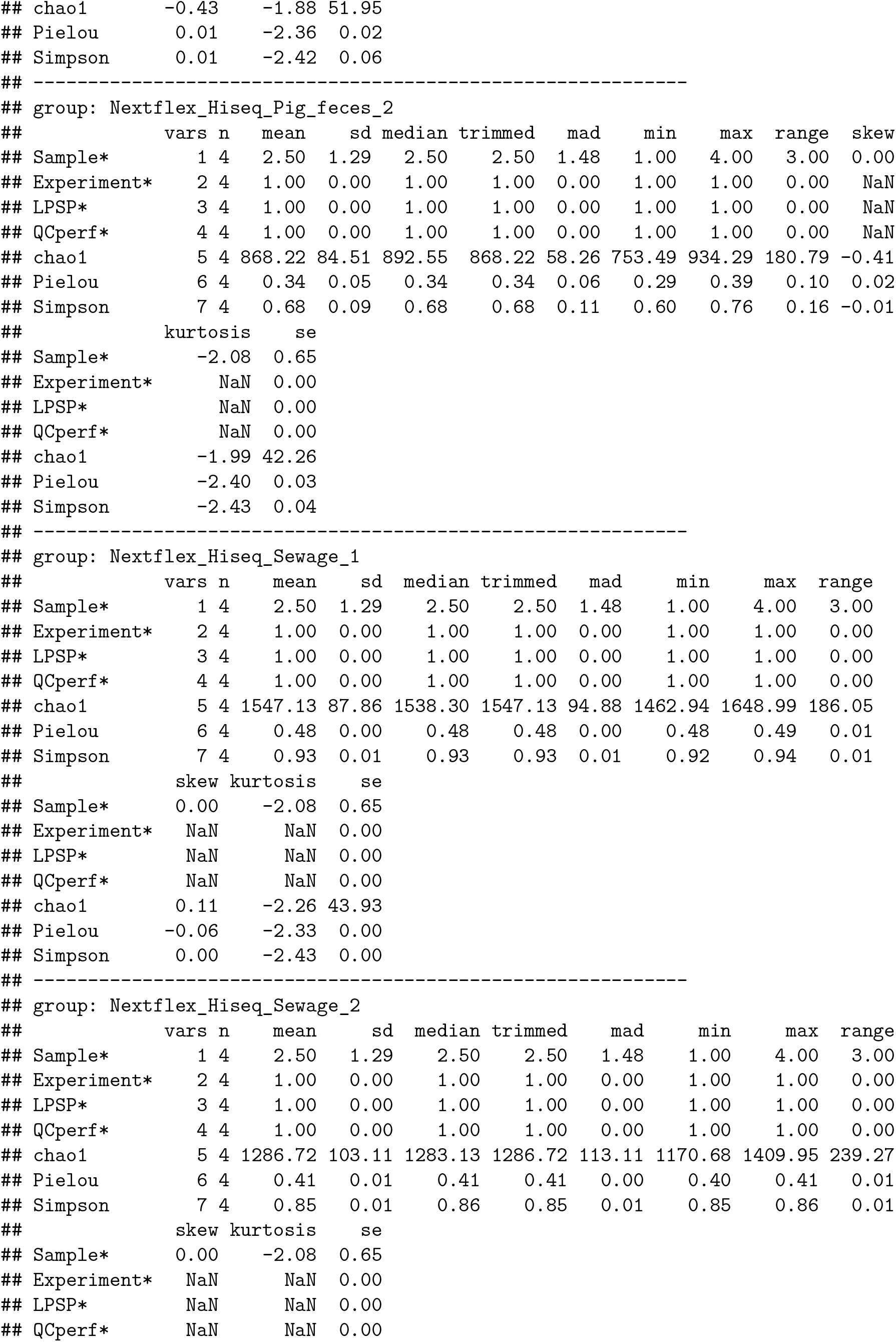
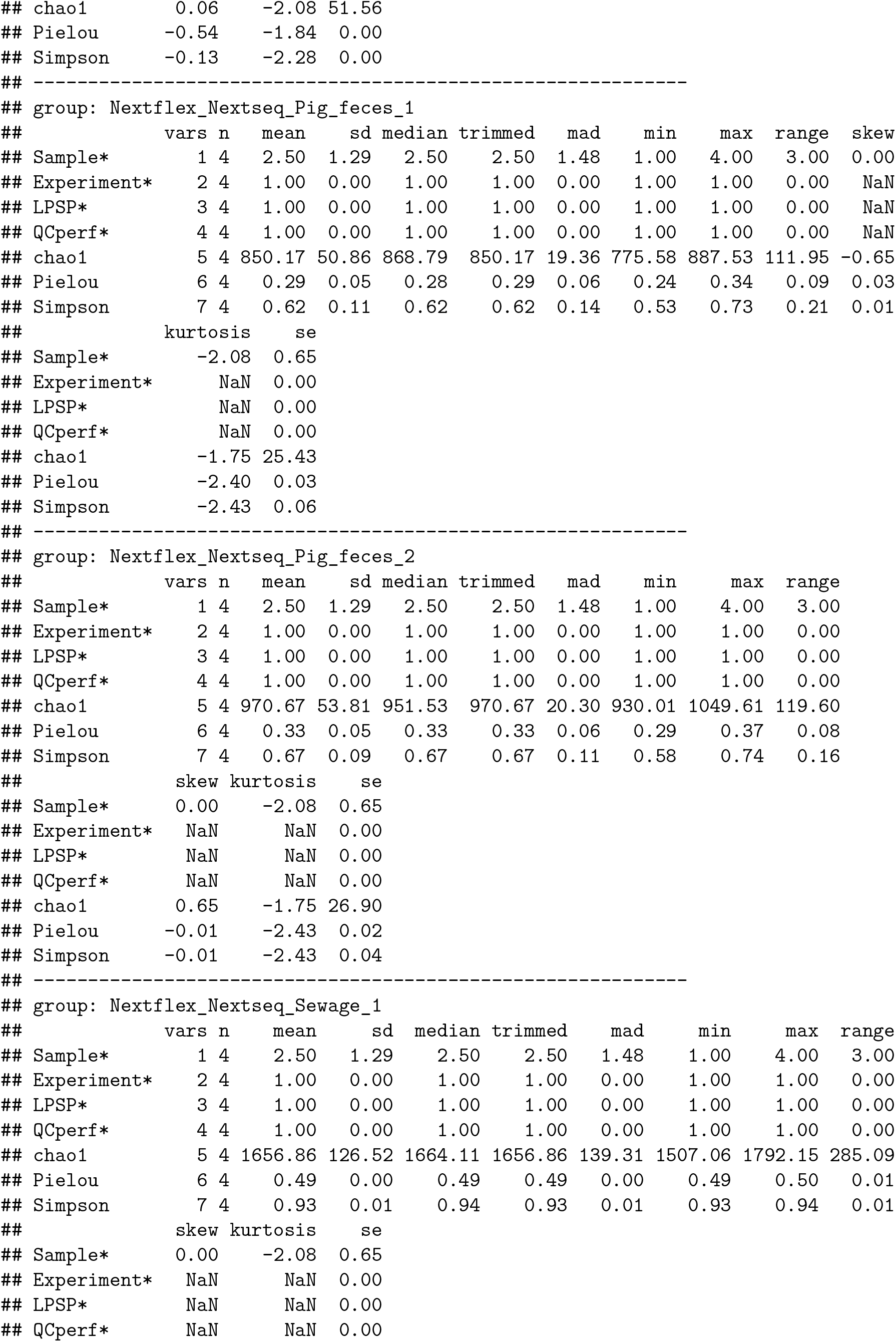
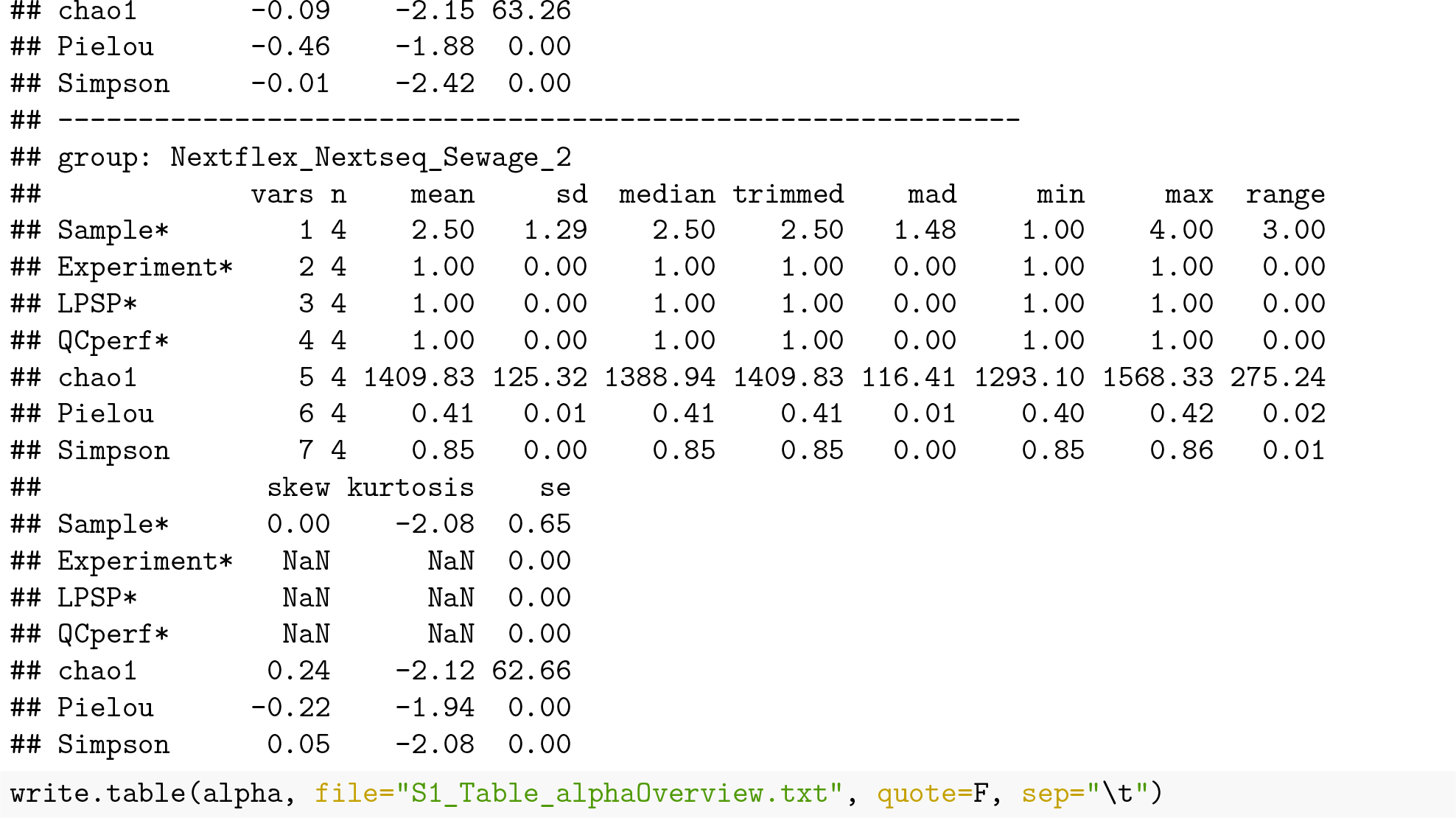

#### PCA

All, pig feces both P1 and P2, and sewage both S1 and S2 creating S3_Fig

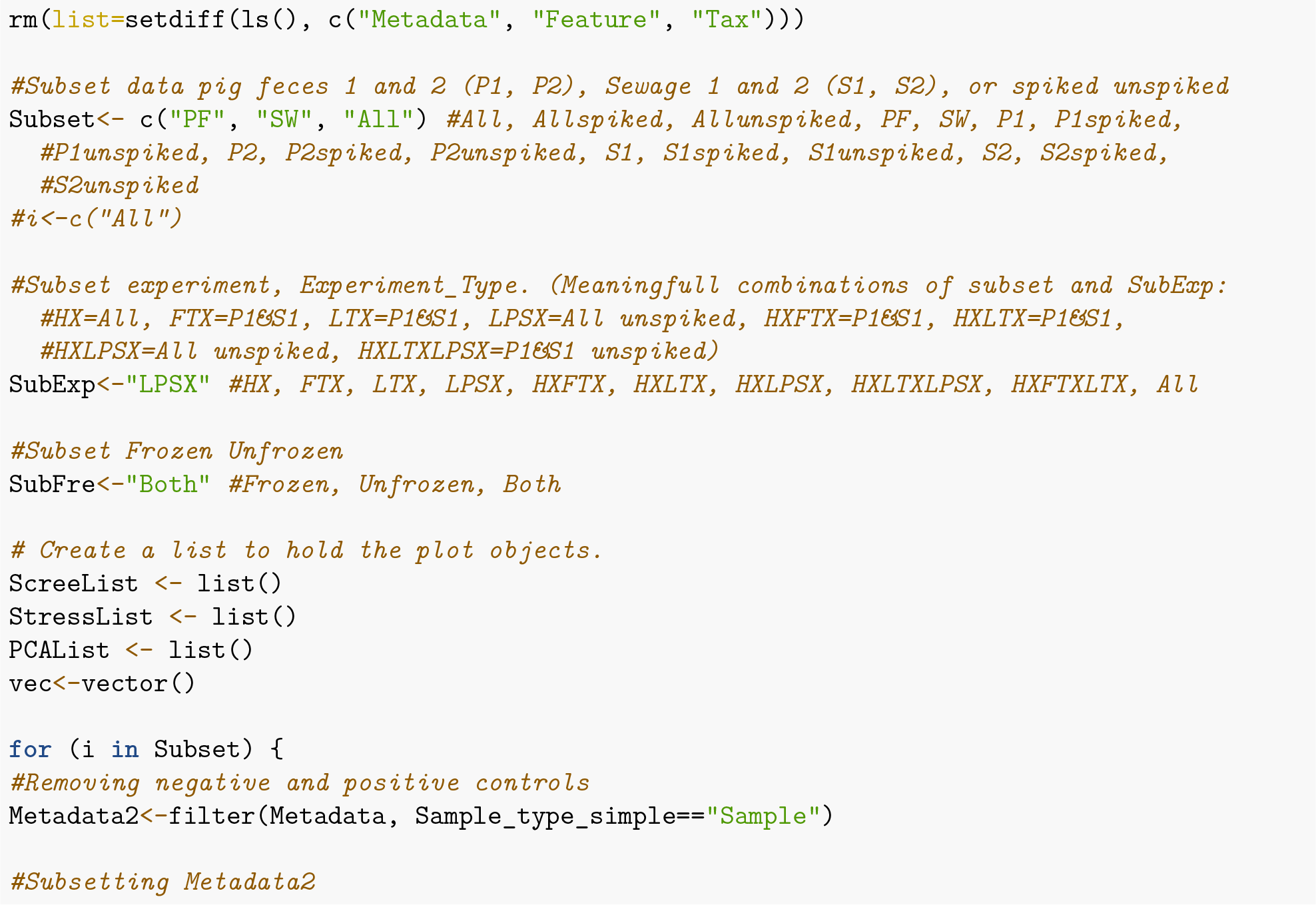
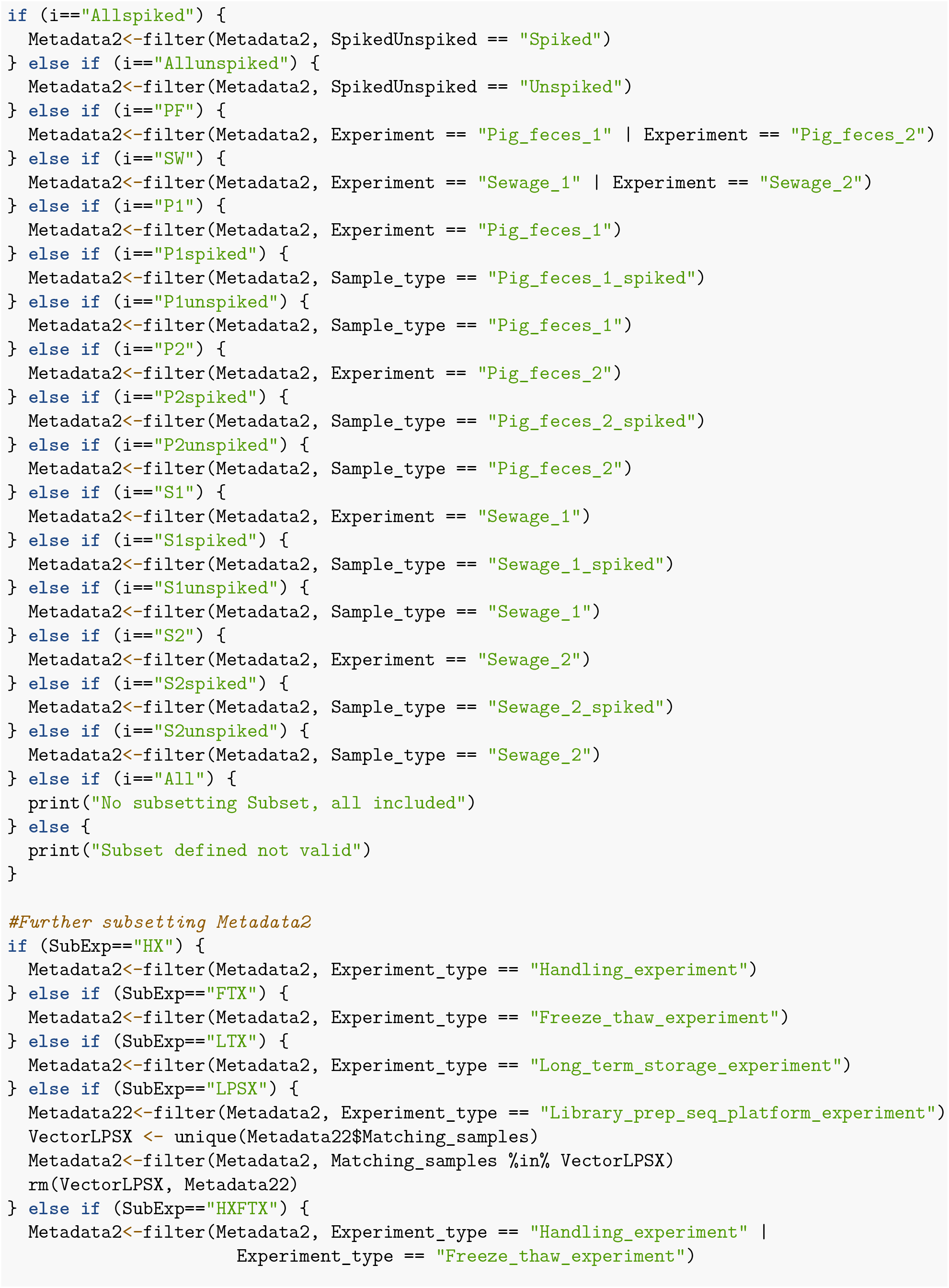
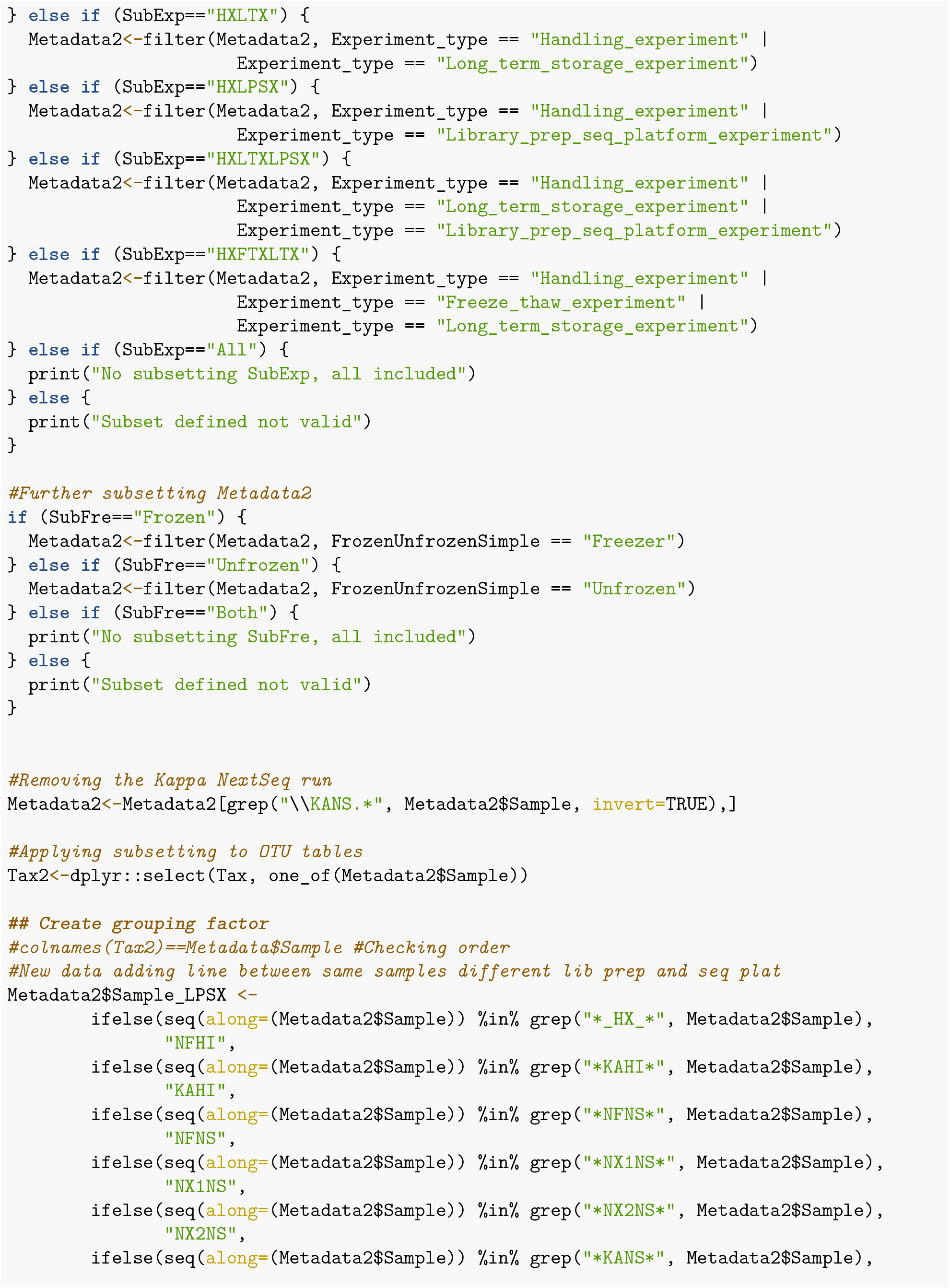
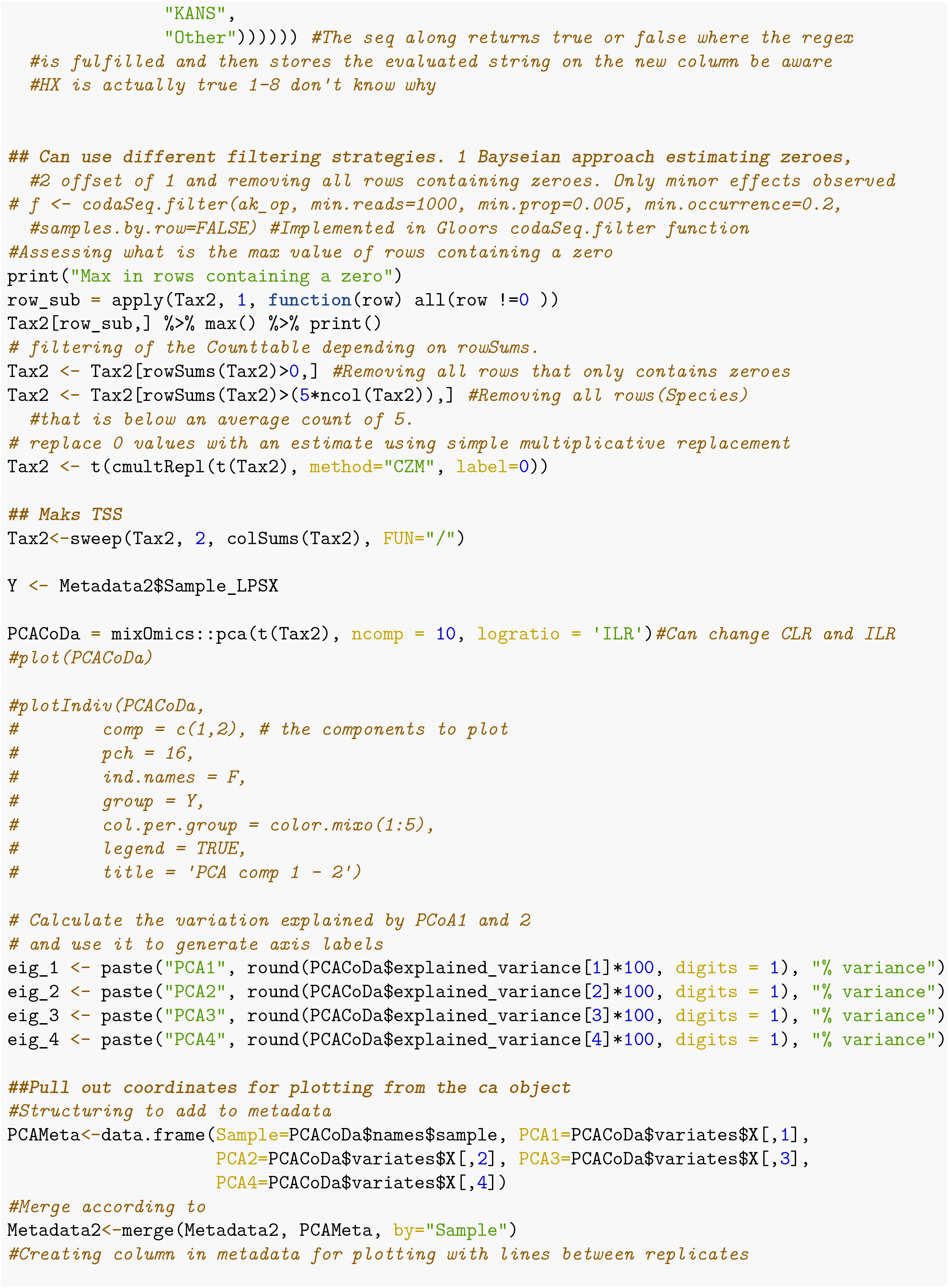
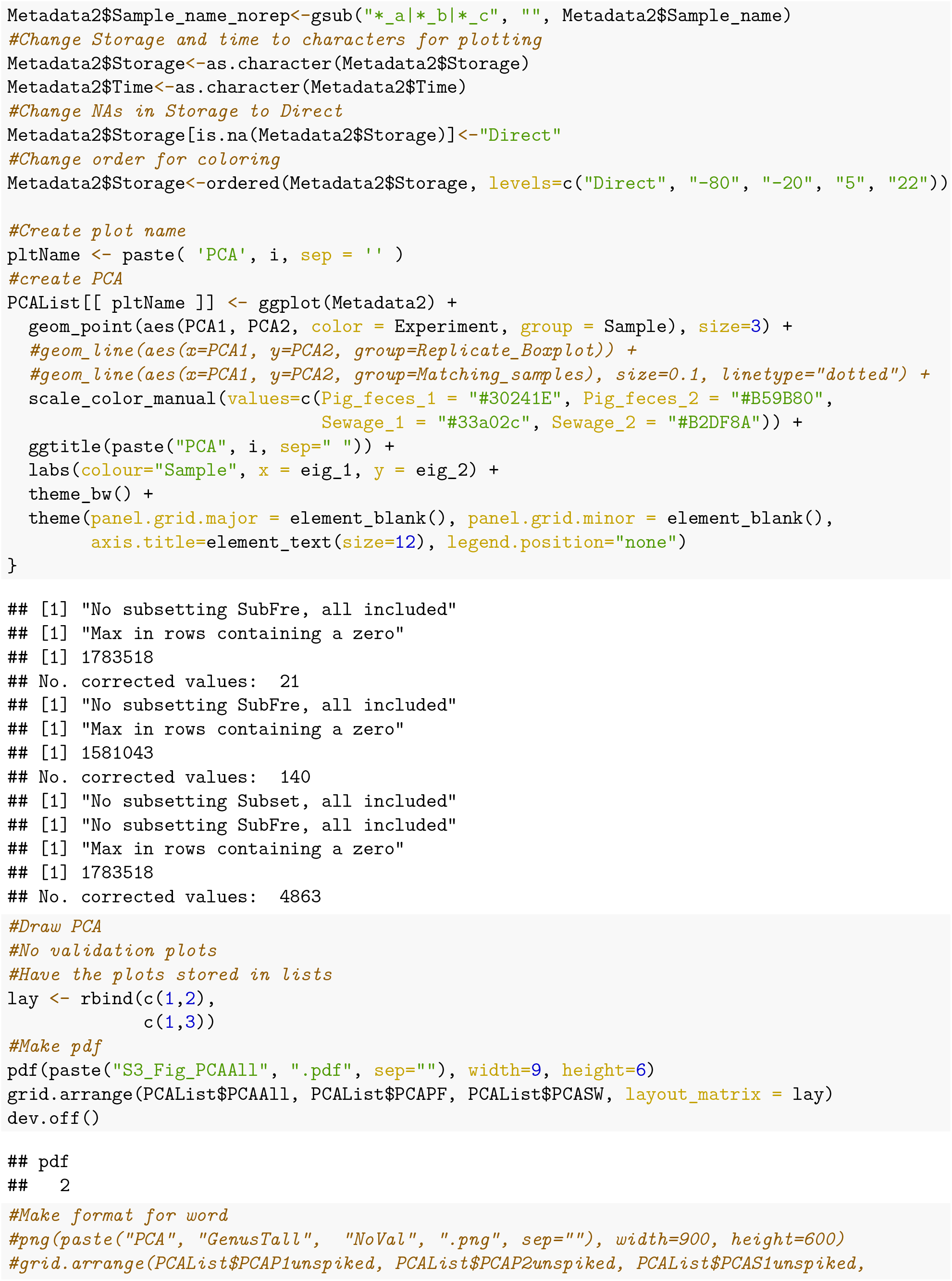
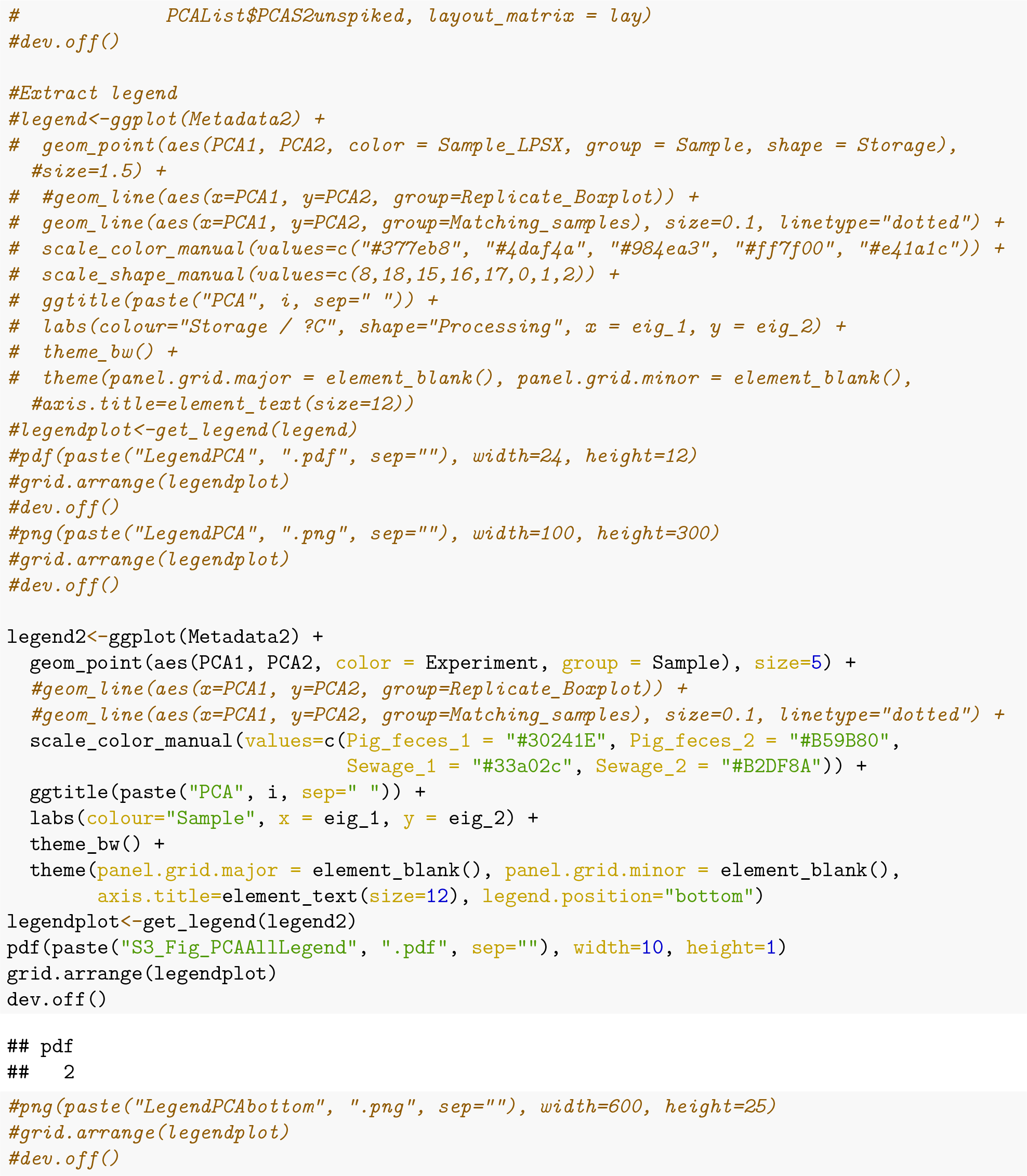

#### PCA Fig_2

Subset to P1, P2, S1 and S2 creating Fig_2

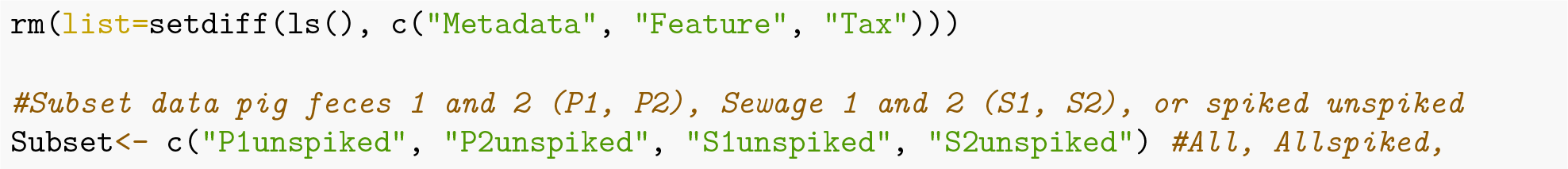
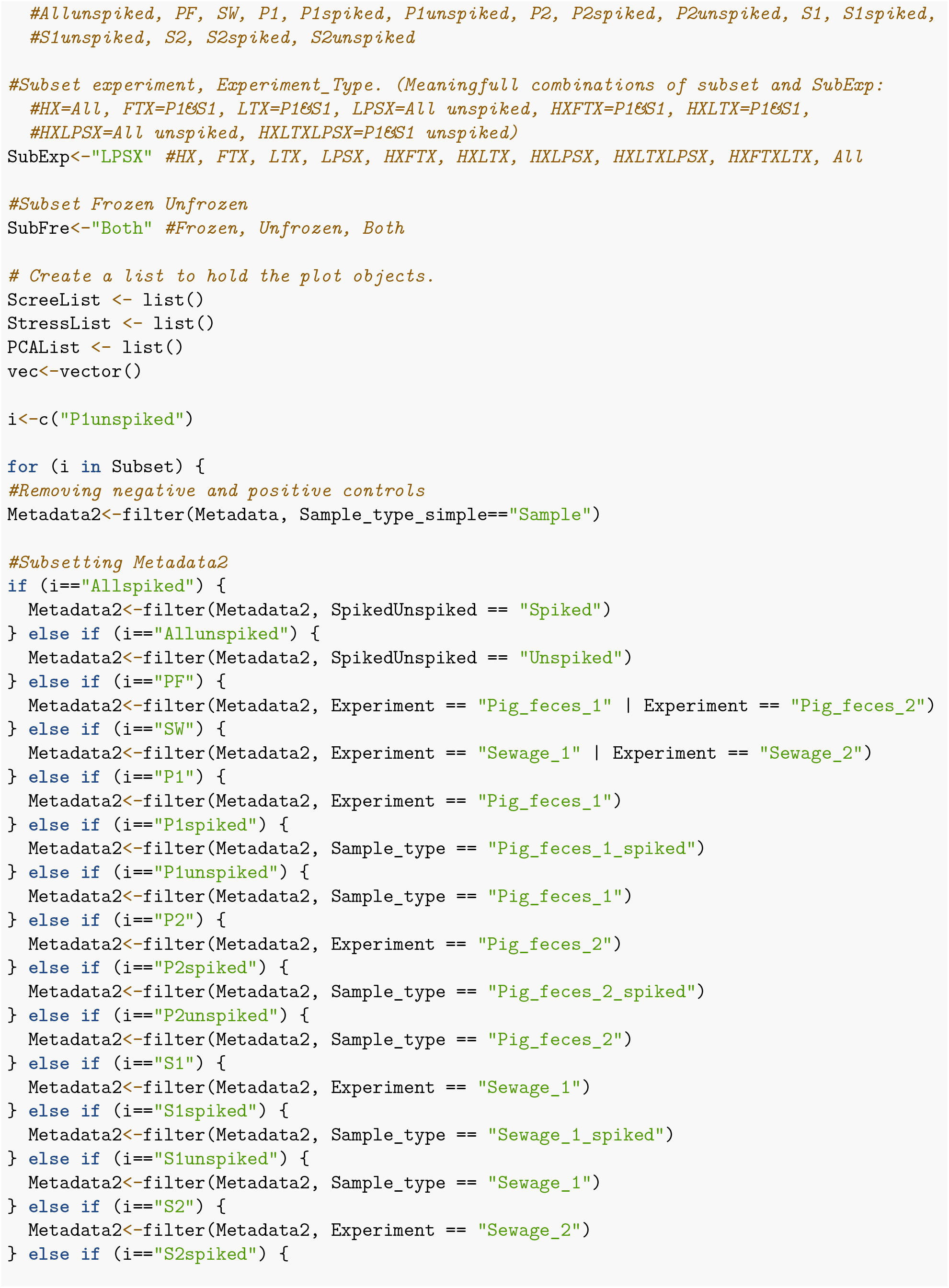
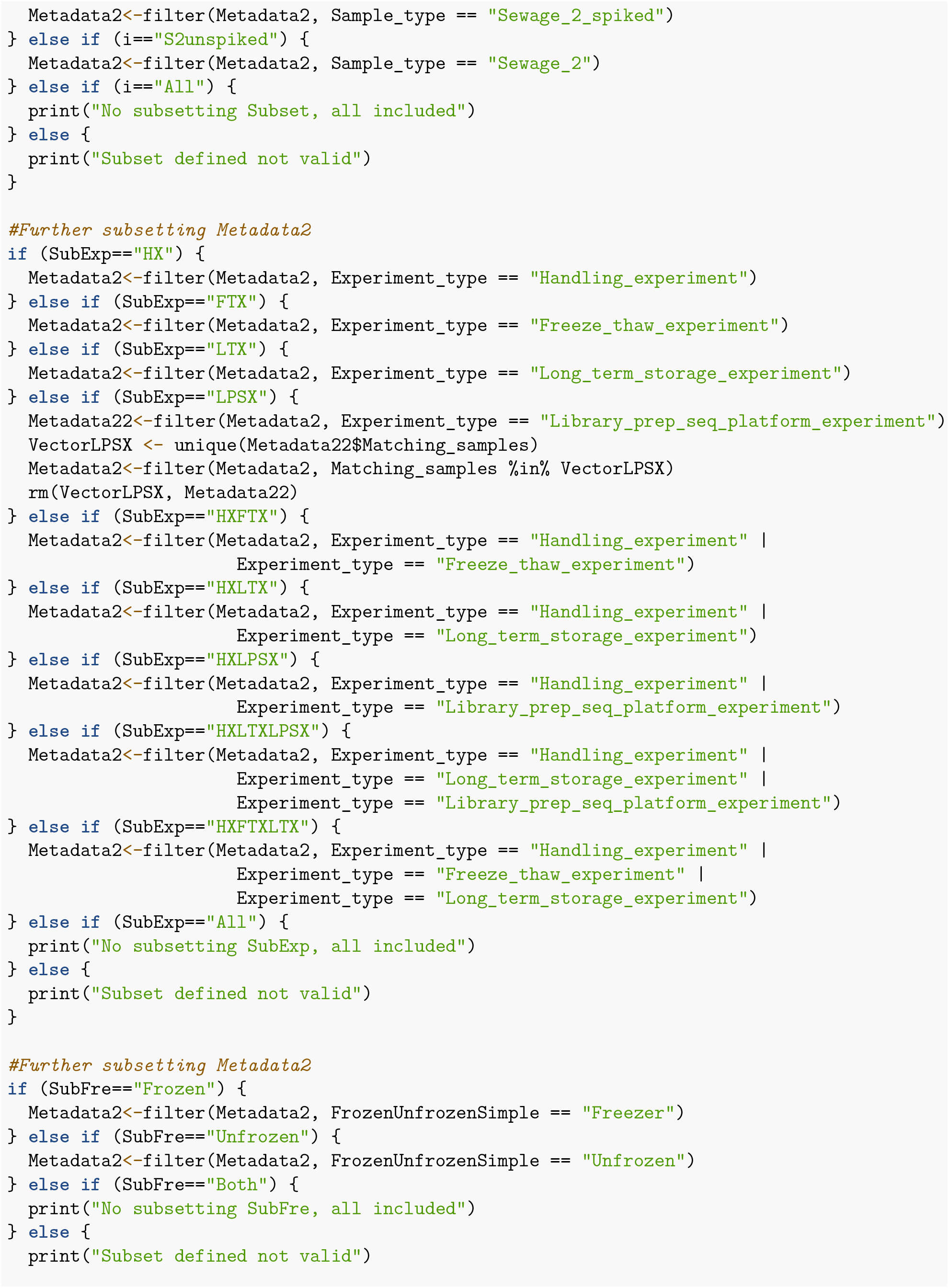
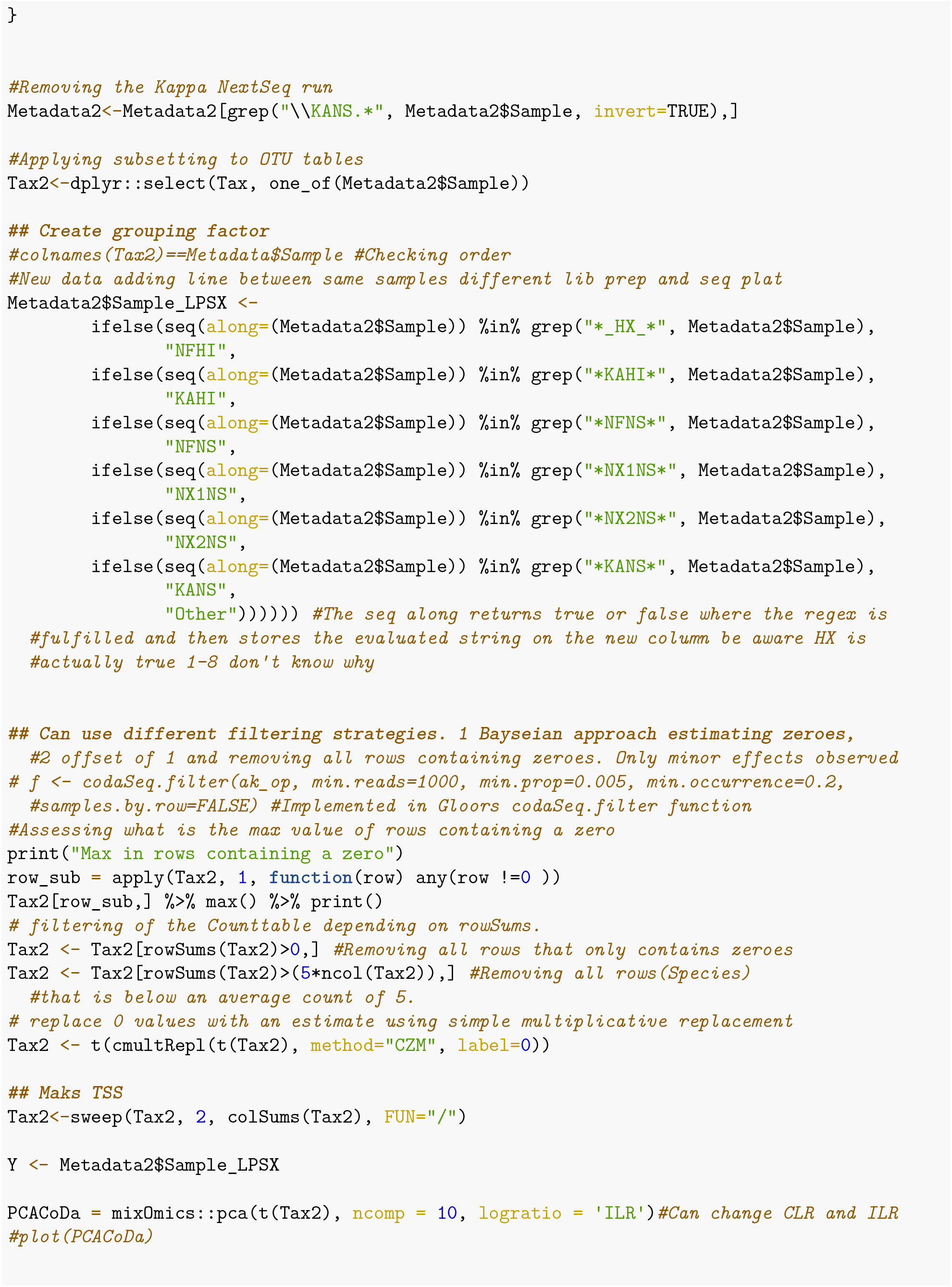
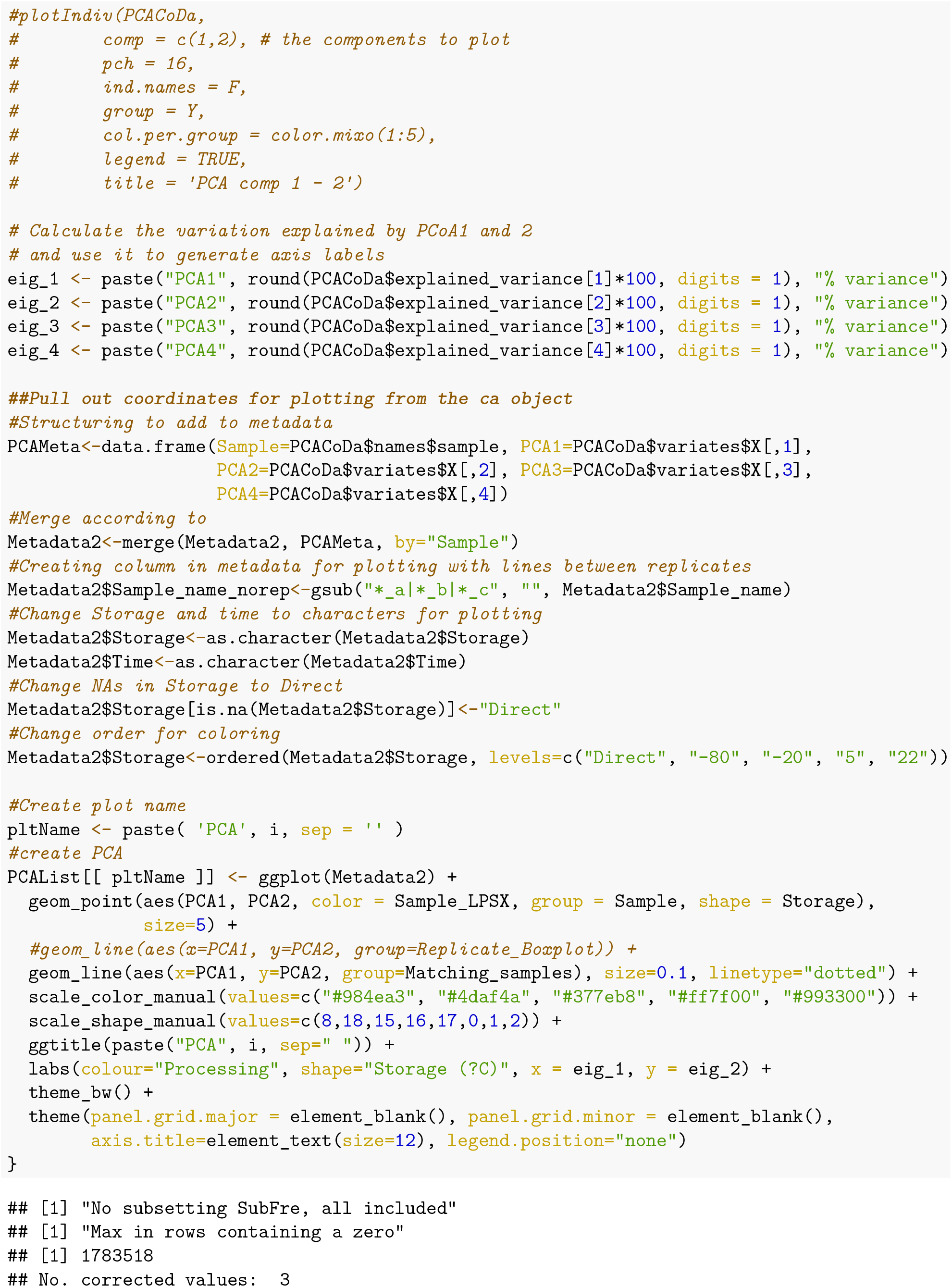
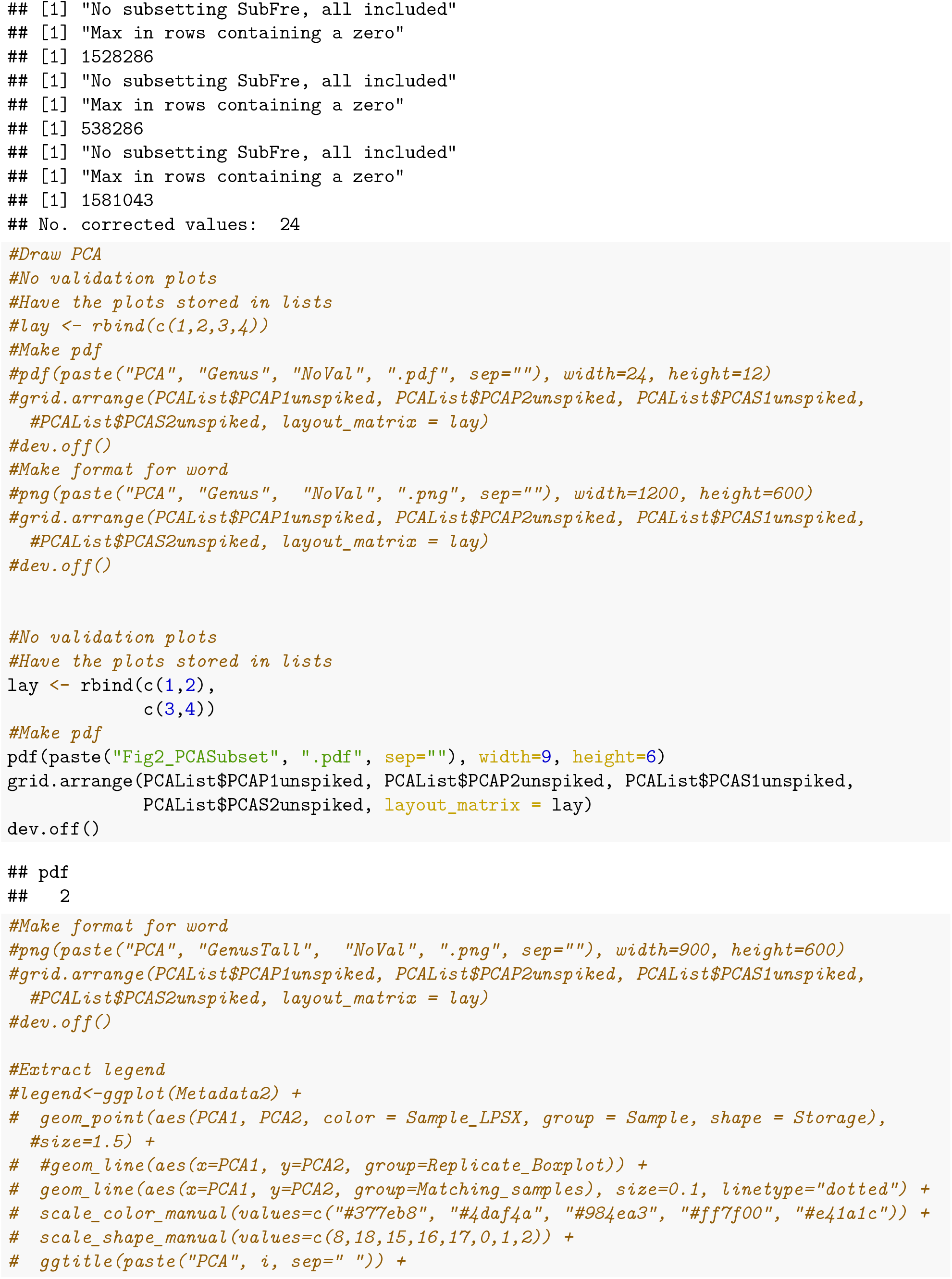
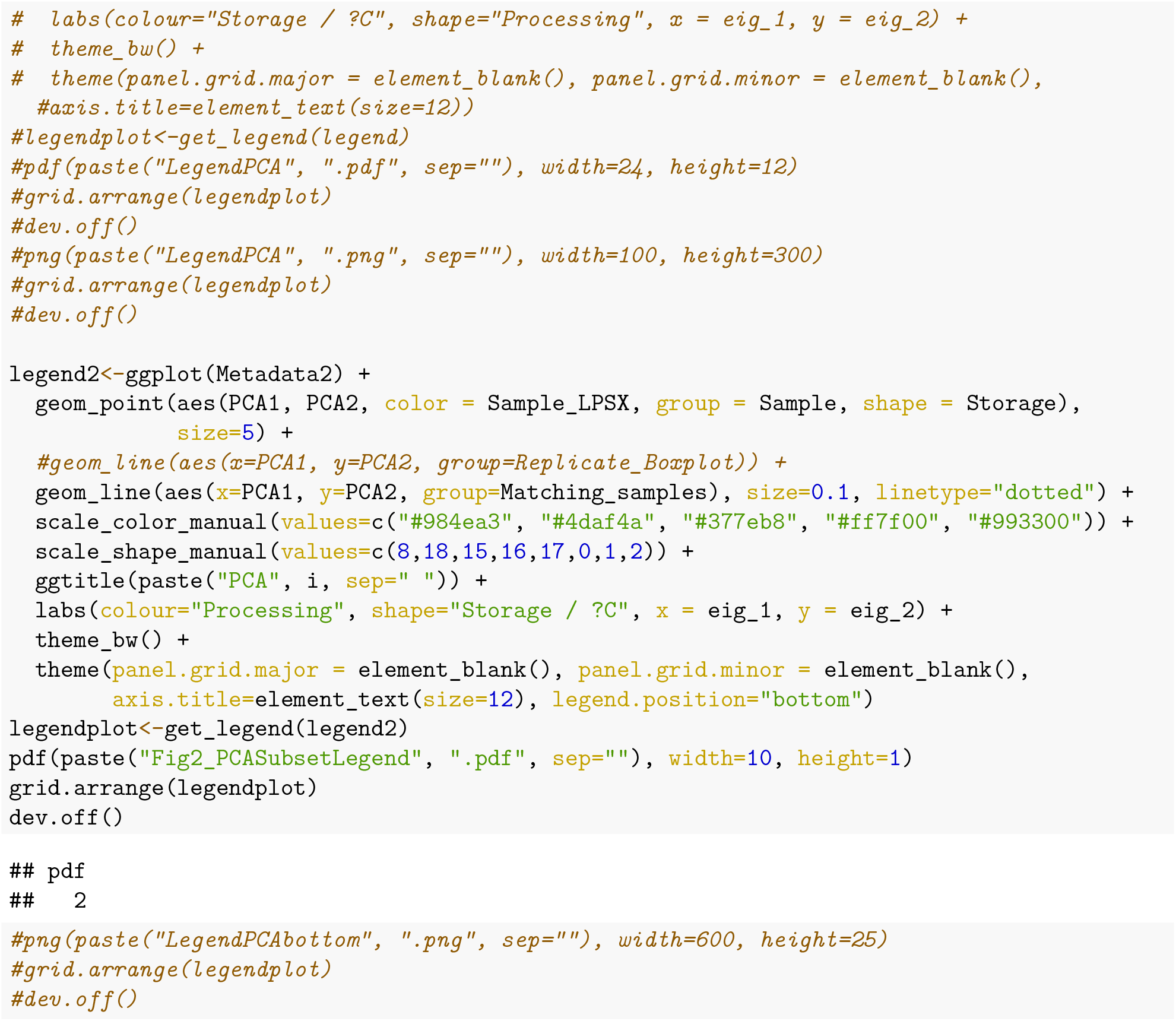

#### Heatmaps

Pig feces both P1 and P2, and sewage both S1 and S2 creating Fig_3. Have added a heatmap of the negative controls S5_Fig

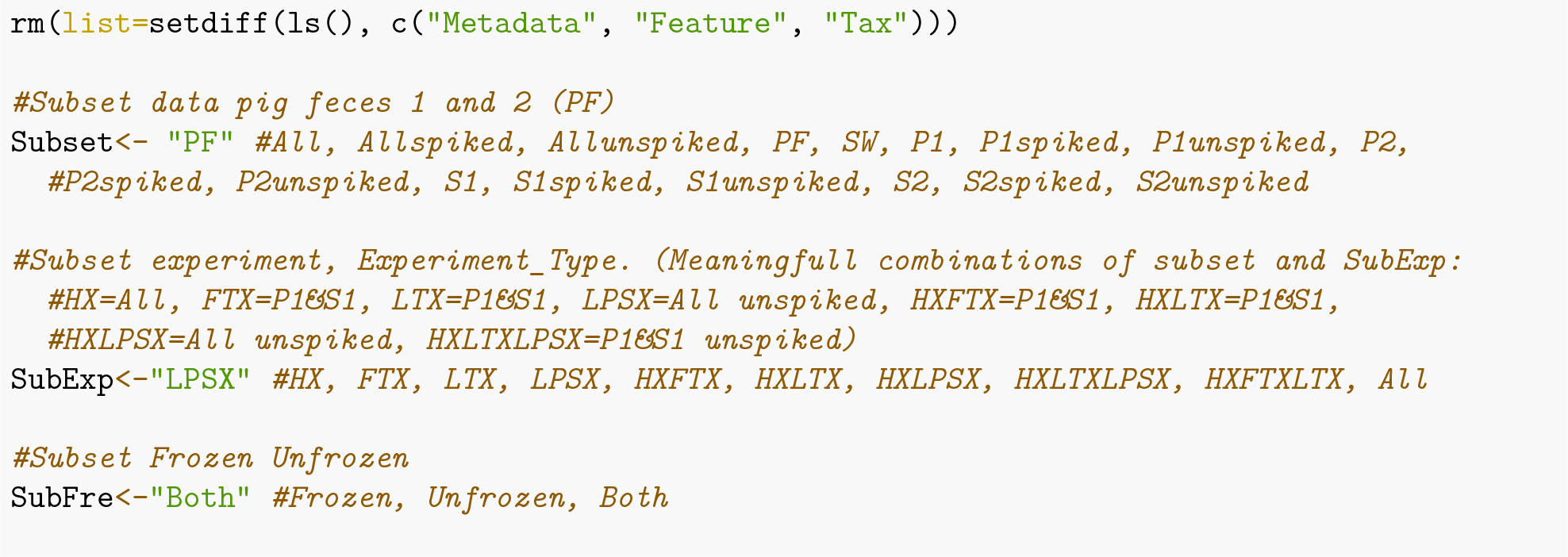
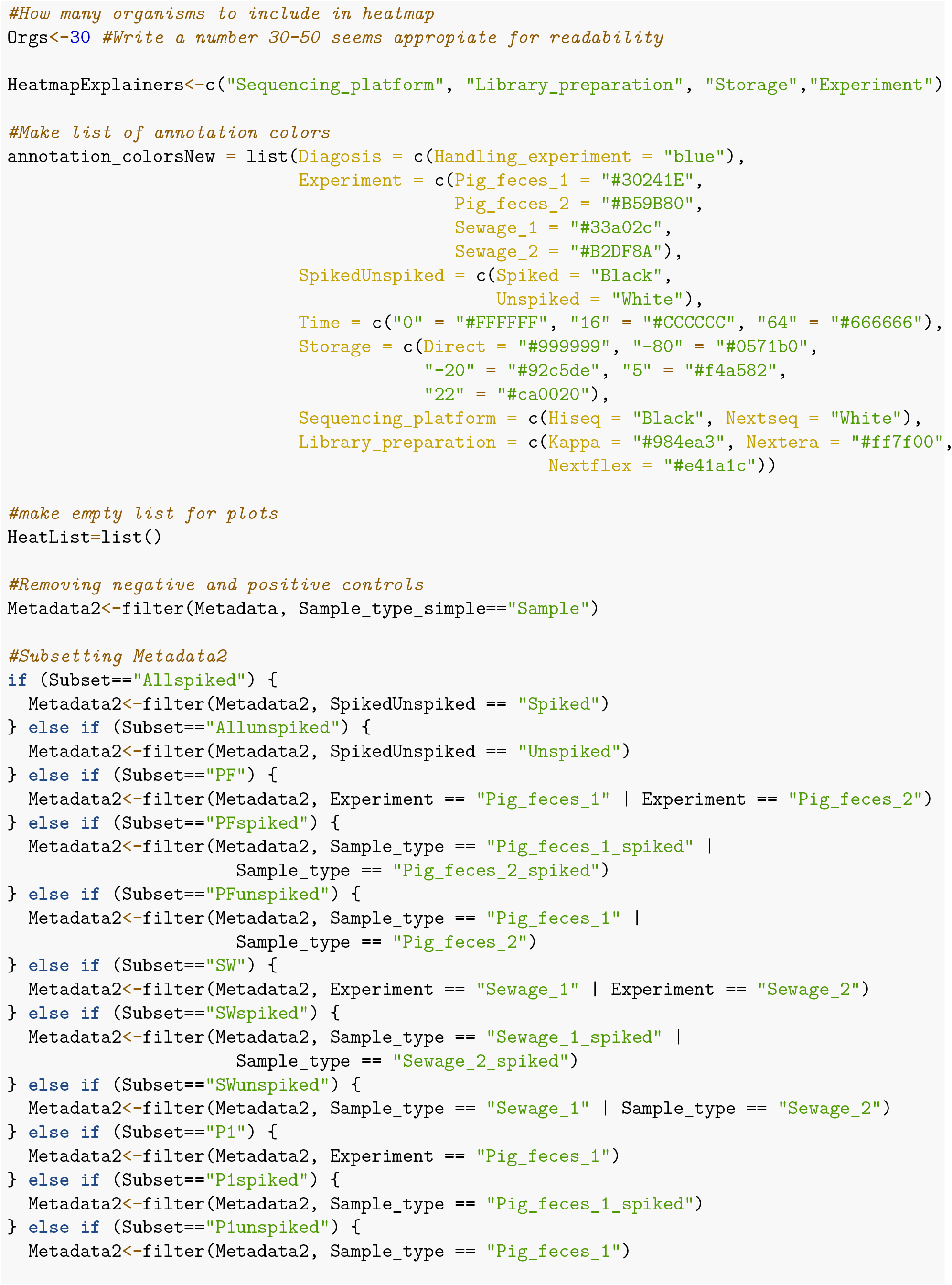
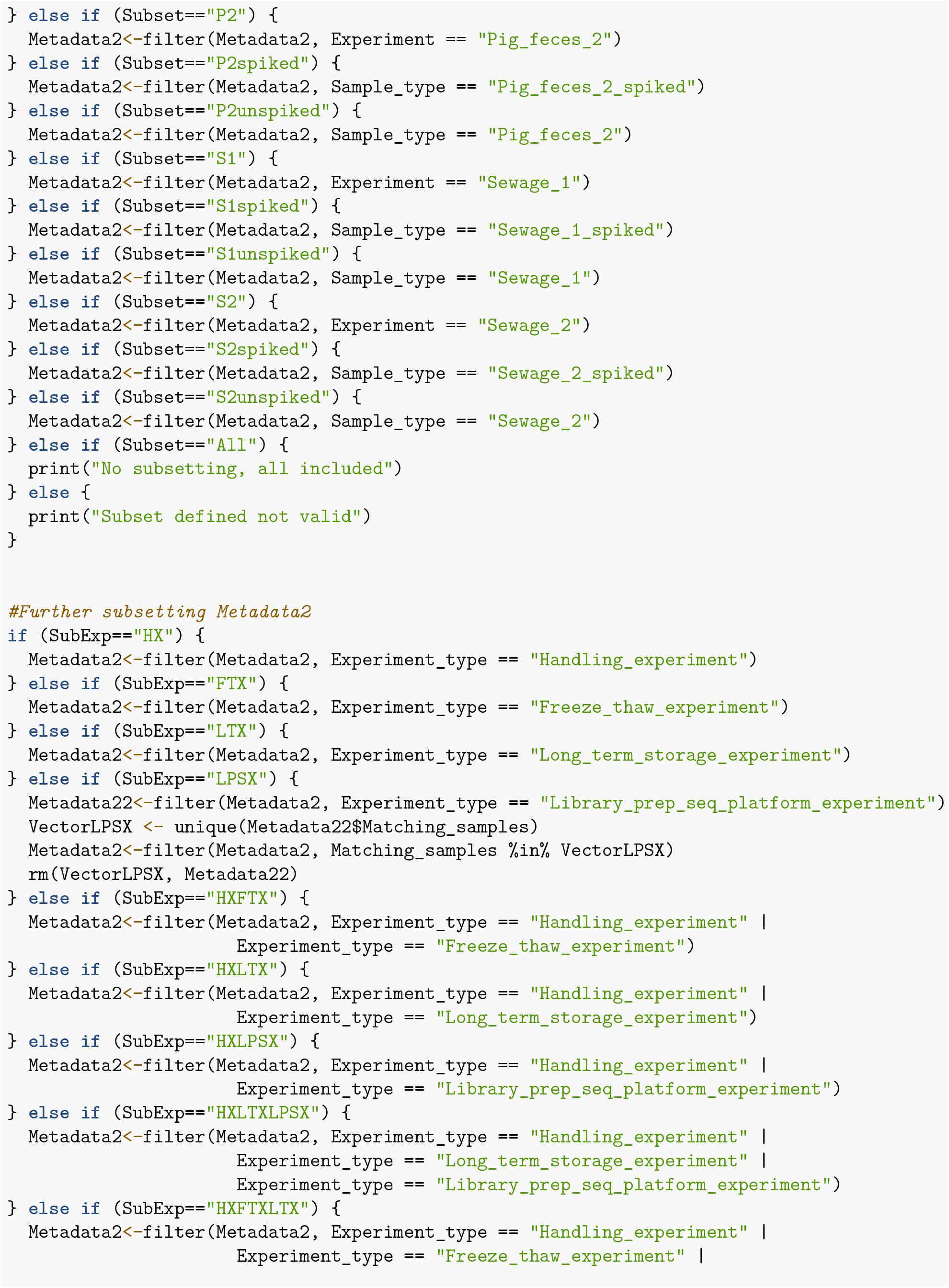
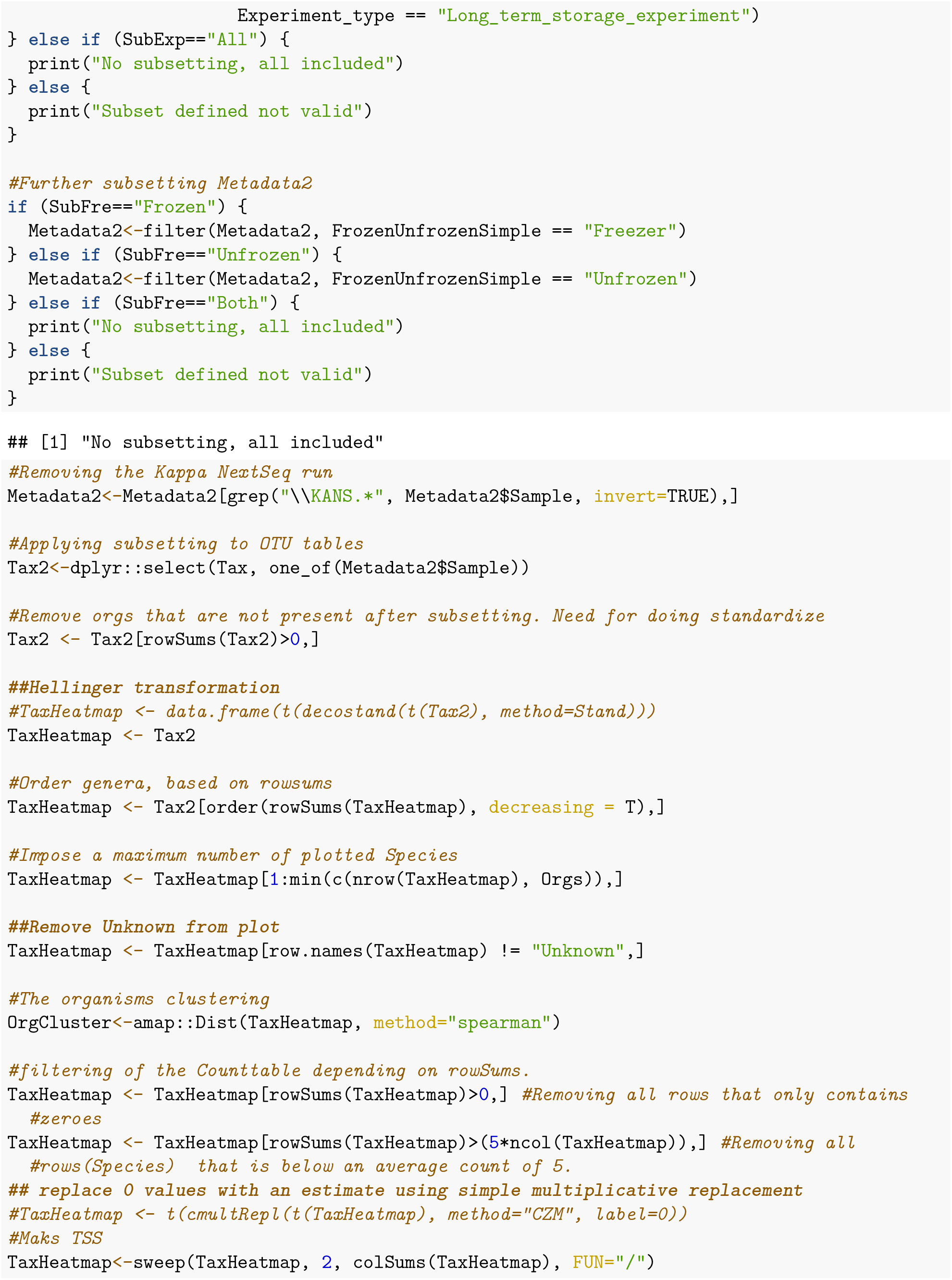
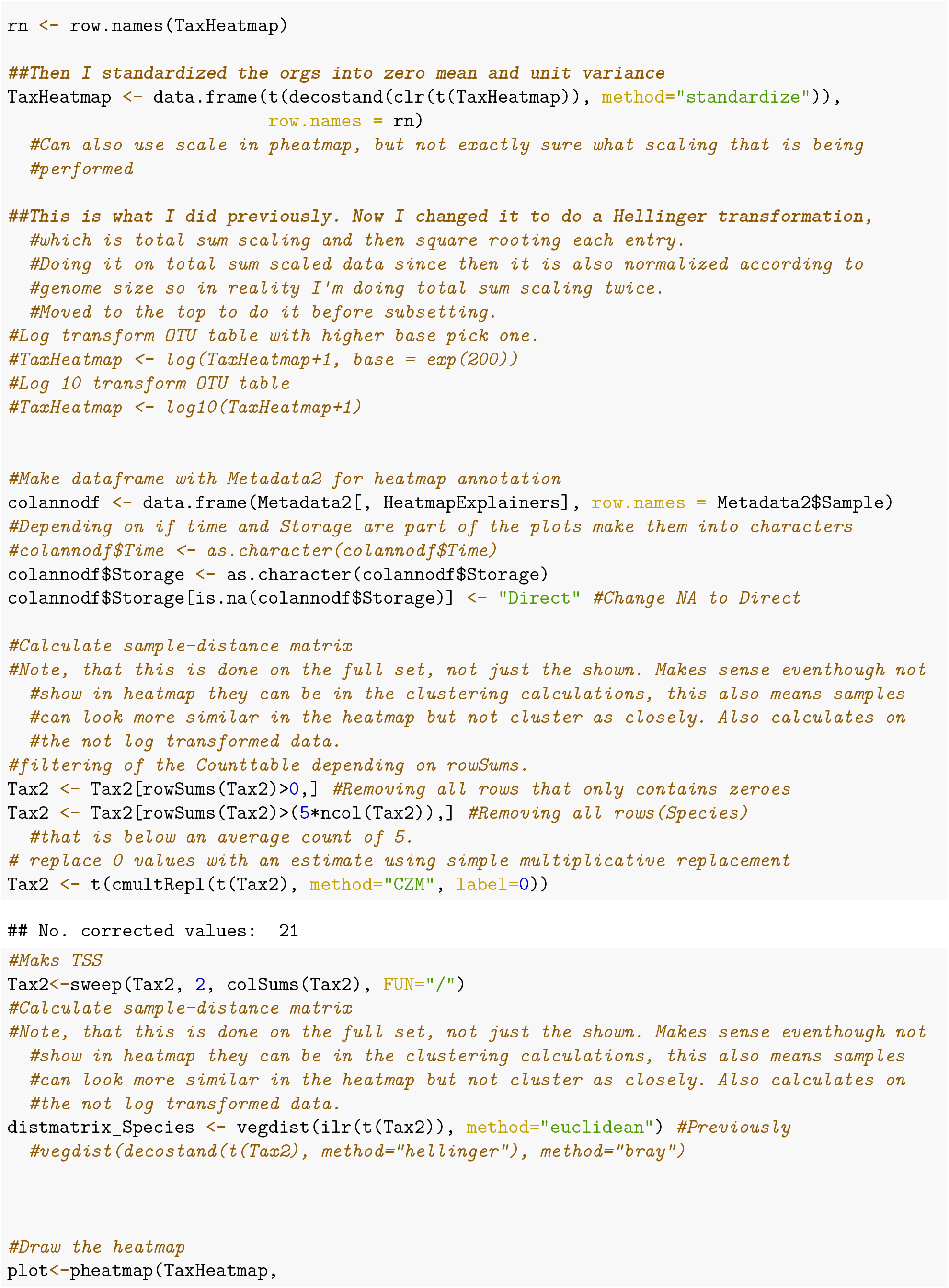
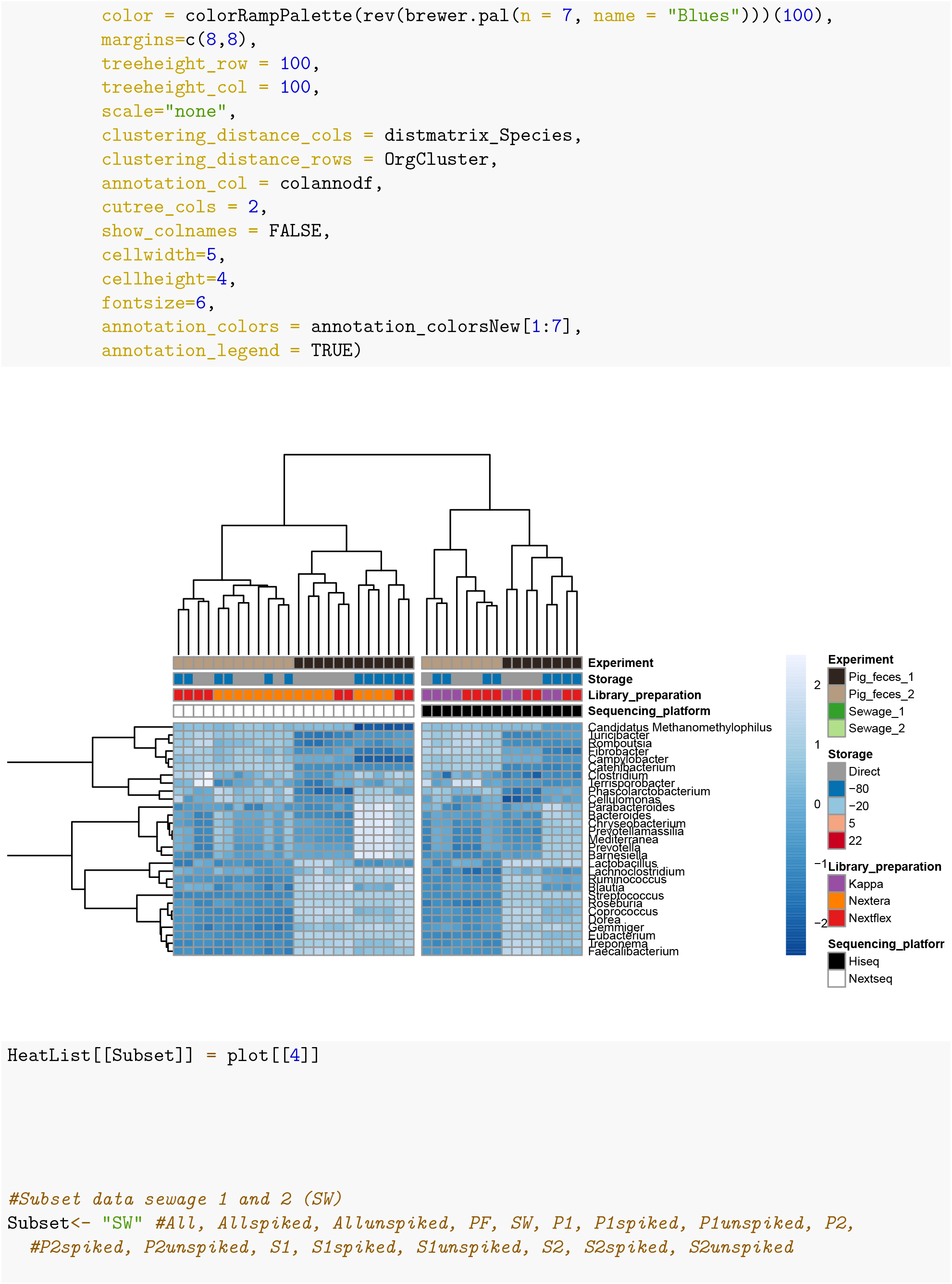
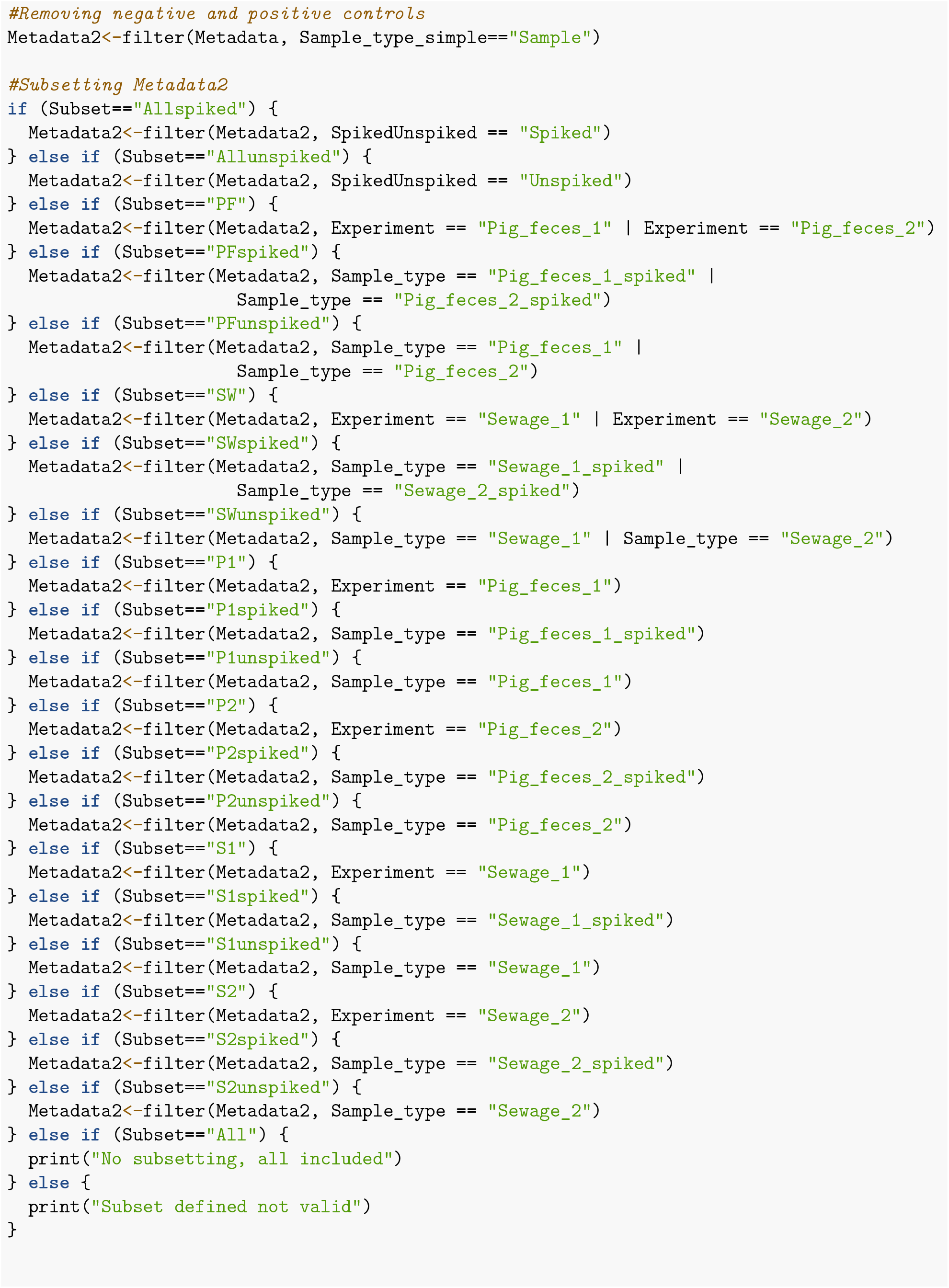
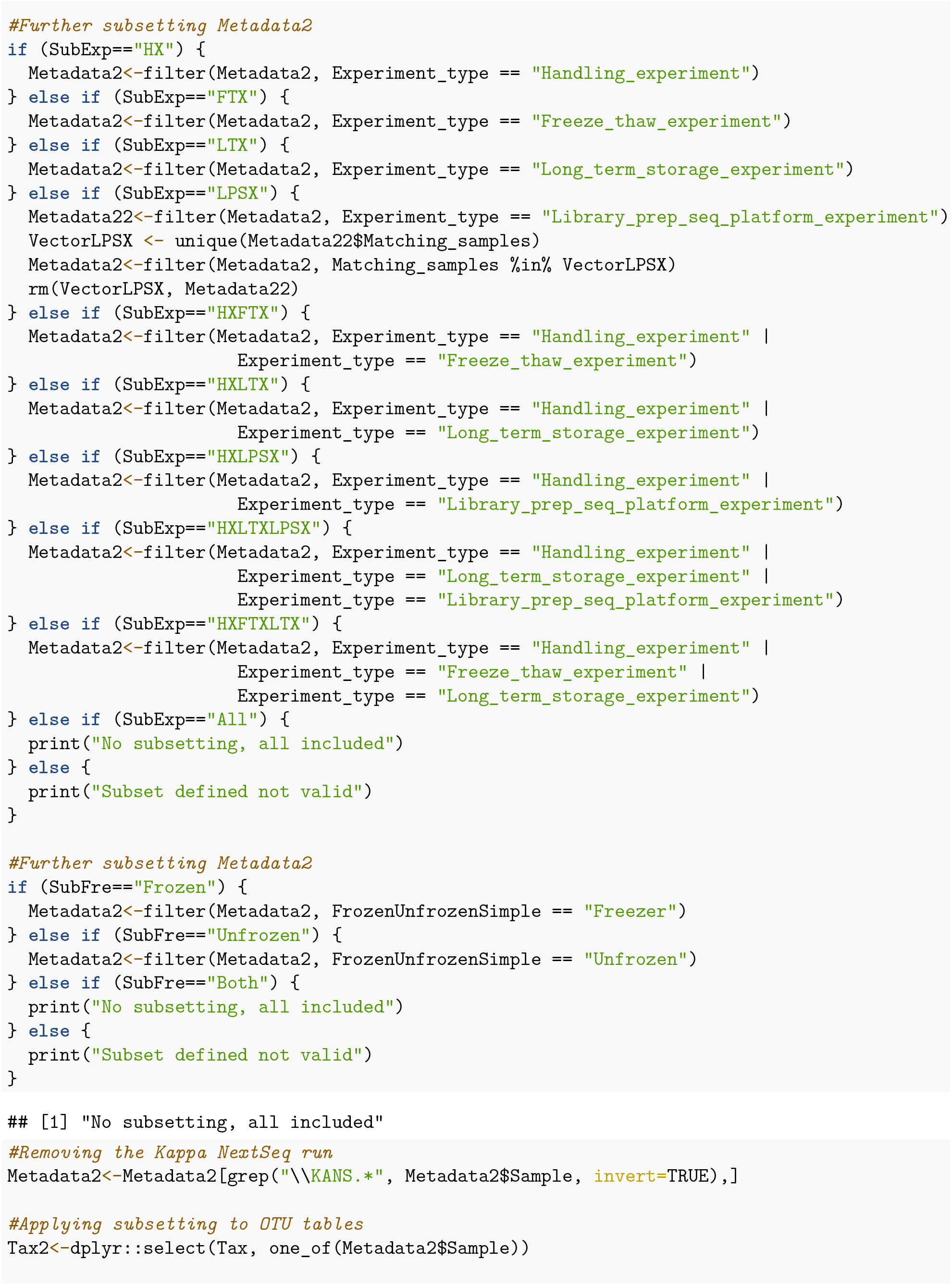
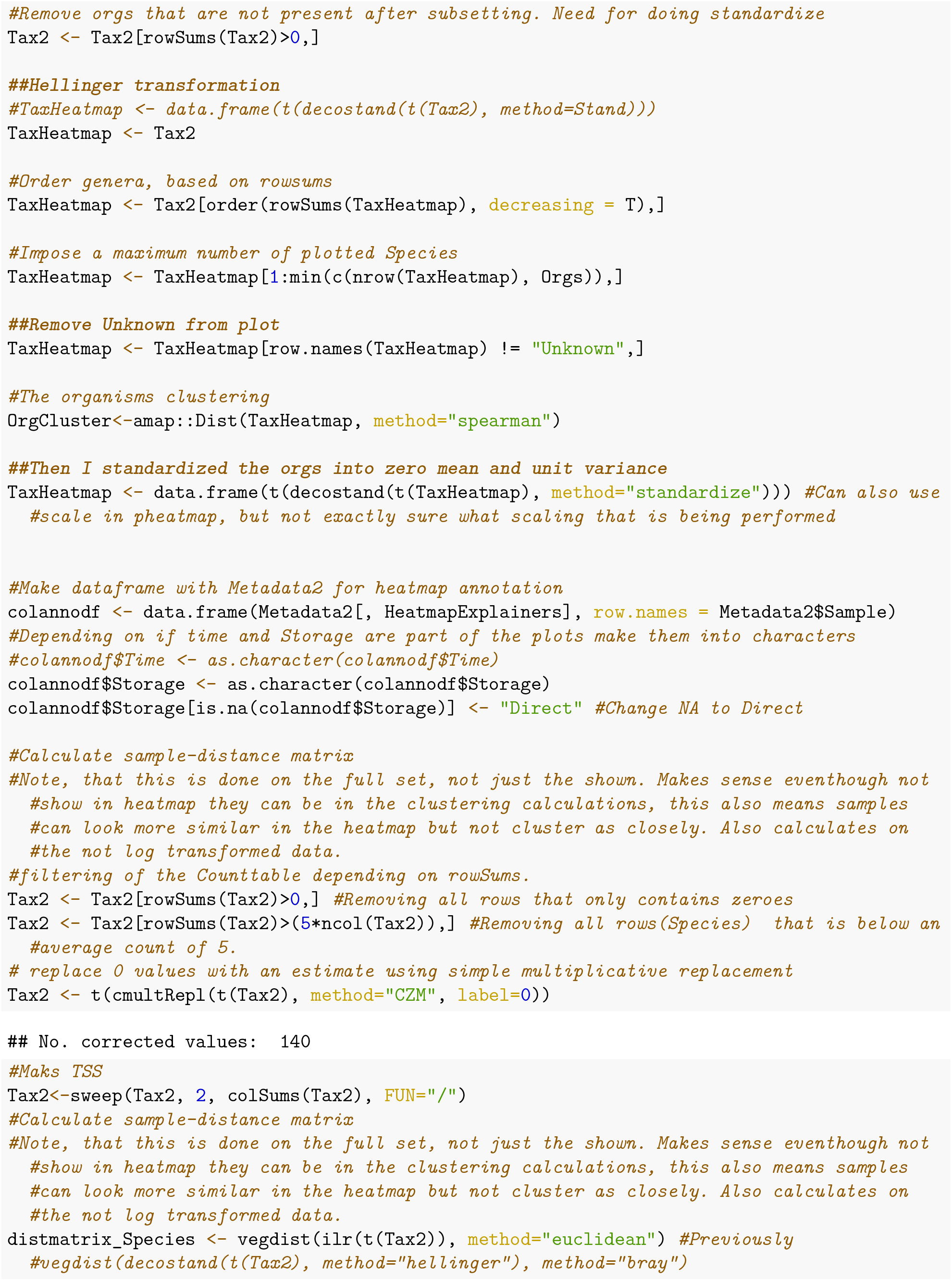
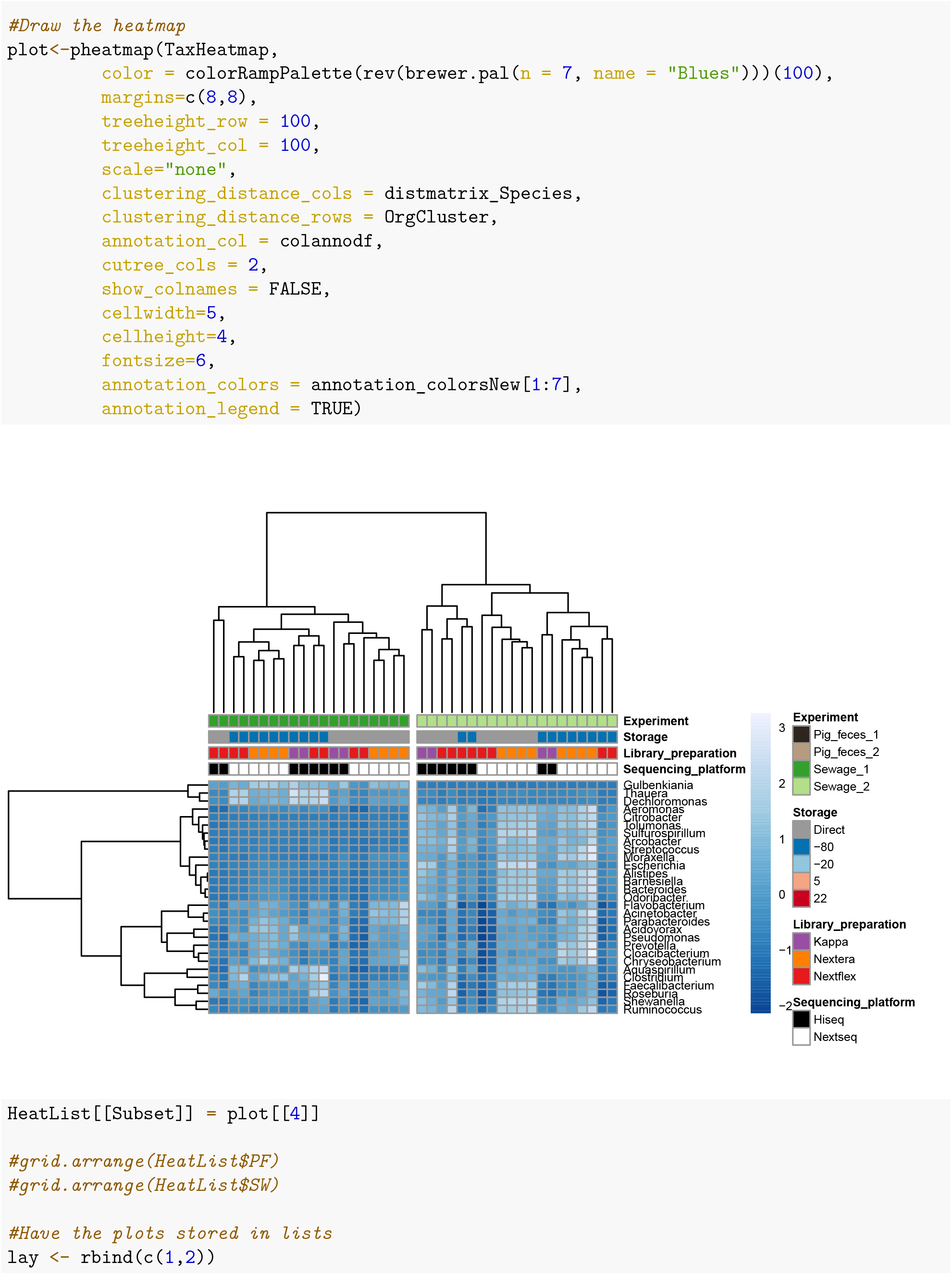
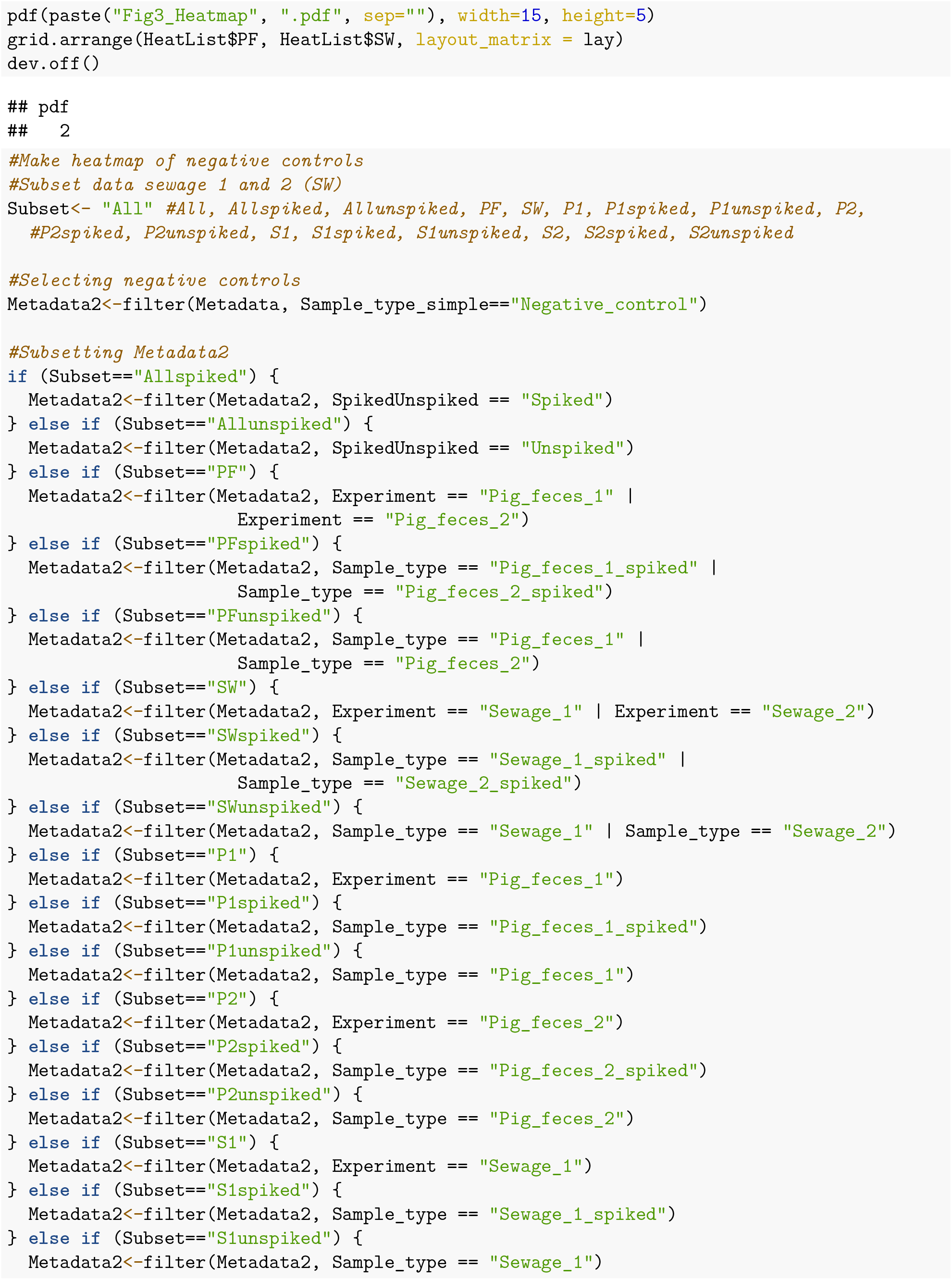
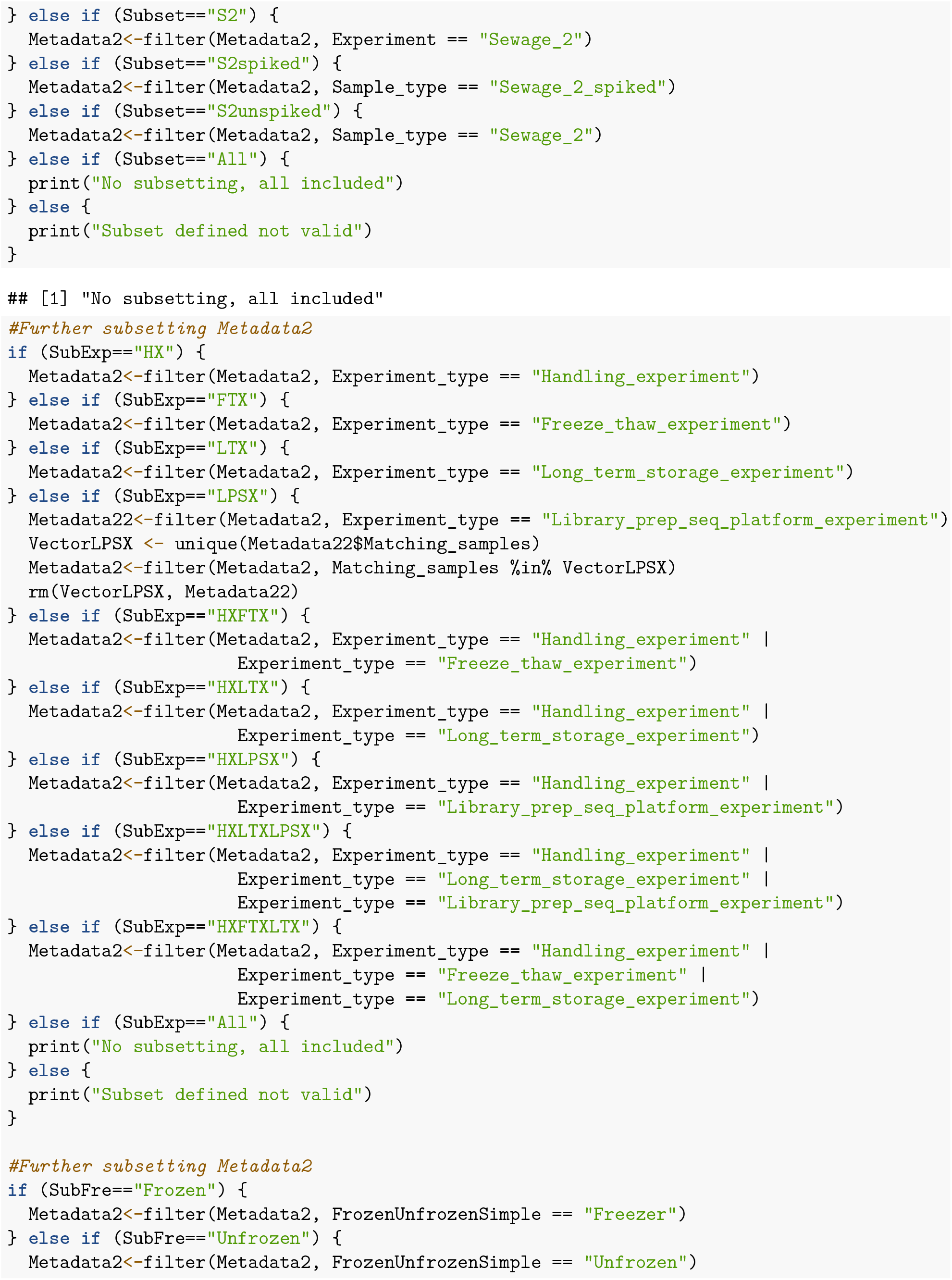
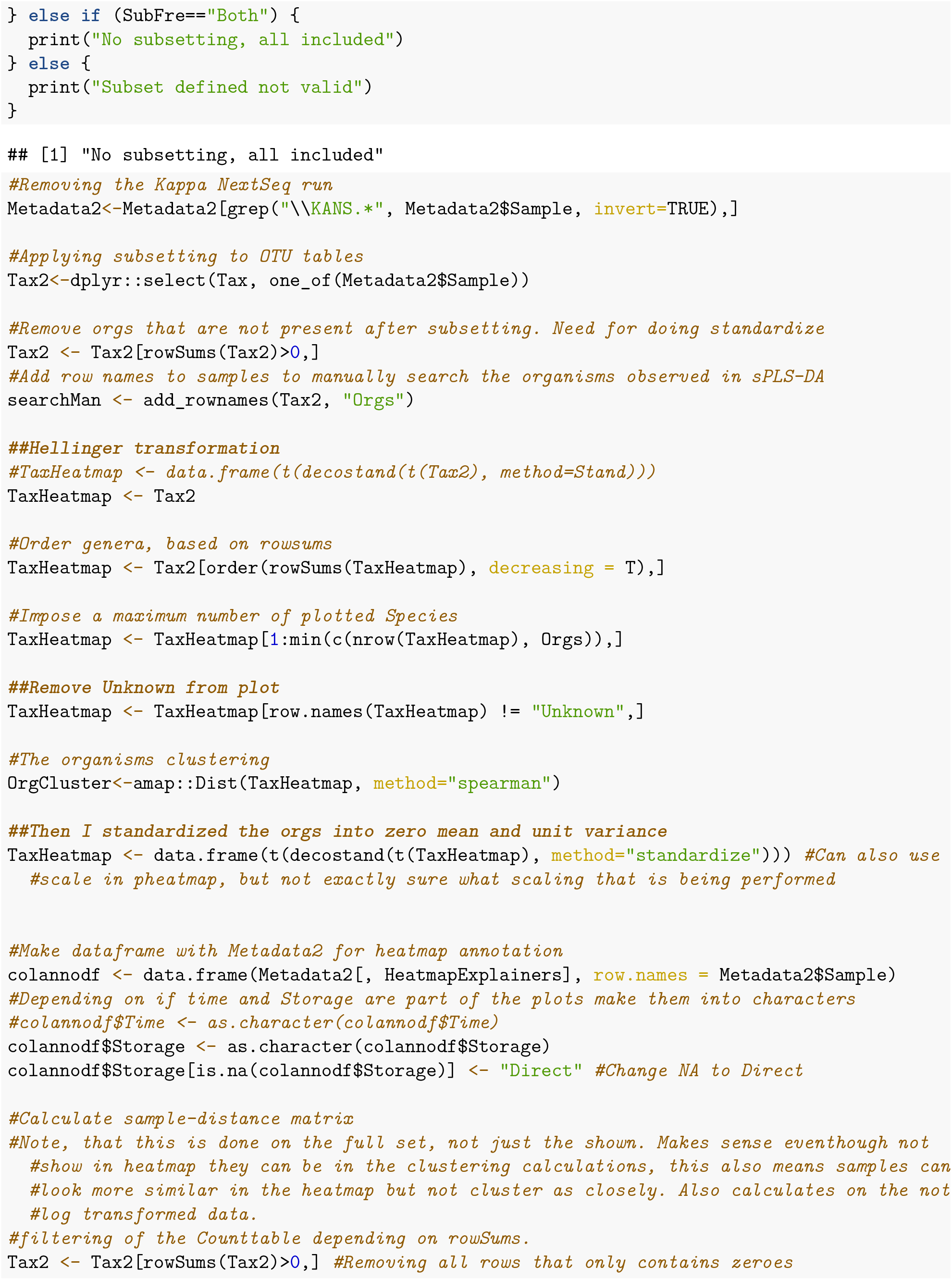
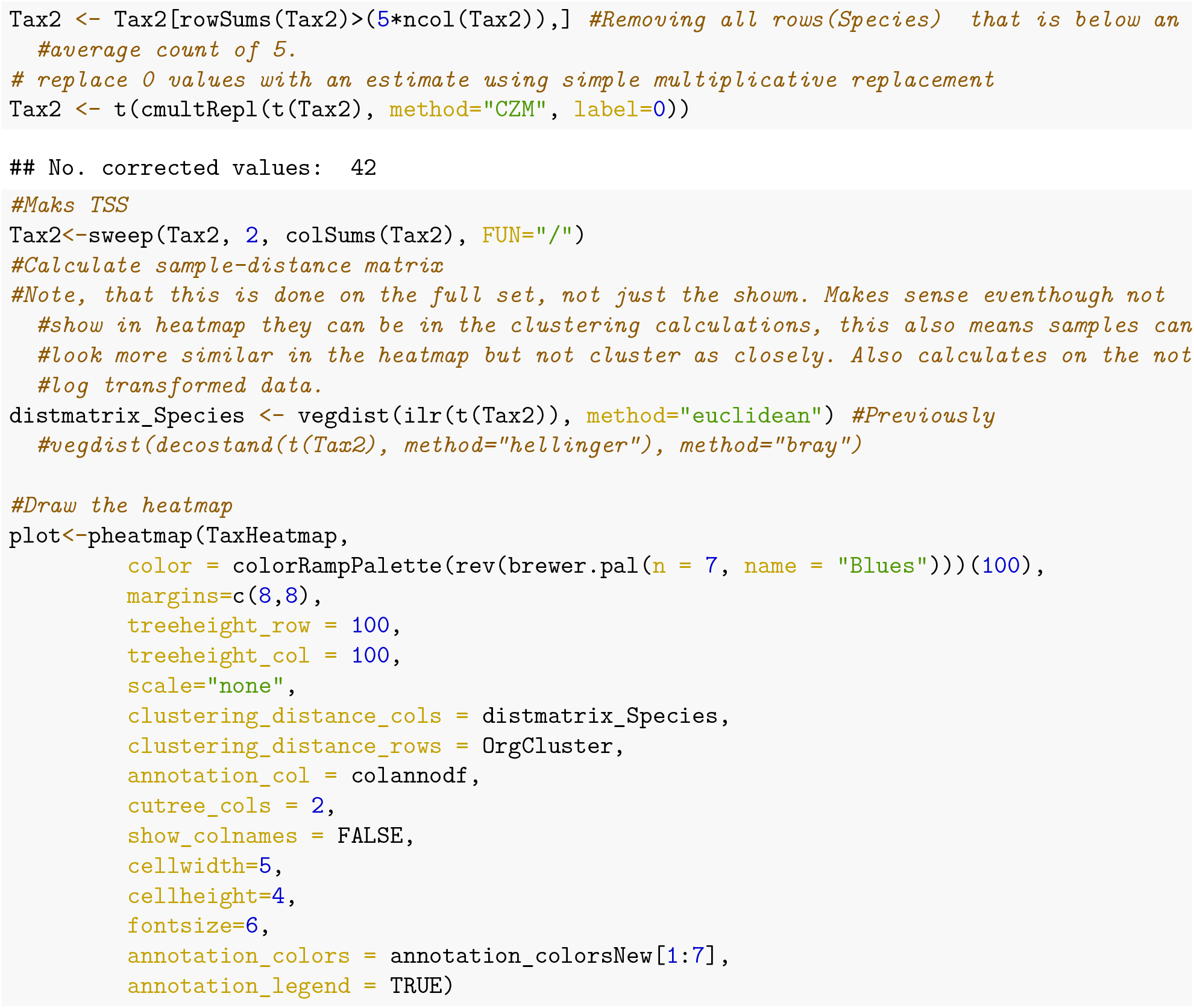
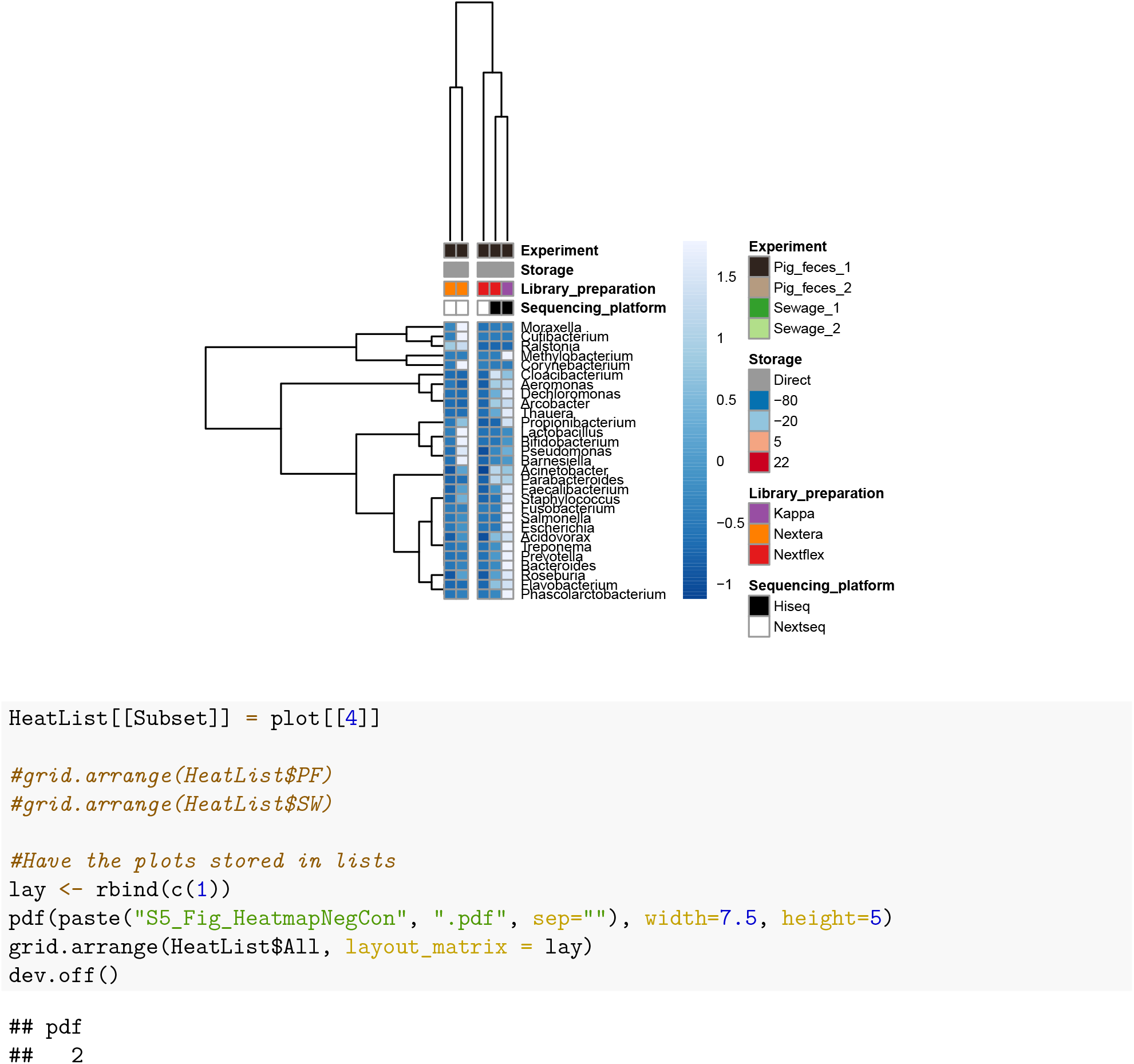

#### Boxplots of distances between samples

Grouped according to different parameters (Sample, Storage, Library preparation, sequencing platform, Replicates) to create Fig_1

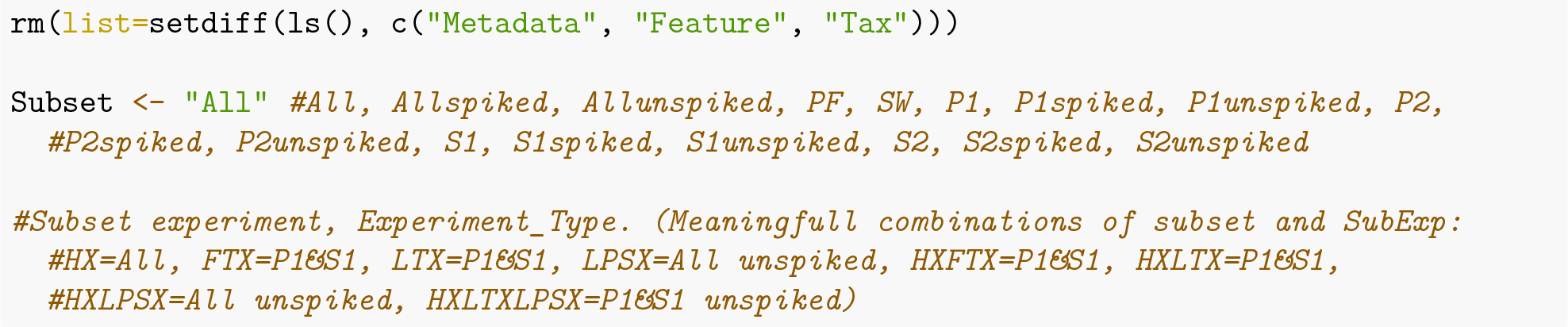
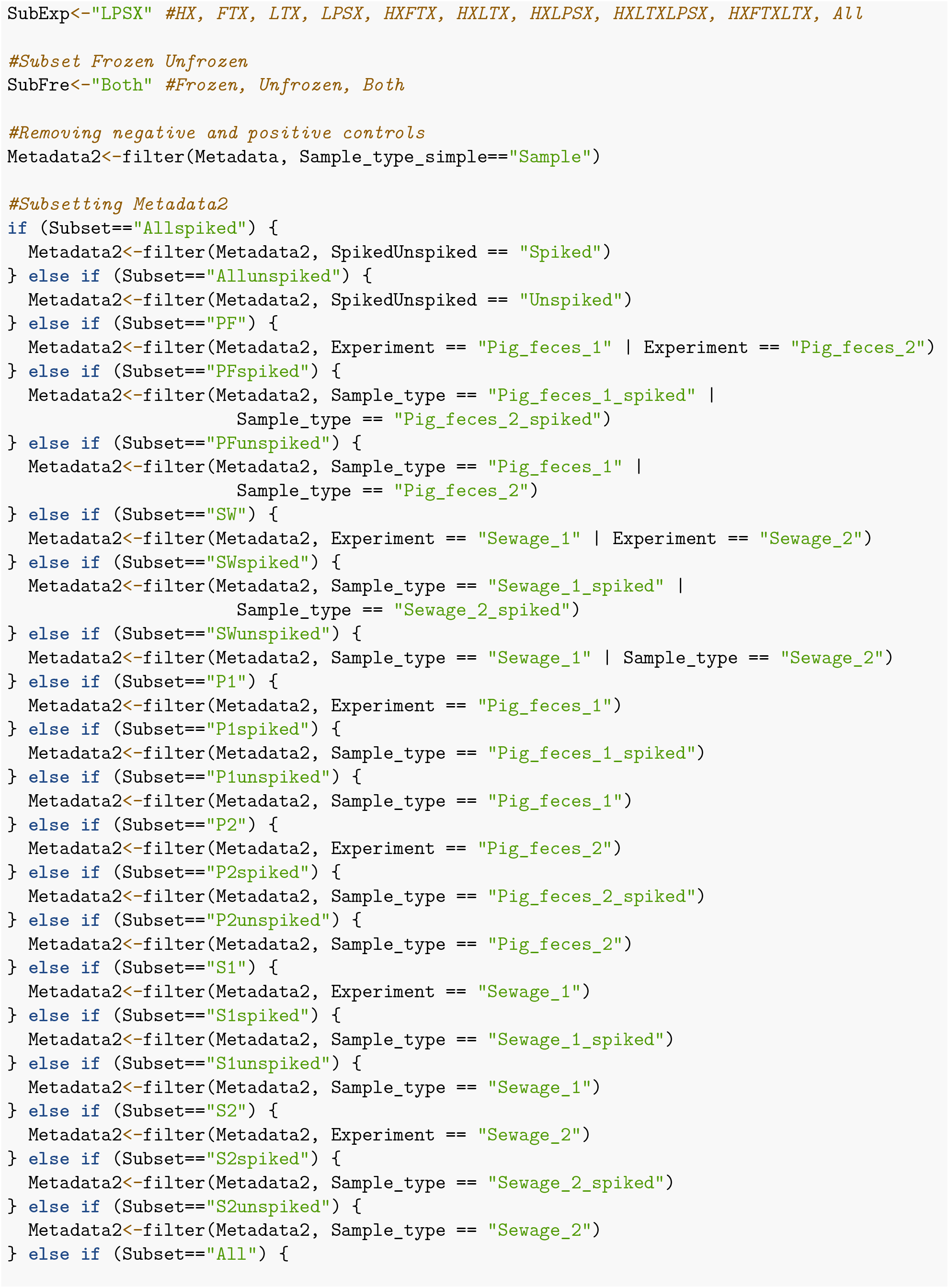
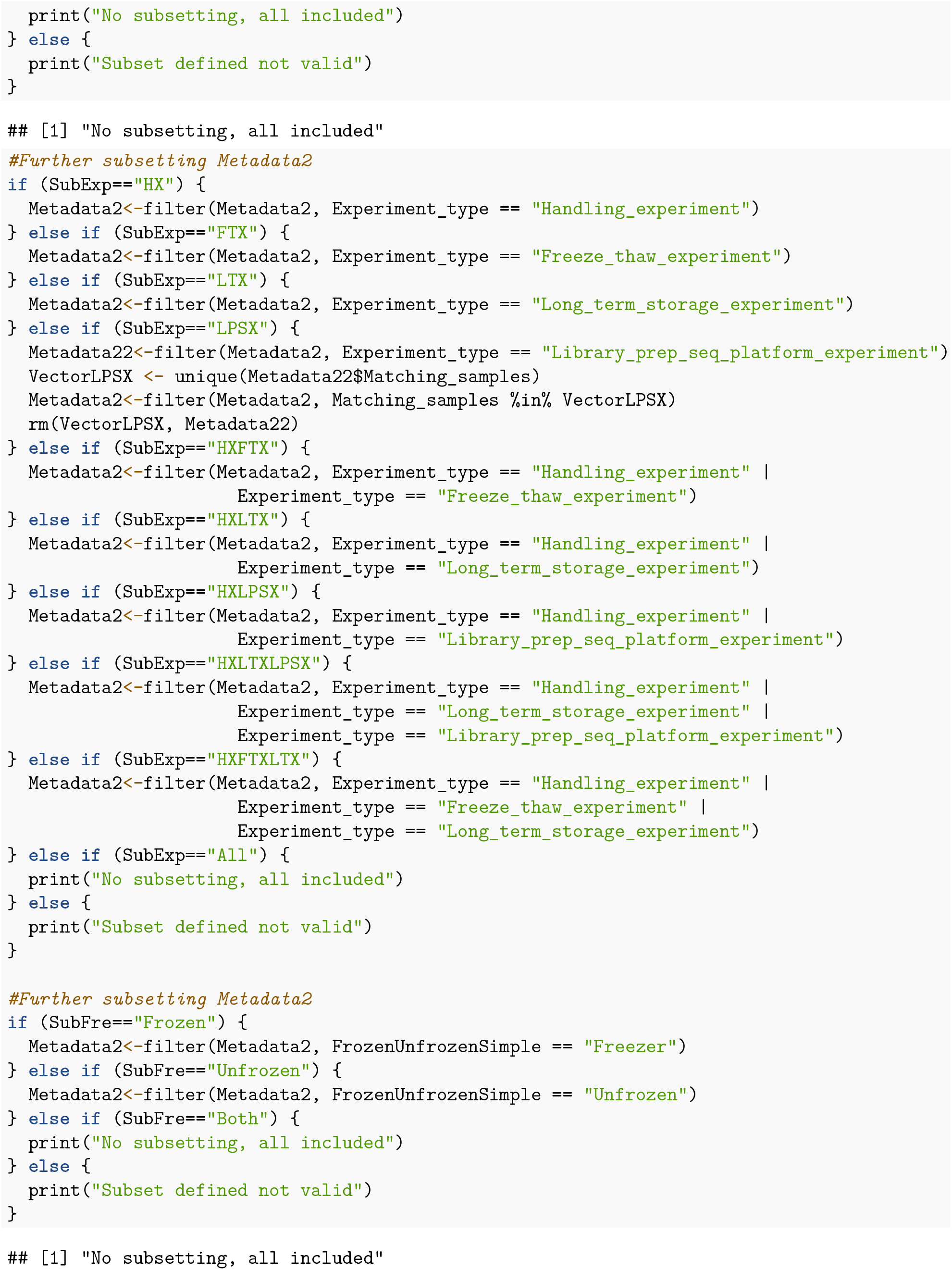
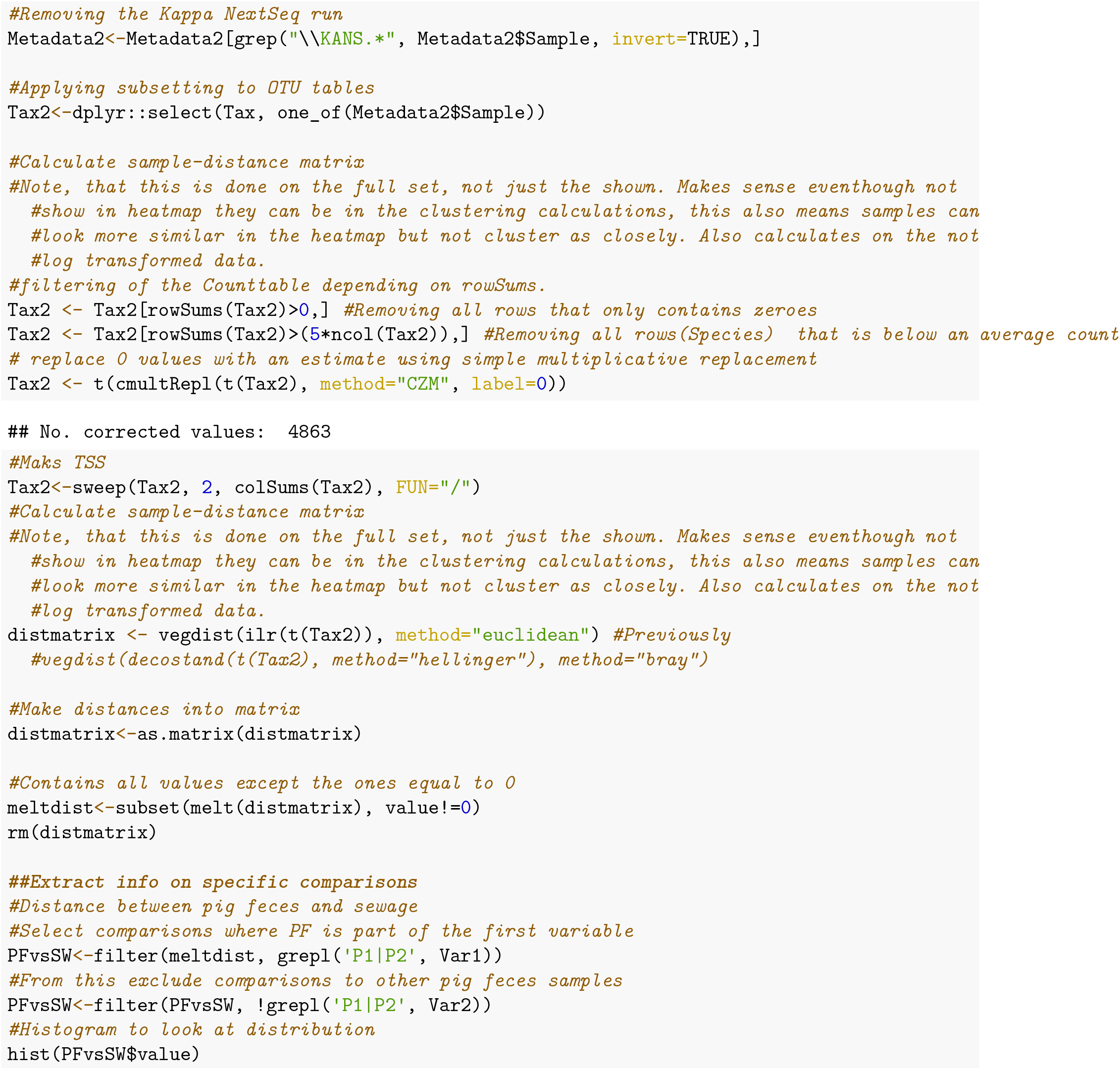
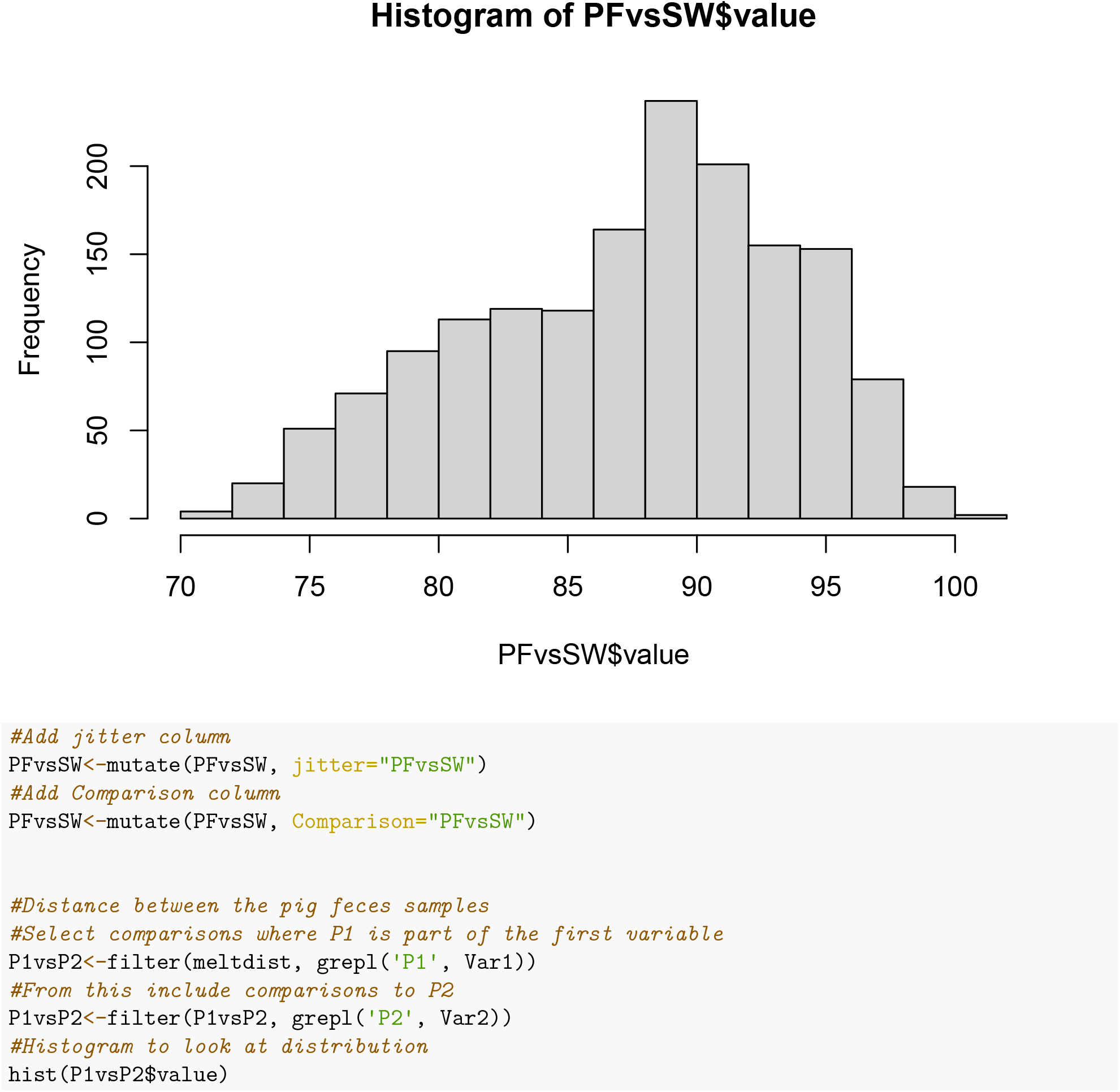
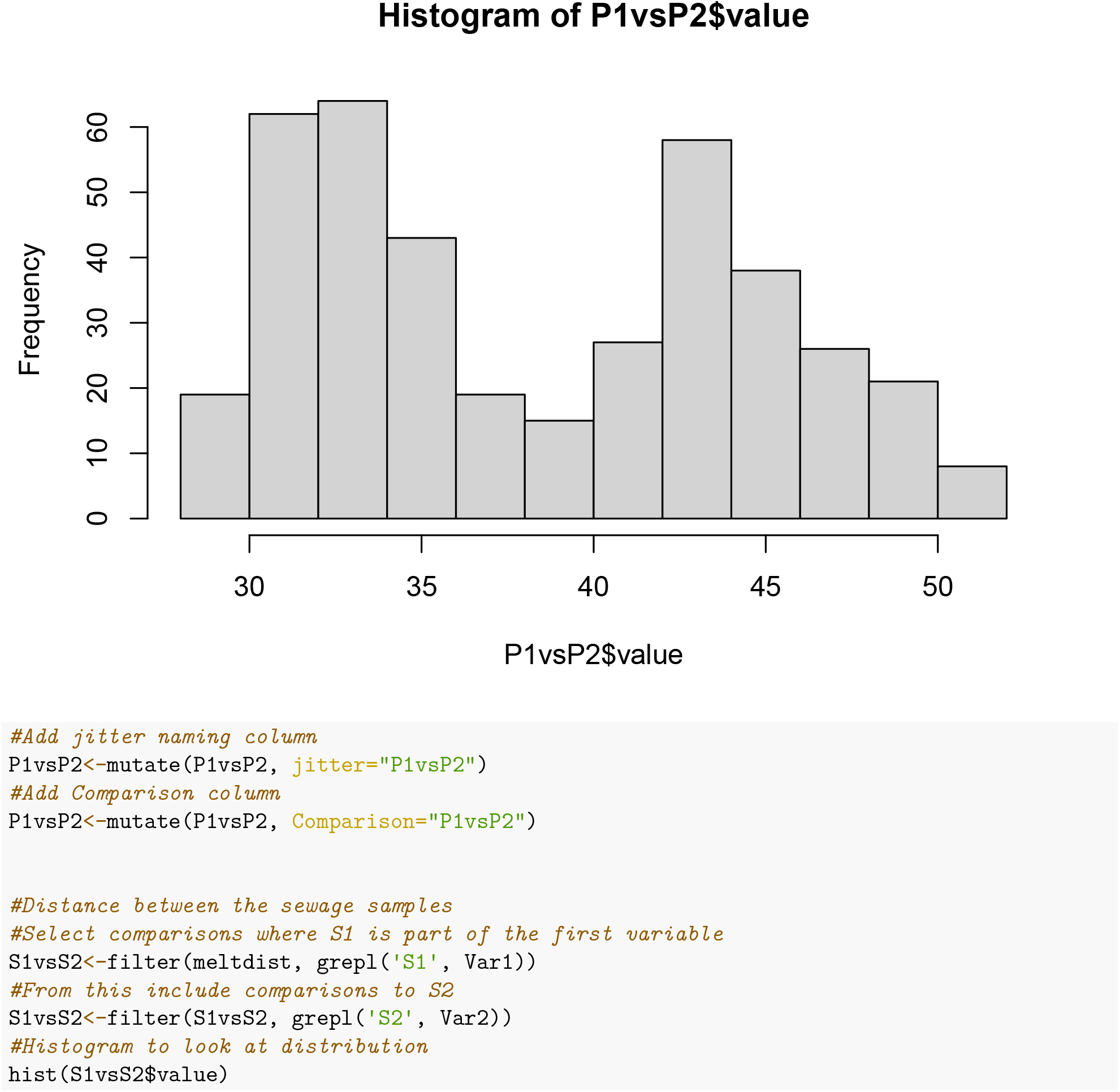
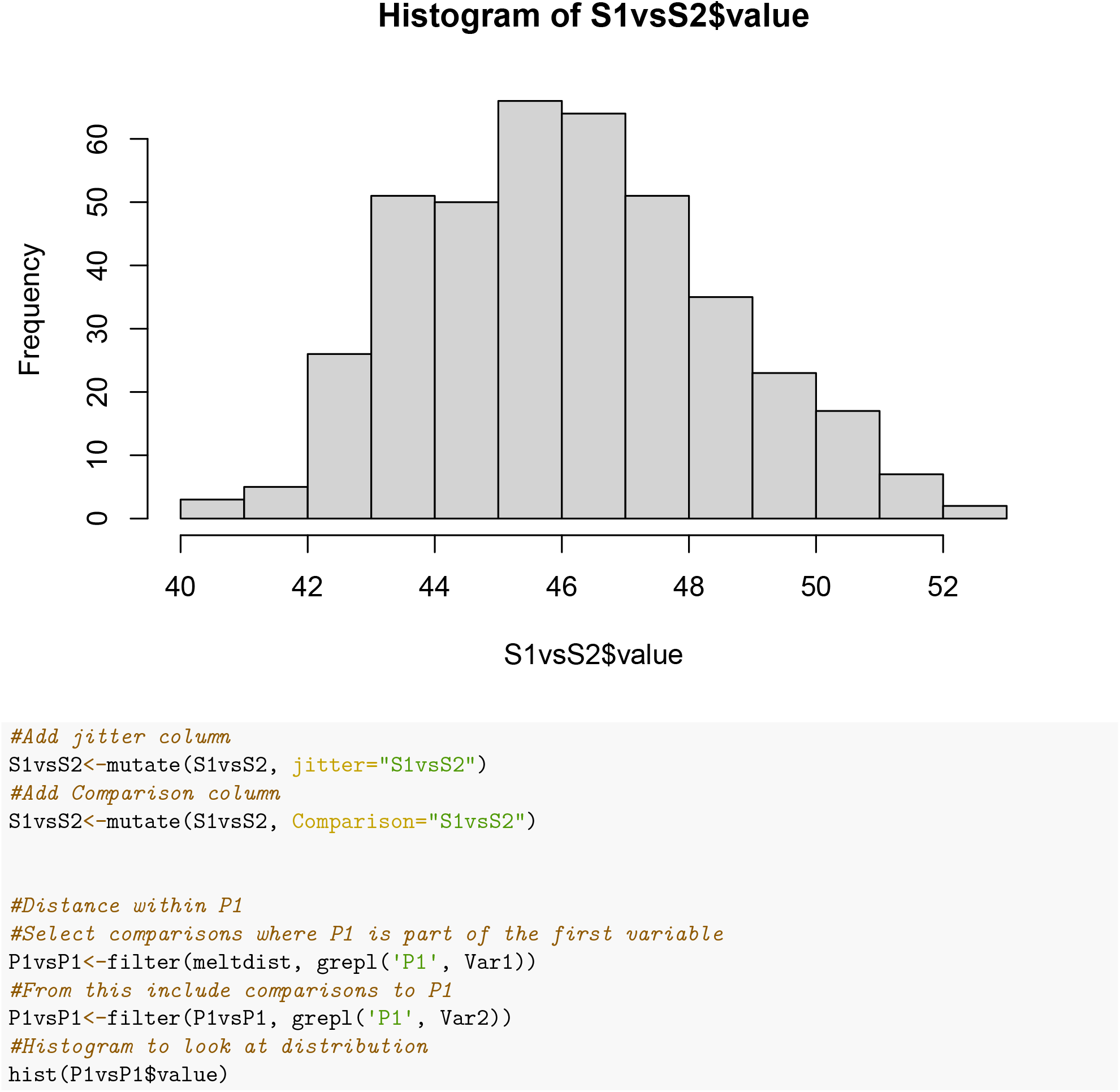
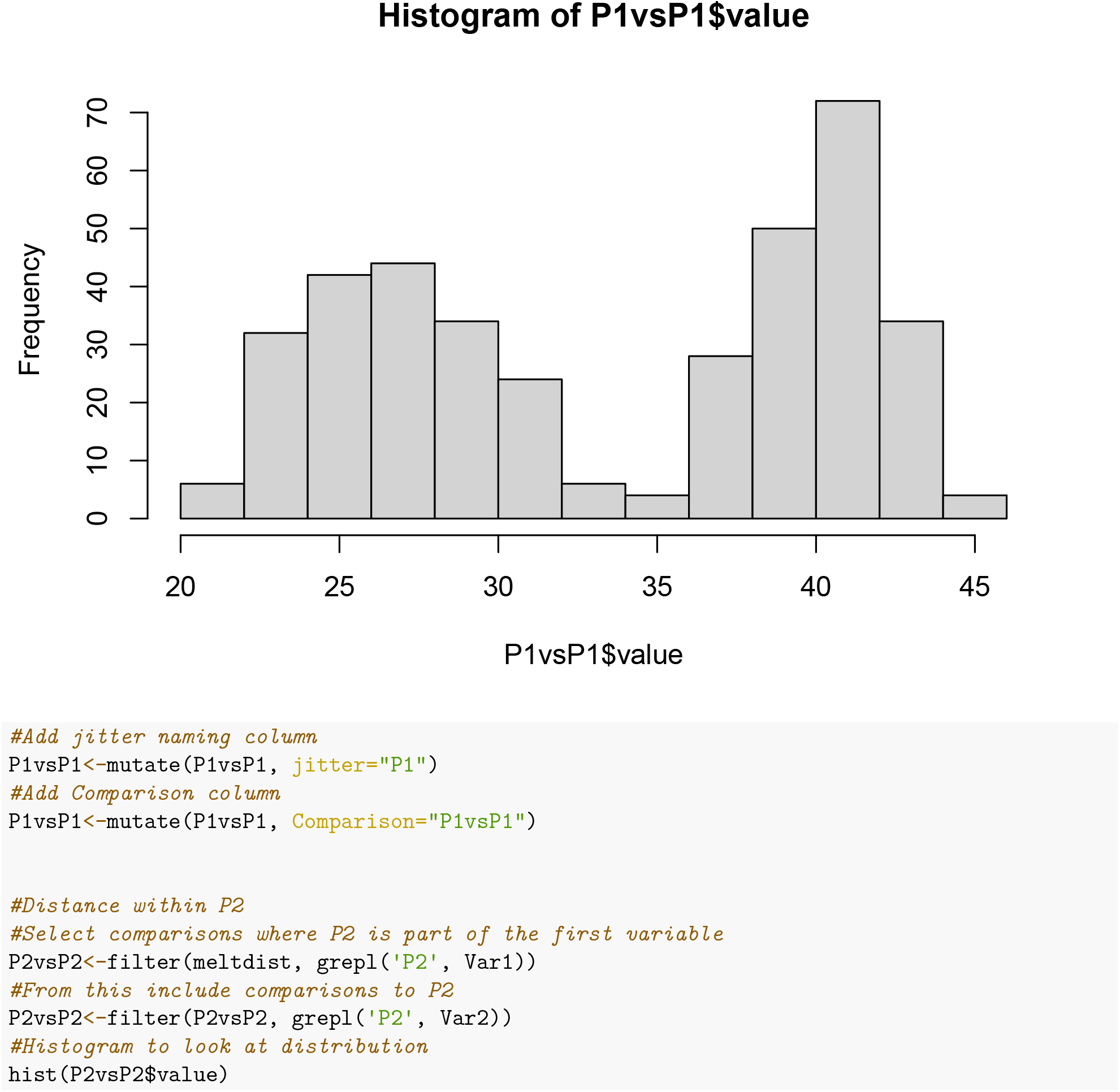
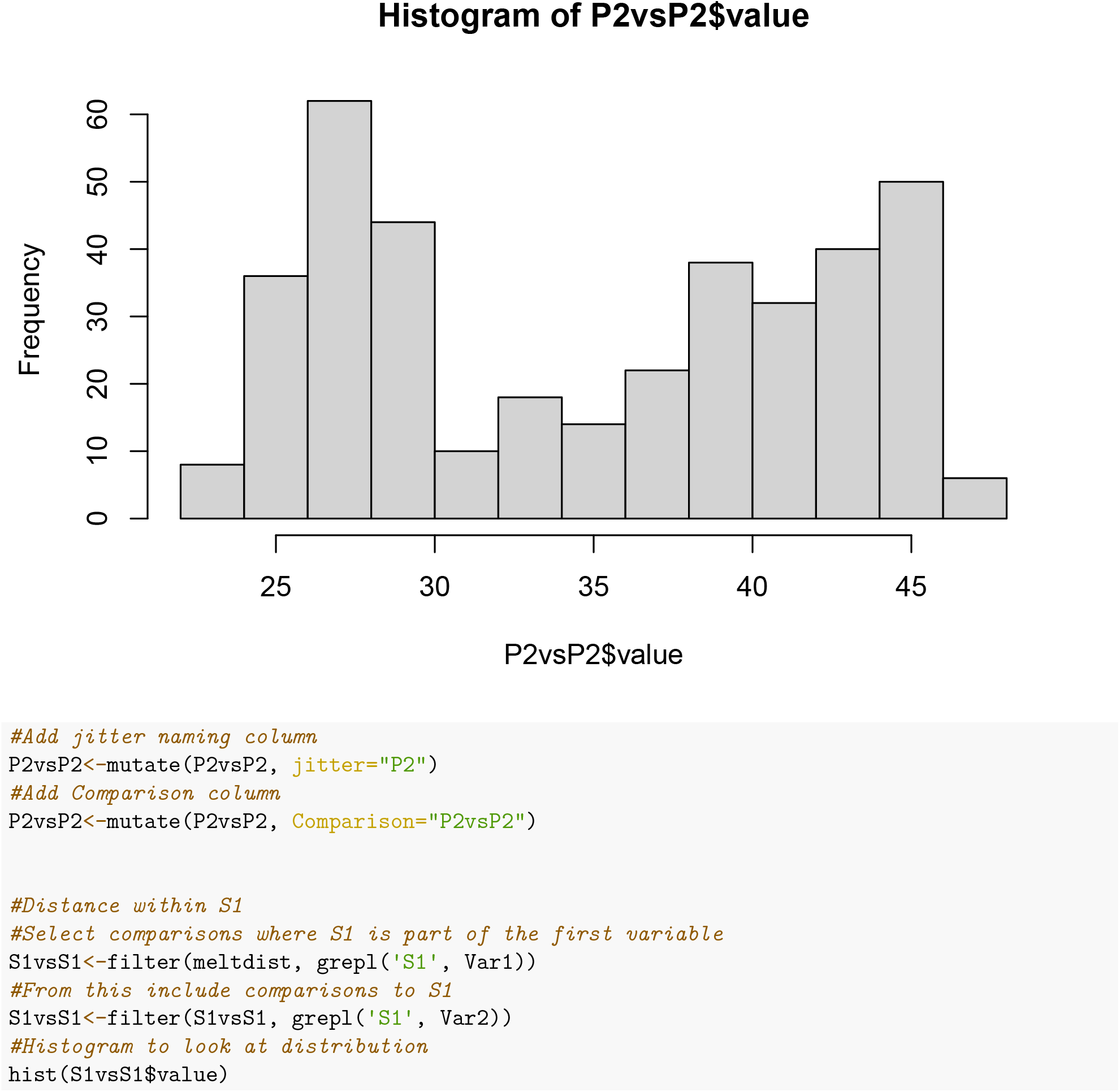
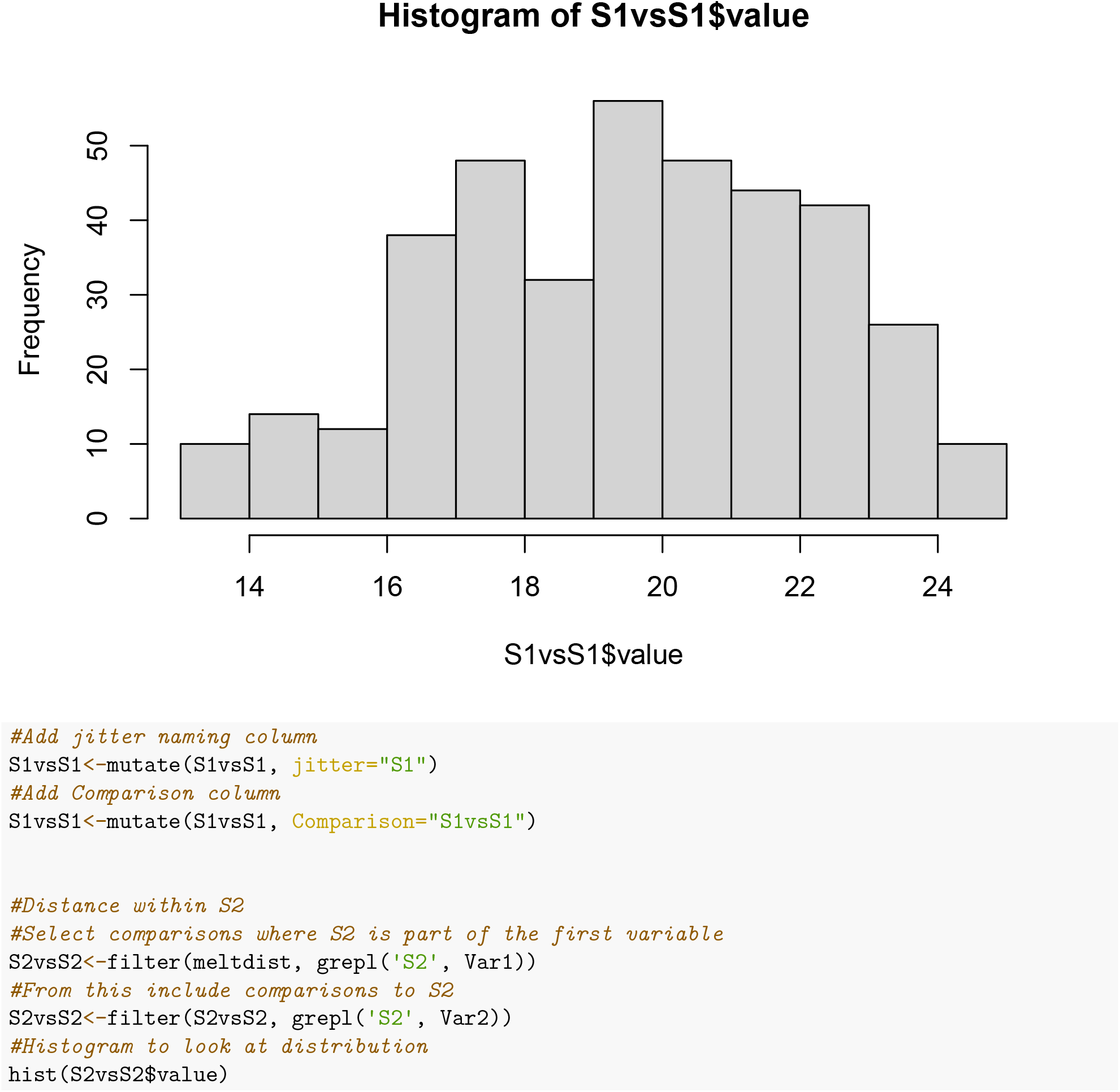
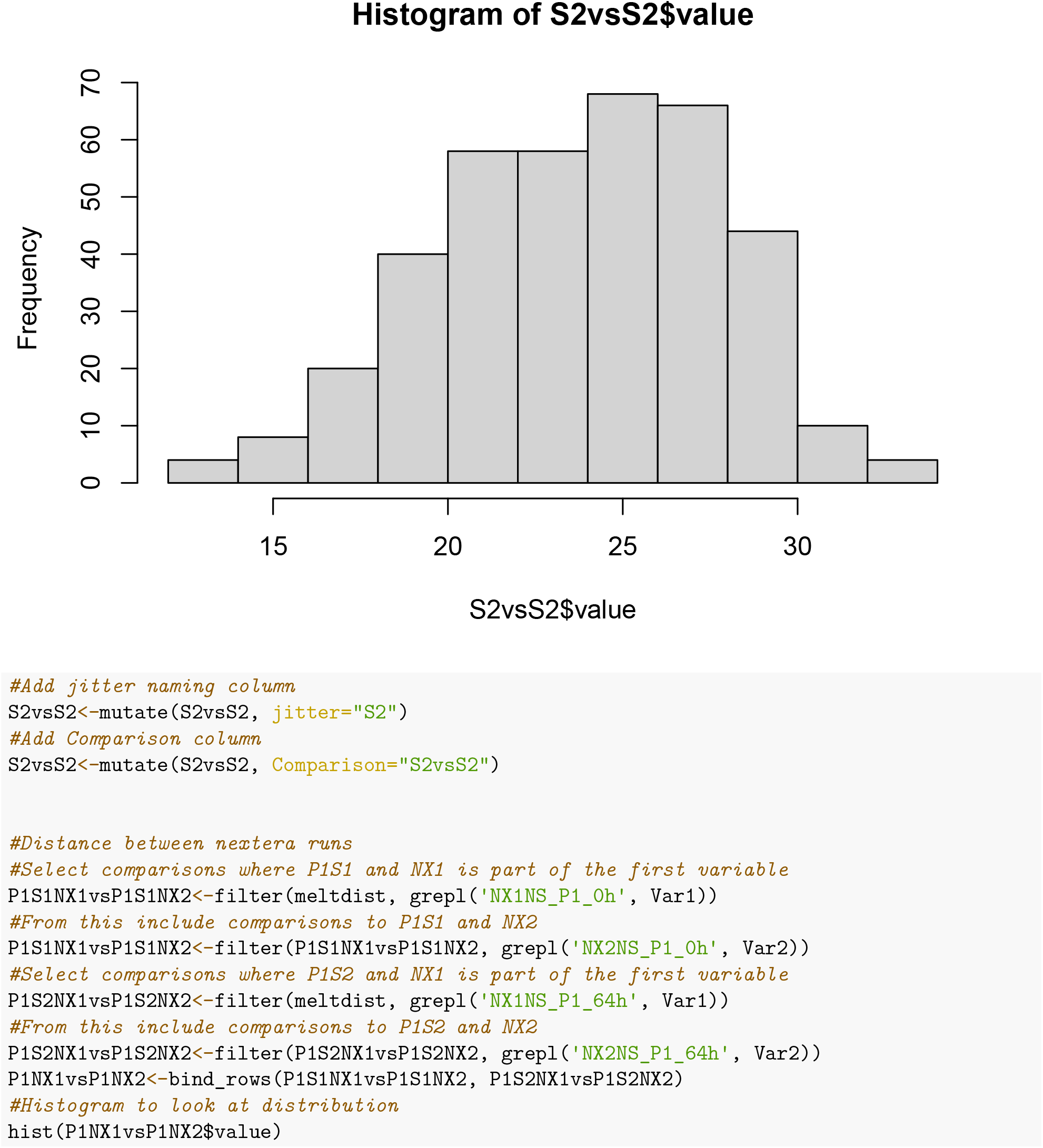
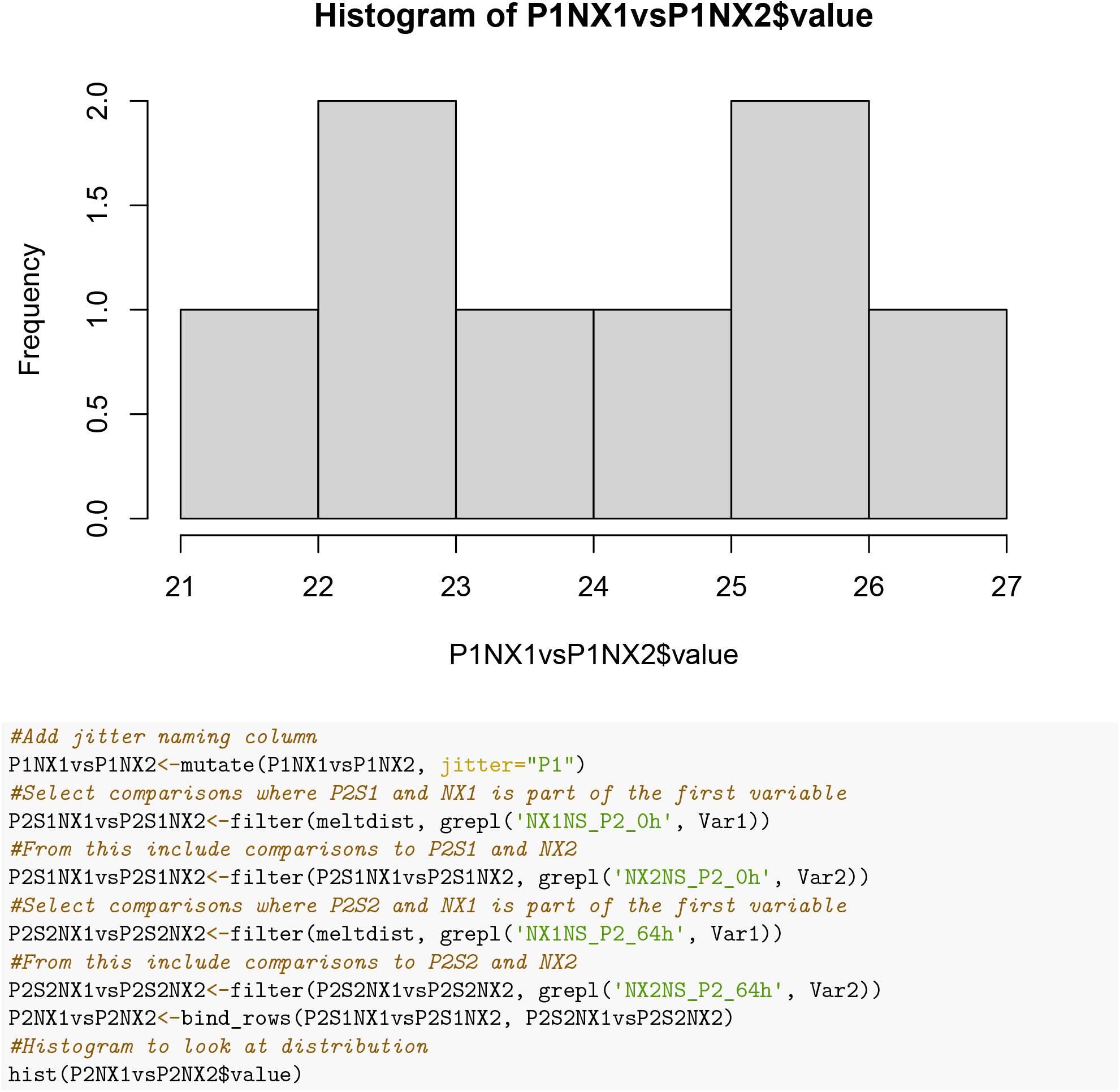
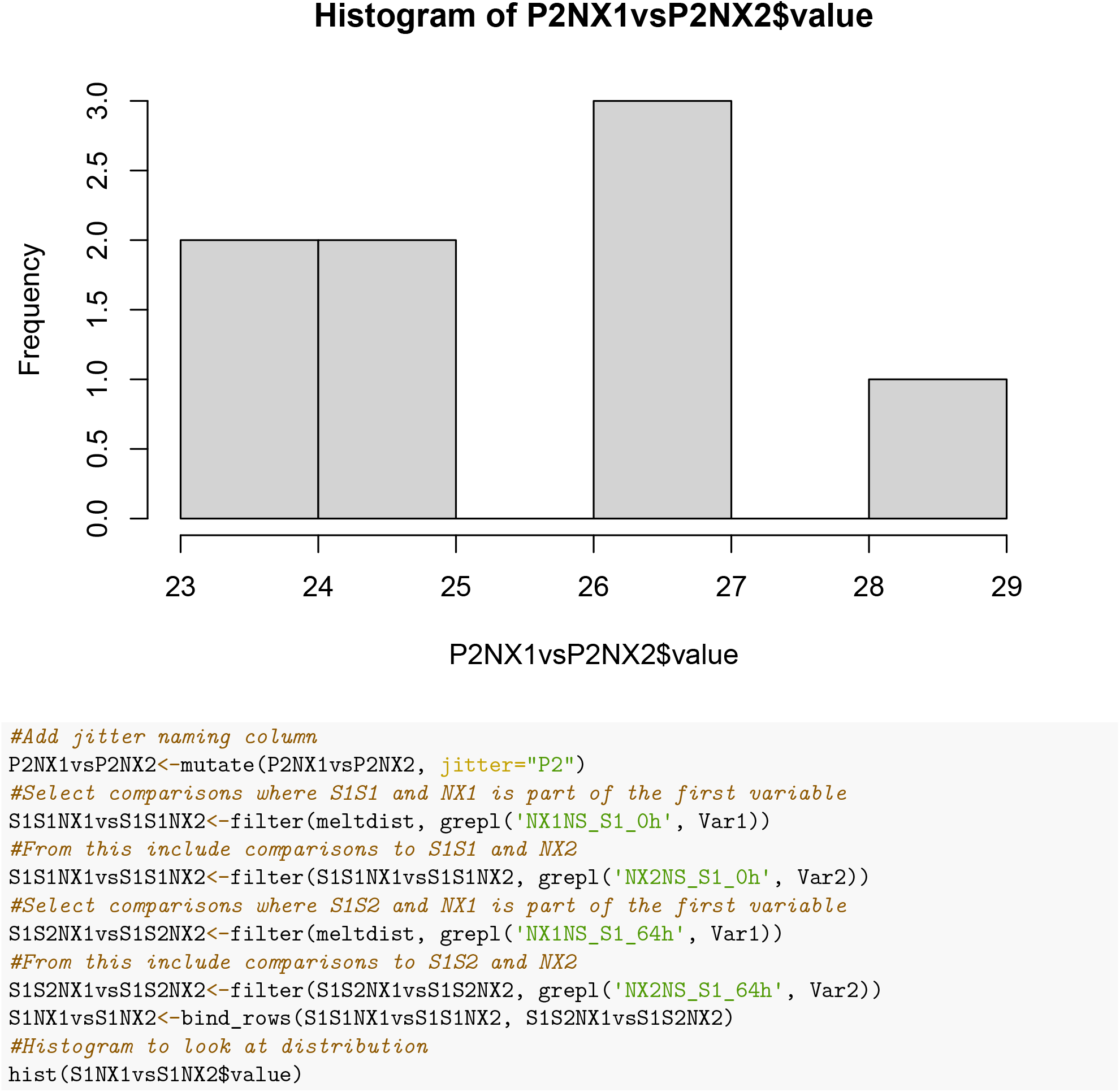
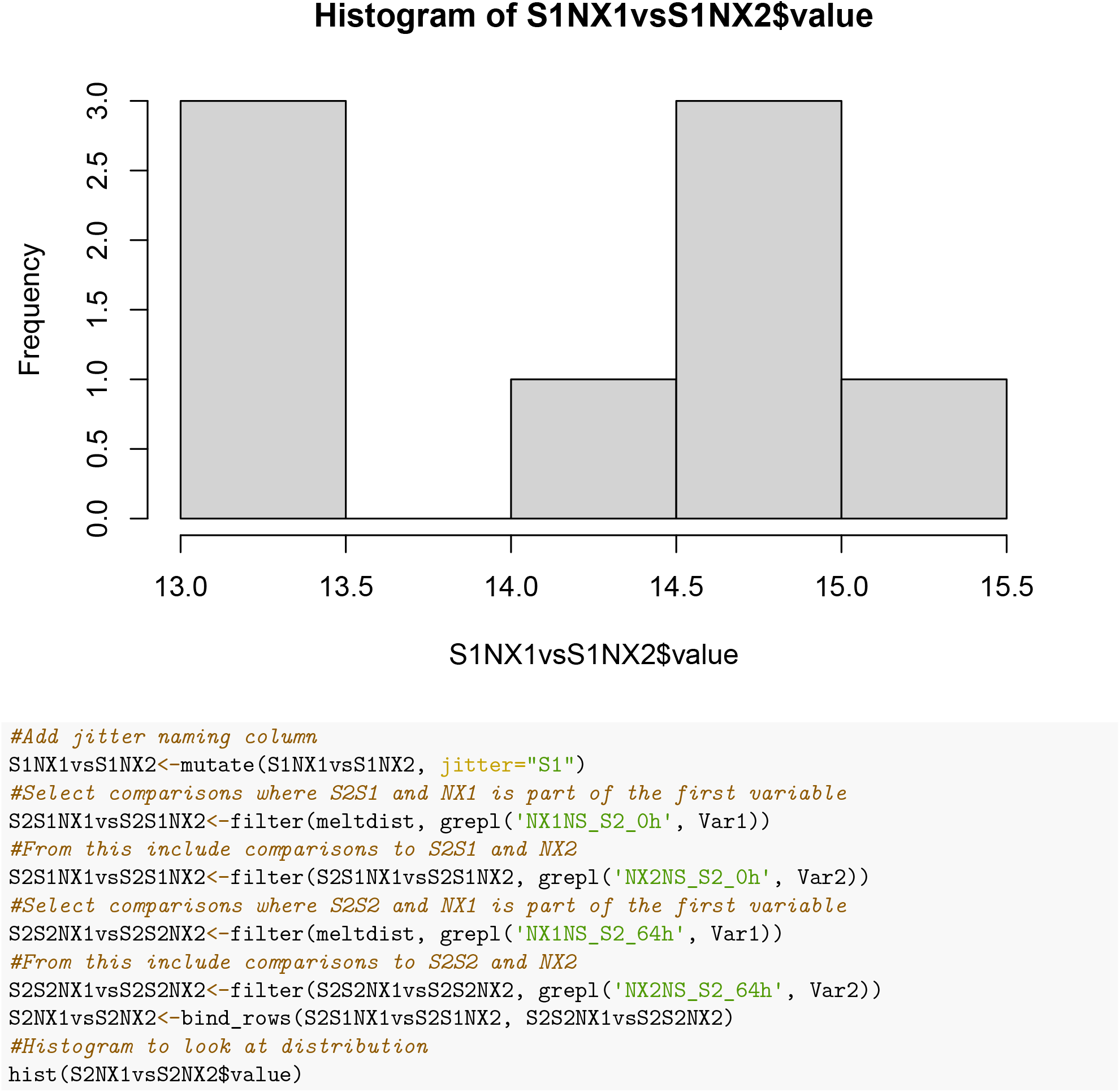
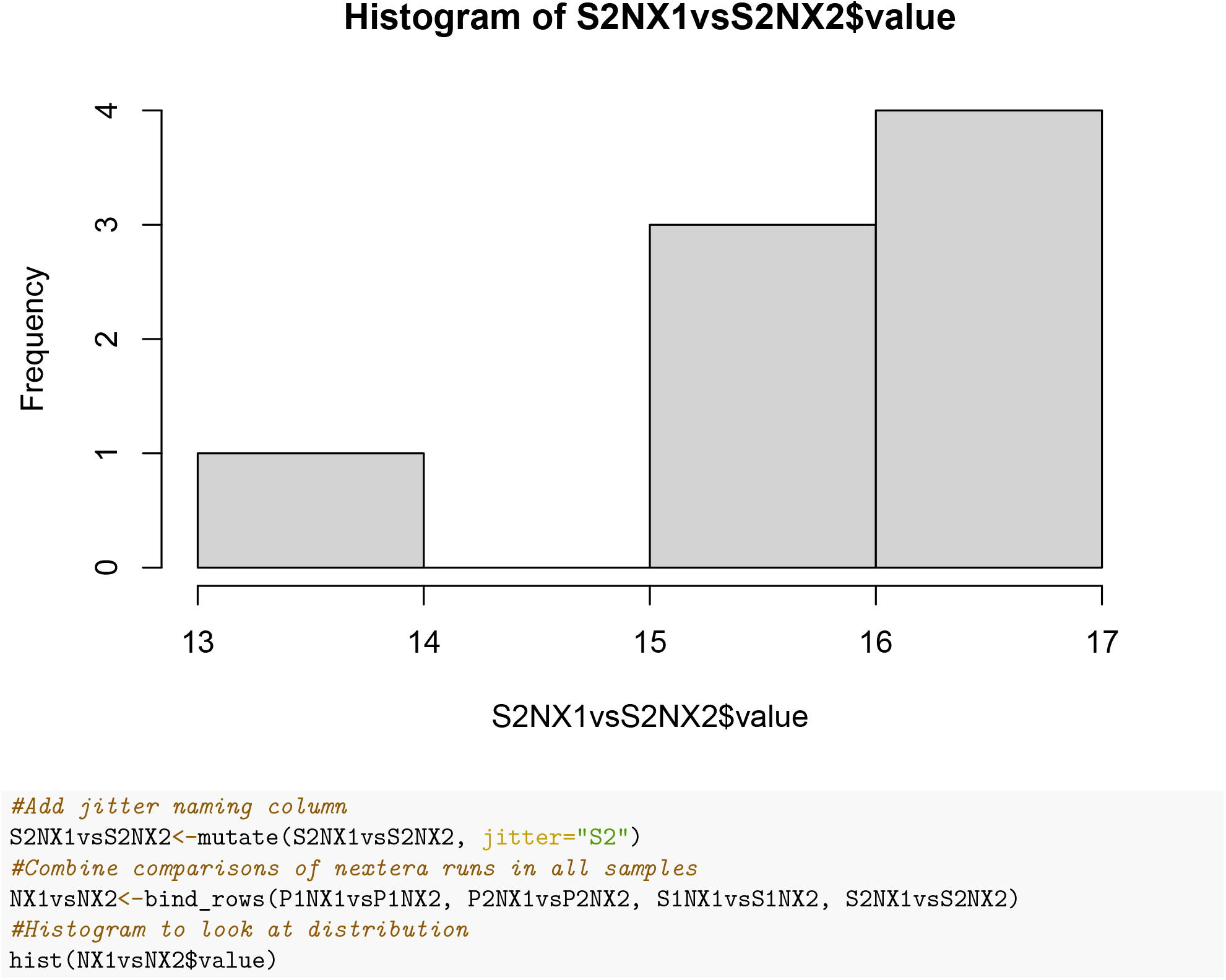
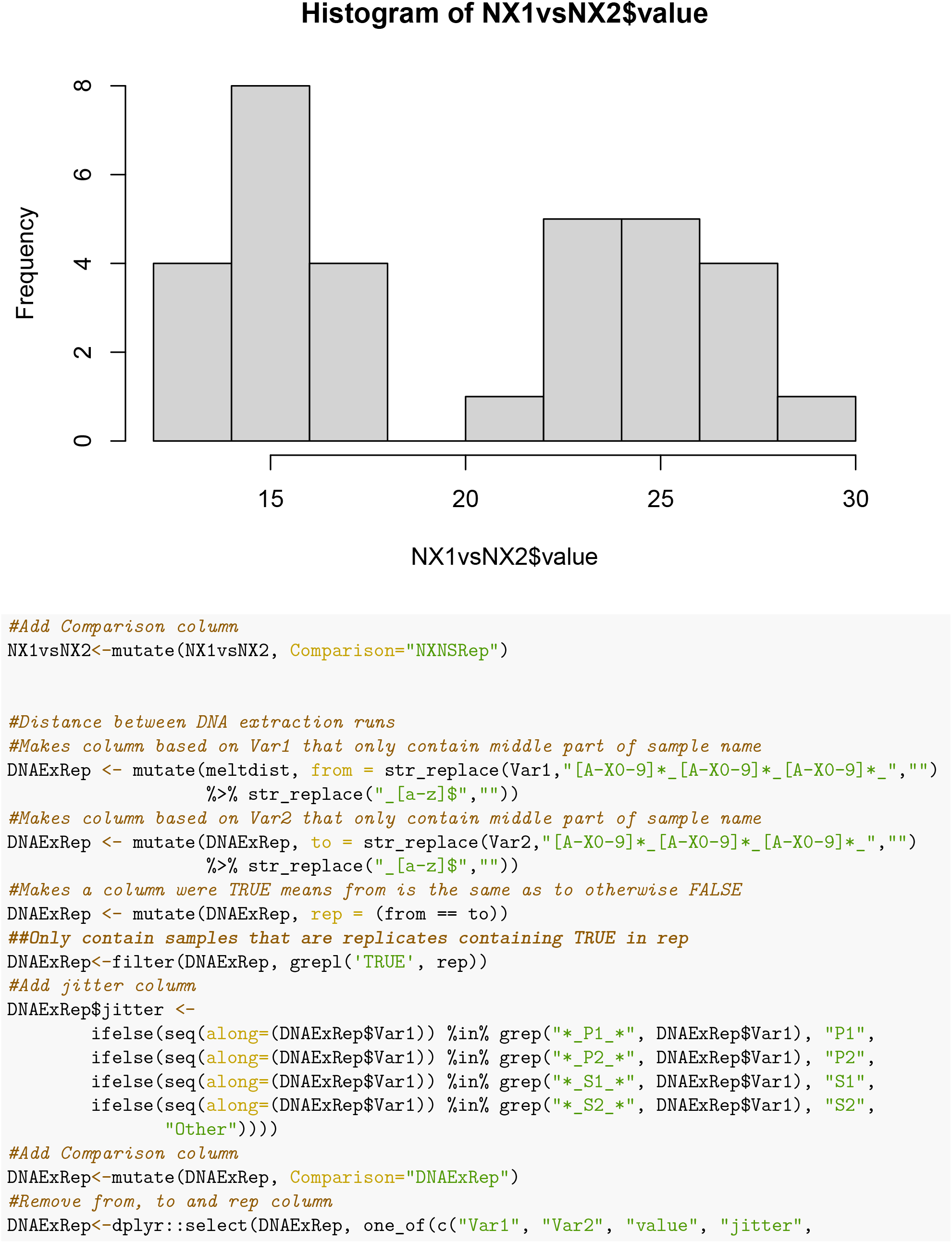
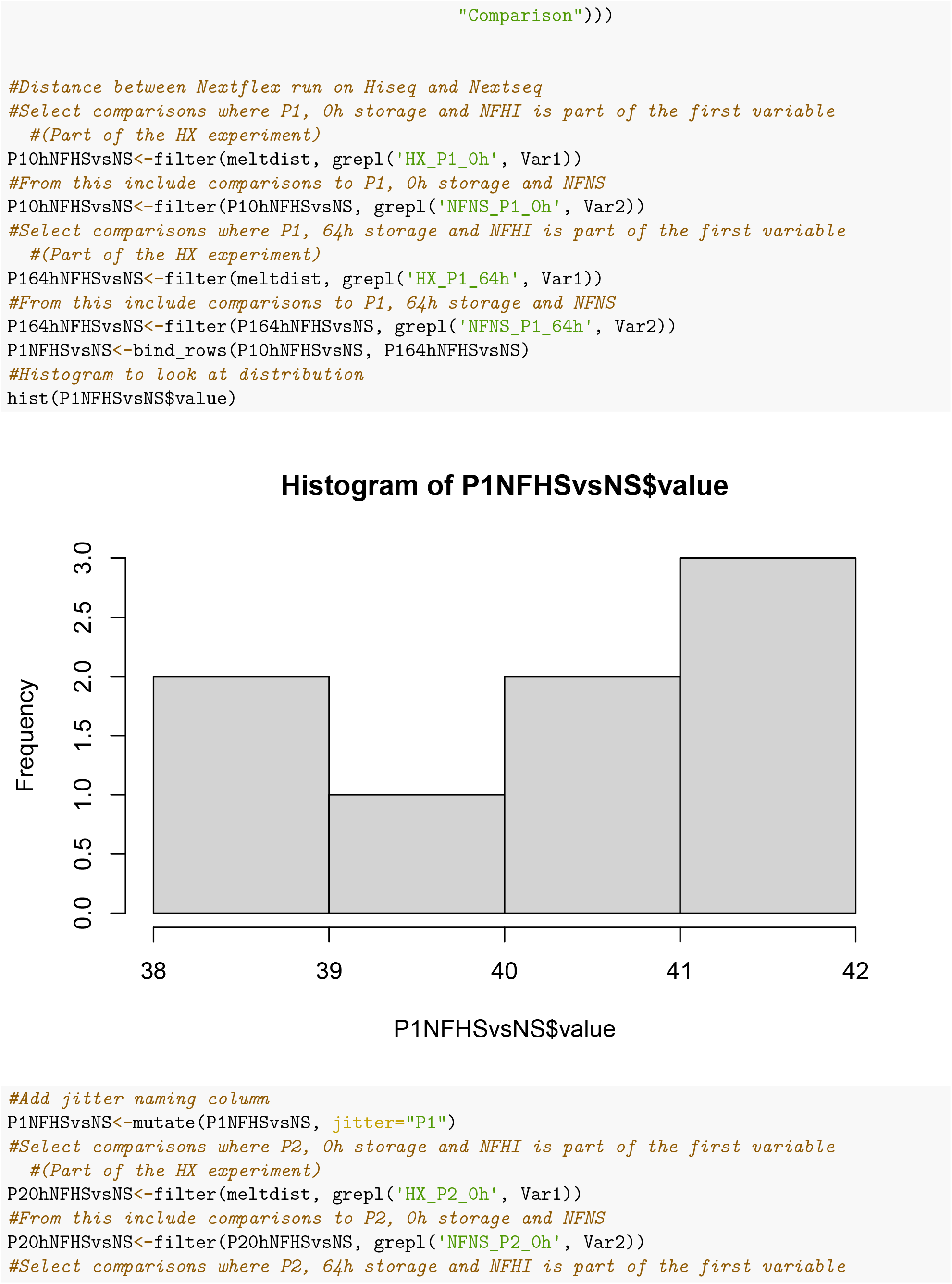
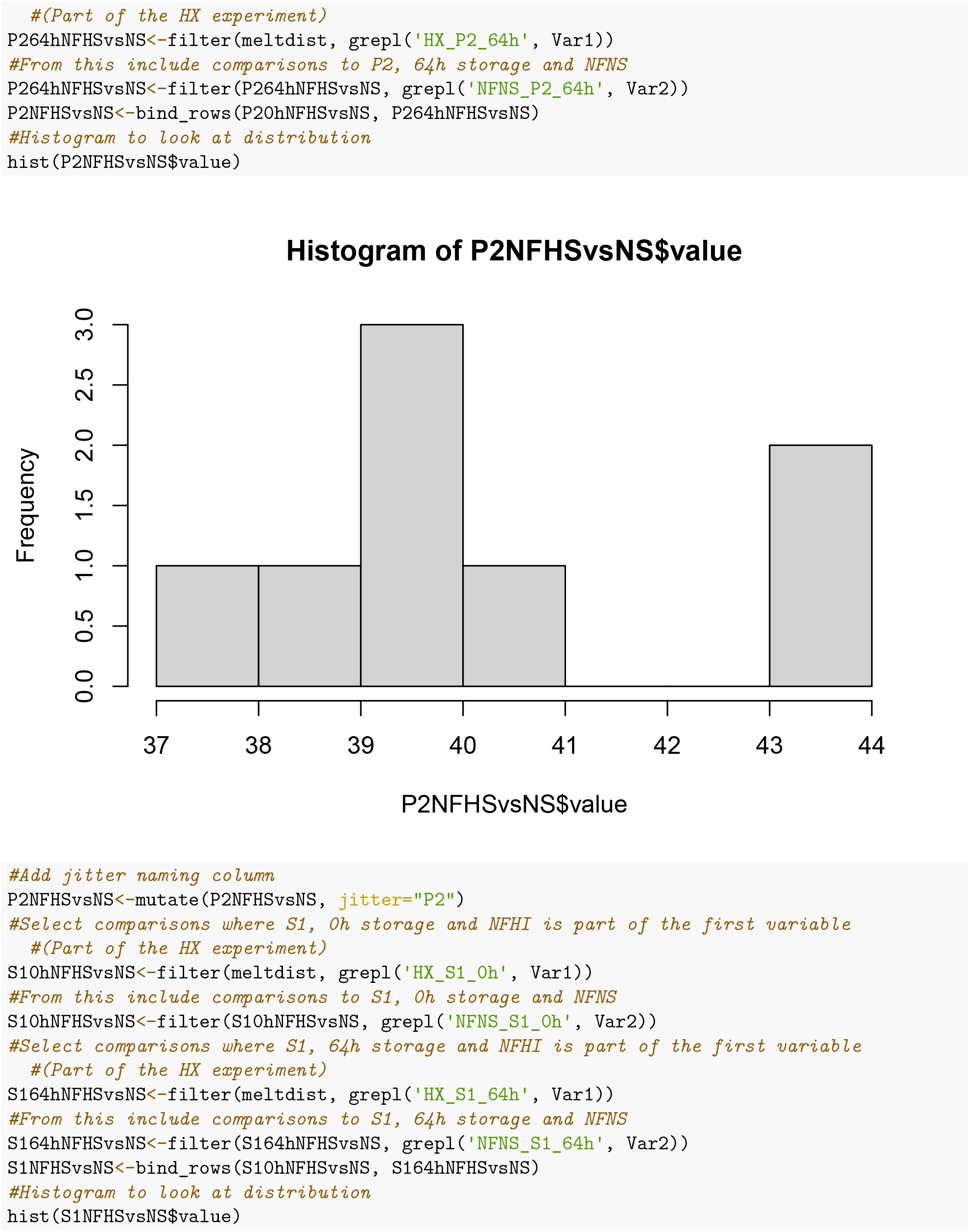
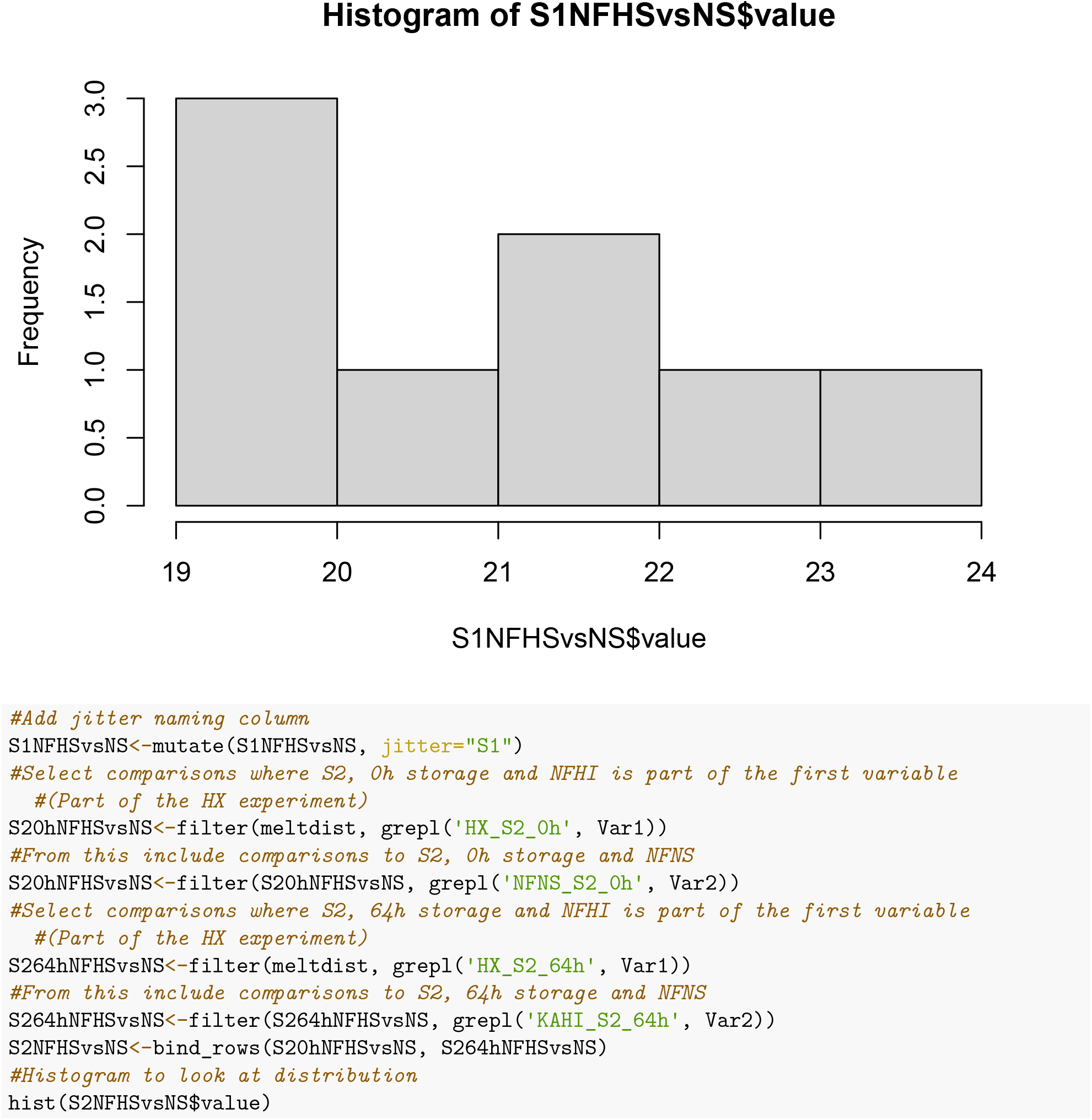
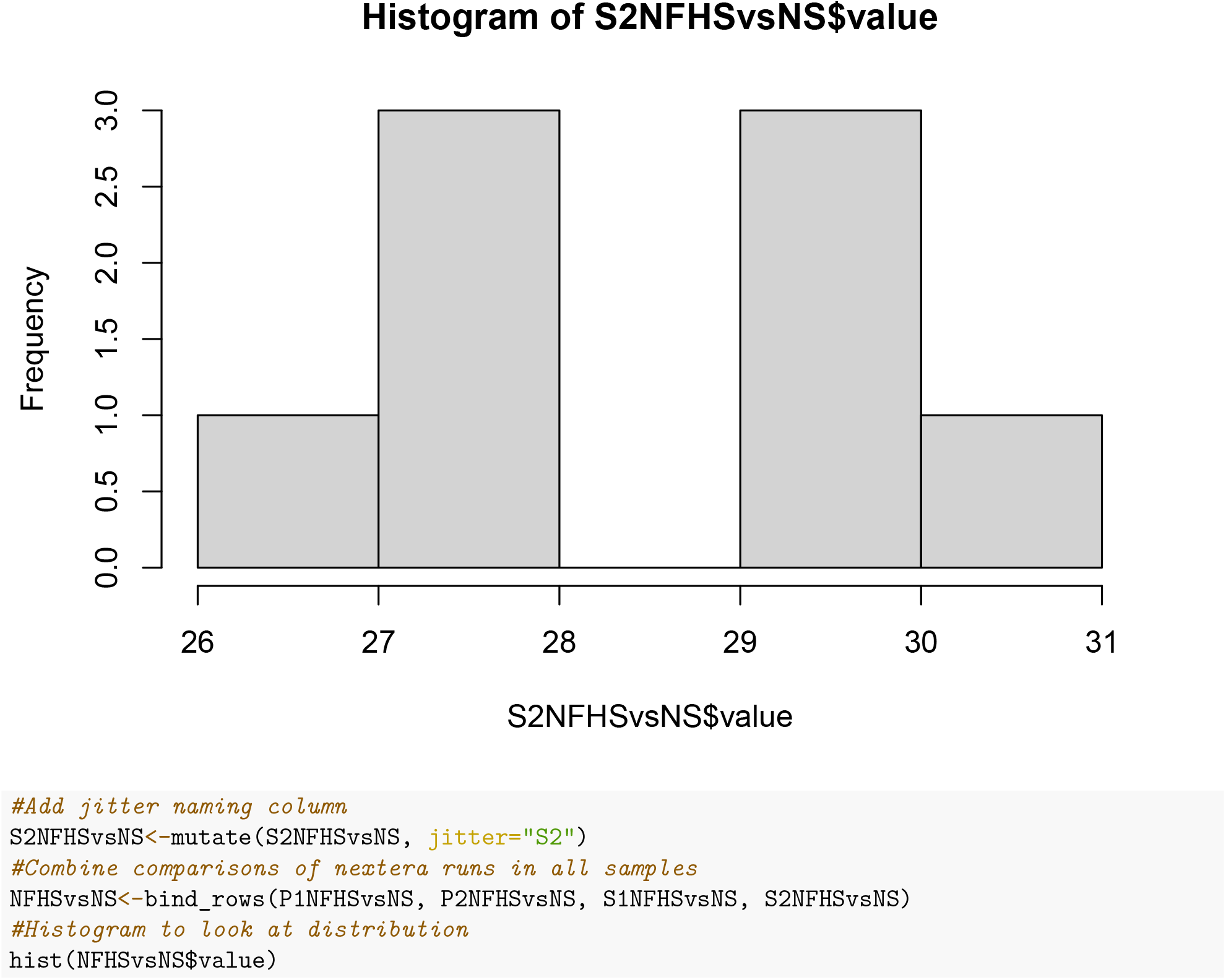
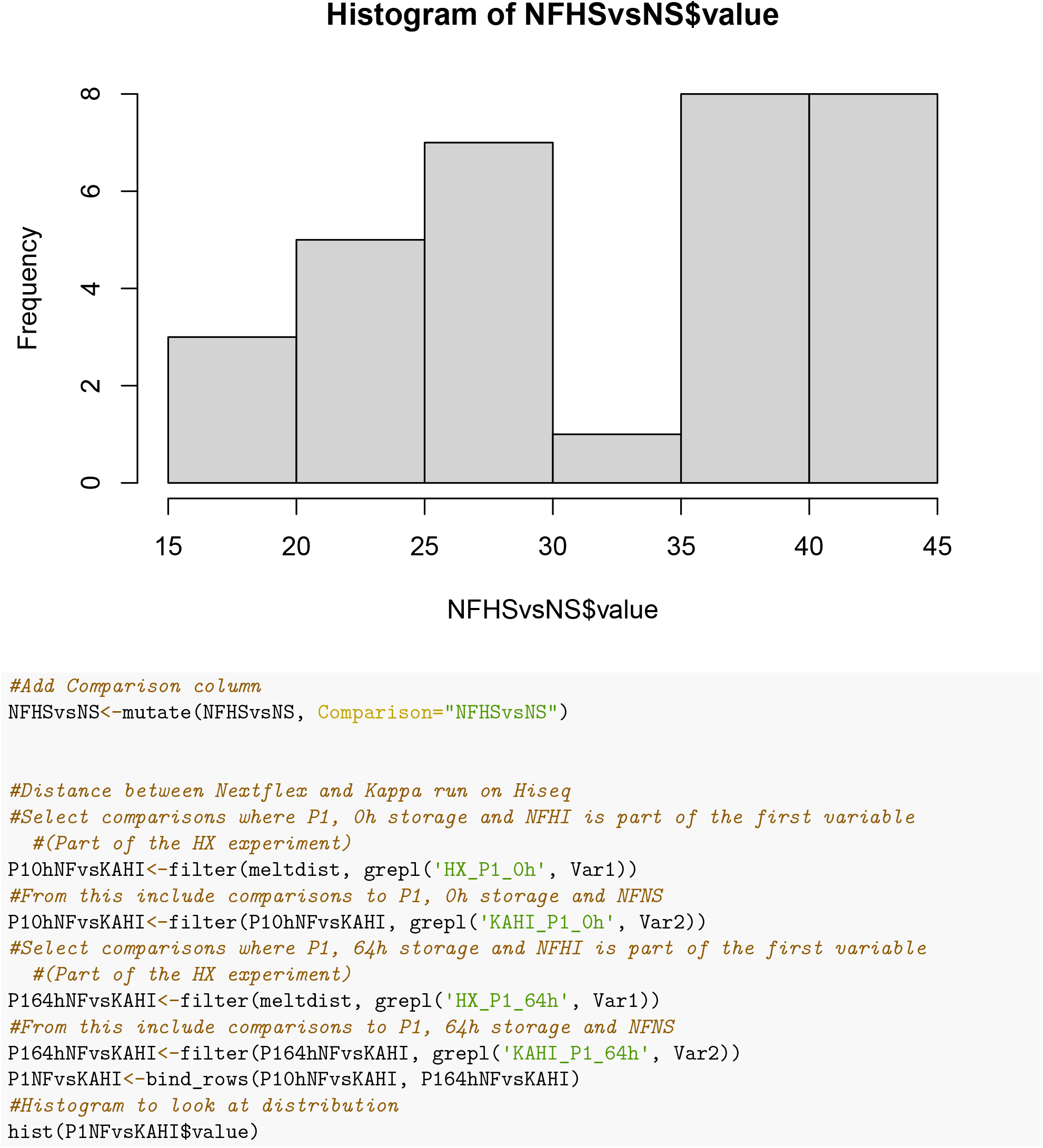
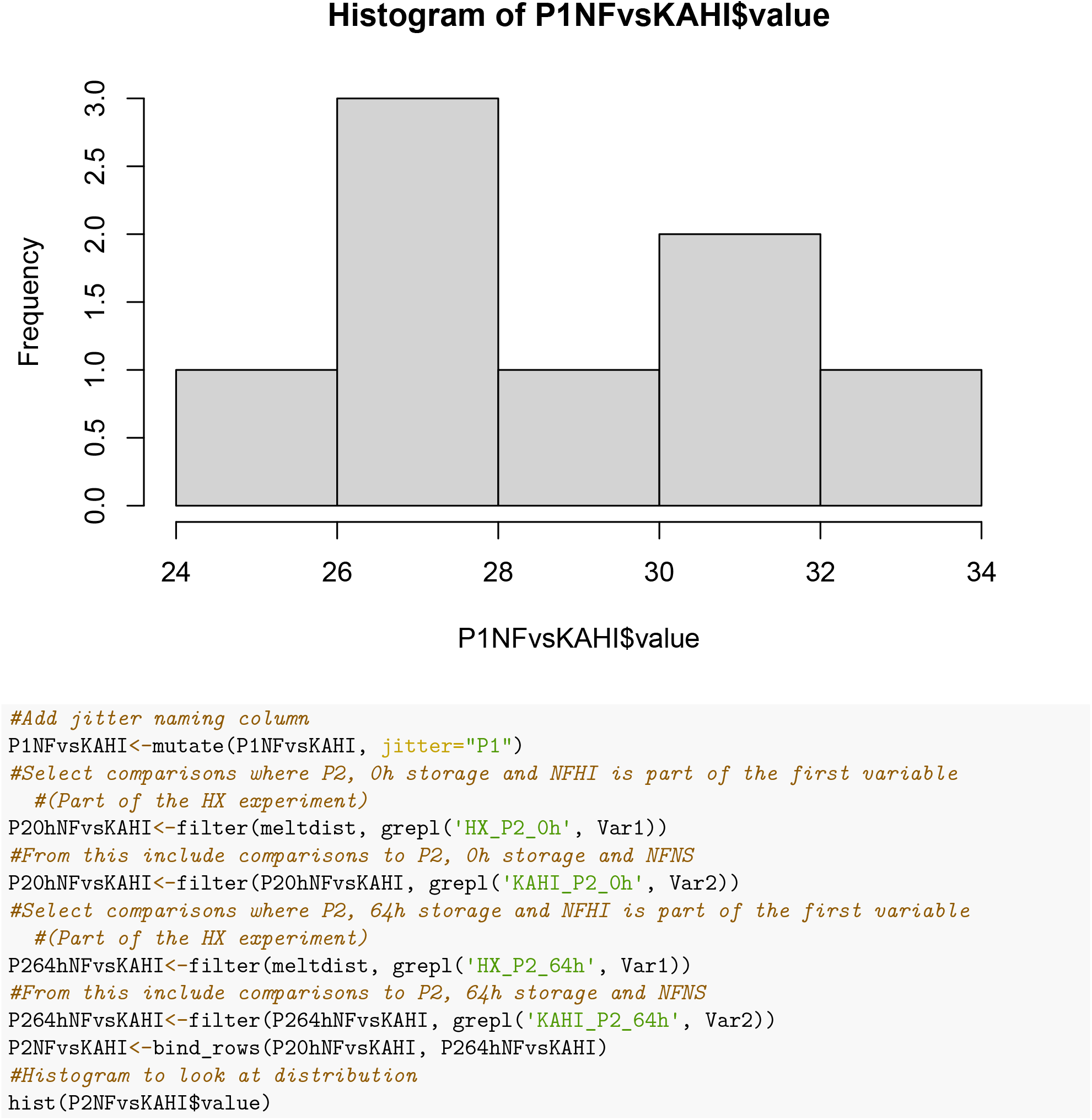
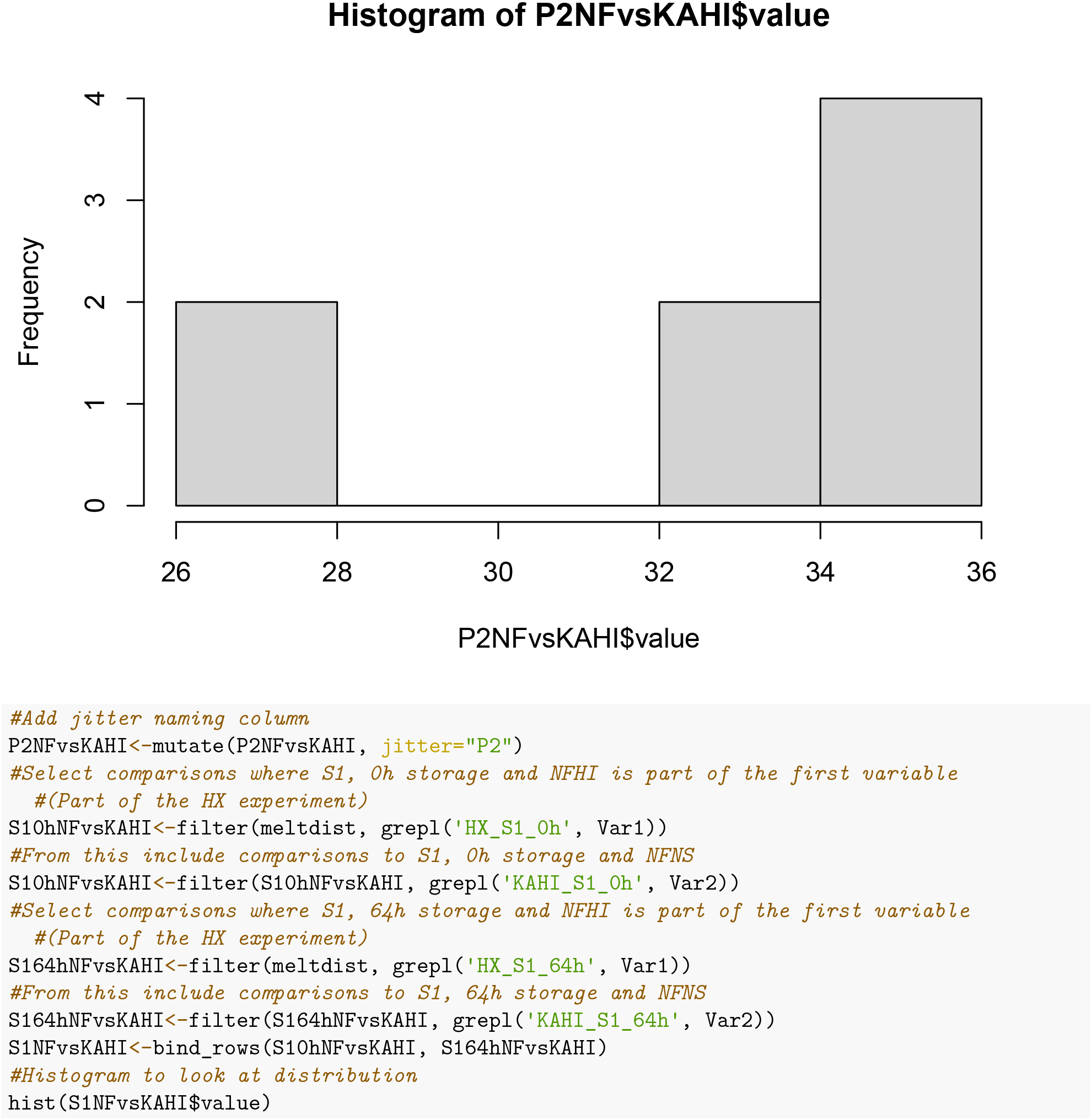
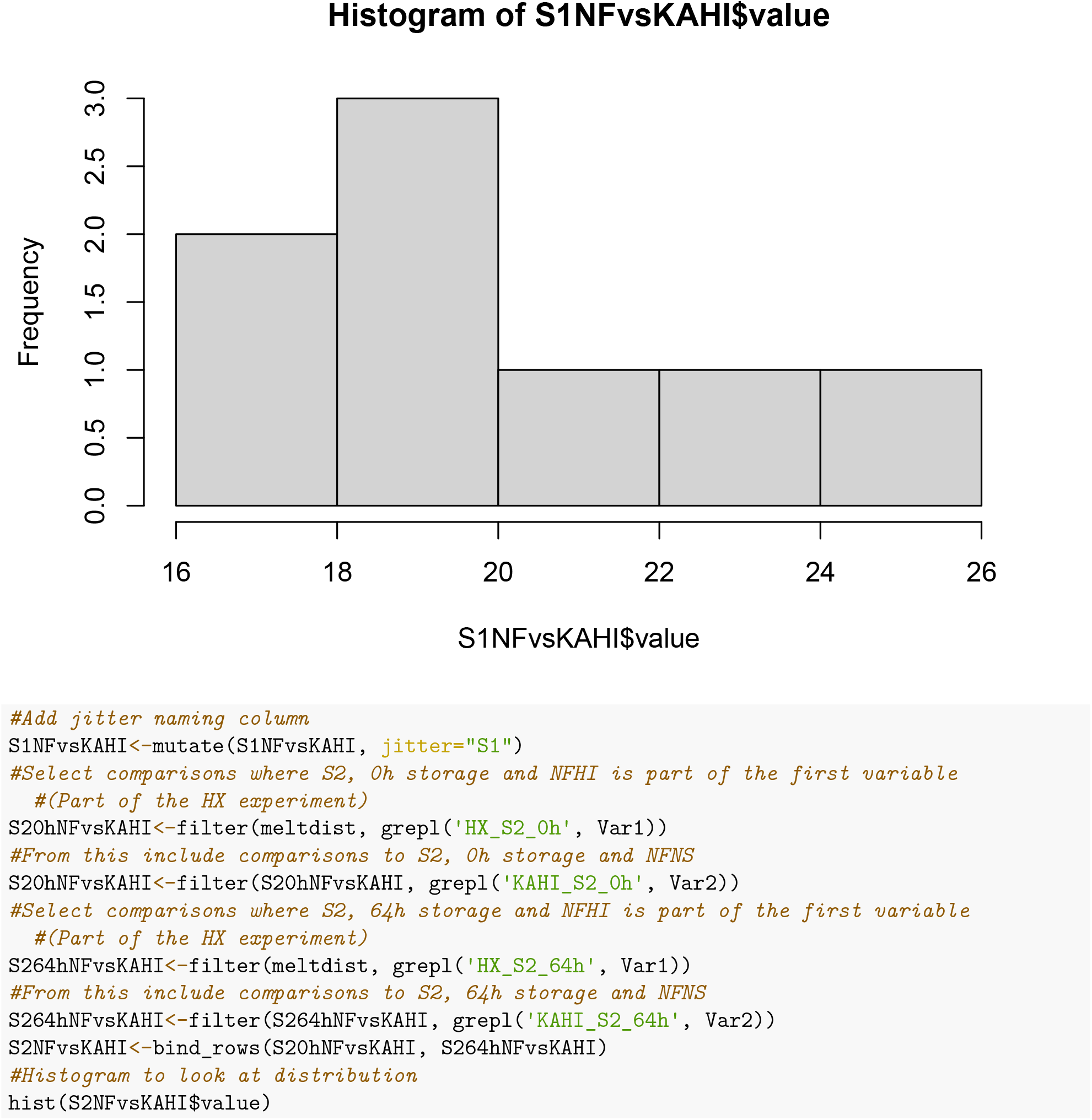
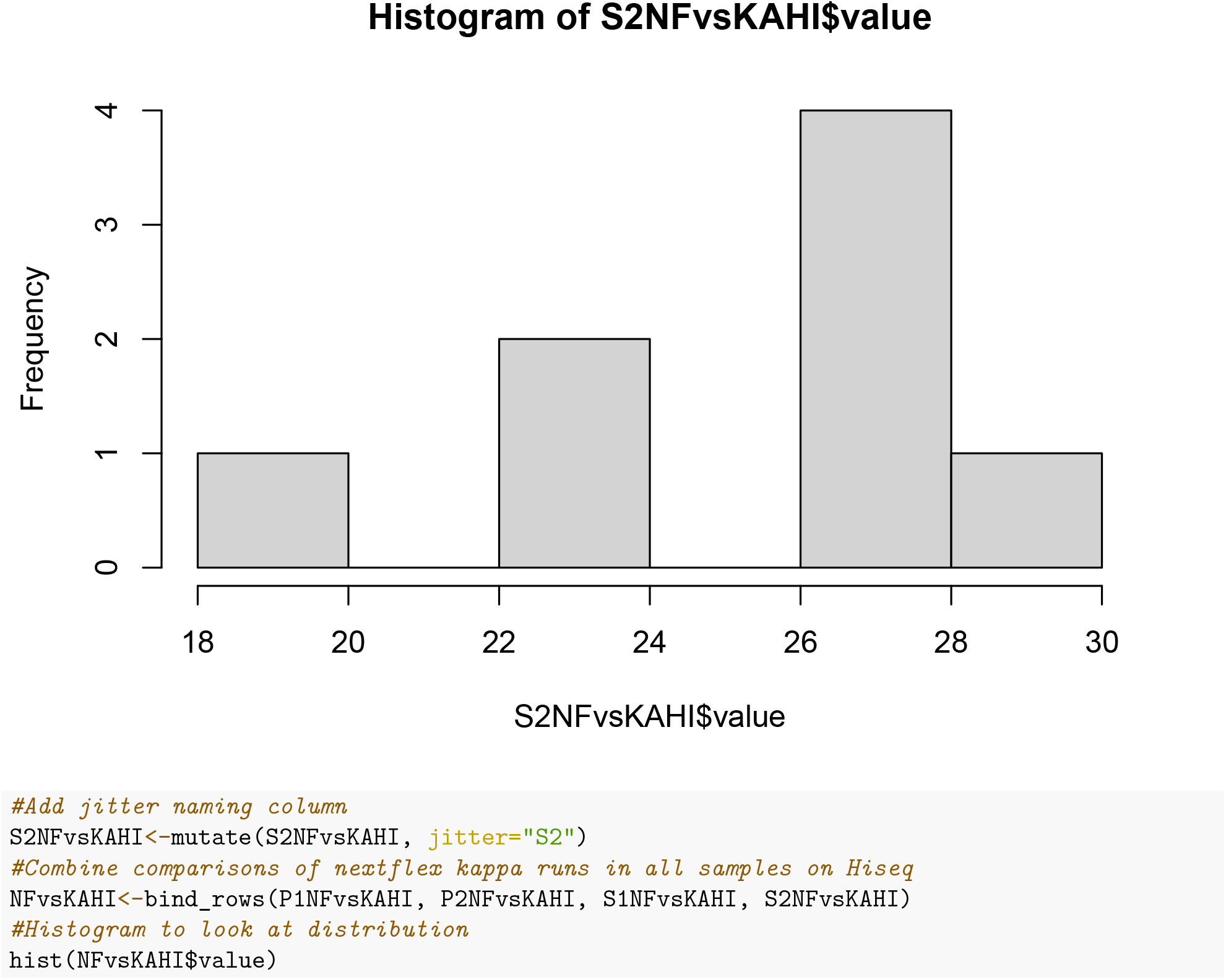
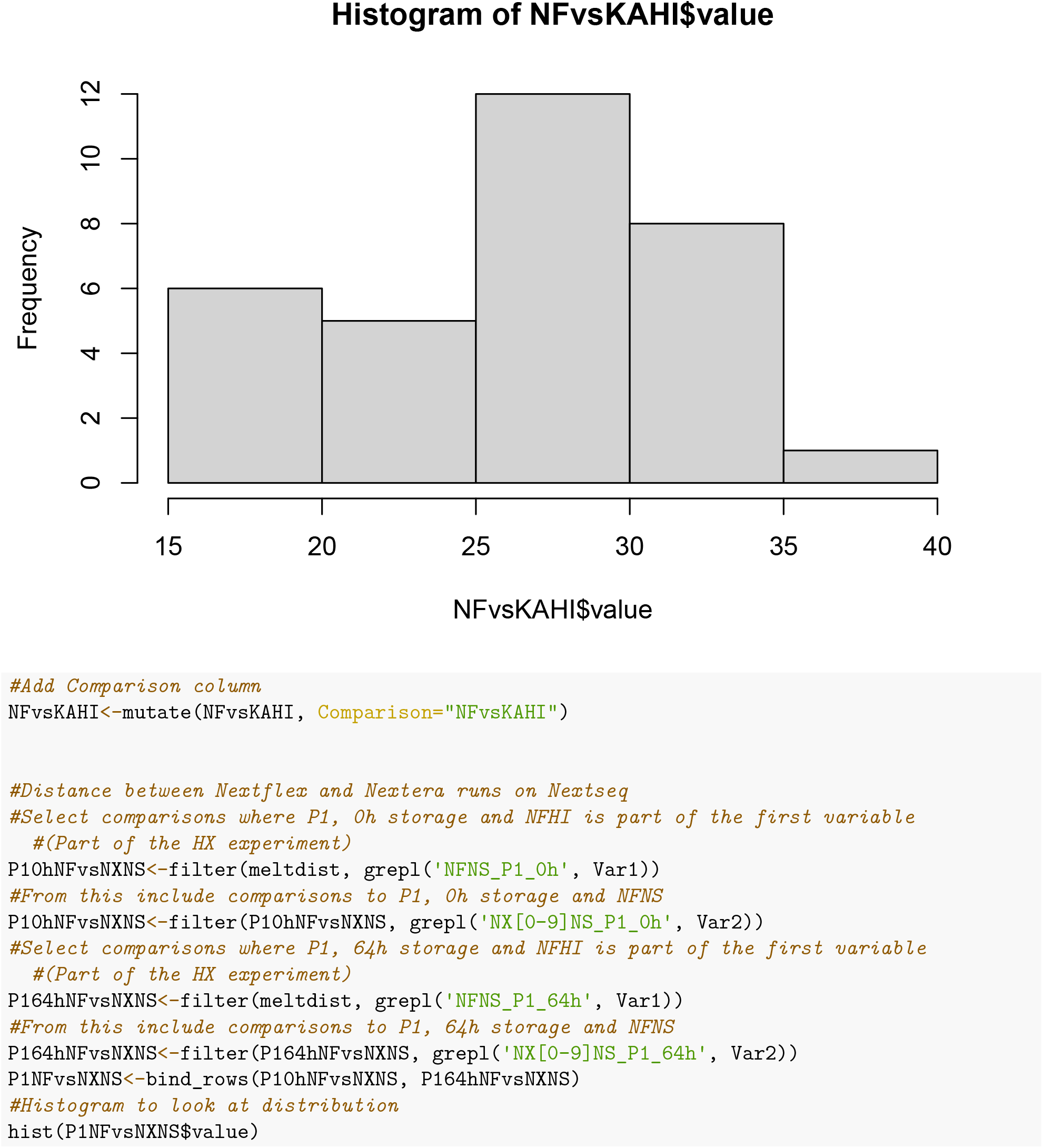
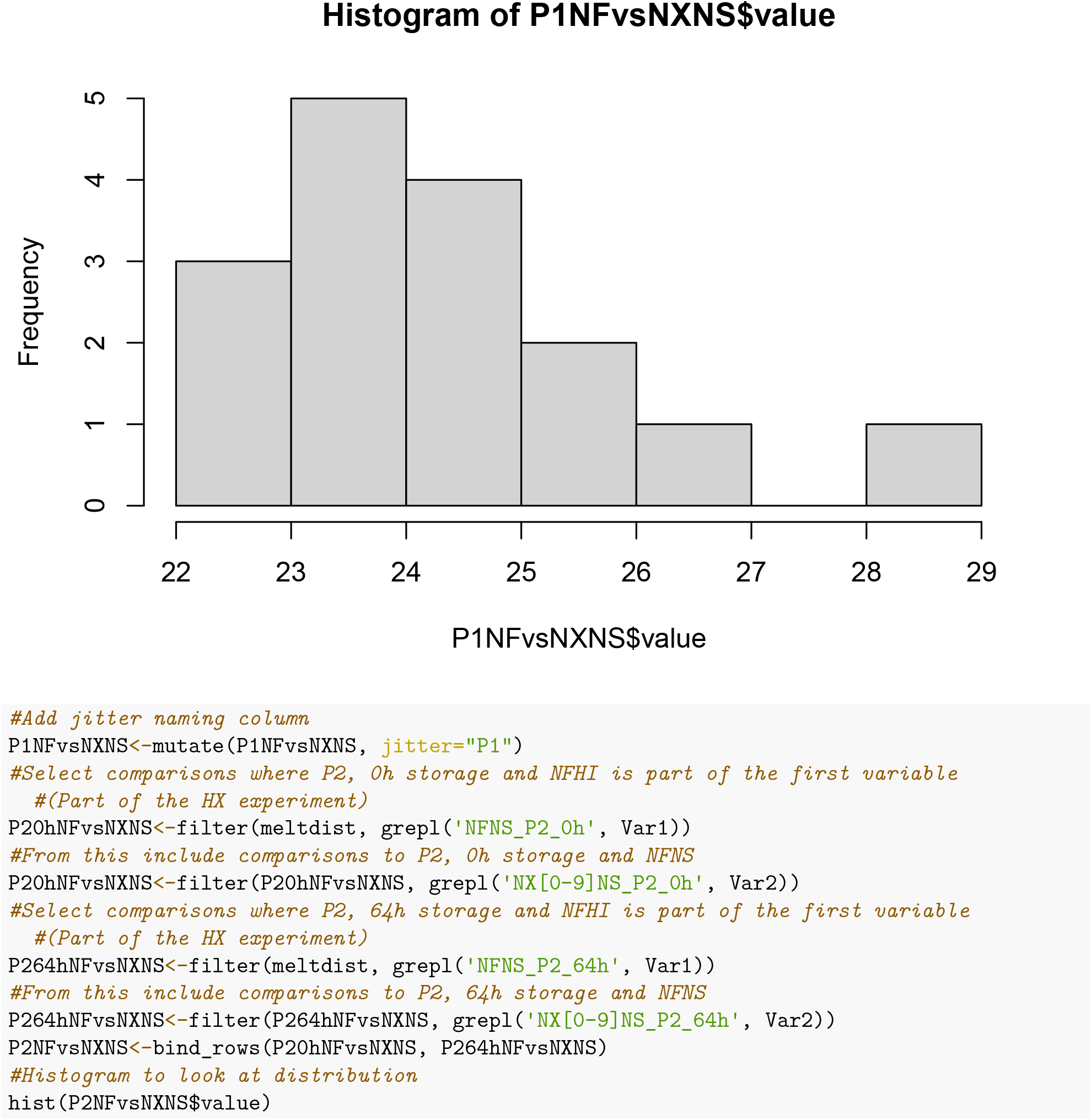
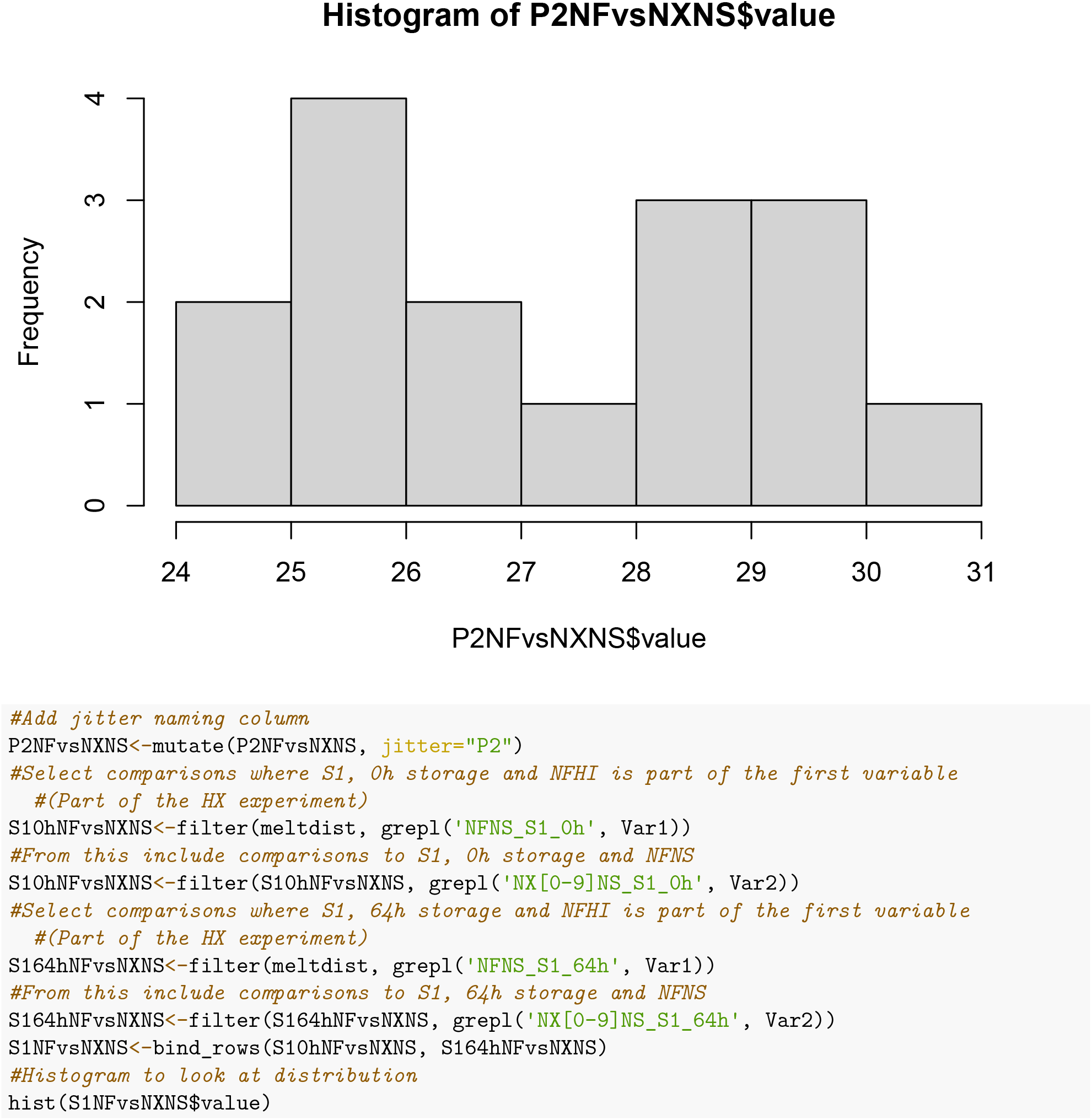
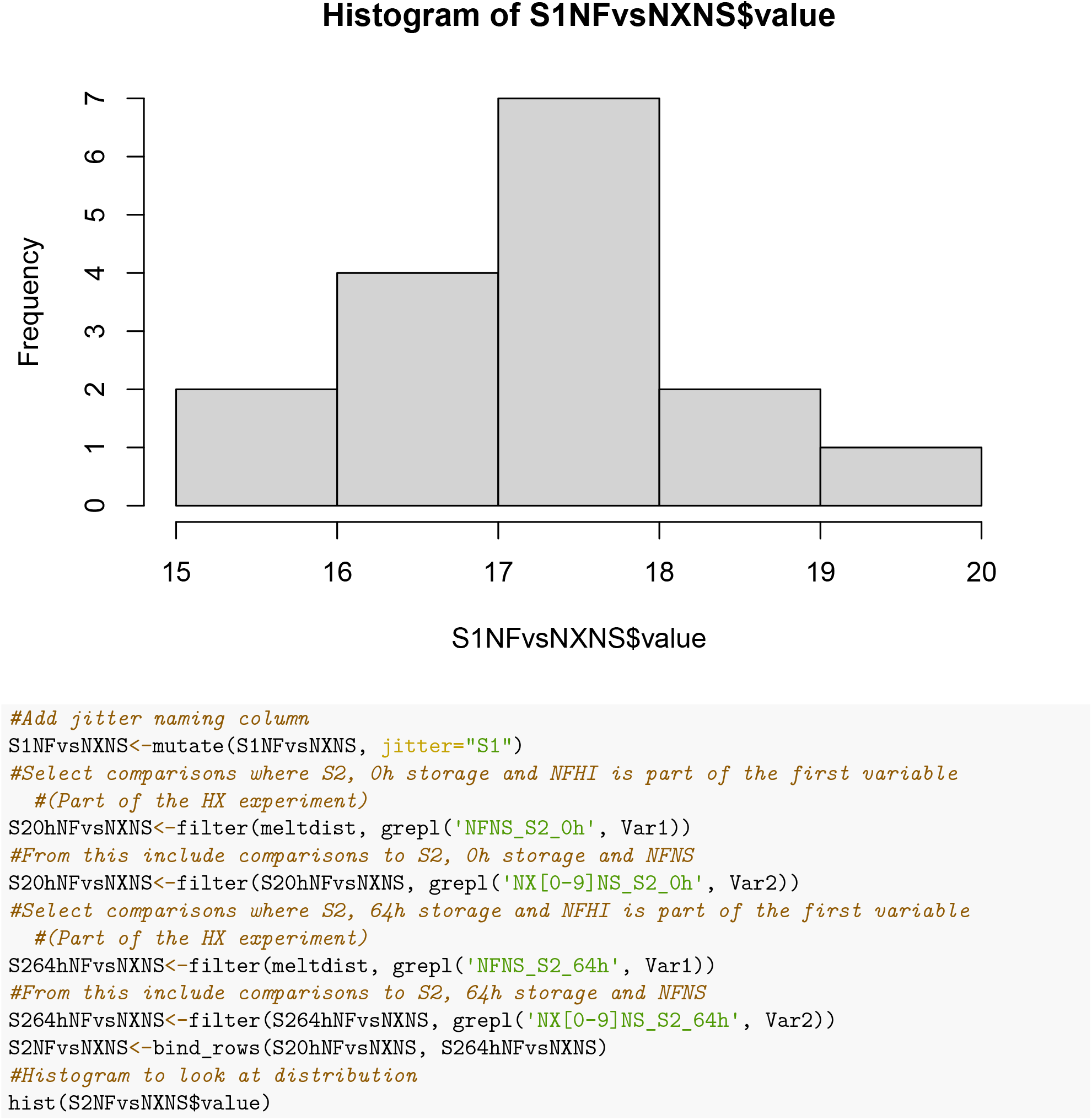
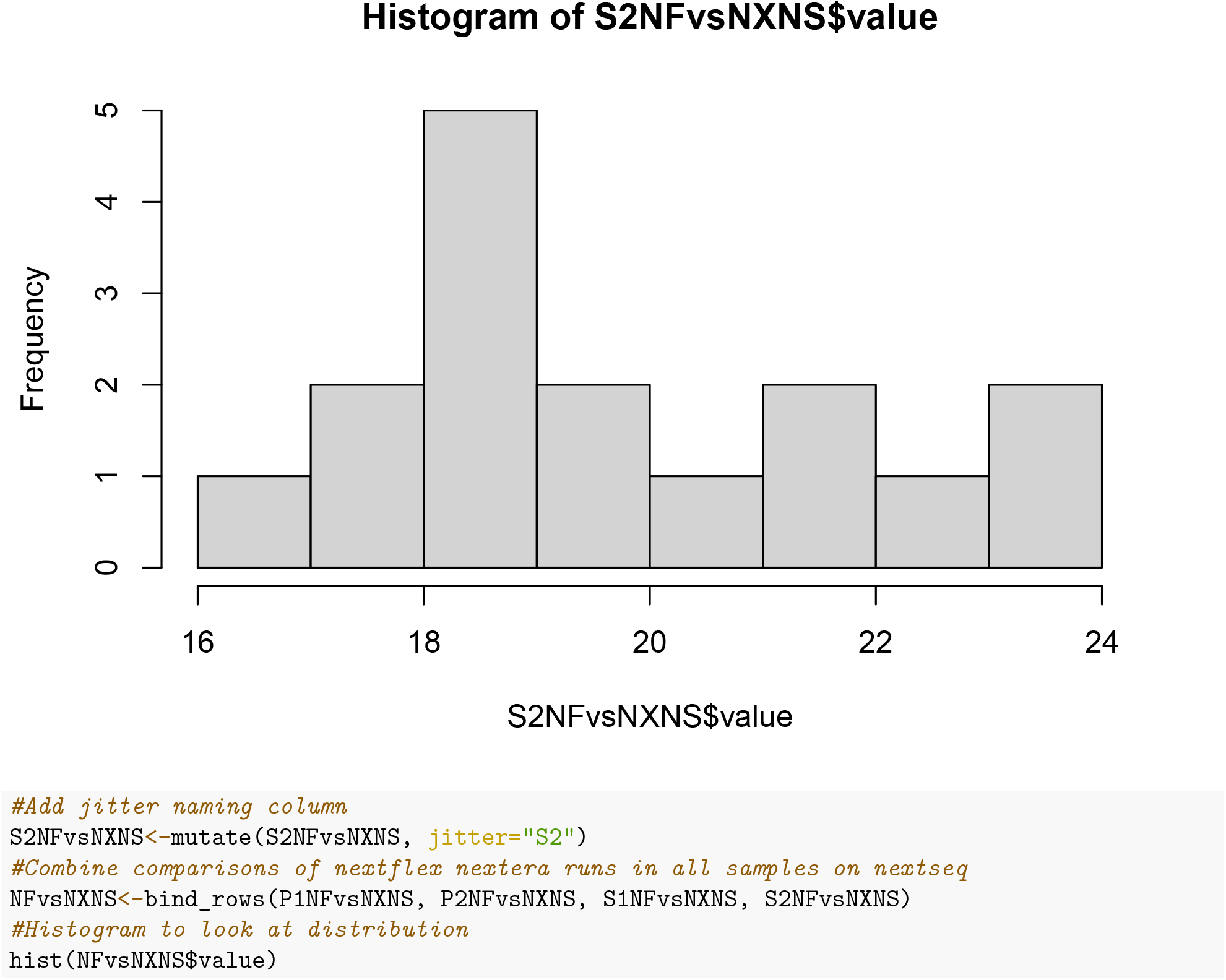
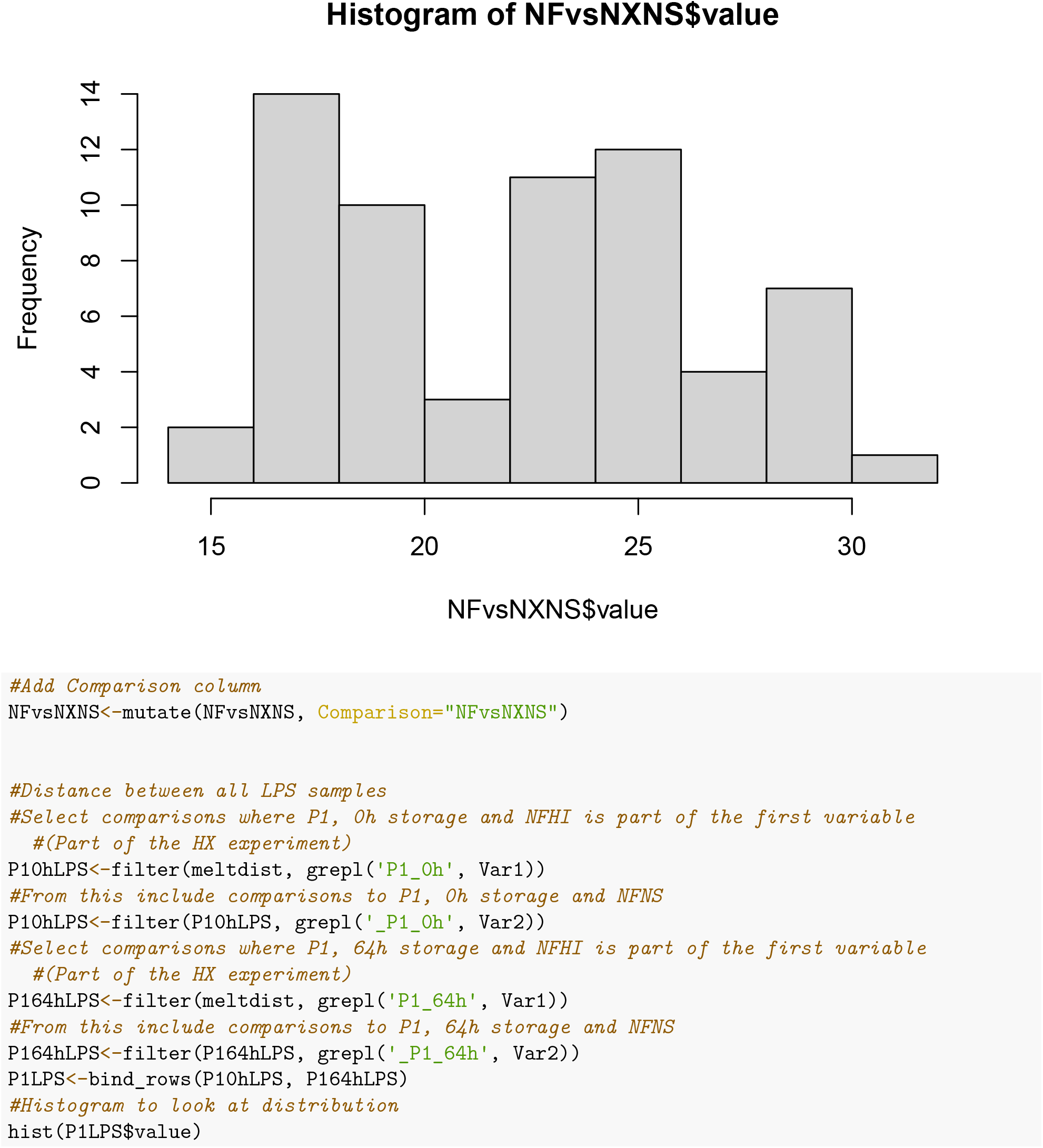
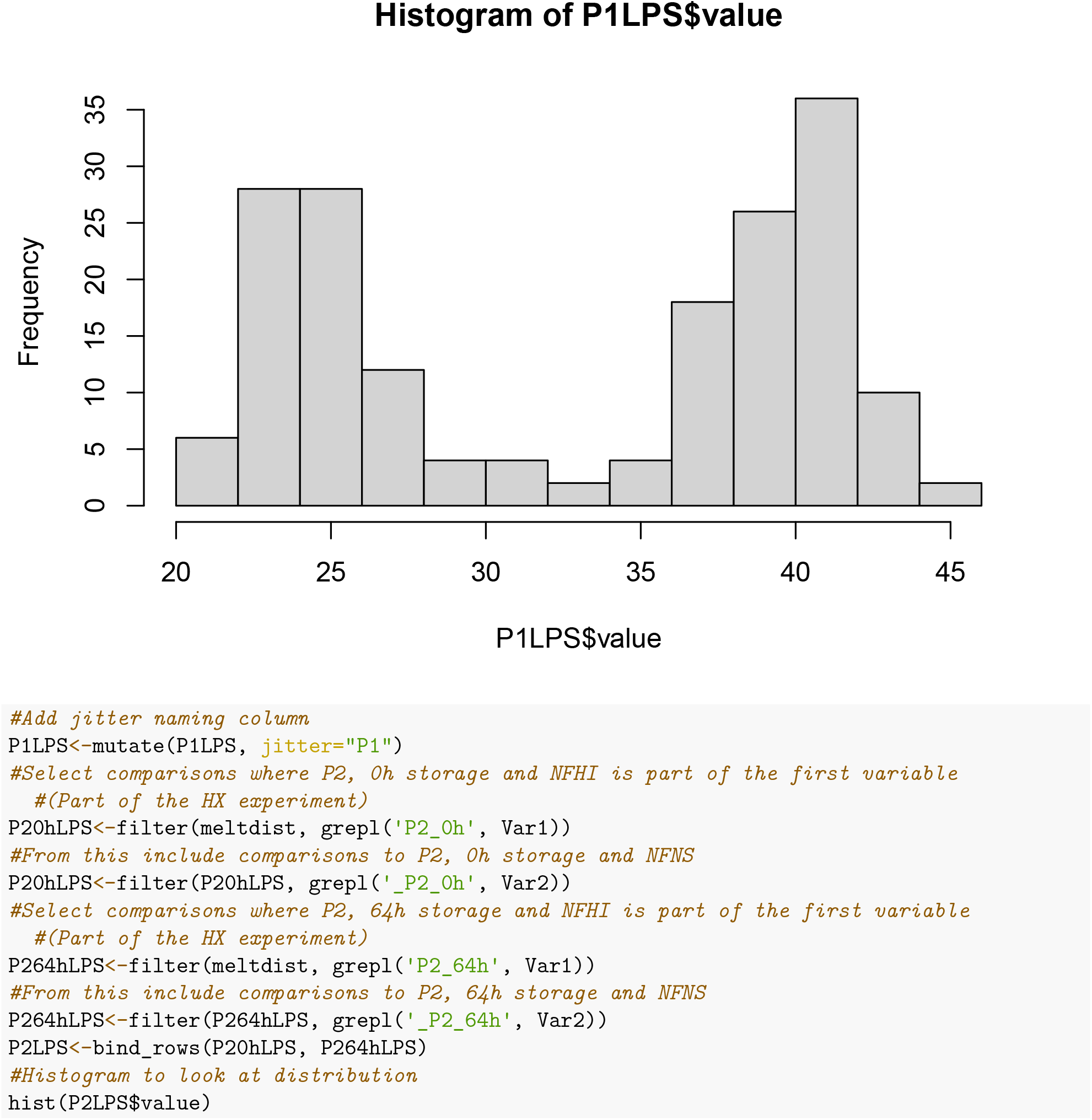
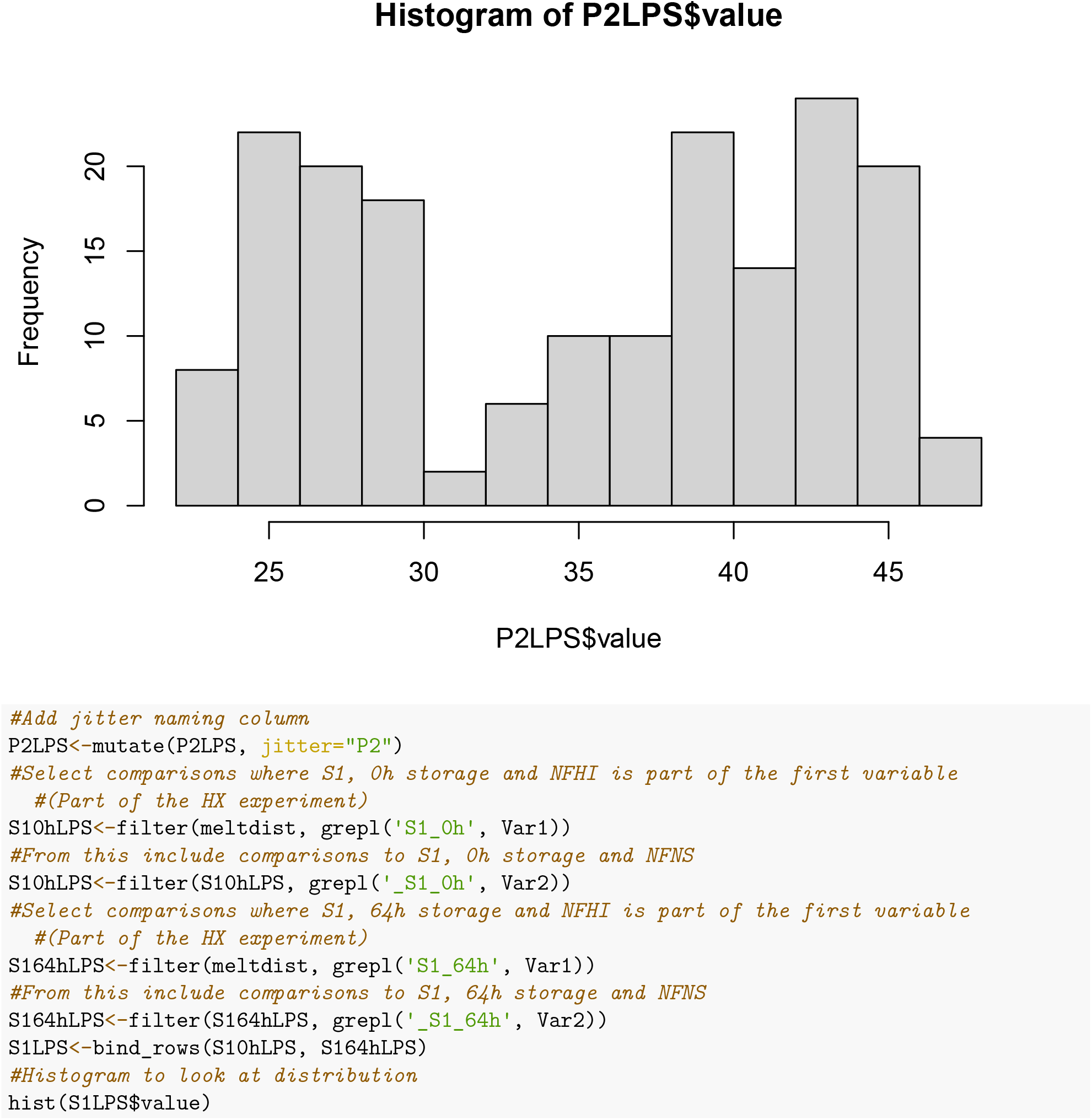
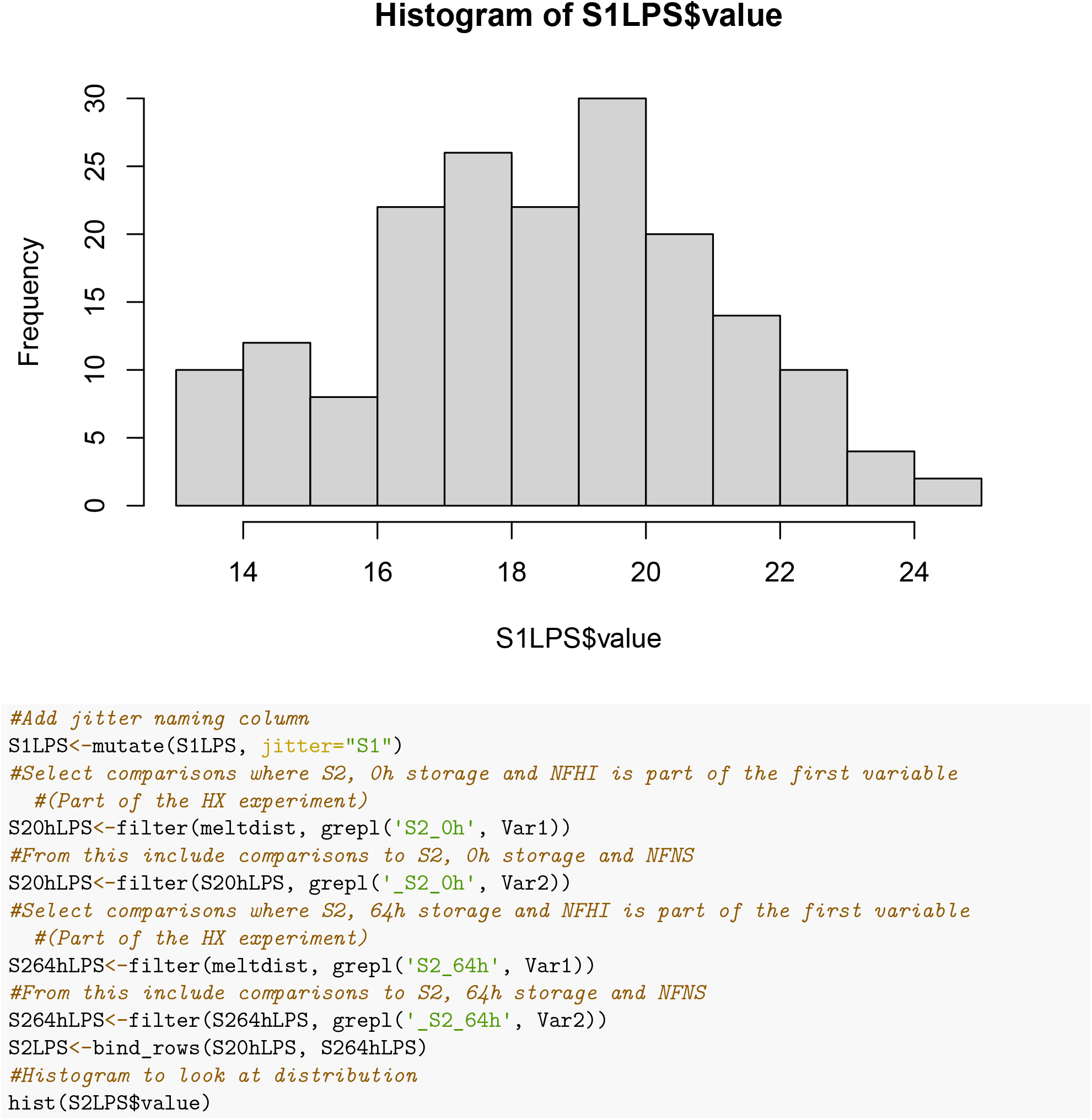
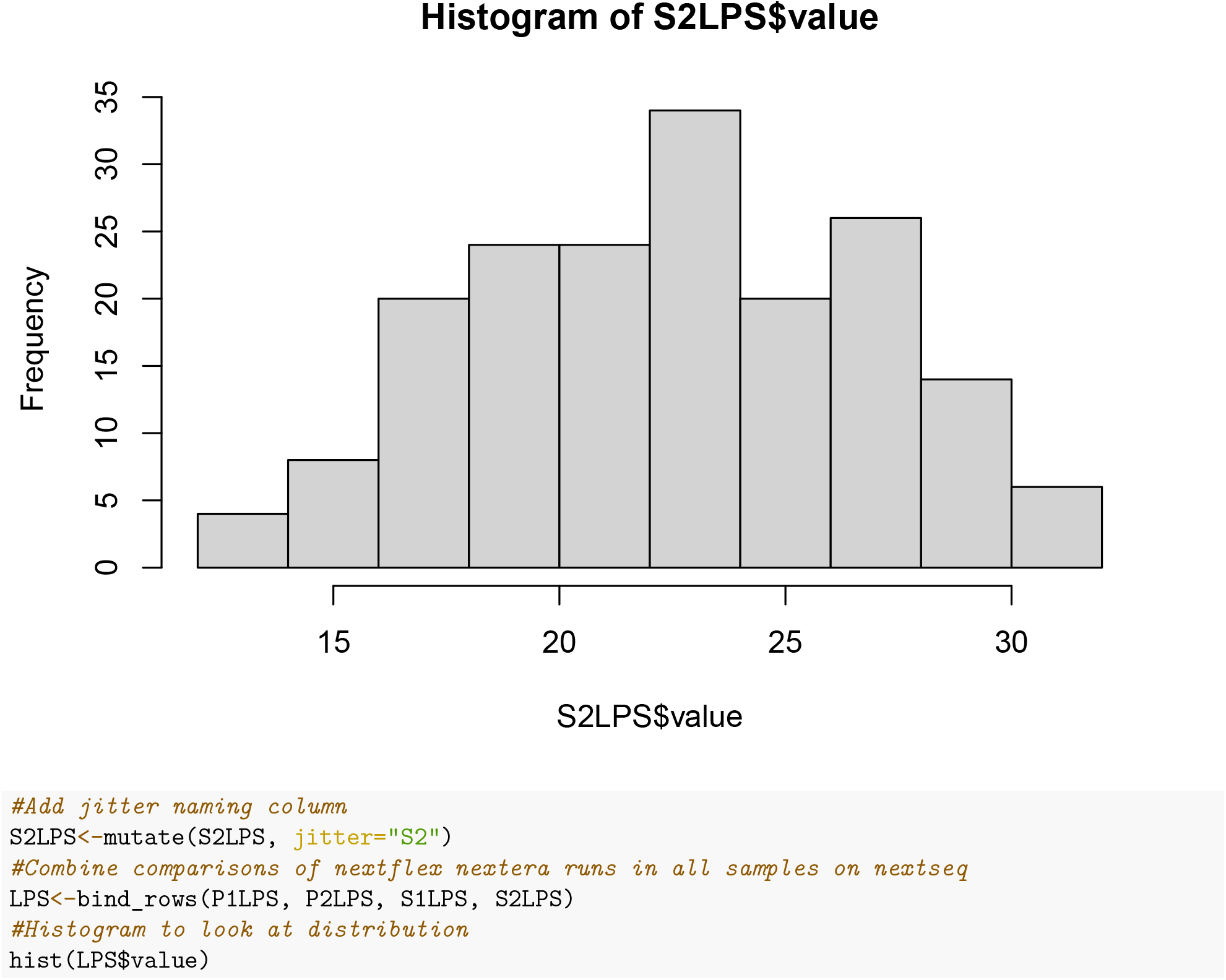
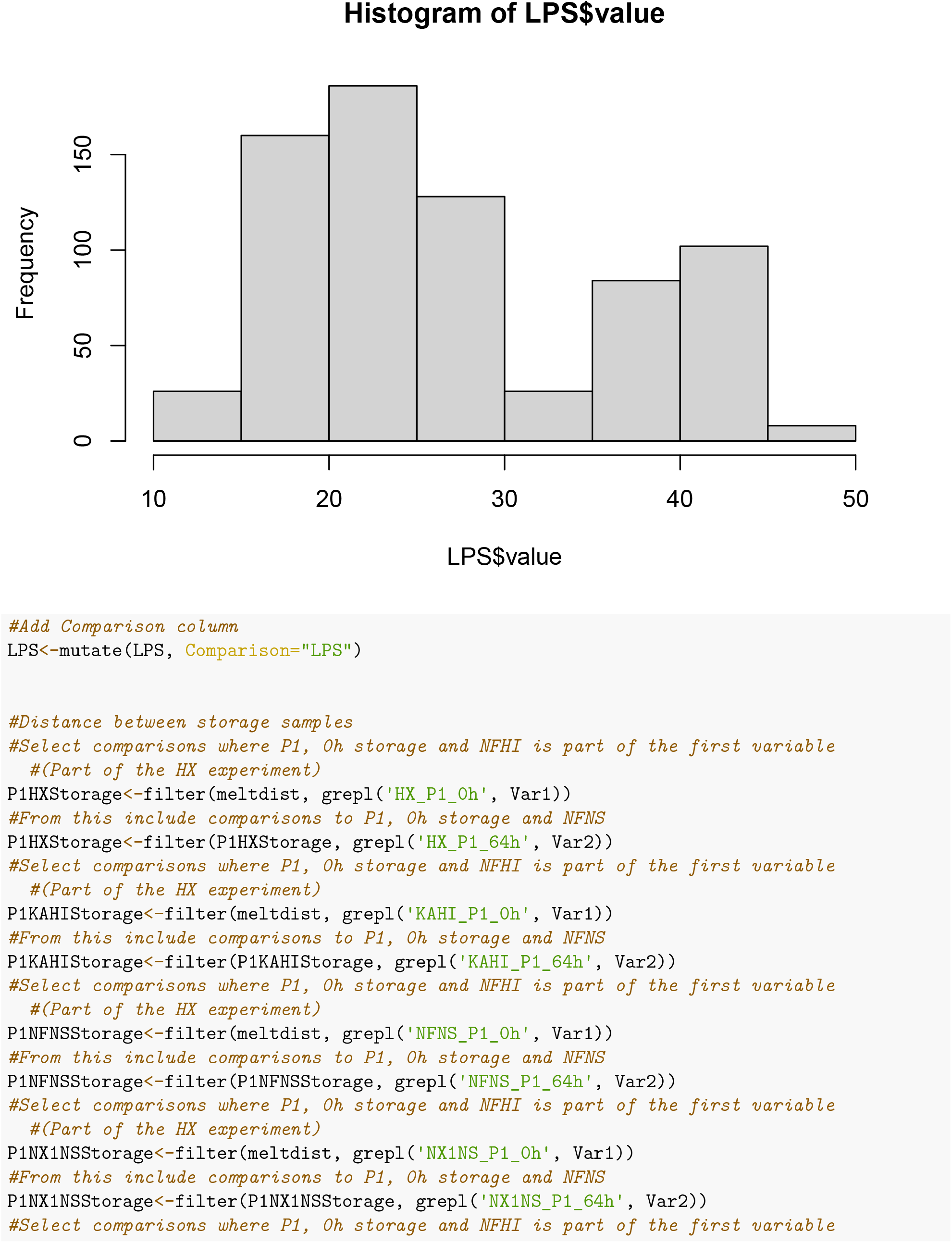
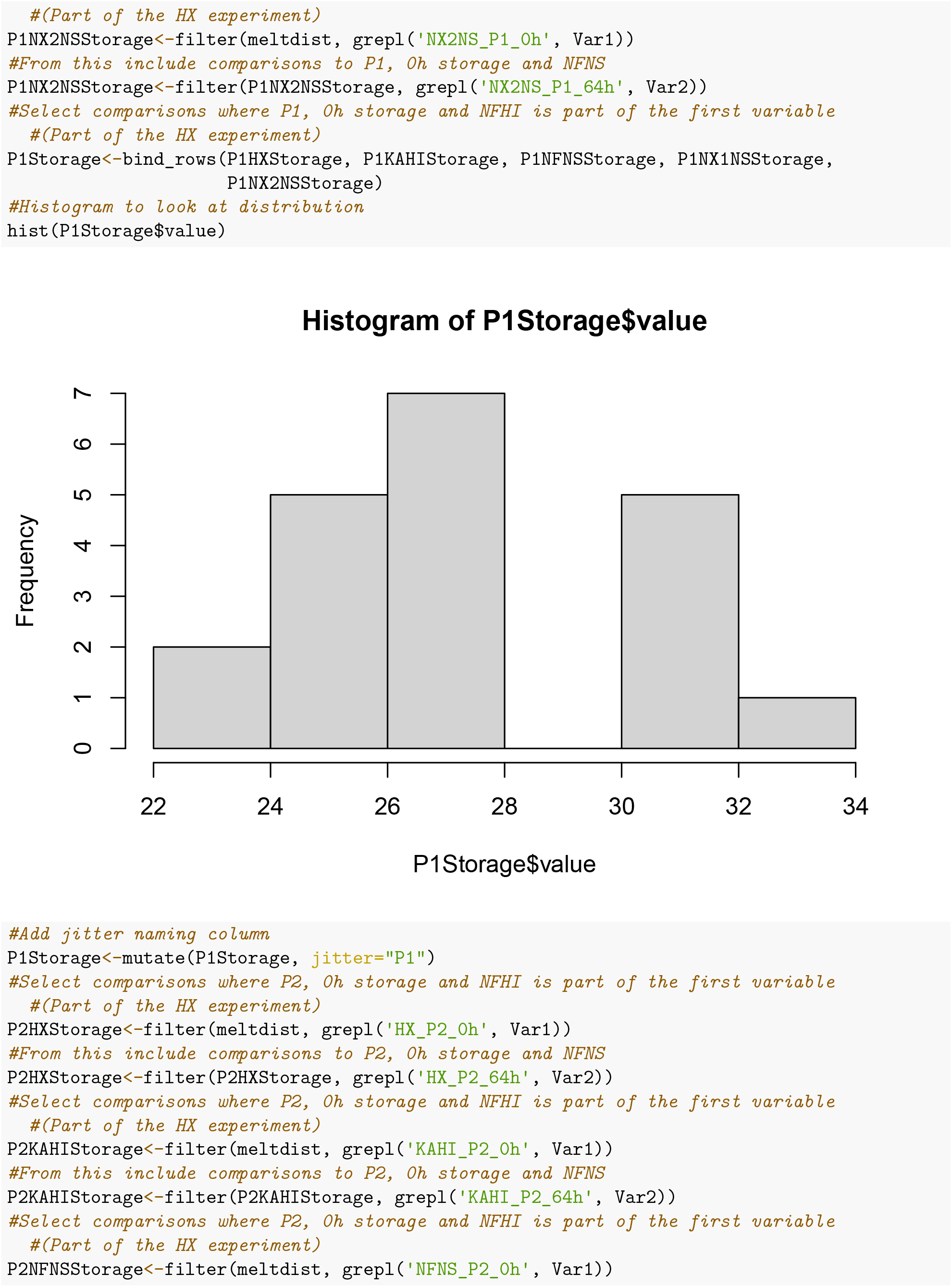
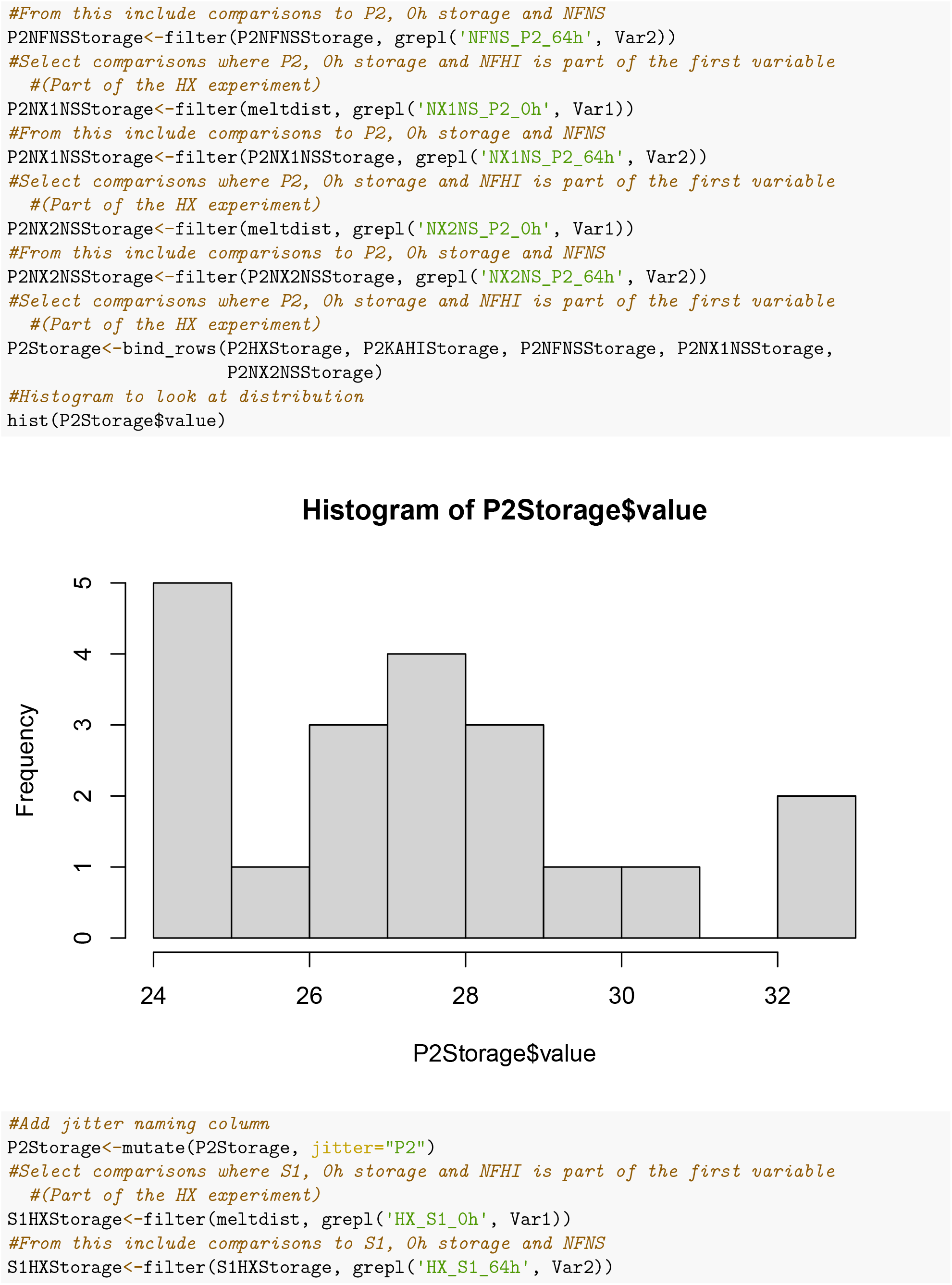
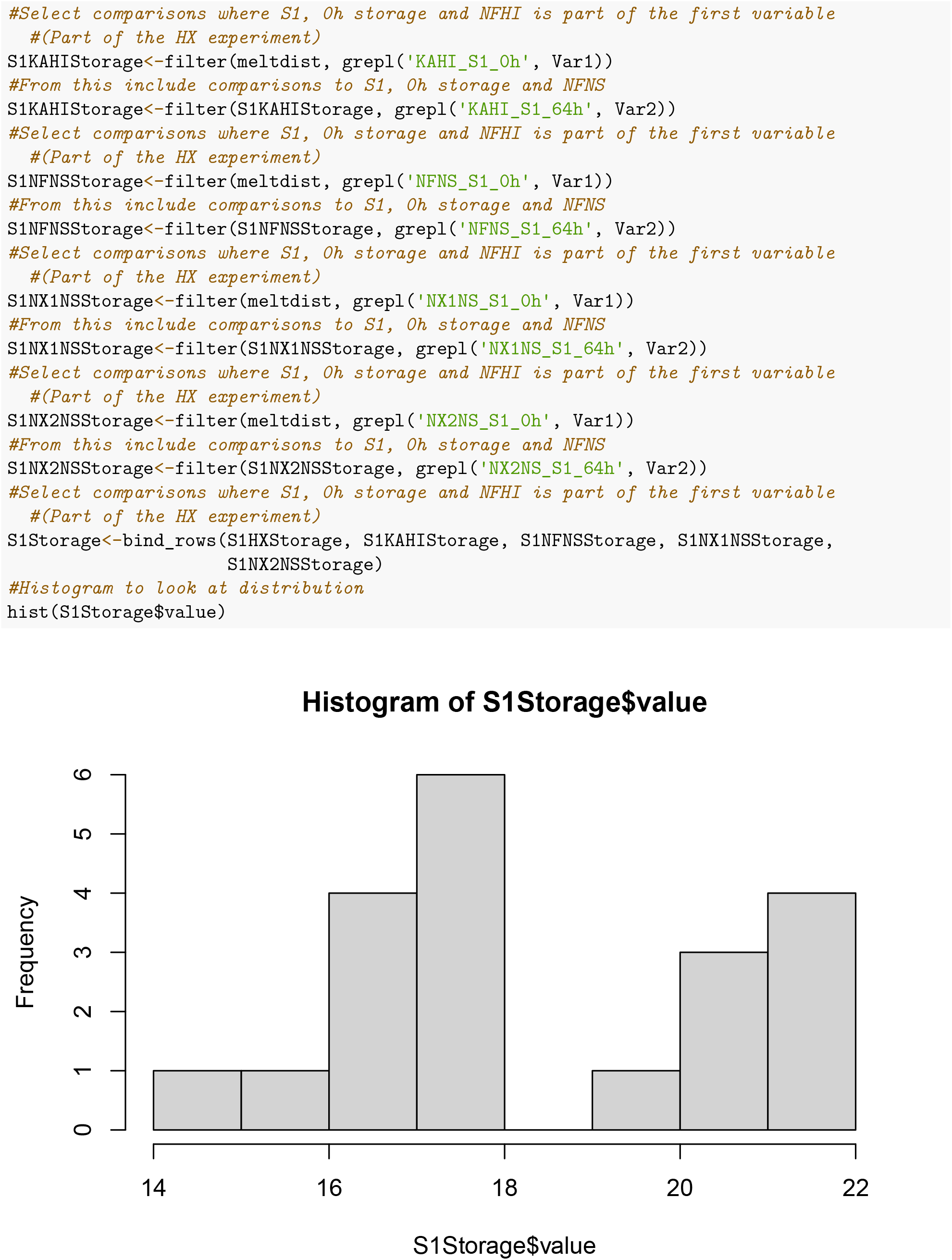
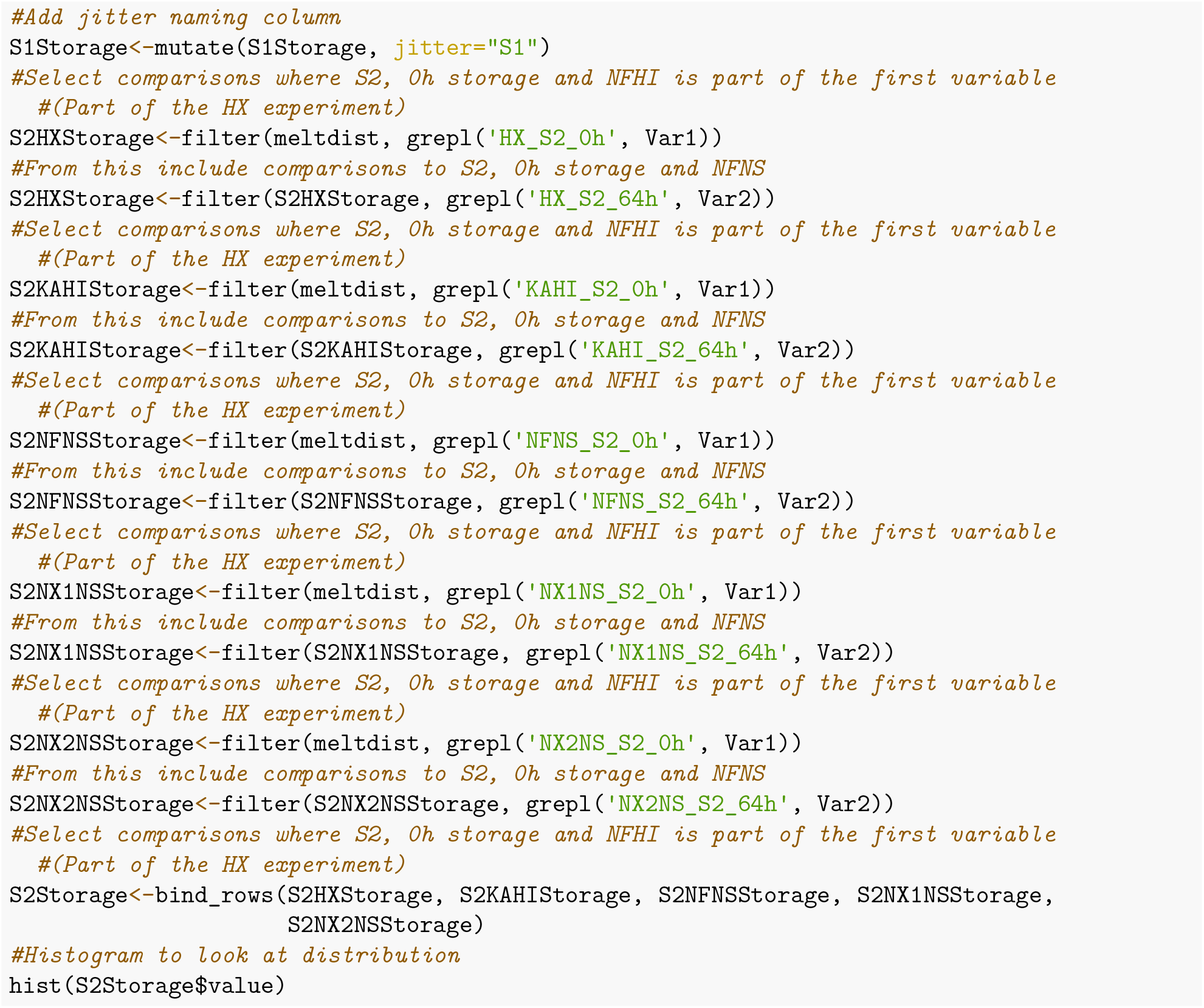
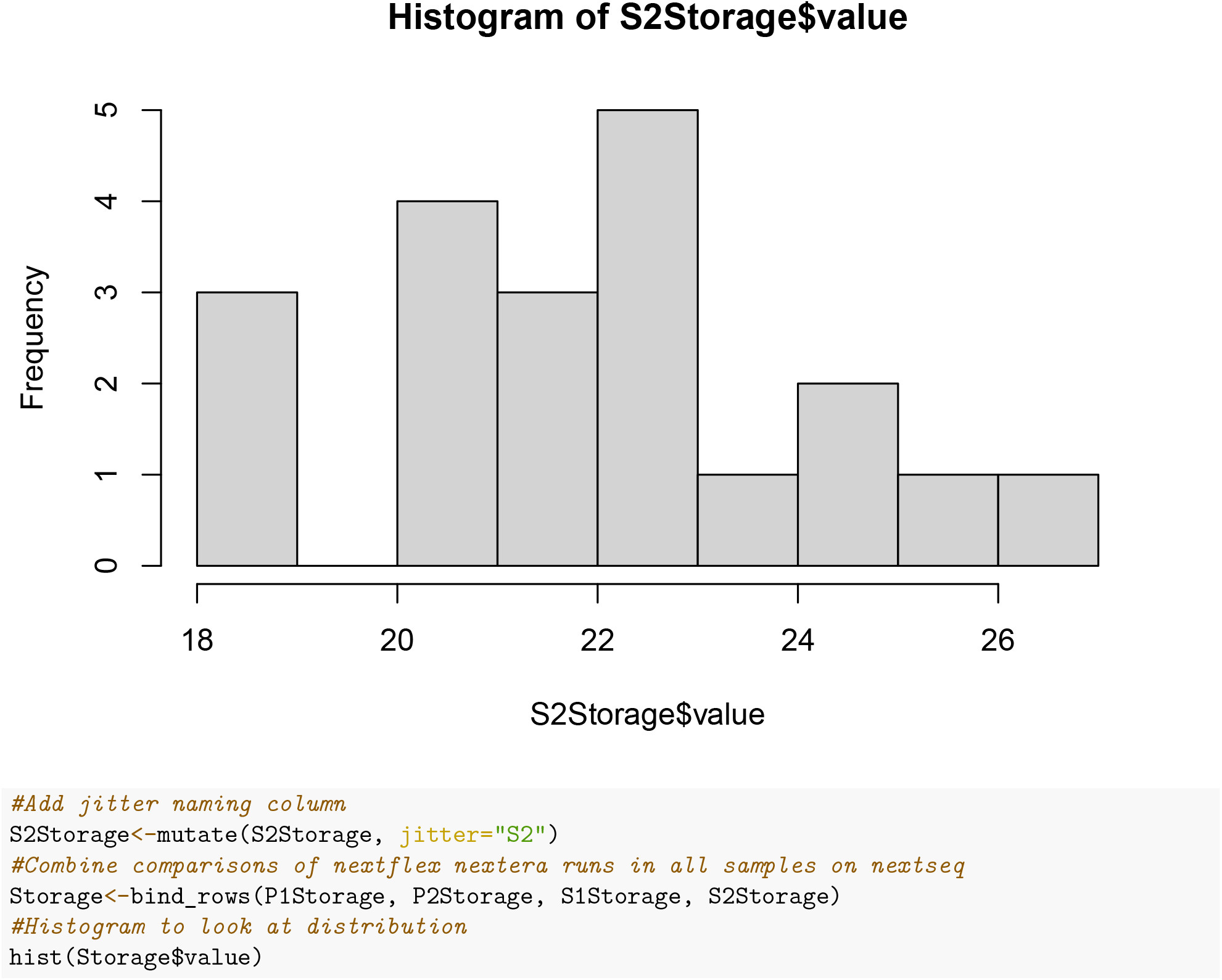
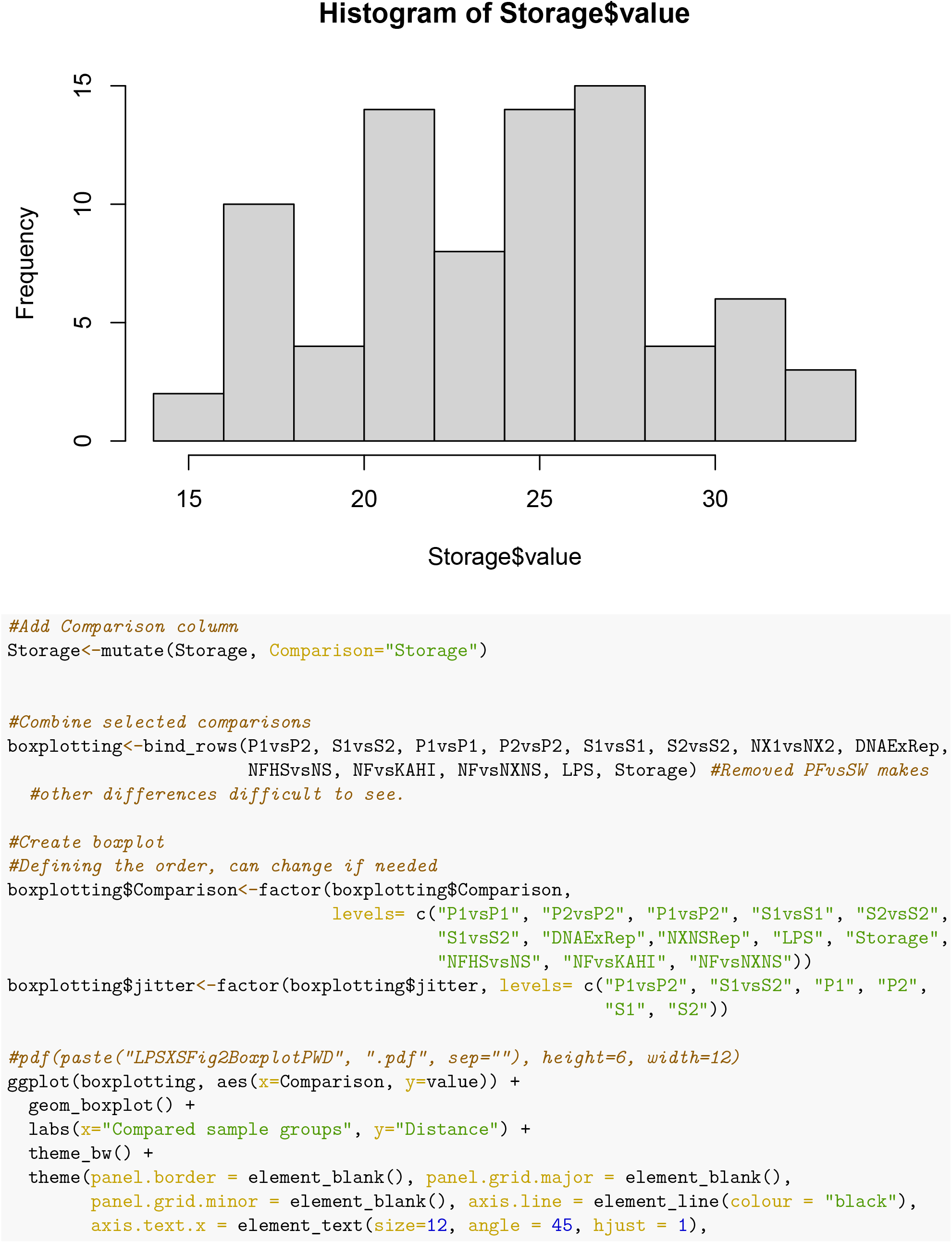
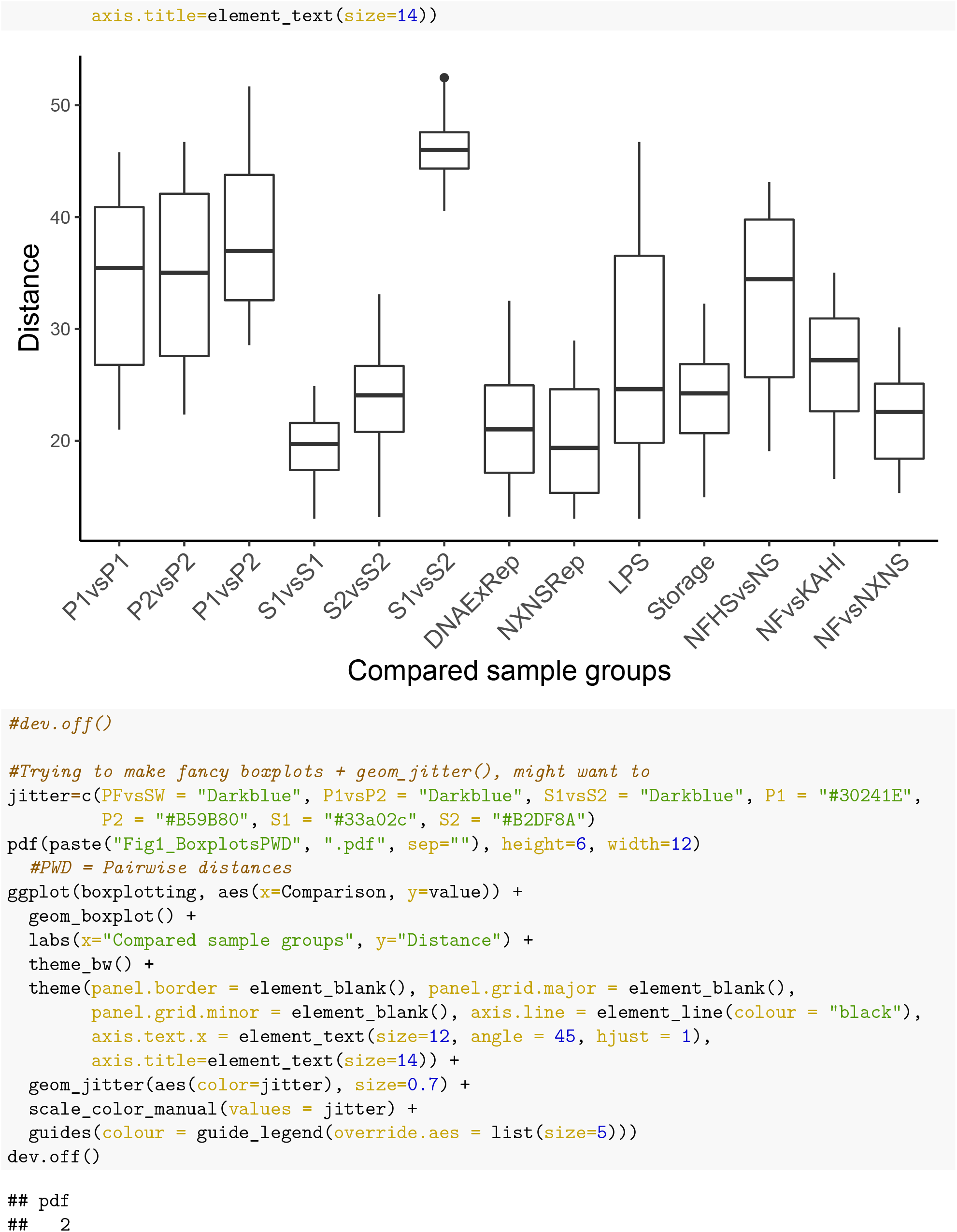

#### sPLS-DA

Used to find most important features for describing library preparation and sequencing platform. Overview represented as S4_Fig

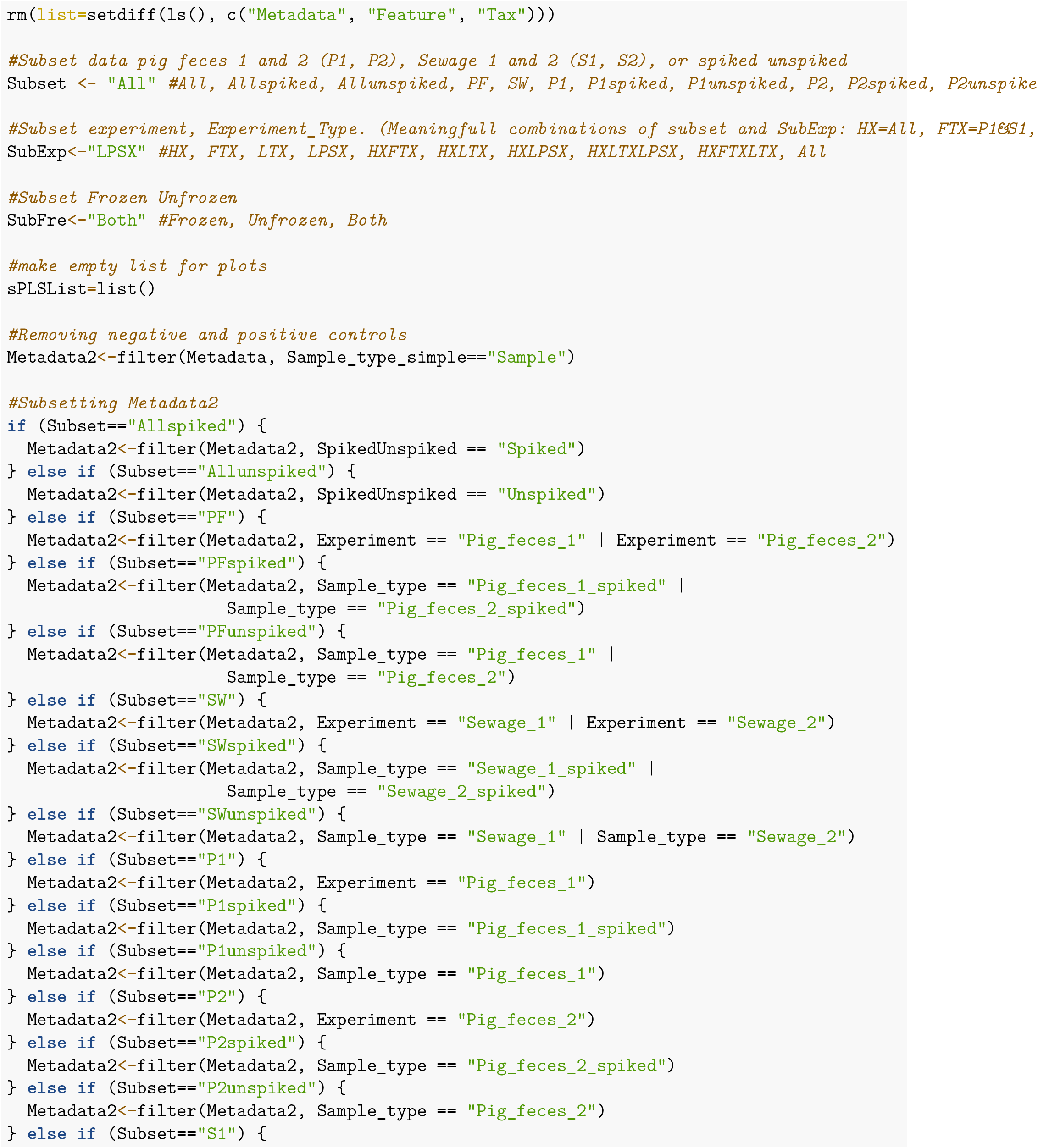
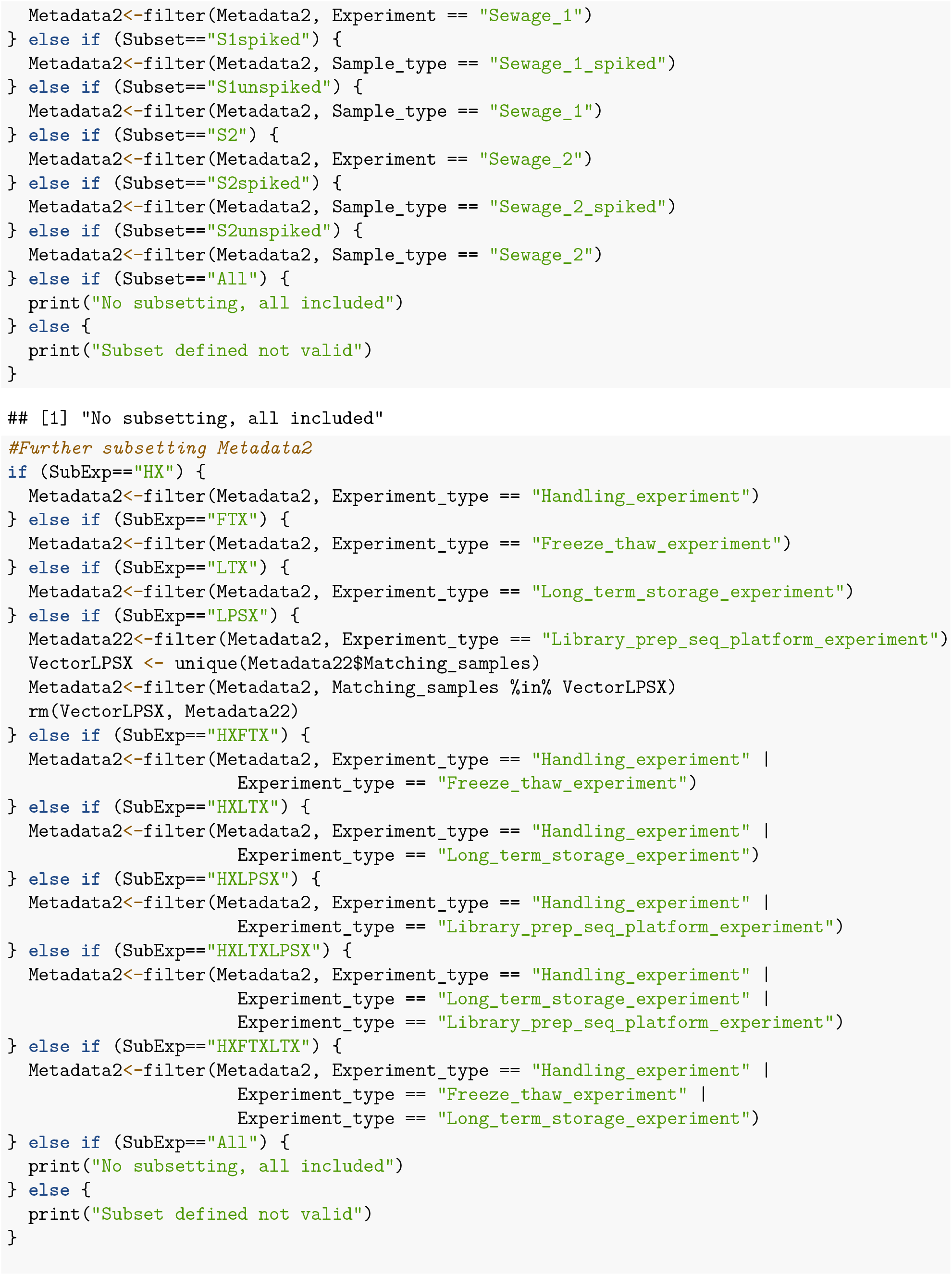
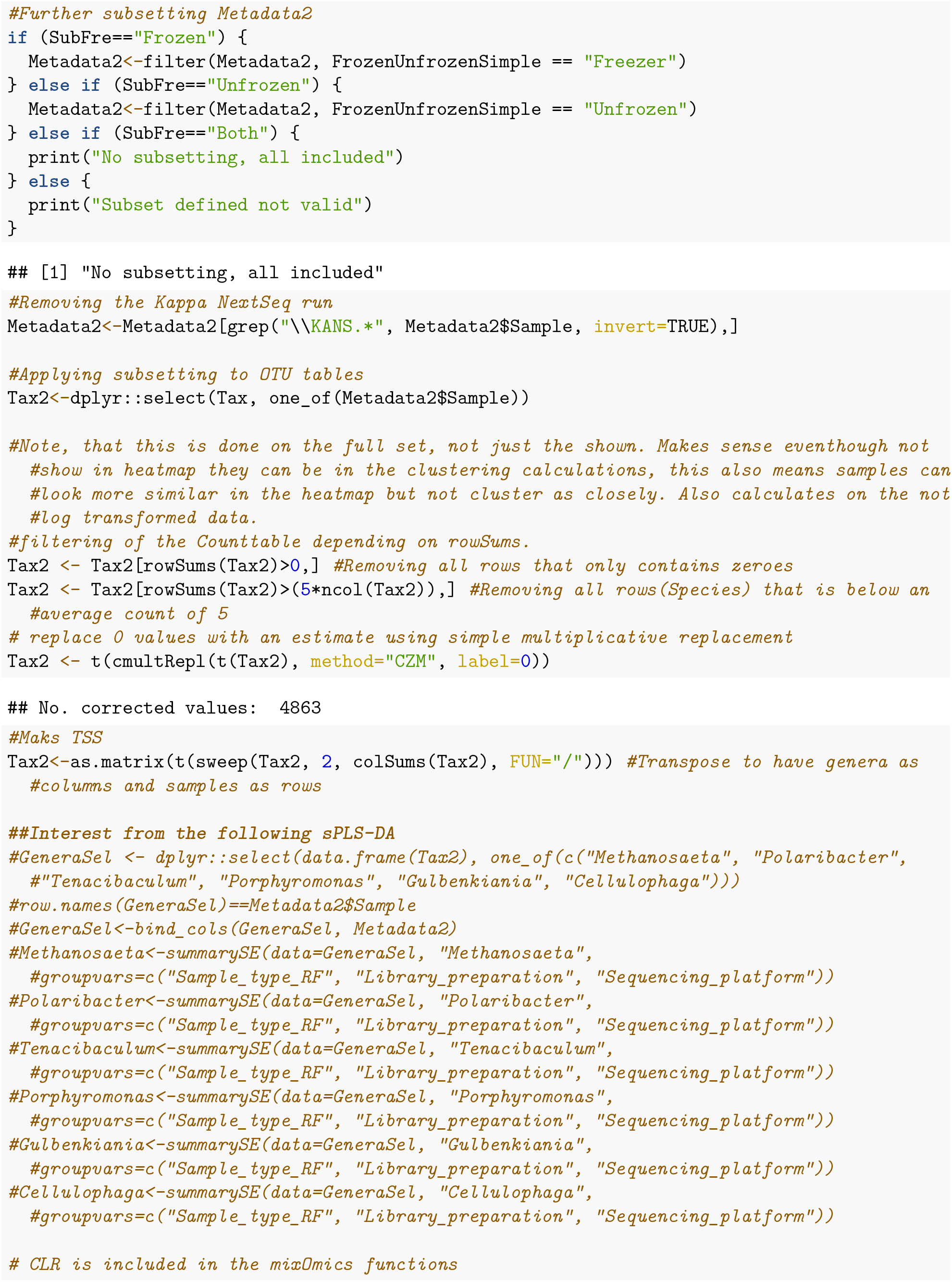
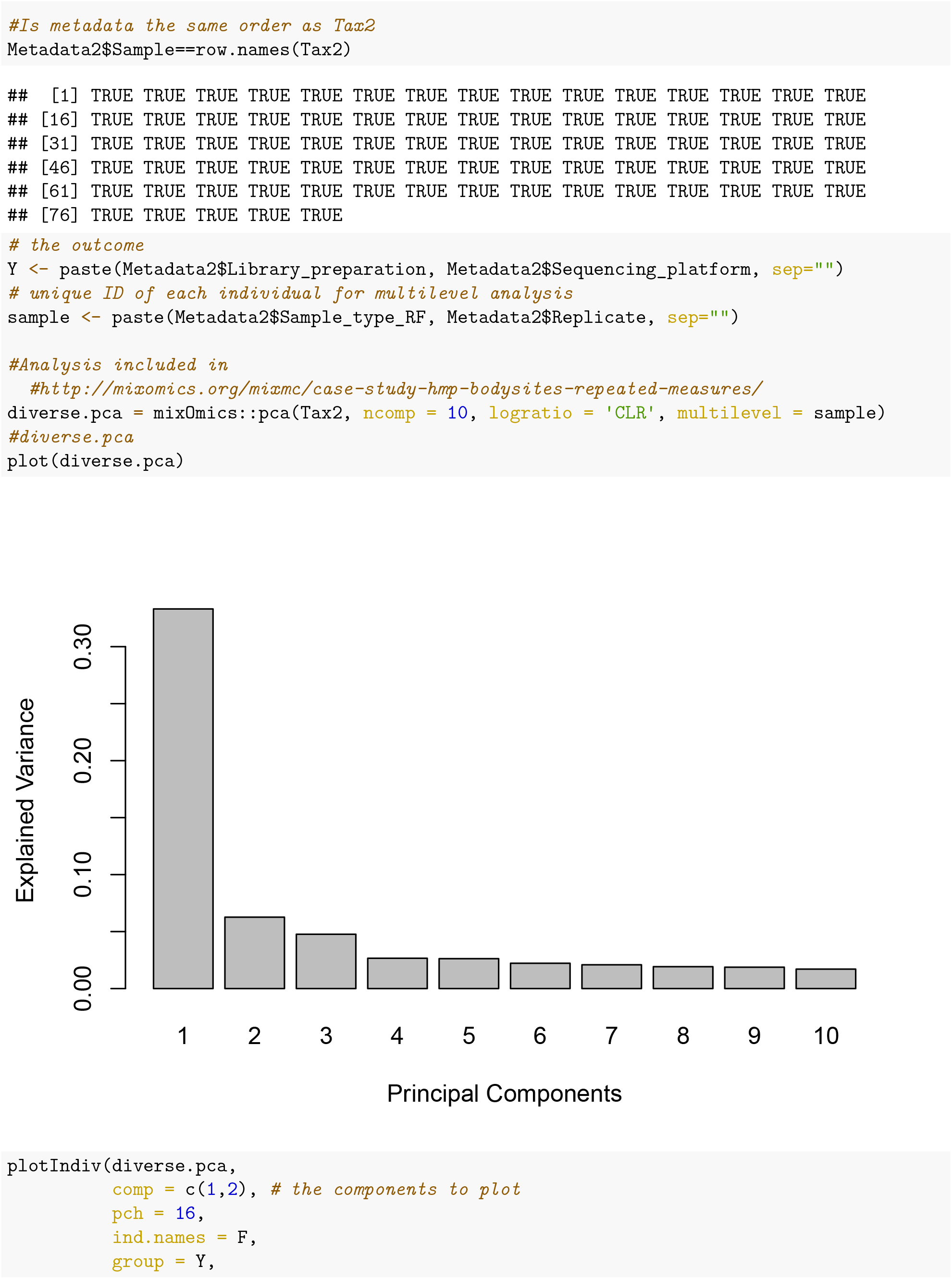
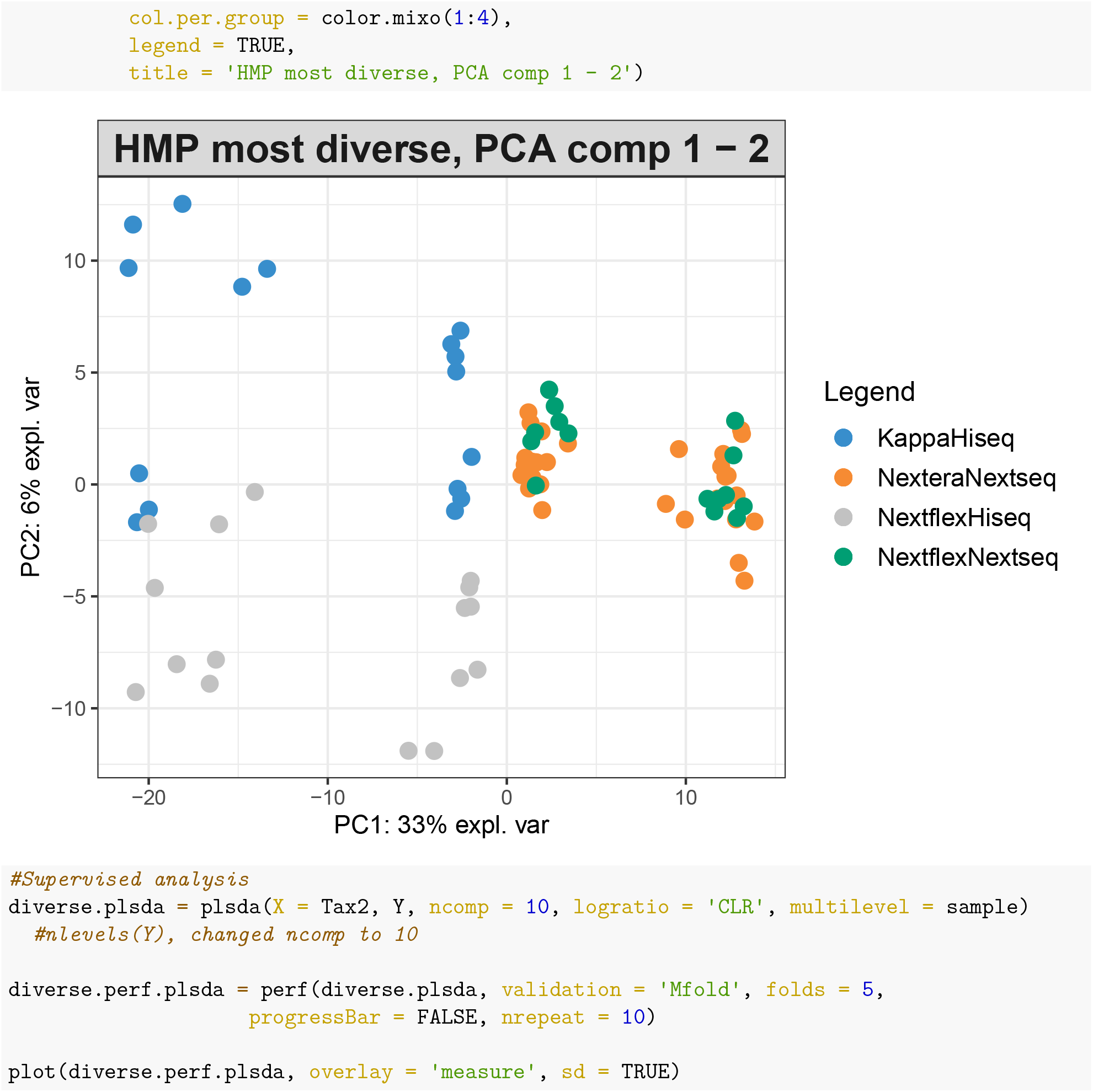
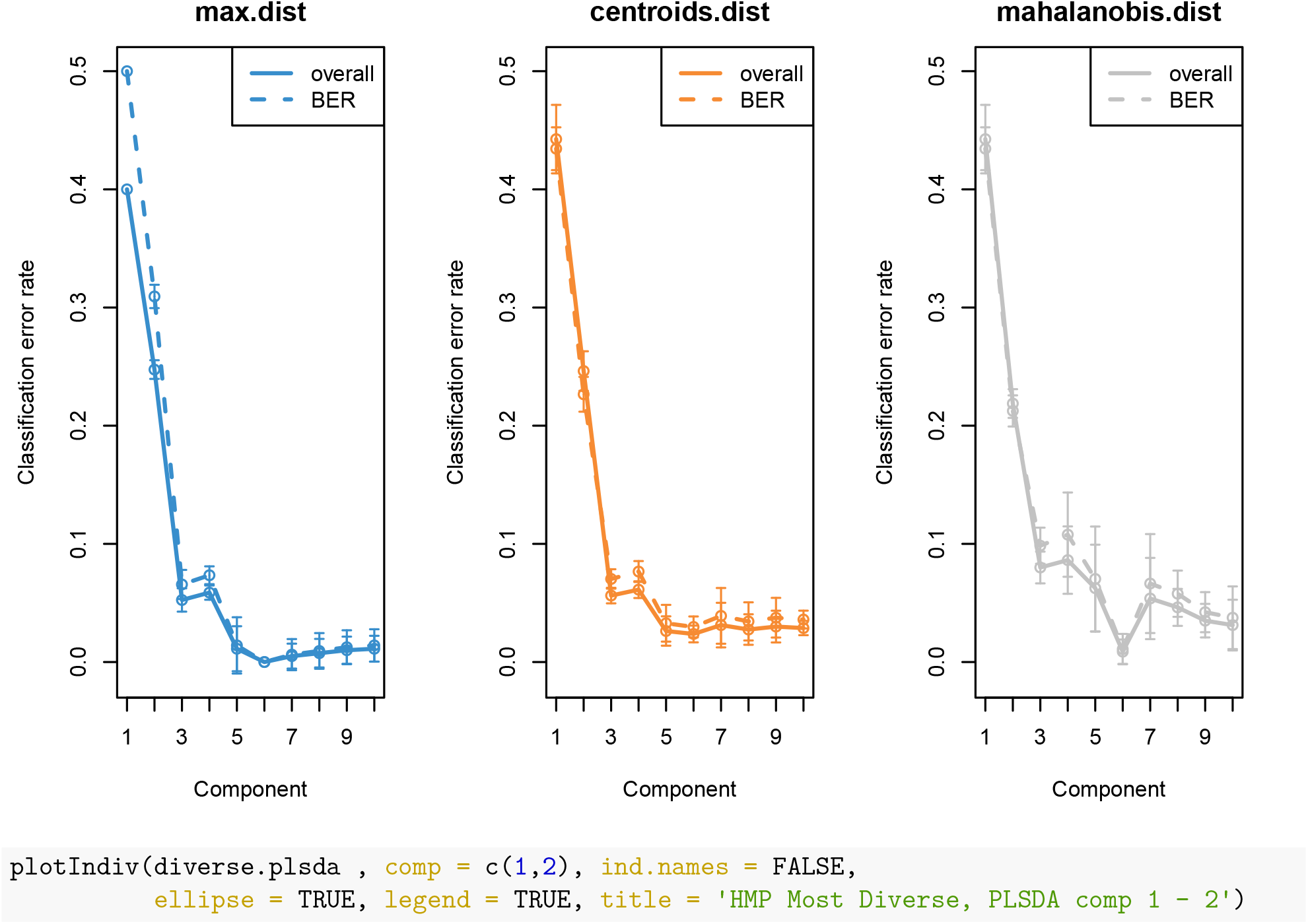
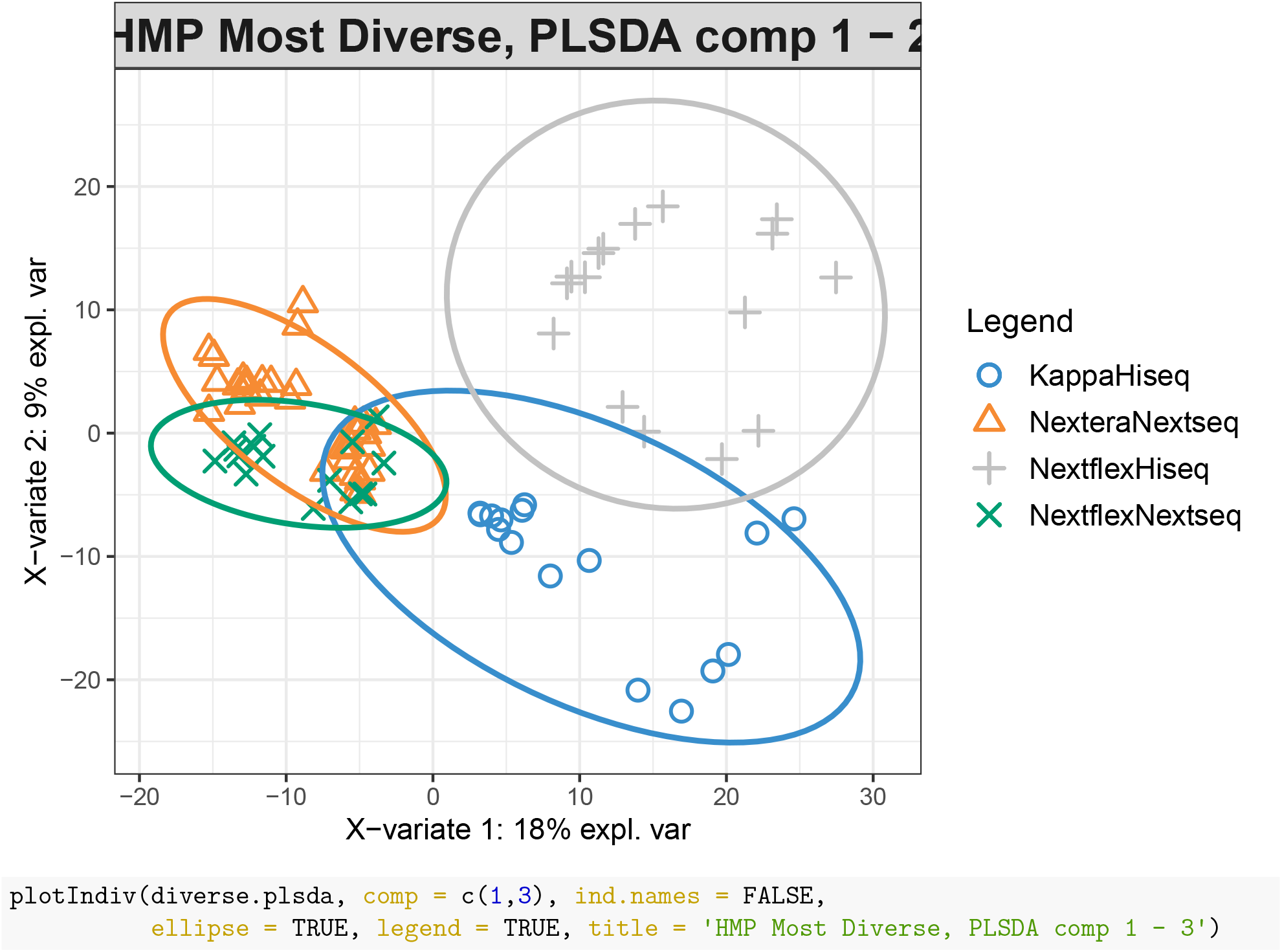
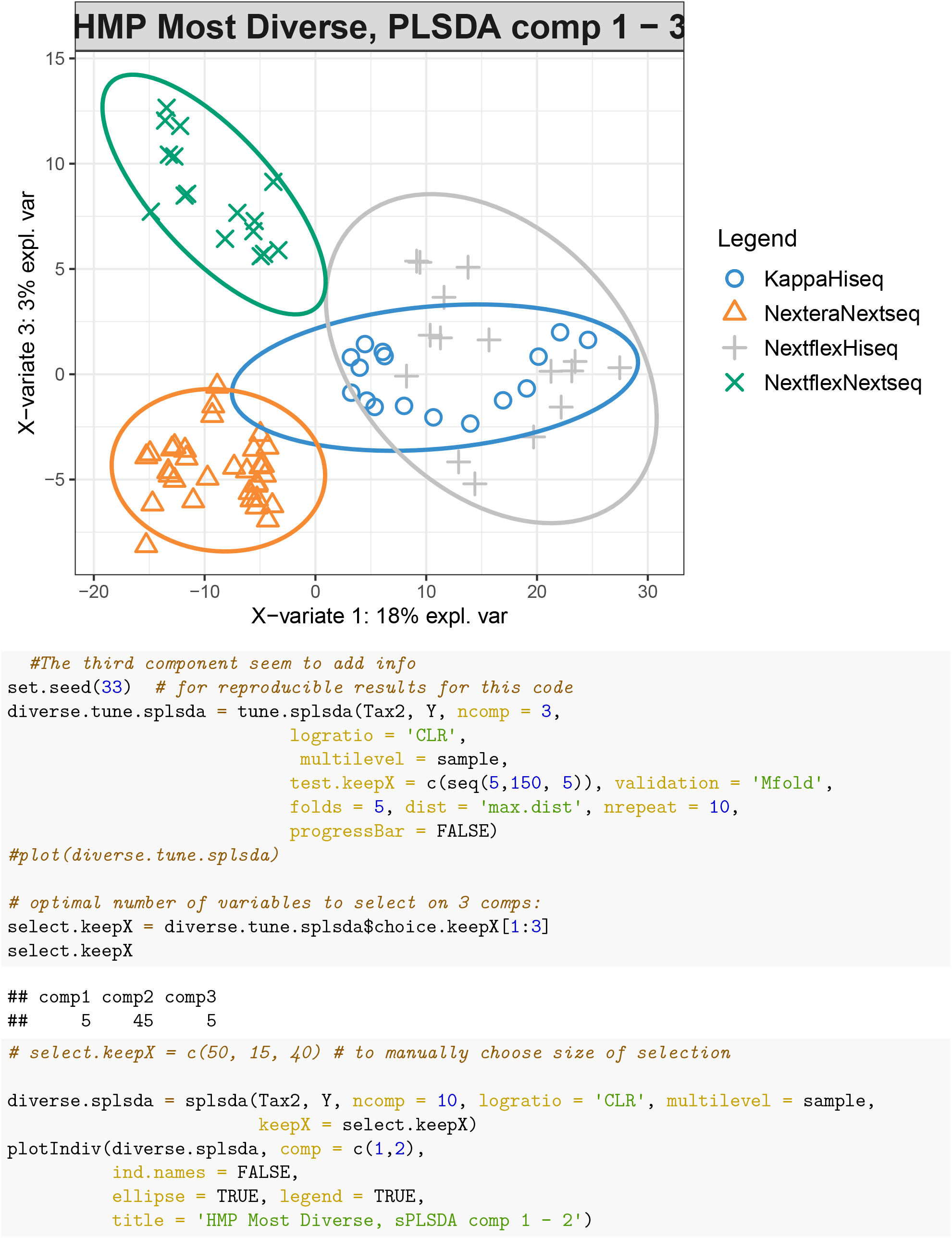
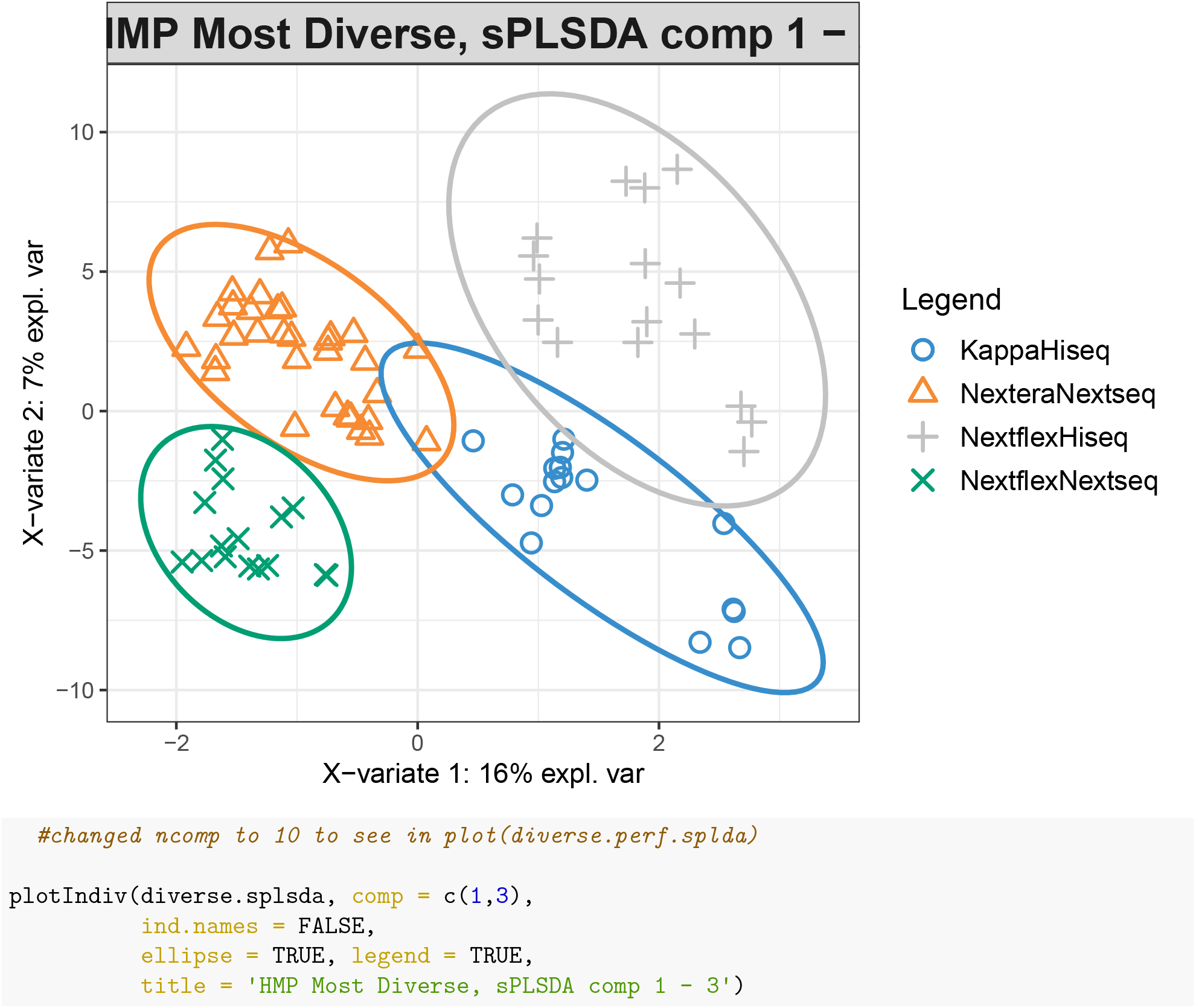
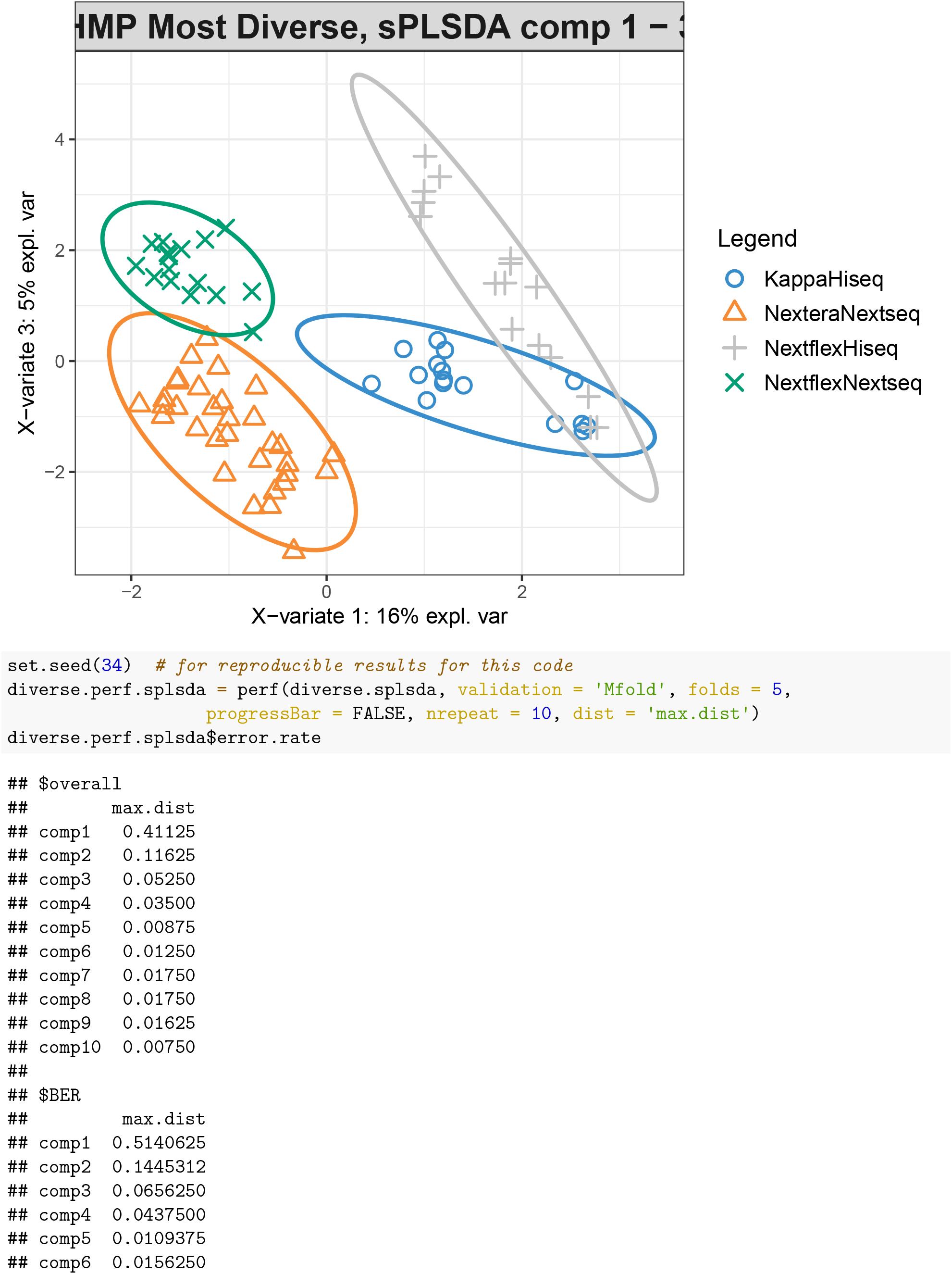
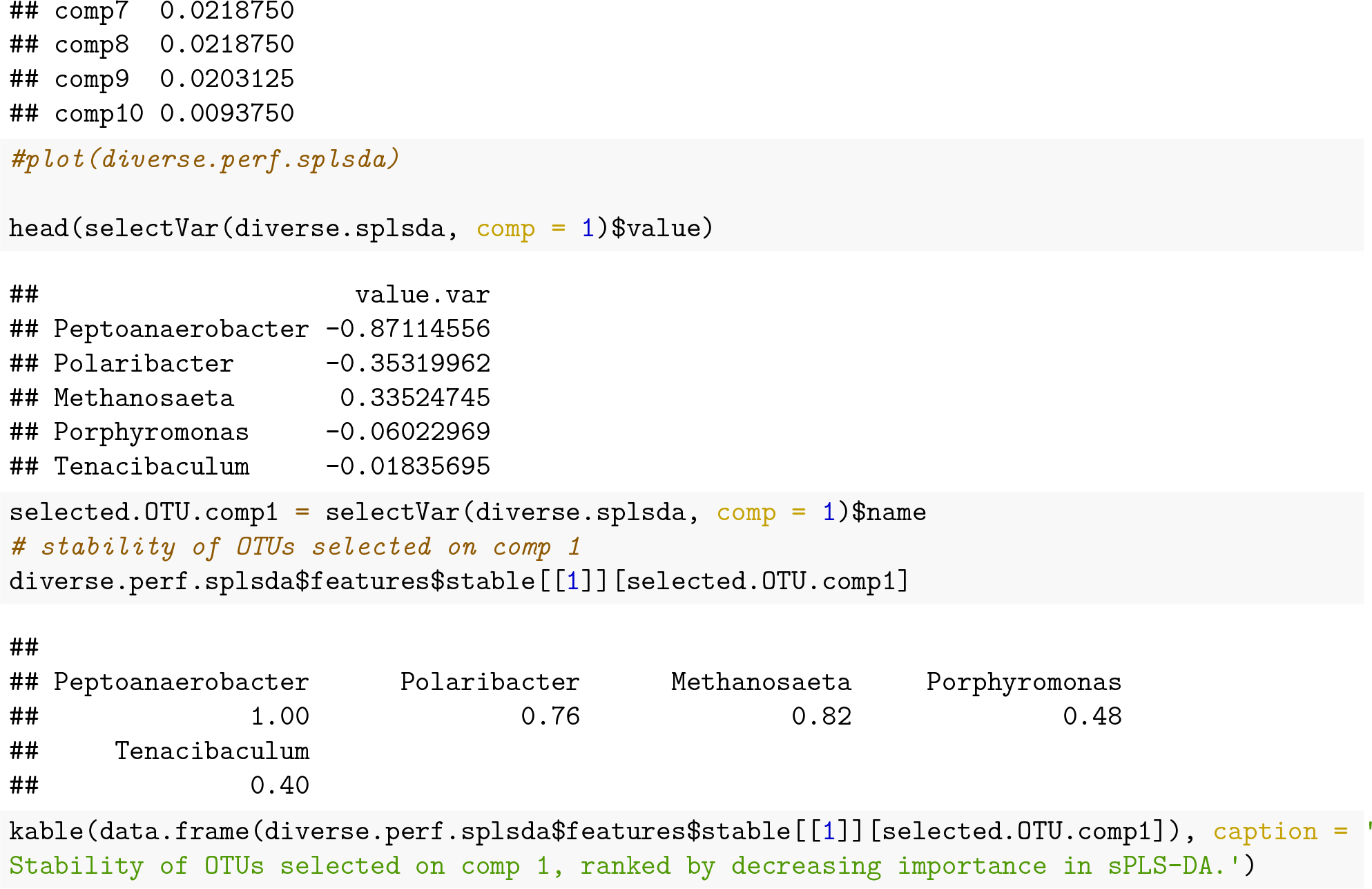

**Table 1:** Stability of OTUs selected on comp 1, ranked by decreasing importance in sPLS-DA.

| Var1 | Freq |
| --- | --- |
| Peptoanaerobacter | 1.00 |
| Polaribacter | 0.76 |
| Methanosaeta | 0.82 |
| Porphyromonas | 0.48 |
| Tenacibaculum | 0.40 |

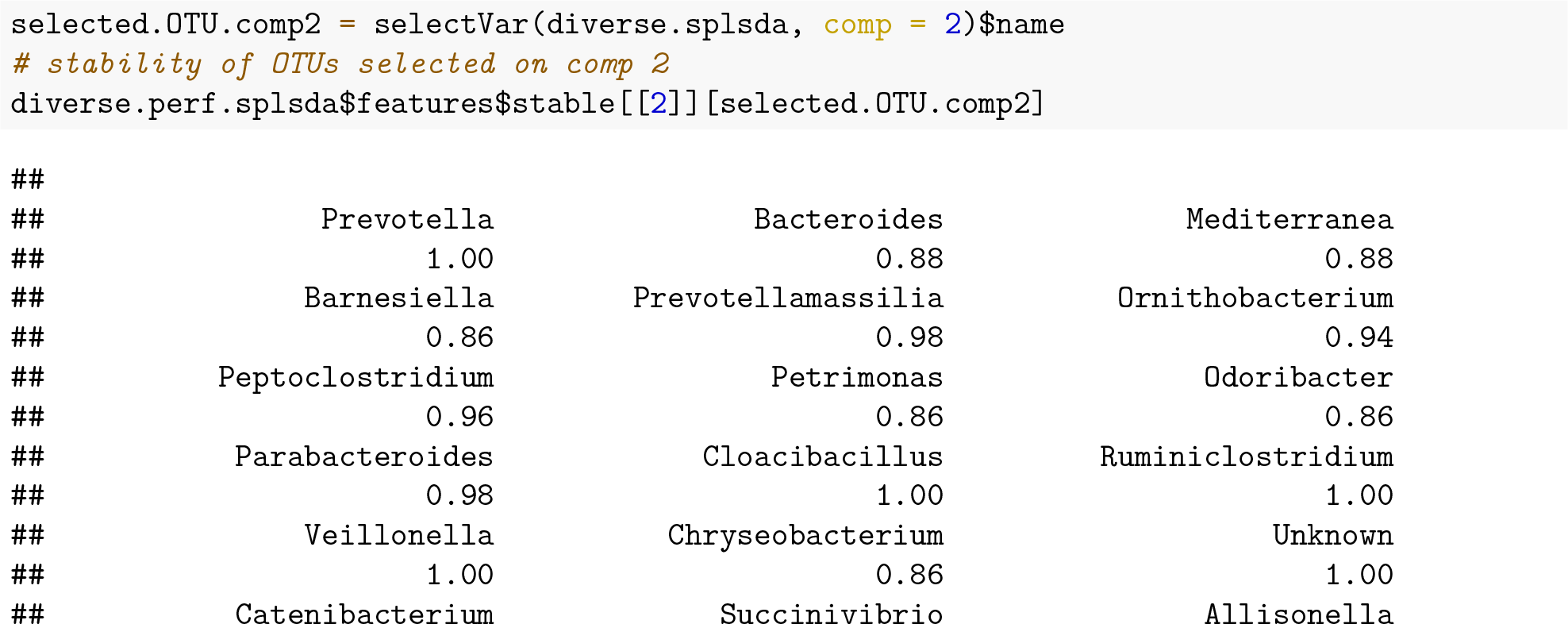
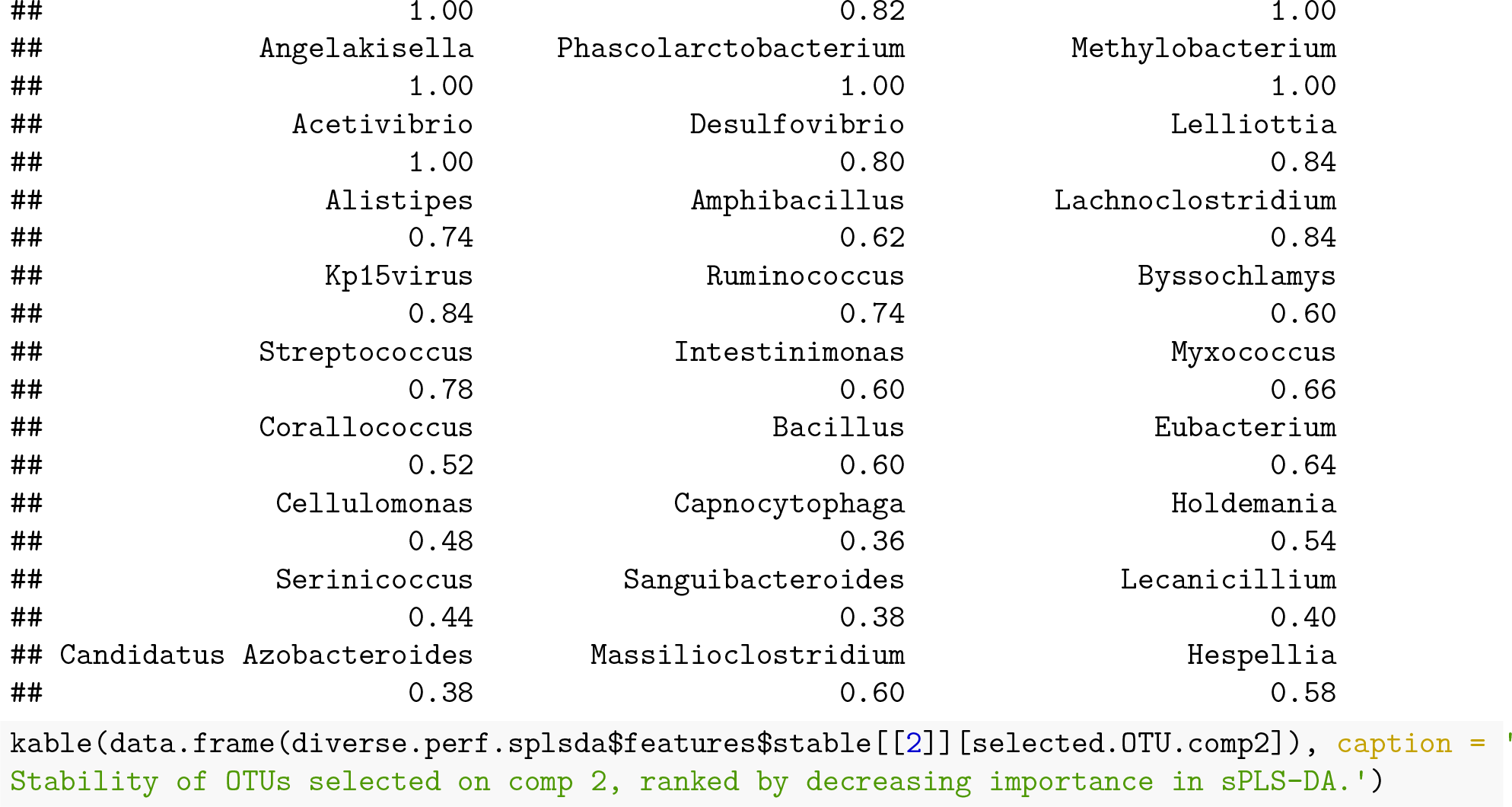

**Table 2:** Stability of OTUs selected on comp 2, ranked by decreasing importance in sPLS-DA.

| Var1 | Freq |
| --- | --- |
| Prevotella | 1.00 |
| Bacteroides | 0.88 |
| Mediterranea | 0.88 |
| Barnesiella | 0.86 |
| Prevotellamassilia | 0.98 |
| Ornithobacterium | 0.94 |
| Peptoclostridium | 0.96 |
| Petrimonas | 0.86 |
| Odoribacter | 0.86 |
| Parabacteroides | 0.98 |
| Cloacibacillus | 1.00 |
| Ruminiclostridium | 1.00 |
| Veillonella | 1.00 |
| Chryseobacterium | 0.86 |
| Unknown | 1.00 |
| Catenibacterium | 1.00 |
| Succinivibrio | 0.82 |
| Allisonella | 1.00 |
| Angelakisella | 1.00 |
| Phascolarctobacterium | 1.00 |
| Methylobacterium | 1.00 |
| Acetivibrio | 1.00 |
| Desulfovibrio | 0.80 |
| Lelliottia | 0.84 |
| Alistipes | 0.74 |
| Amphibacillus | 0.62 |
| Lachnoclostridium | 0.84 |
| Kp15virus | 0.84 |
| Ruminococcus | 0.74 |
| Byssochlamys | 0.60 |
| Streptococcus | 0.78 |
| Intestinimonas | 0.60 |
| Myxococcus | 0.66 |
| Corallococcus | 0.52 |
| Bacillus | 0.60 |
| Eubacterium | 0.64 |
| Cellulomonas | 0.48 |
| Capnocytophaga | 0.36 |
| Holdemania | 0.54 |
| Serinicoccus | 0.44 |
| Sanguibacteroides | 0.38 |
| Lecanicillium | 0.40 |
| Candidatus Azobacteroides | 0.38 |
| Massilioclostridium | 0.60 |
| Hespellia | 0.58 |

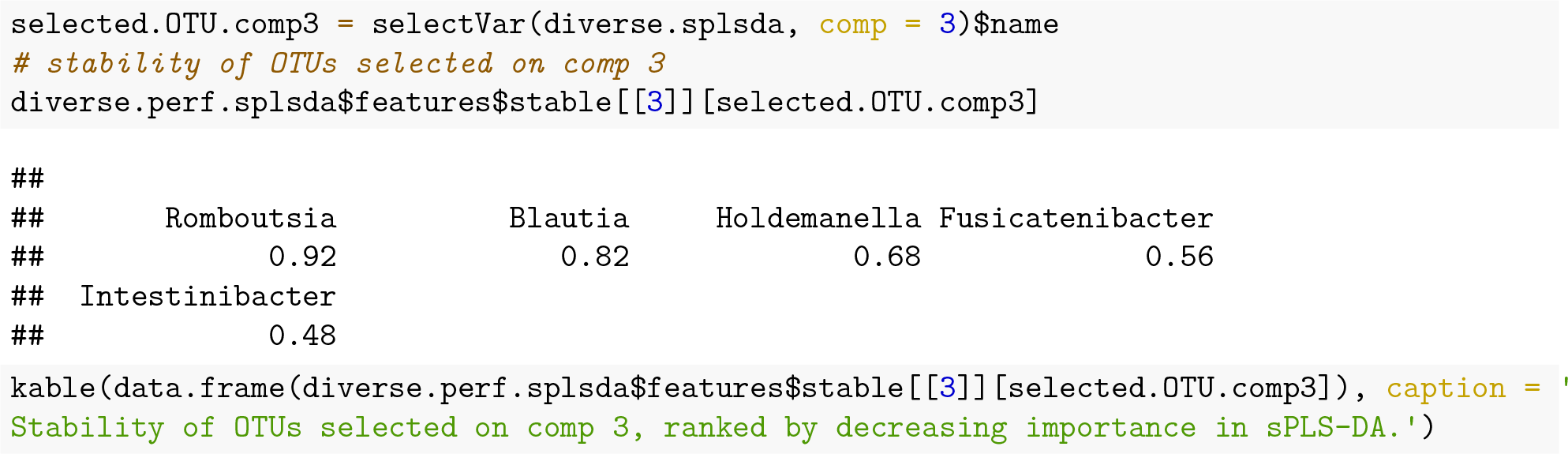

**Table 3:**
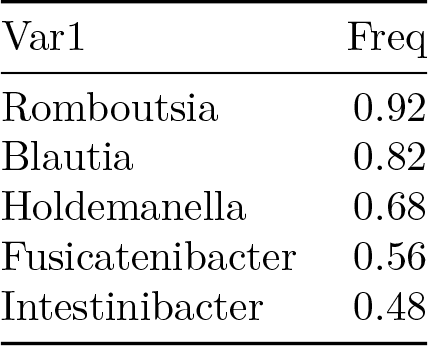
Stability of OTUs selected on comp 3, ranked by decreasing importance in sPLS-DA.

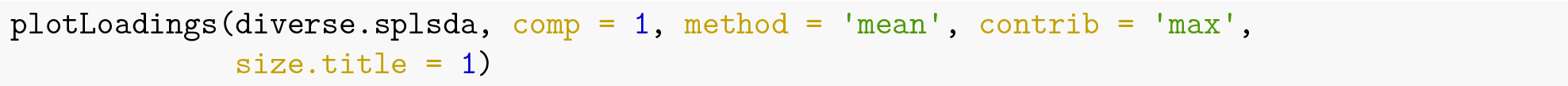
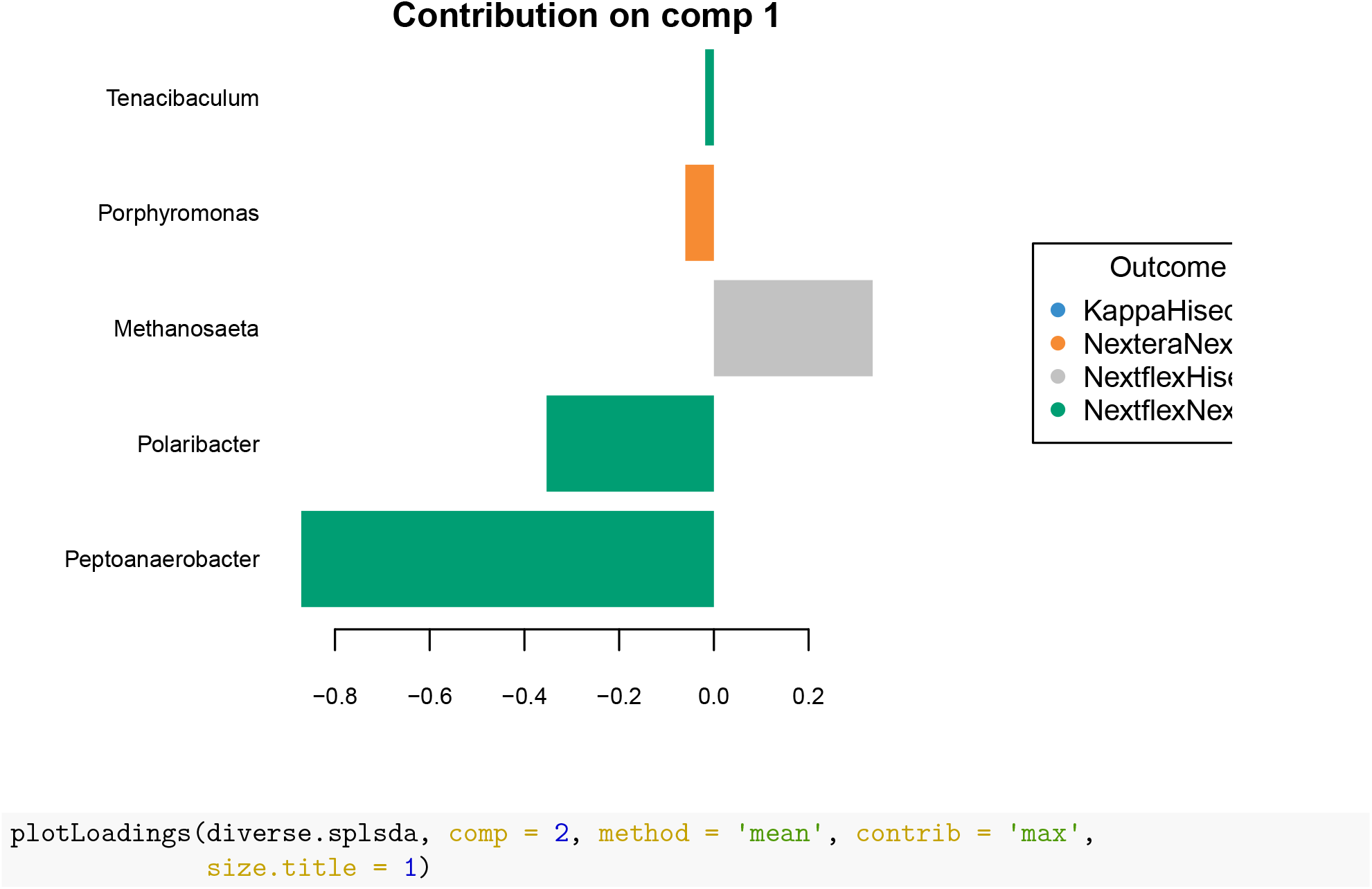
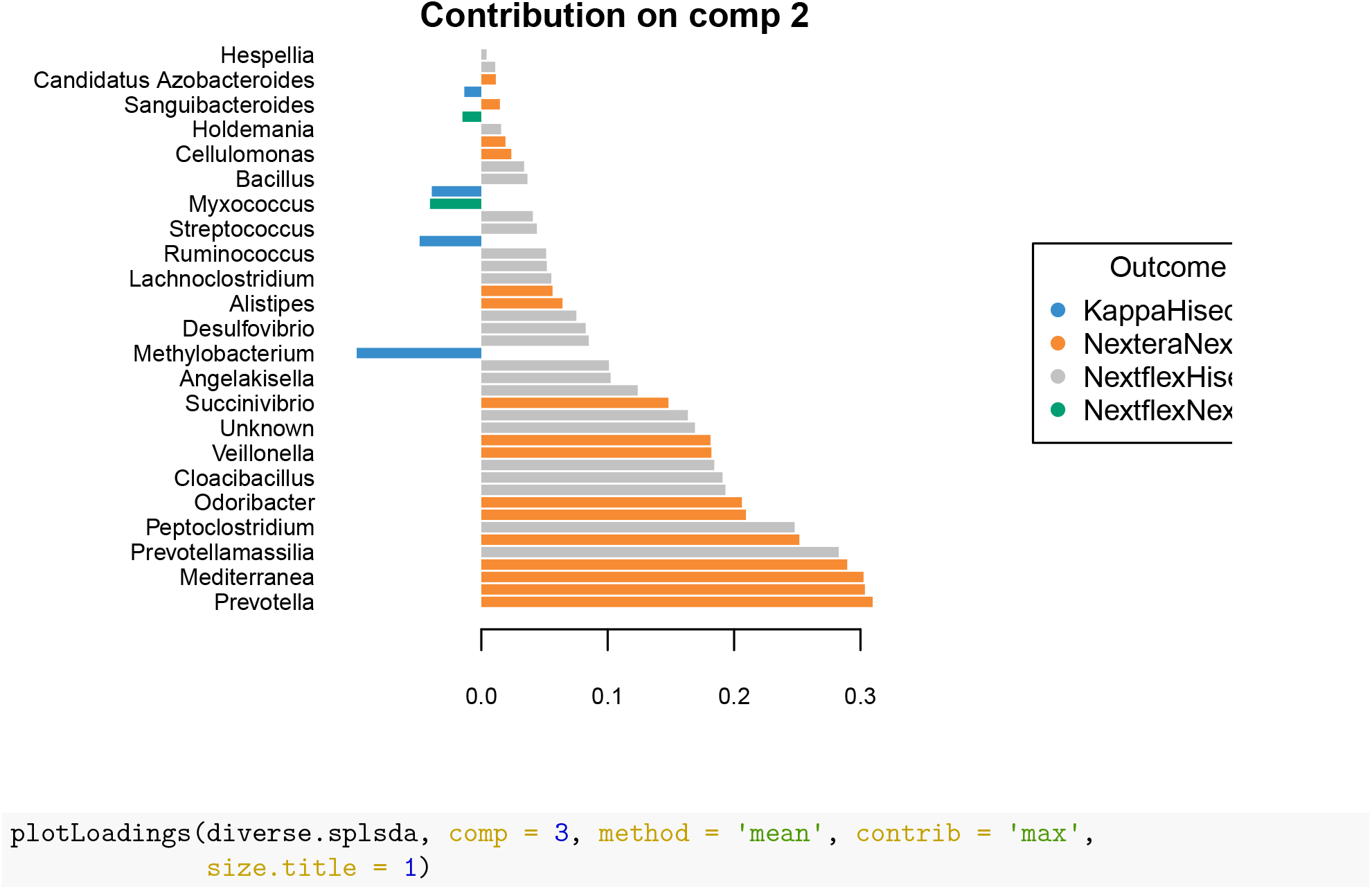
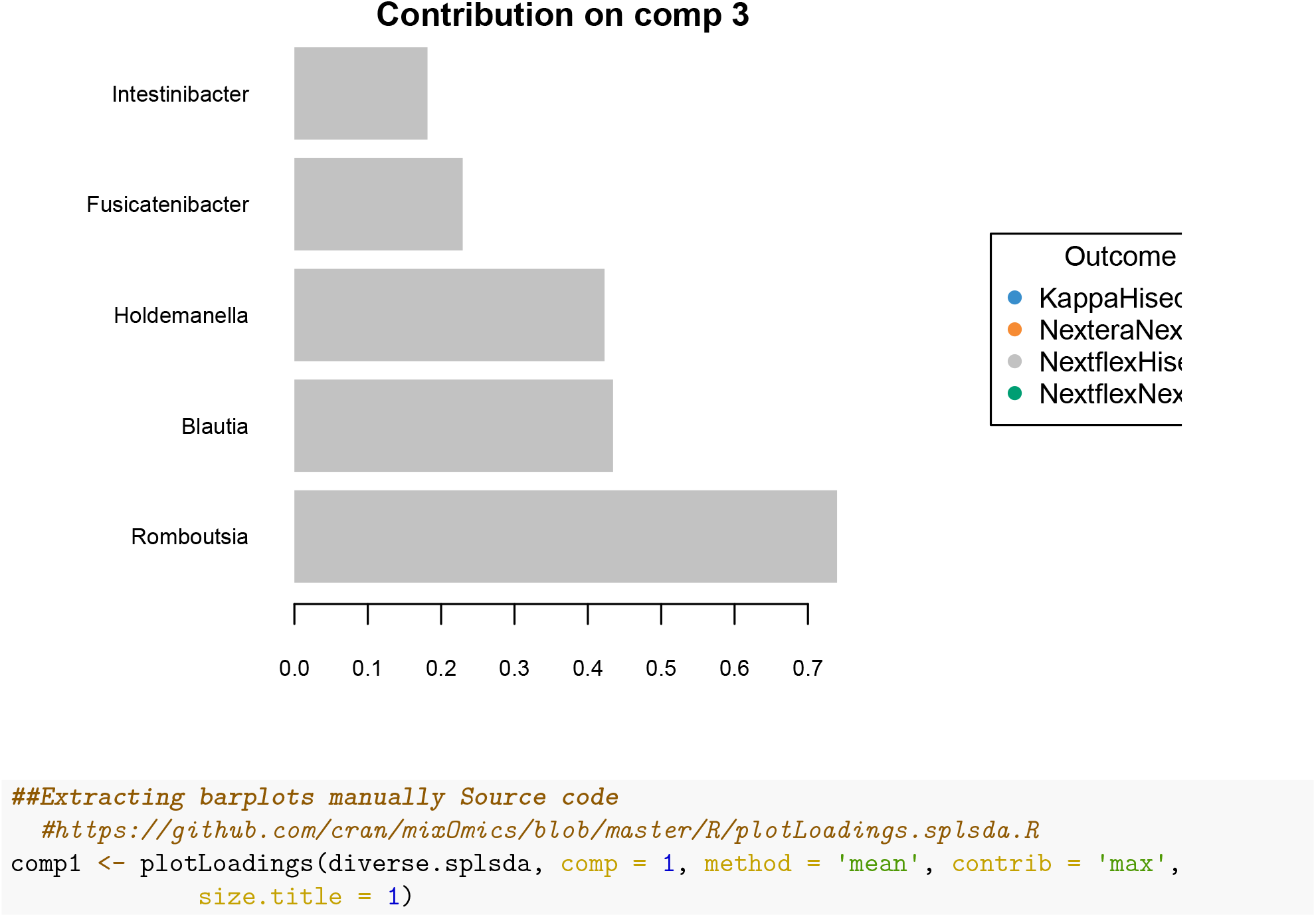
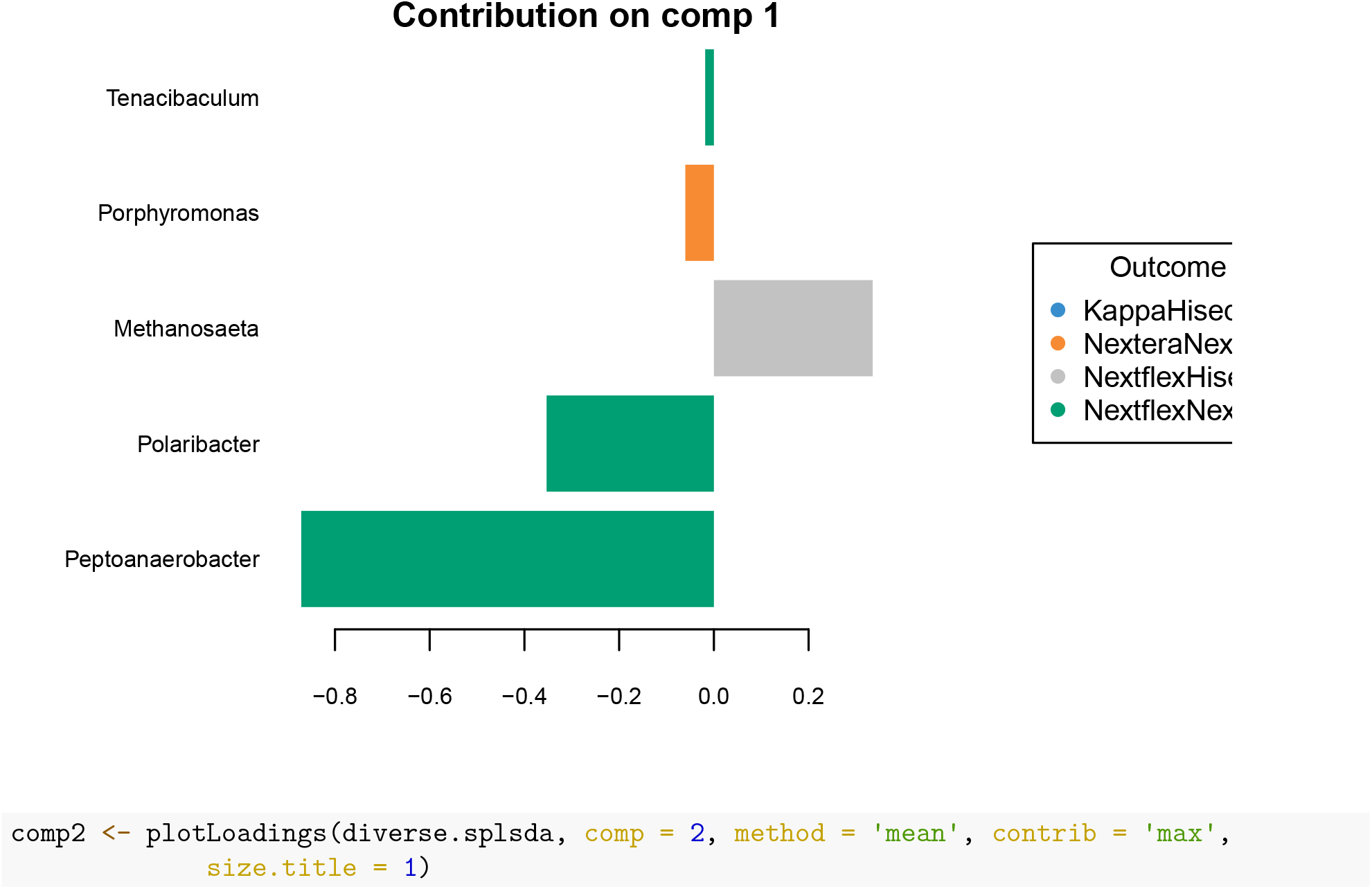
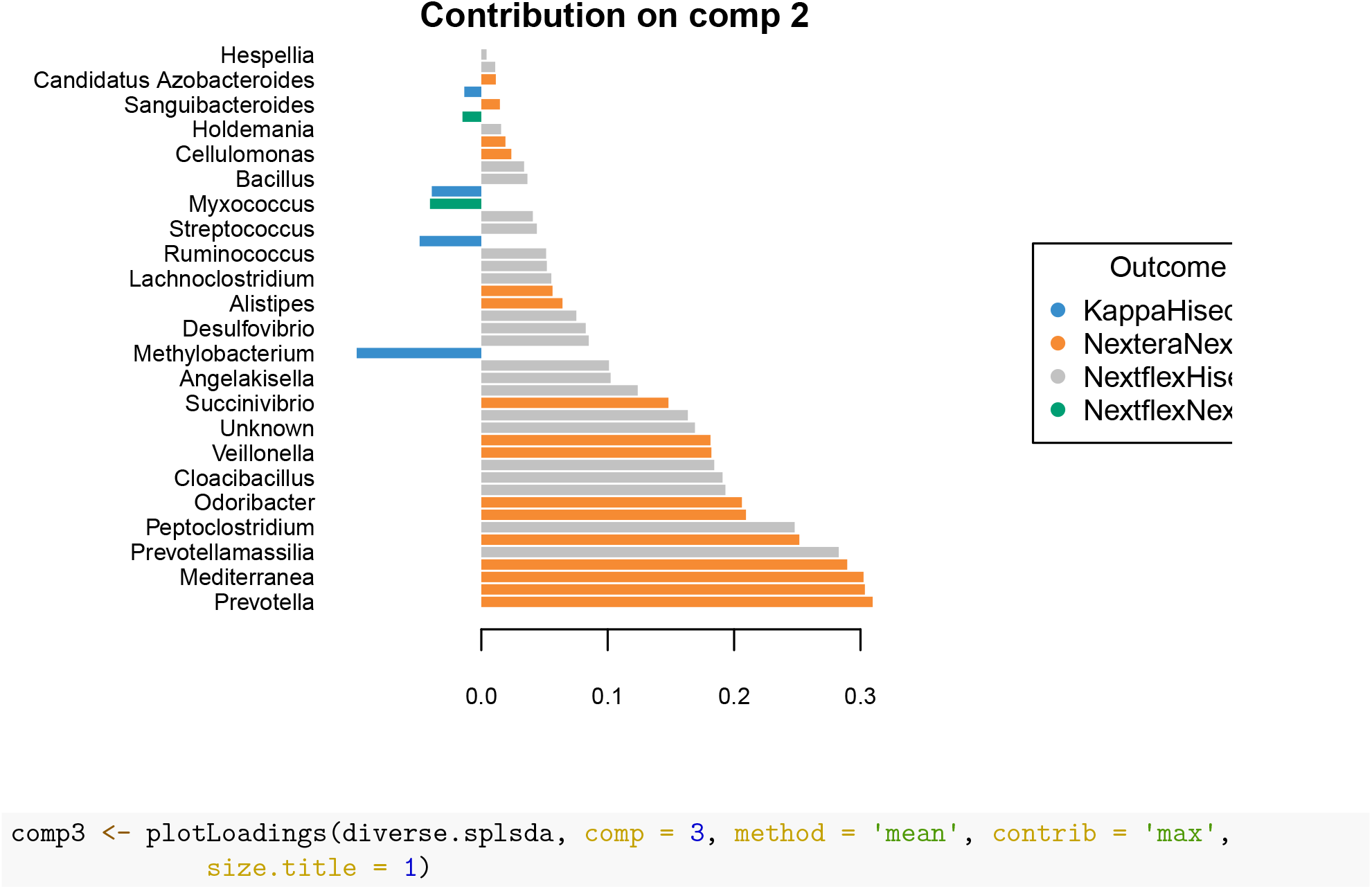
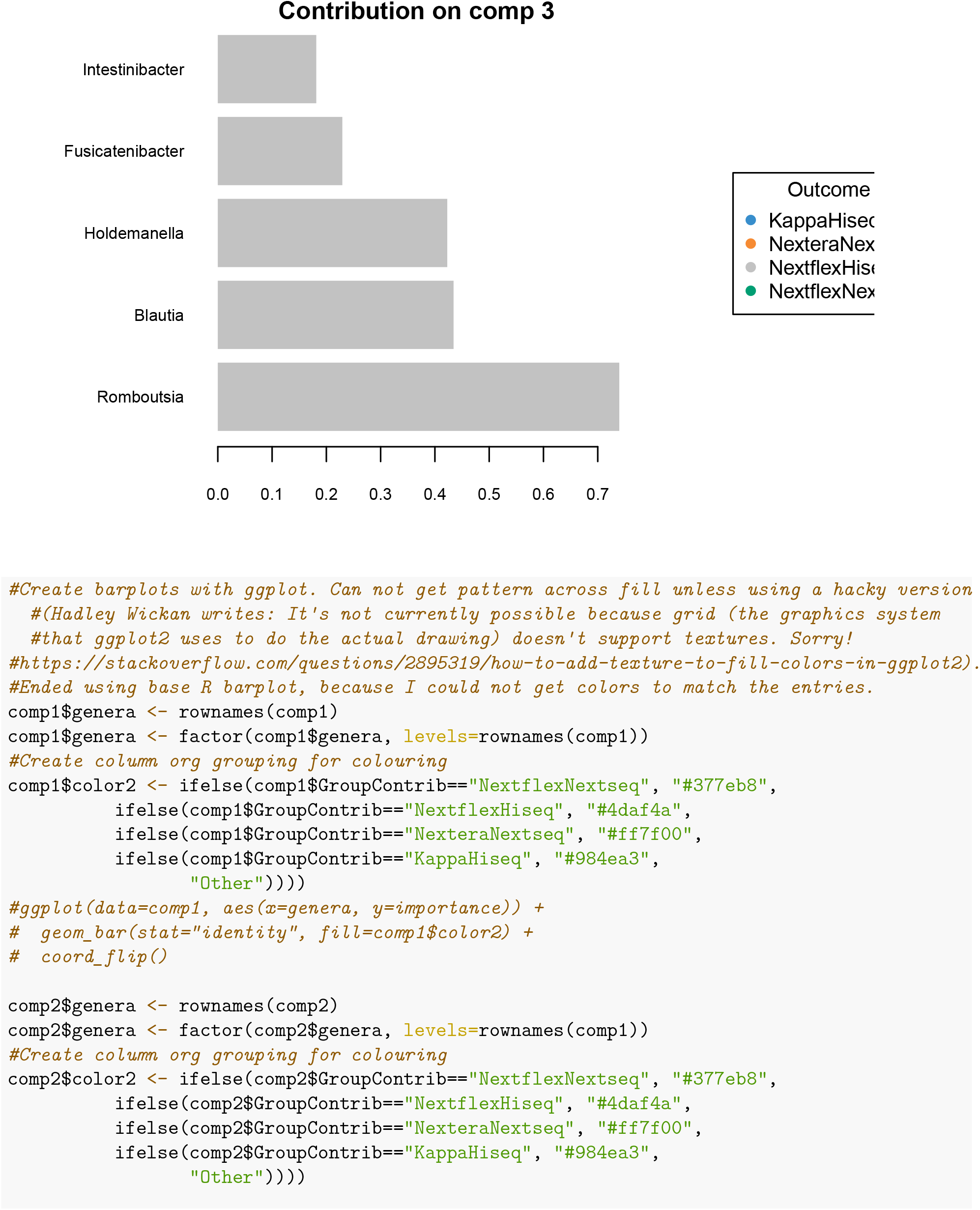
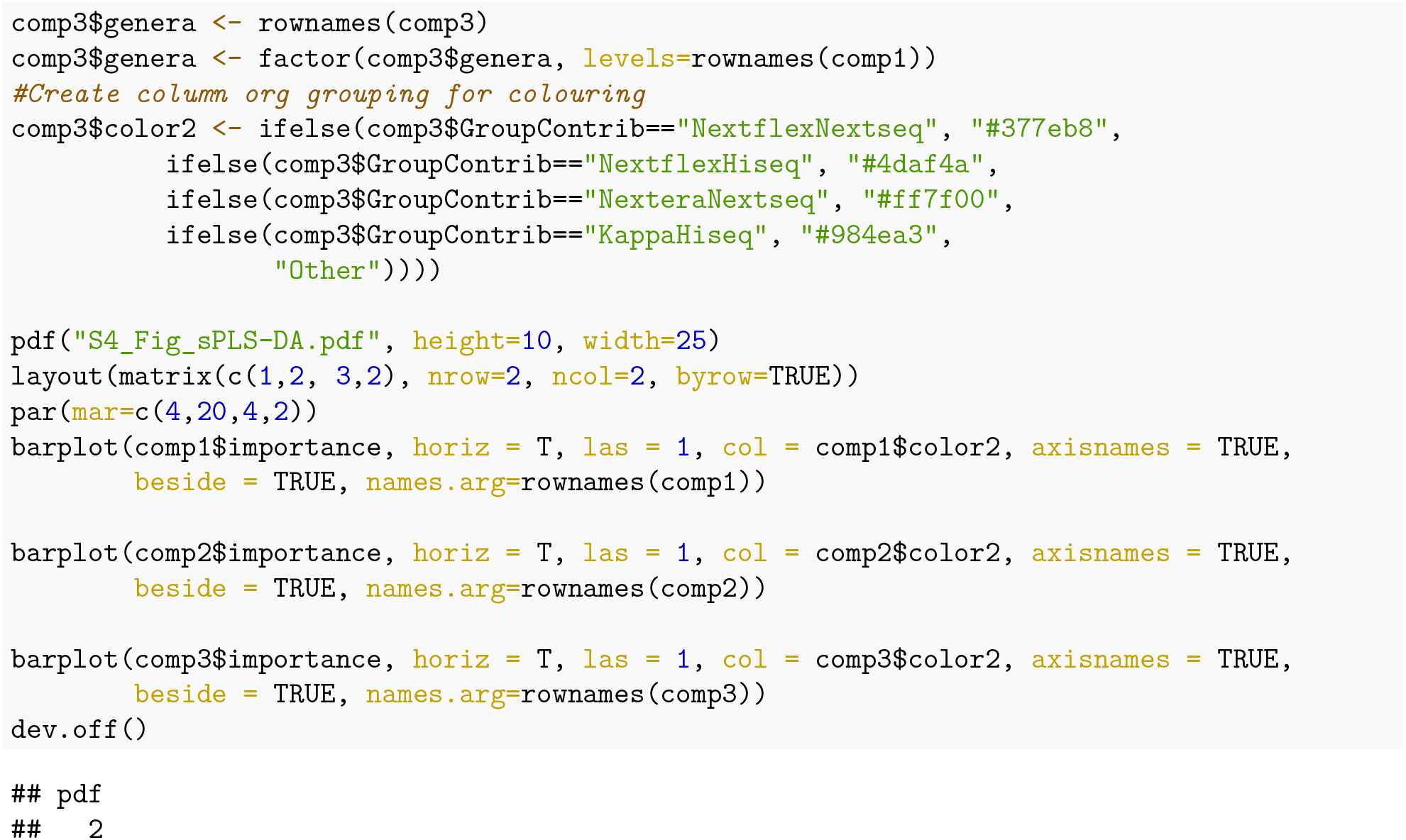

#### Multivariate ANOVA based on dissimilarities (Adonis)

Statistical output is presented in Table 1 and S2_Table

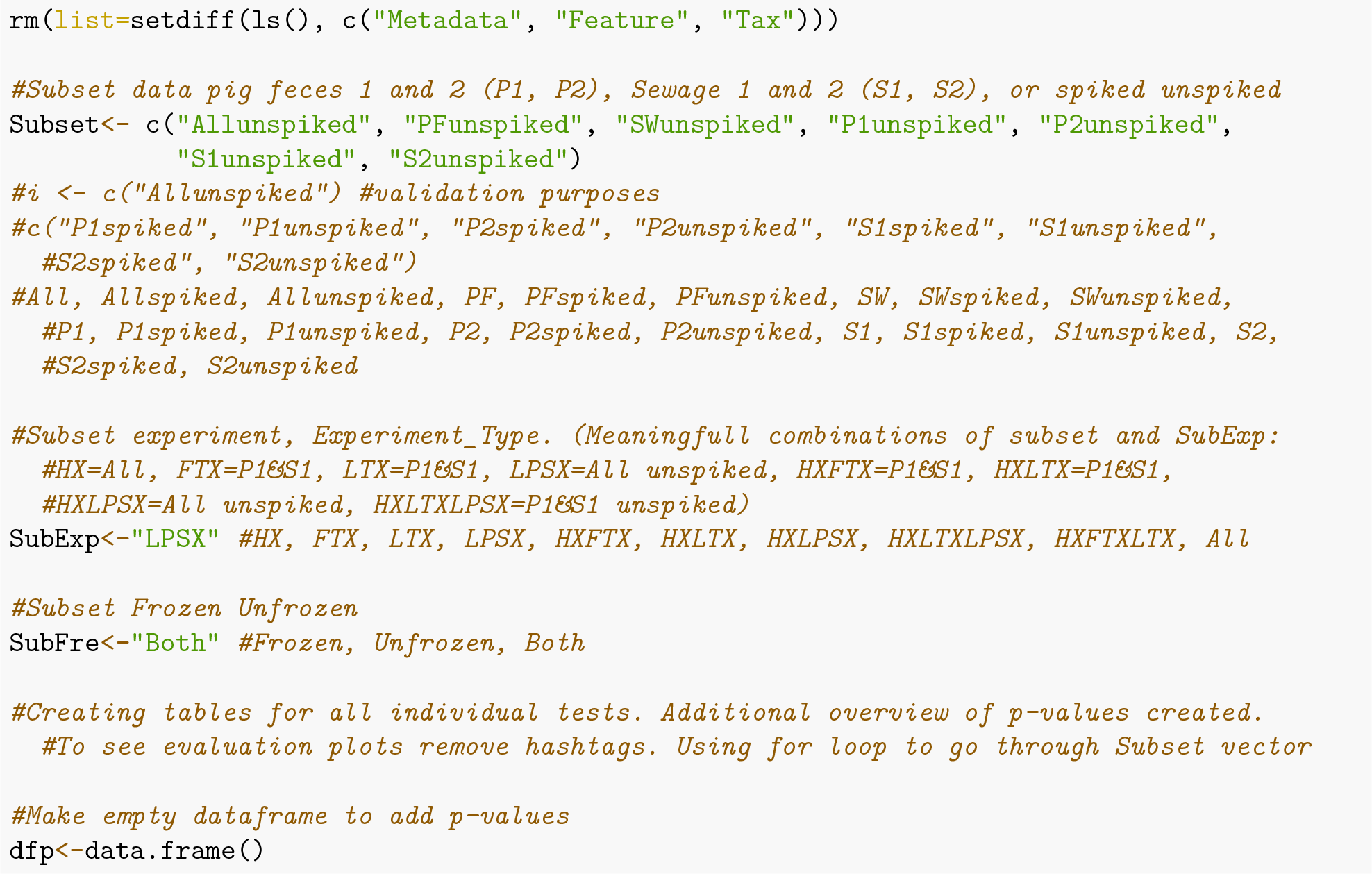
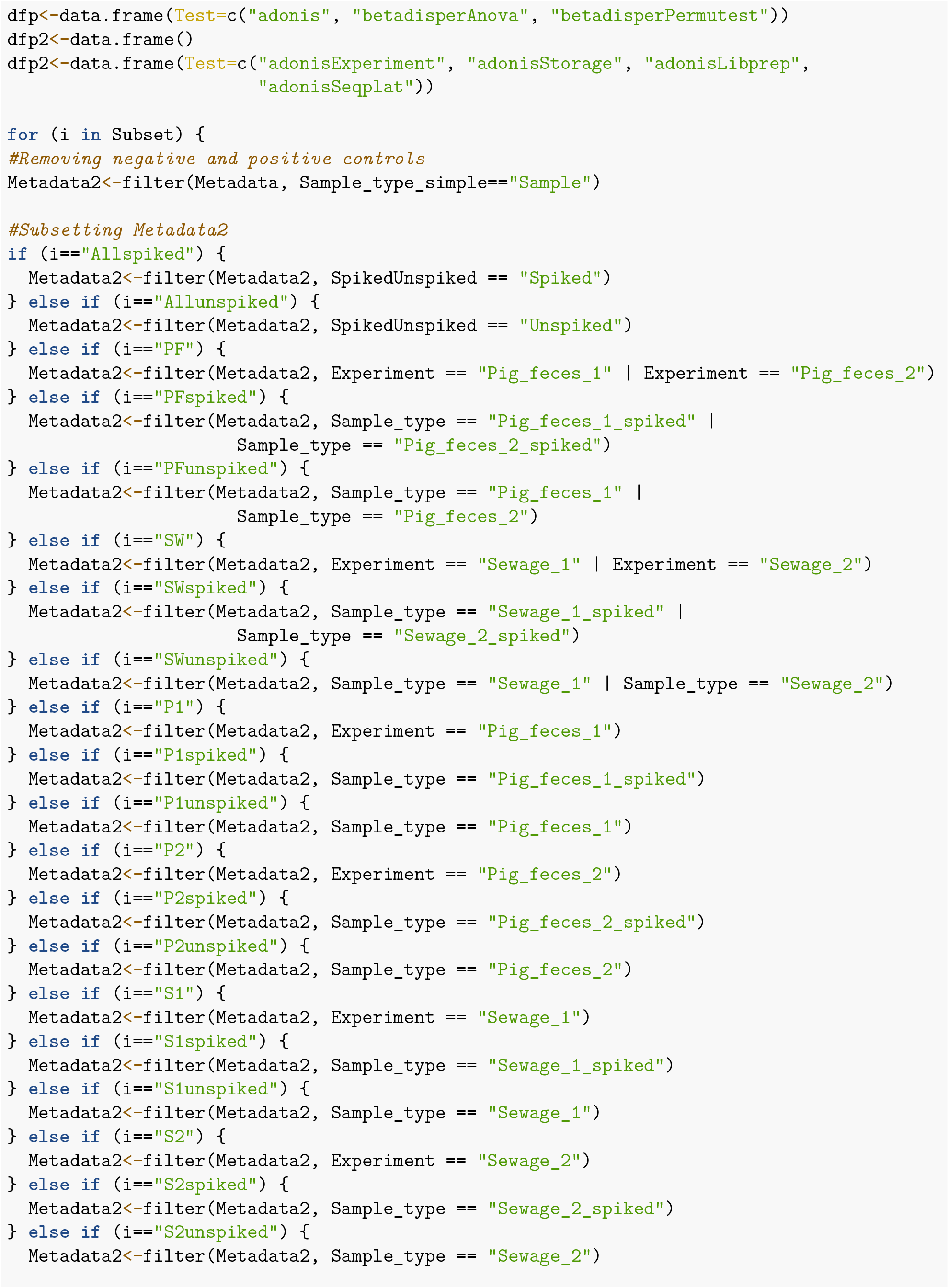
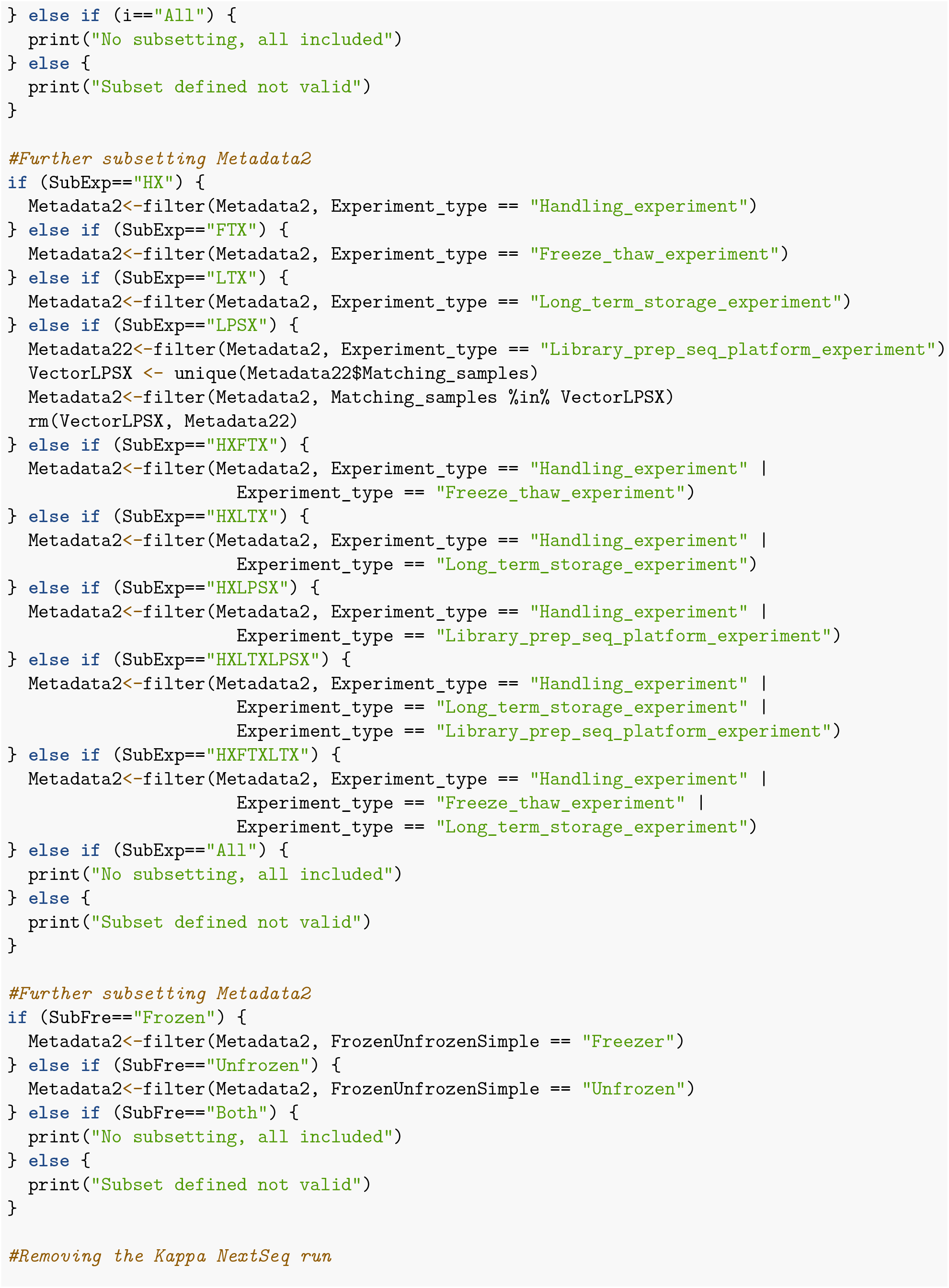
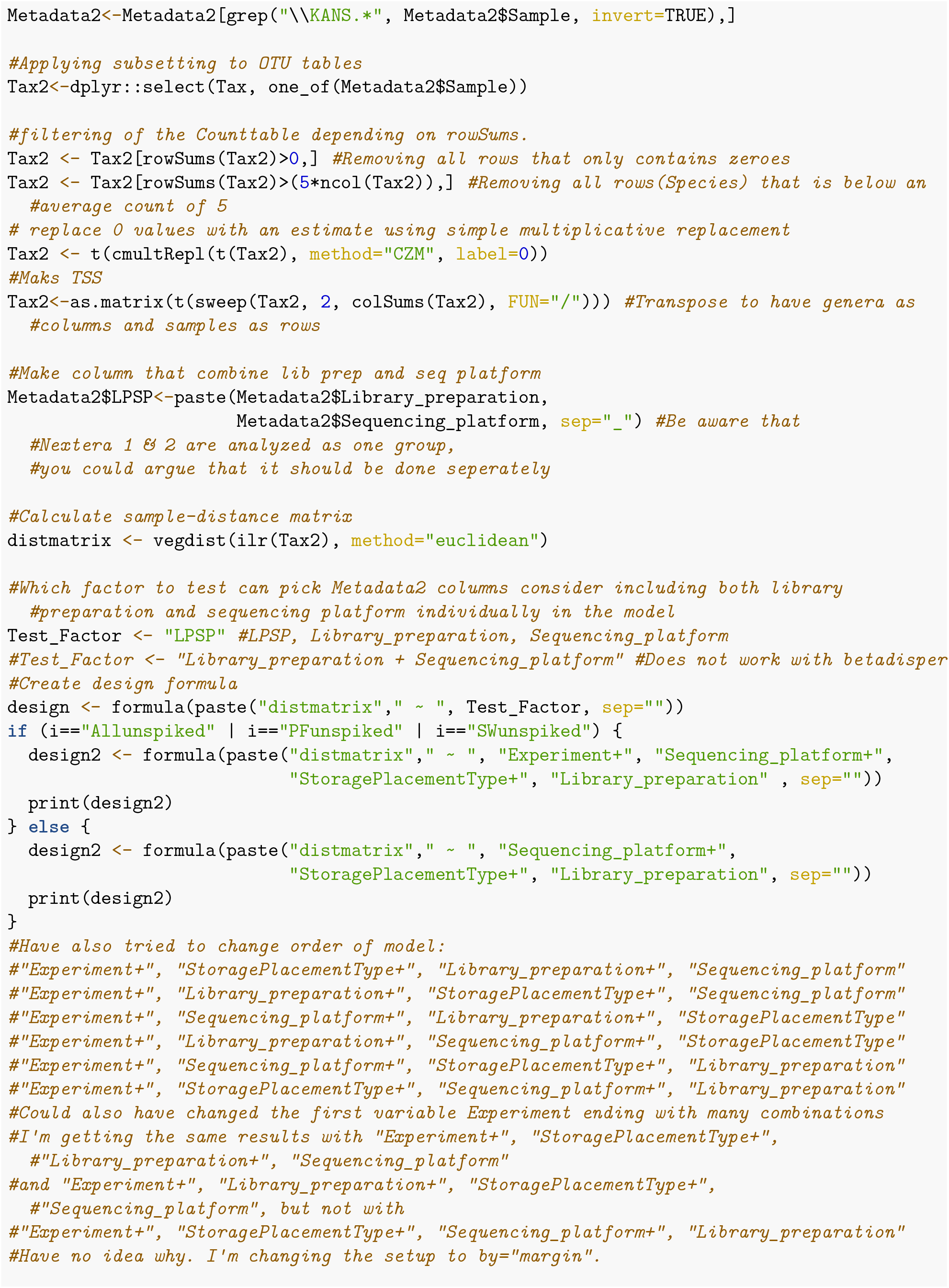
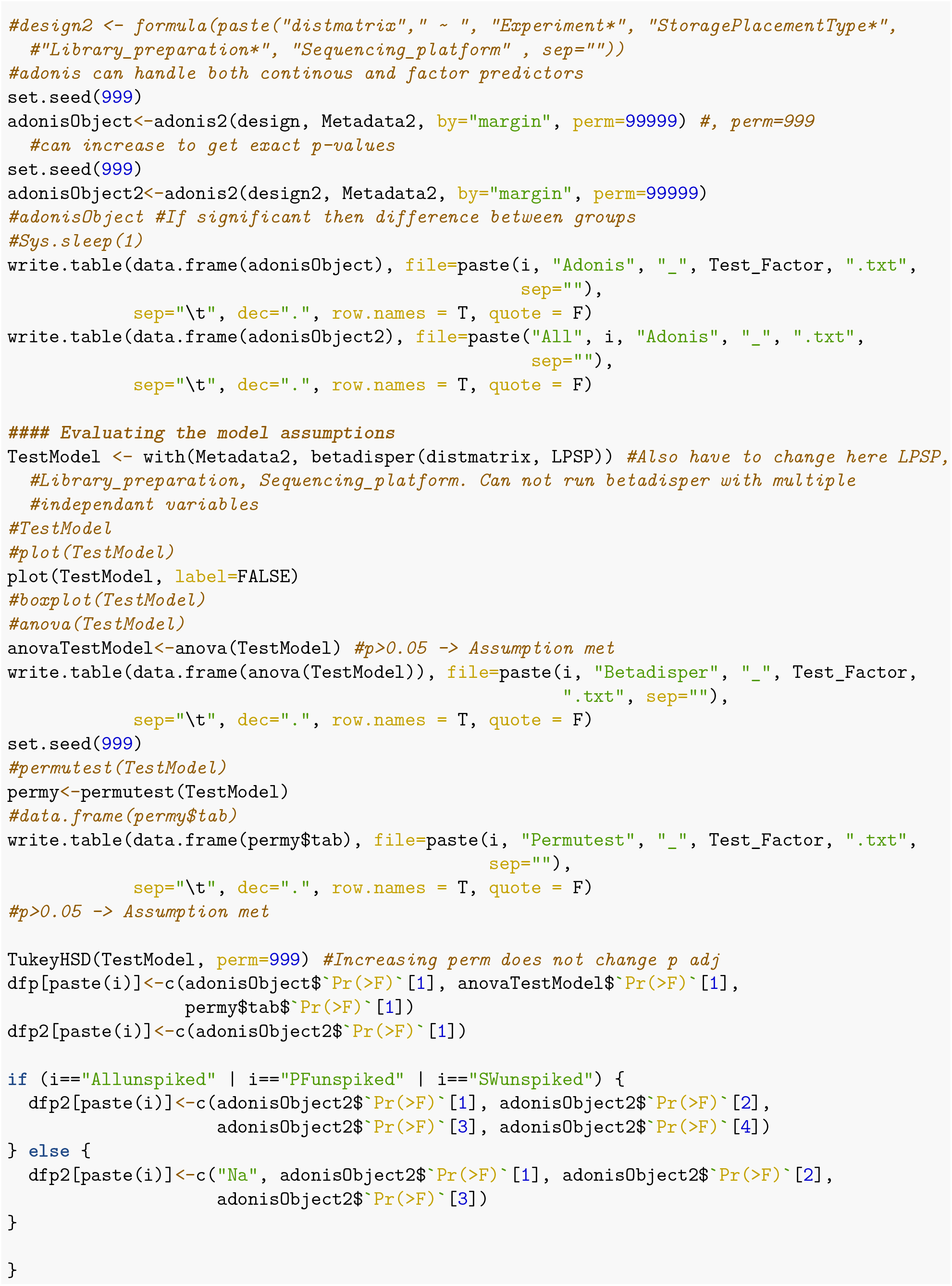
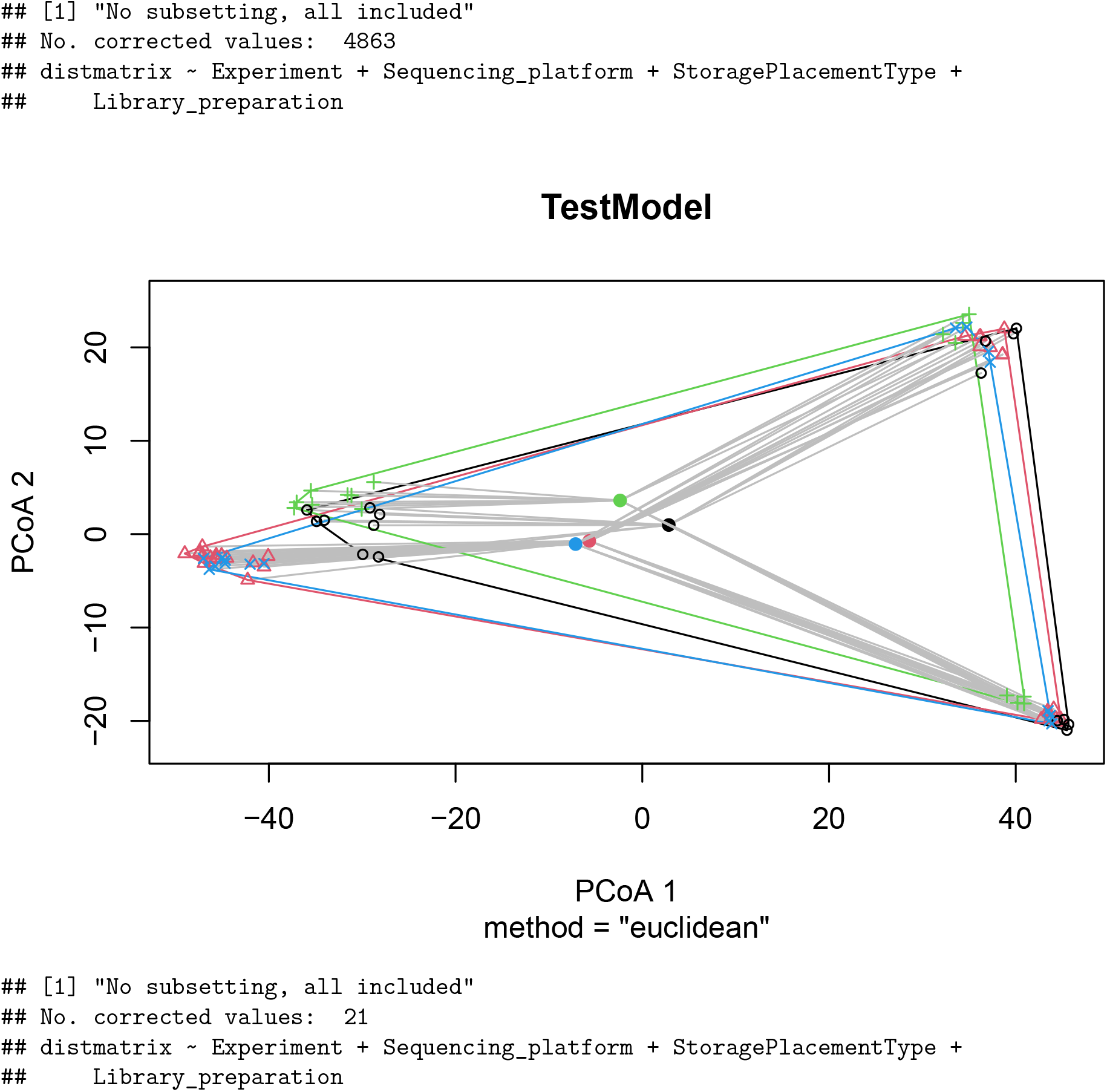
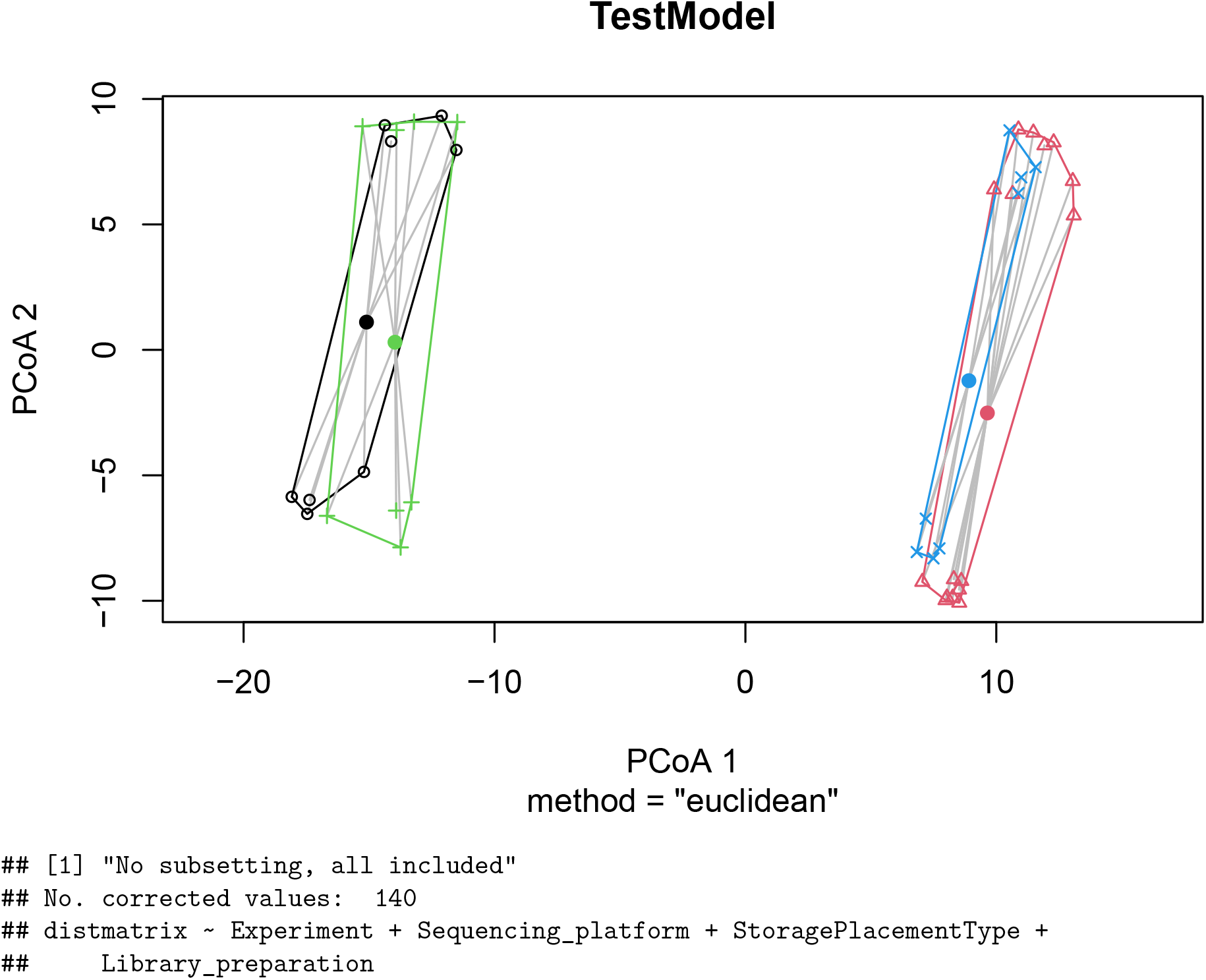
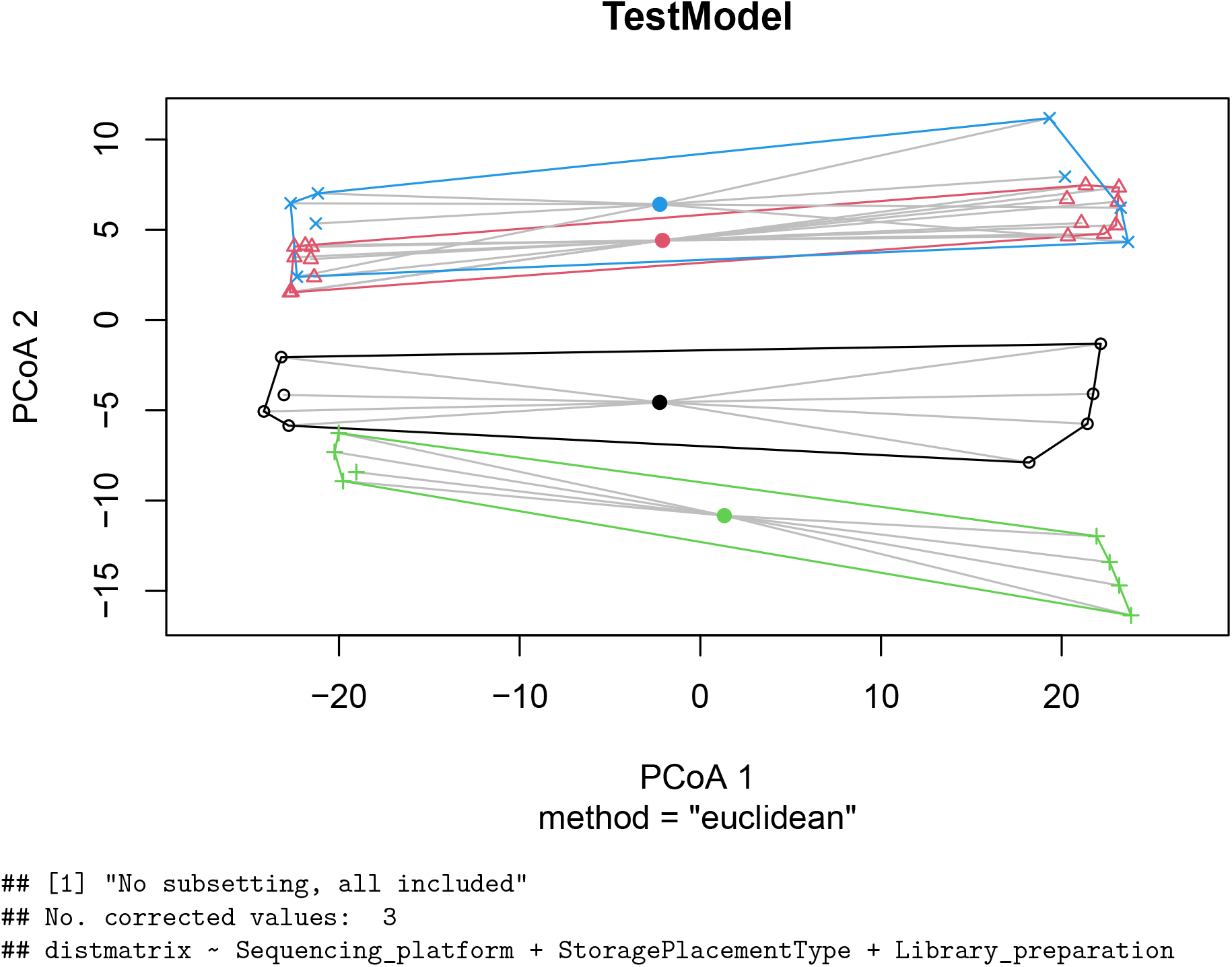
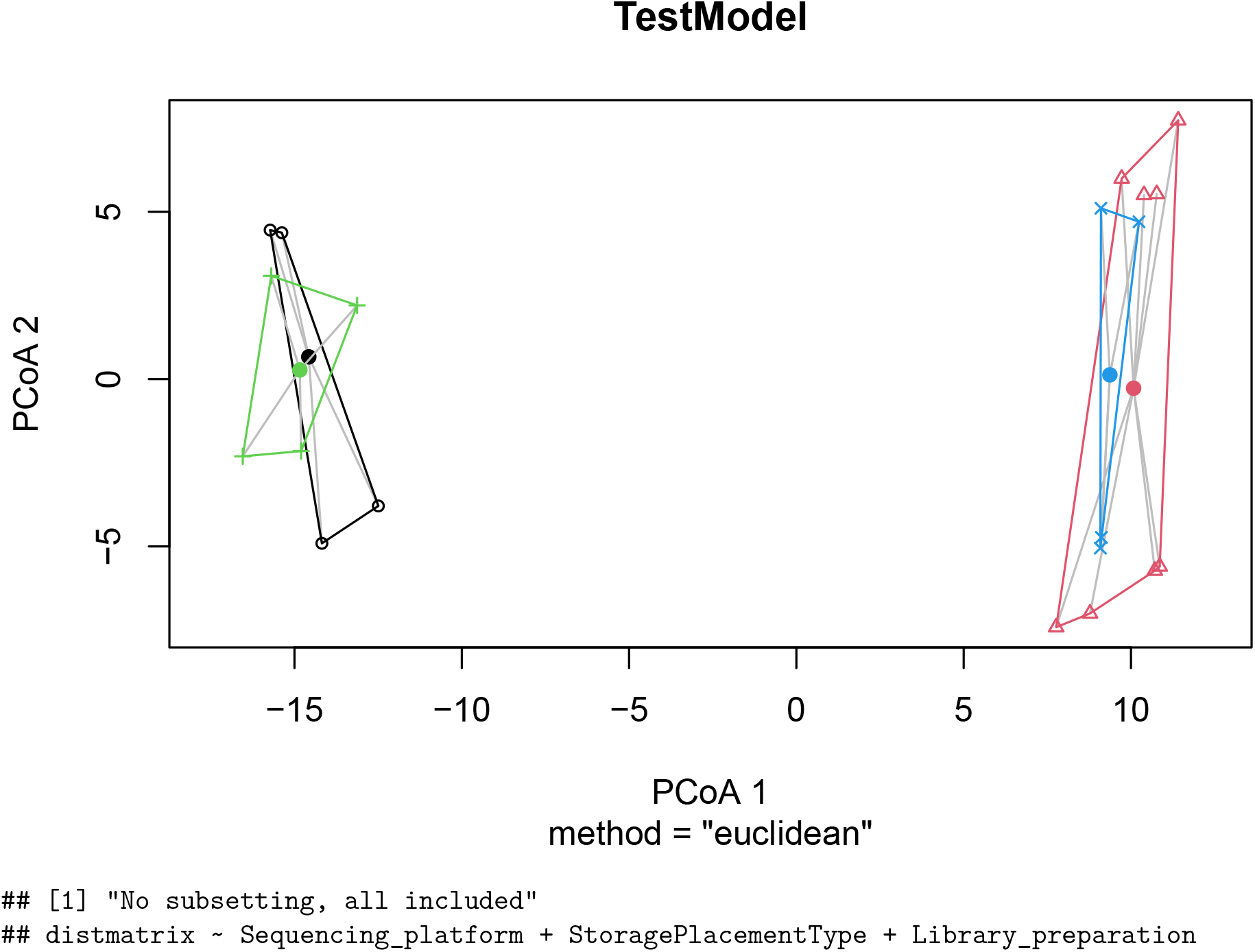
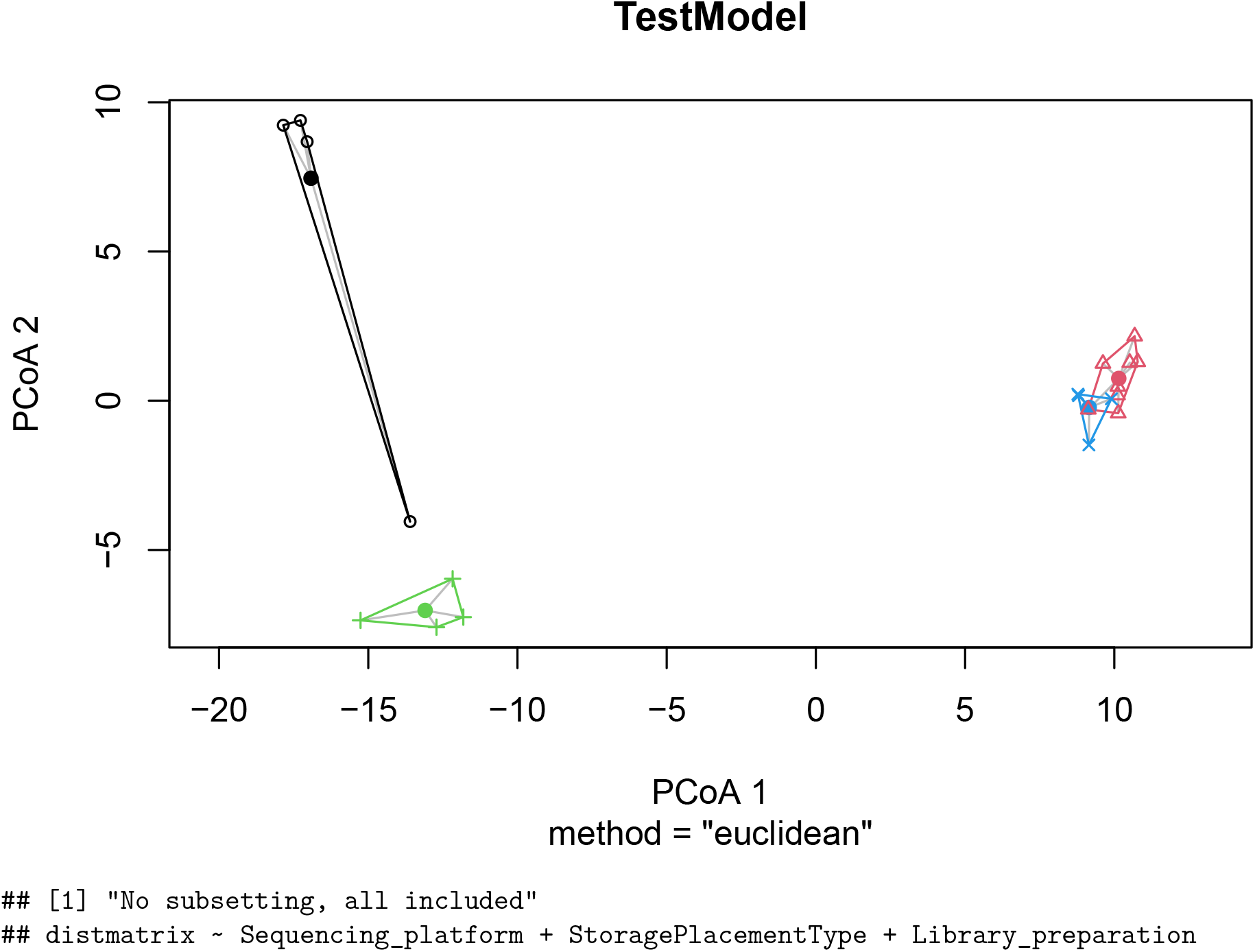
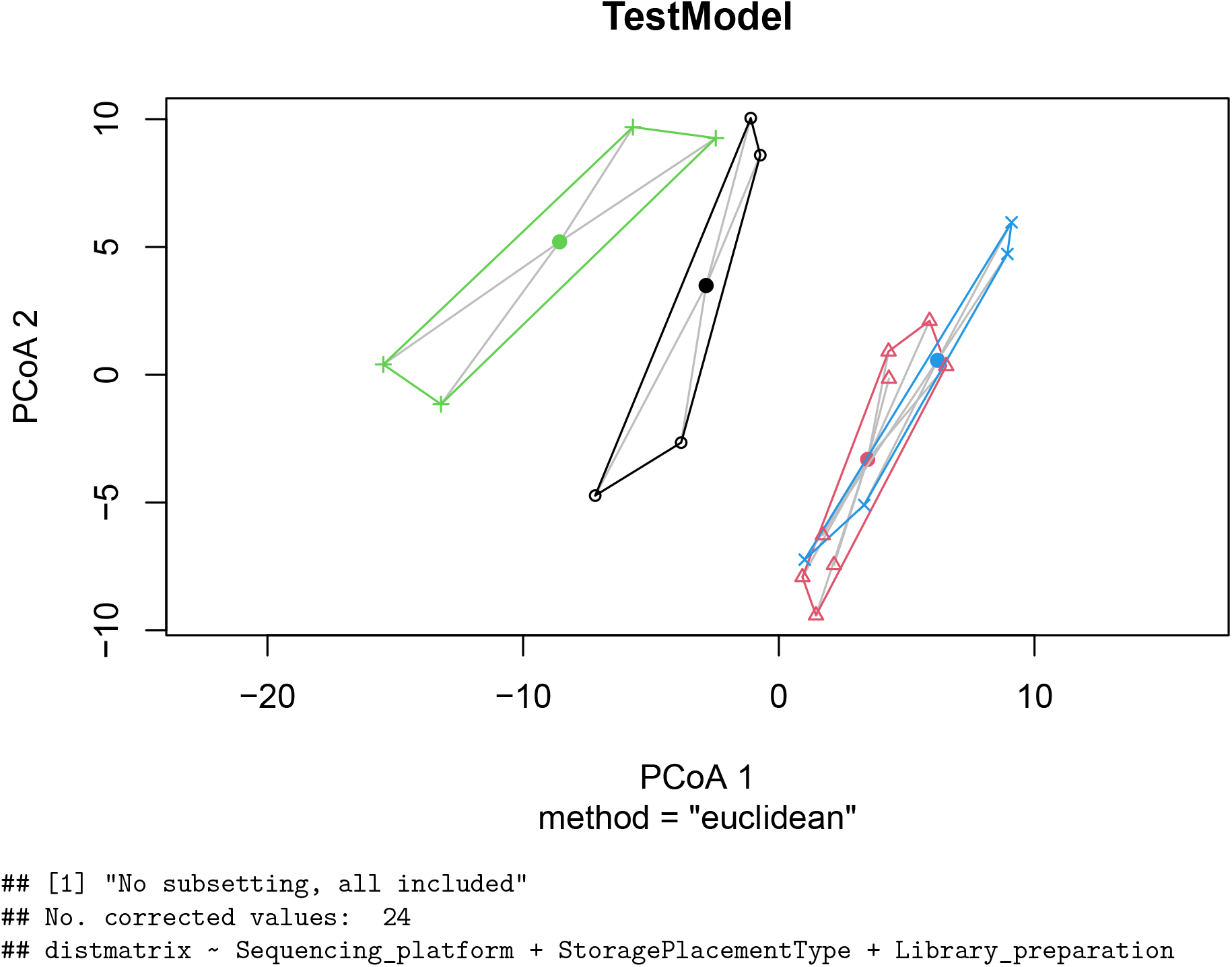
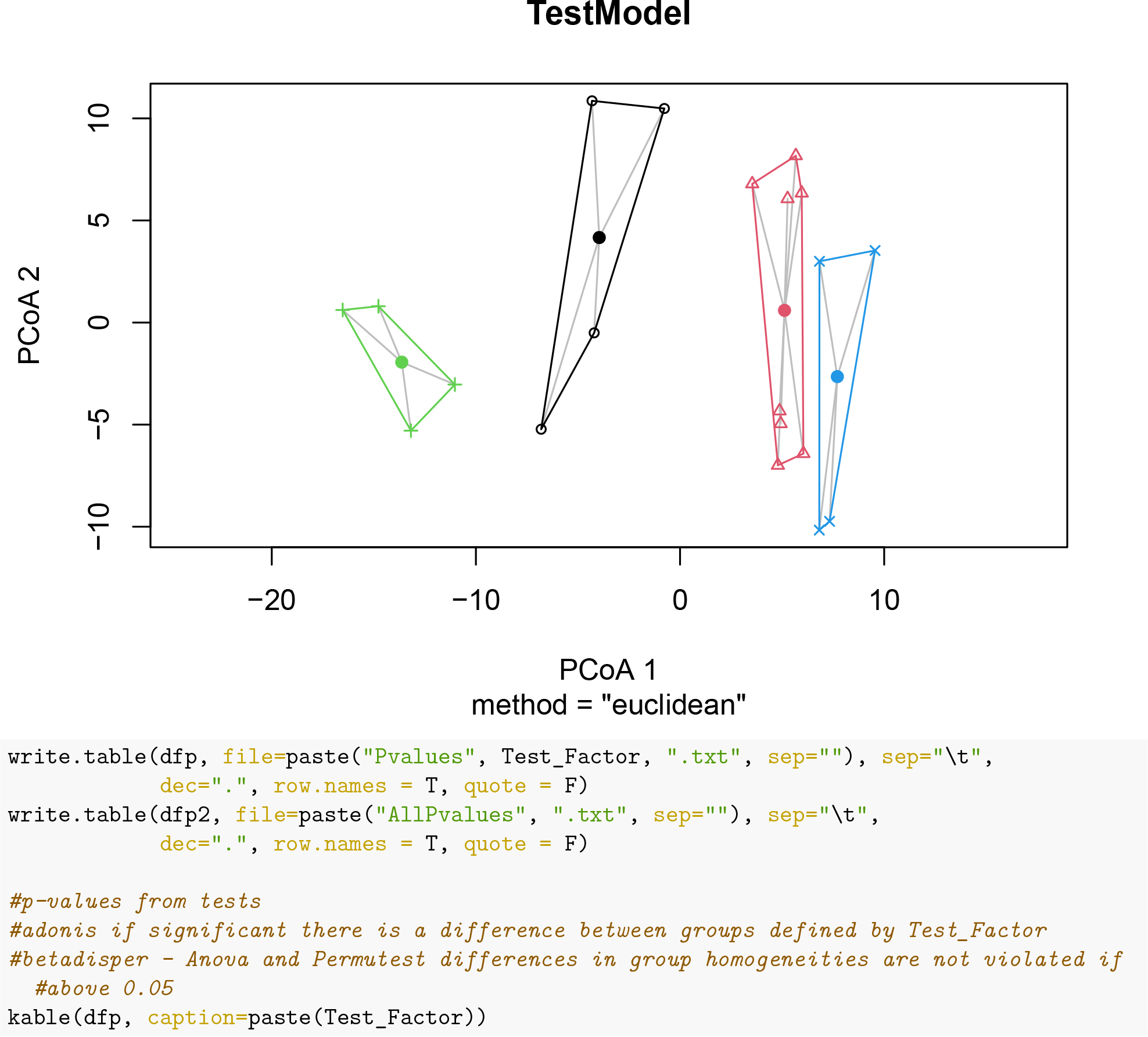

**Table 4:** LPSP

| Test | Allunspiked | PFunspiked | SWunspiked | P1unspiked | P2unspiked | S1unspiked | S2unspiked |
| --- | --- | --- | --- | --- | --- | --- | --- |
| adonis | 0.299100 | 0.000010 | 0.2353700 | 0.0000100 | 0.0000100 | 0.0000100 | 0.0000100 |
| betadisperAnova | 0.000817 | 0.081415 | 0.6165621 | 0.0061238 | 0.2675113 | 0.1009037 | 0.3100214 |
| betadisperPermutest | 0.002000 | 0.089000 | 0.6150000 | 0.0110000 | 0.2400000 | 0.0930000 | 0.2900000 |

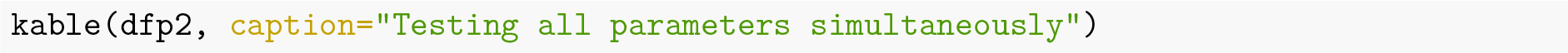

**Table 5:** Testing all parameters simultaneously

| Test | Allunspiked | PFunspiked | SWunspiked | P1unspiked | P2unspiked | S1unspiked | S2unspiked |
| --- | --- | --- | --- | --- | --- | --- | --- |
| adonisExperiment | 0.00001 | 0.00001 | 0.00001 | Na | Na | Na | Na |
| adonisStorage | 0.00030 | 0.00001 | 0.00444 | 1e-05 | 1e-05 | 1e-05 | 1e-05 |
| adonisLibprep | 0.06384 | 0.00381 | 0.02524 | 0.00284 | 0.16807 | 1e-05 | 1e-05 |
| adonisSeqplat | 0.04171 | 0.00057 | 0.02953 | 0.02804 | 0.0054 | 0.00036 | 6e-05 |

### Constrained ordination with redundancy analysis (rda)

The vegan function capscale were used to create the rda

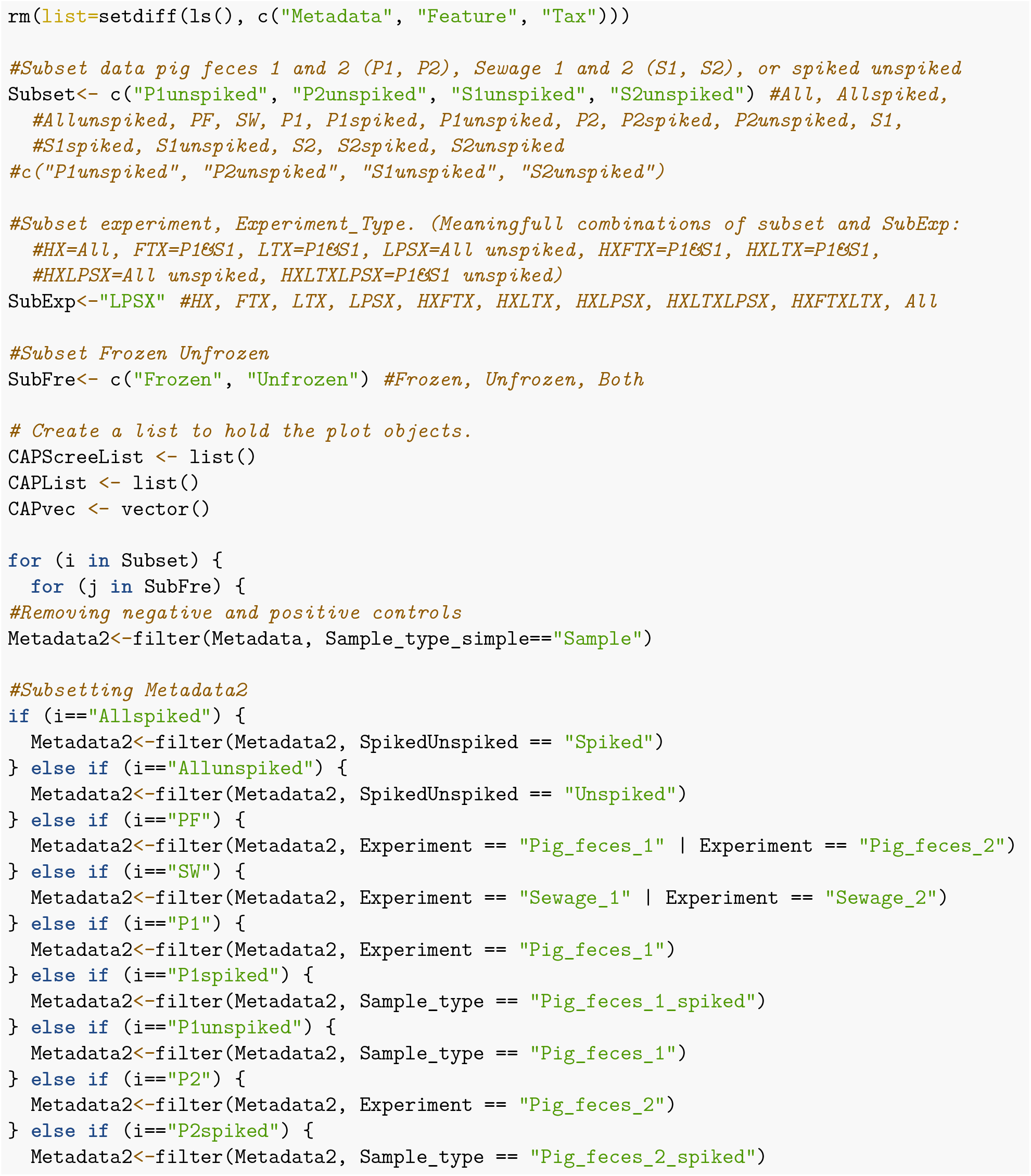
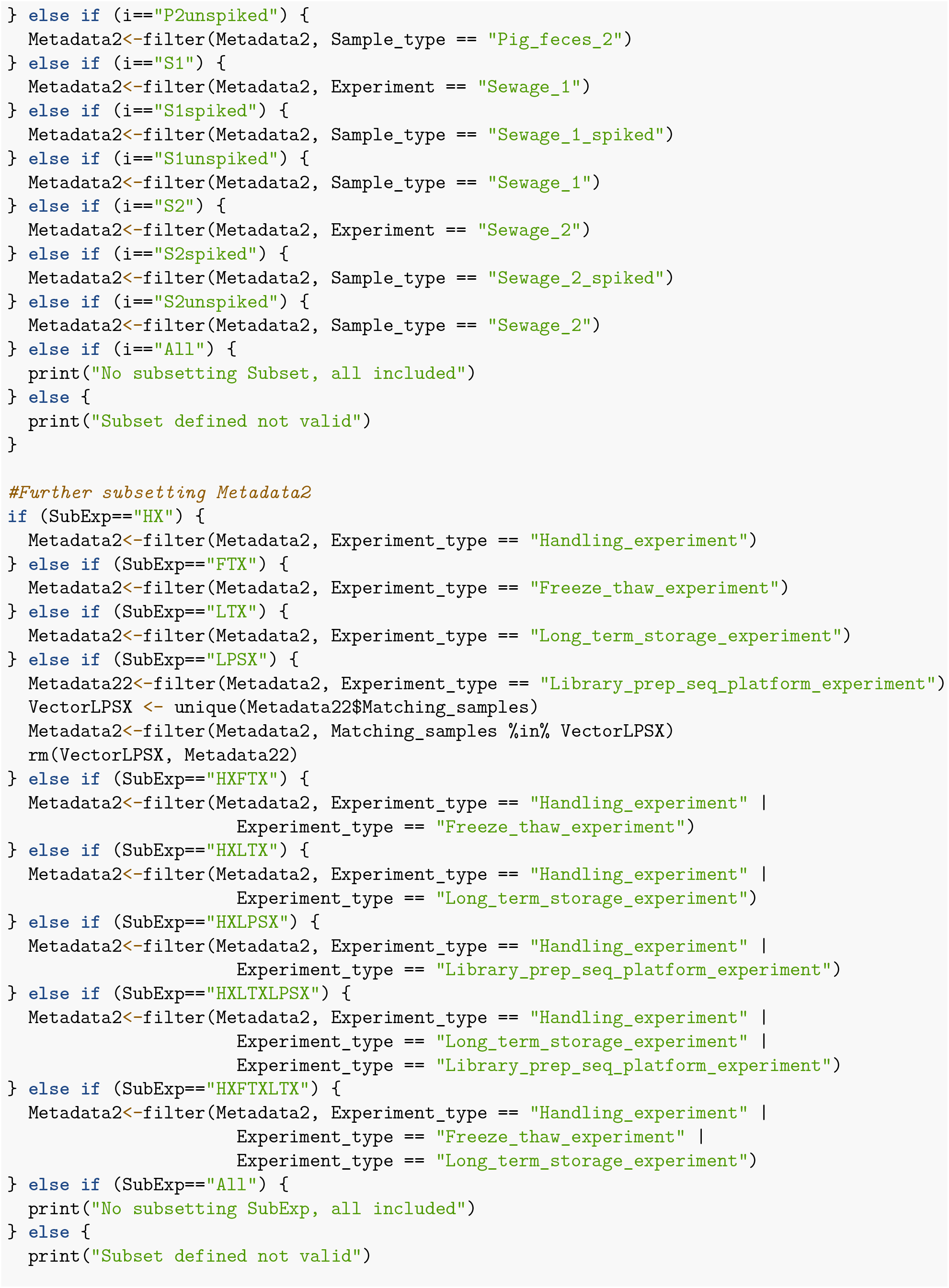
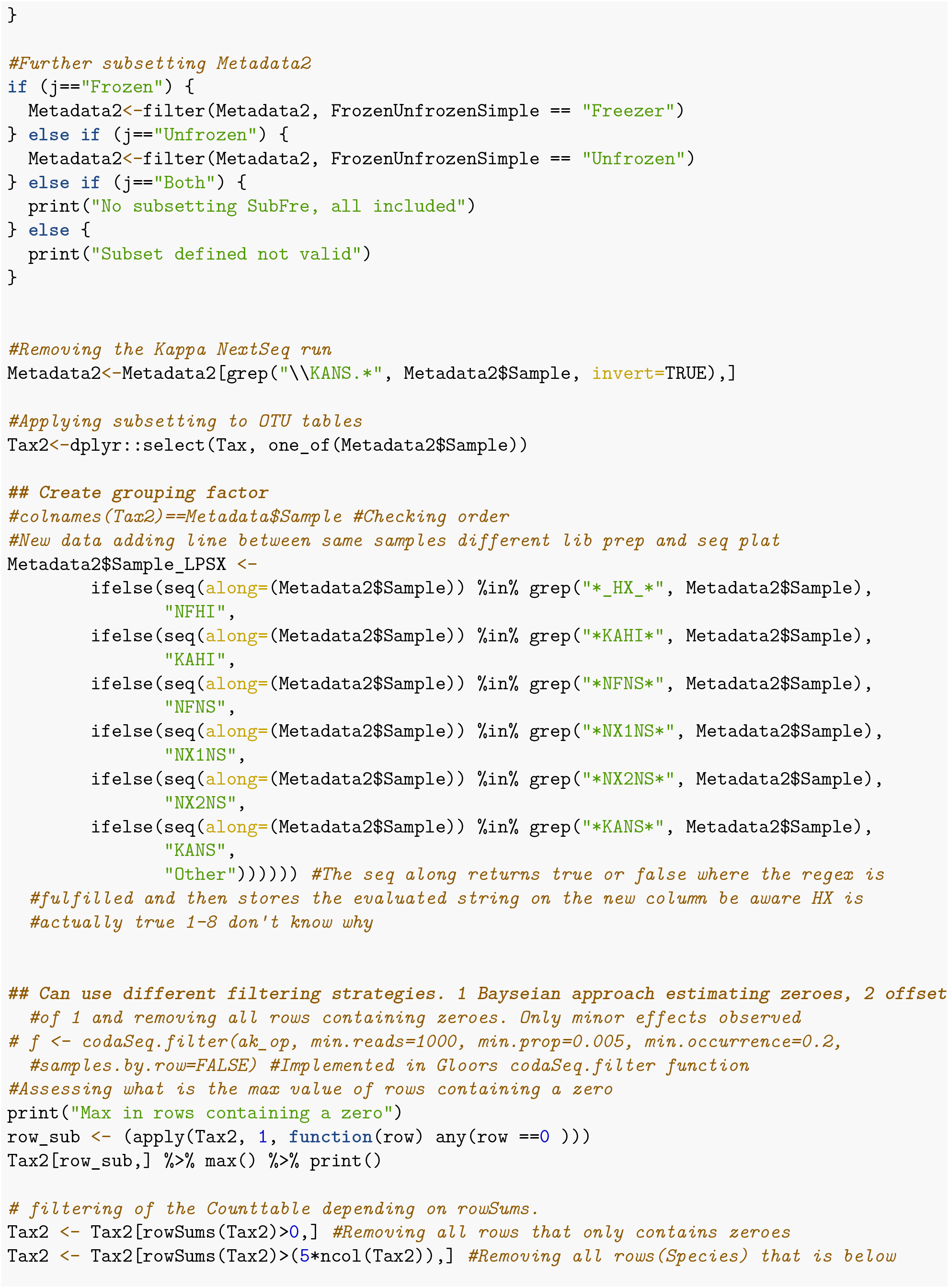
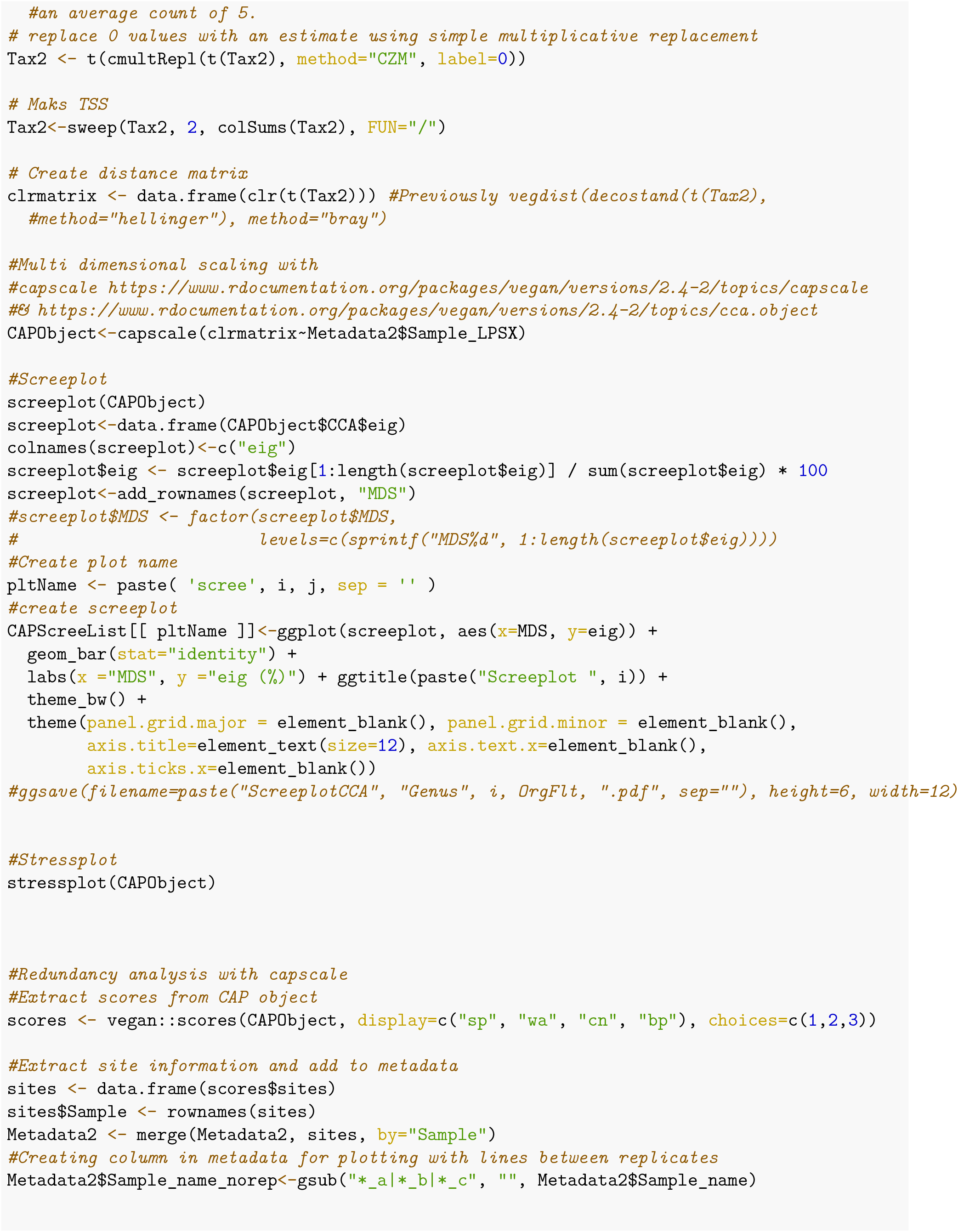
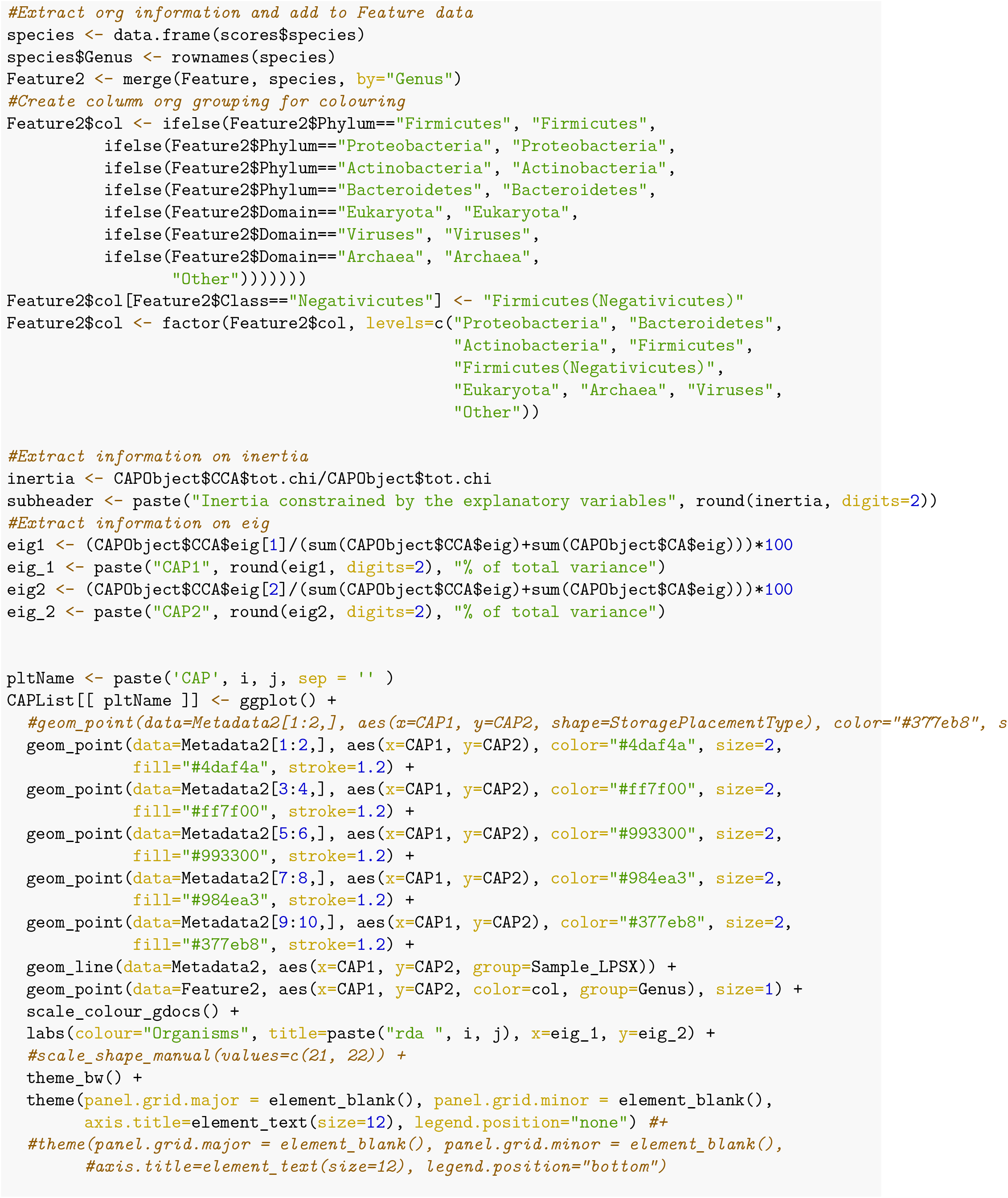
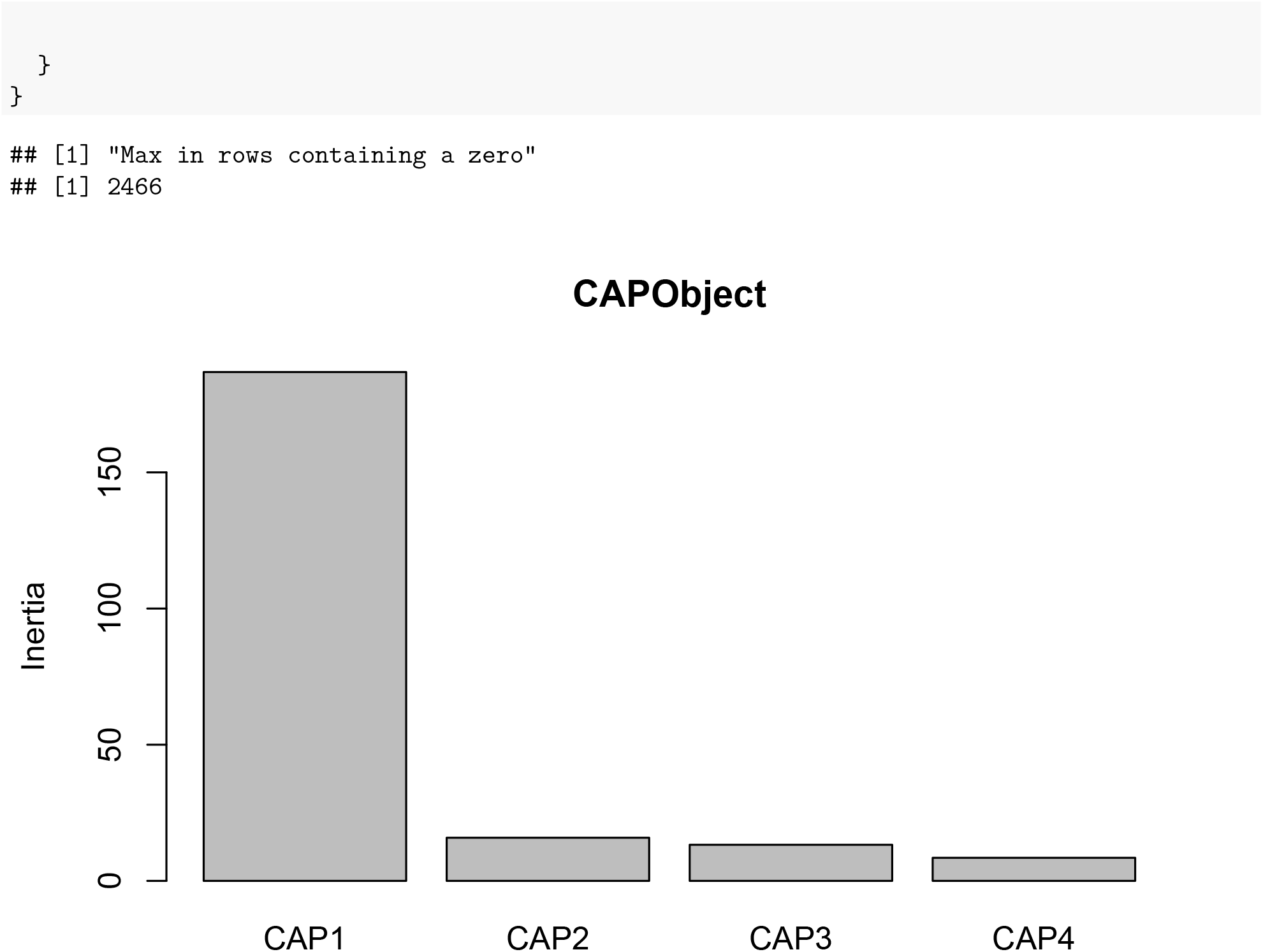
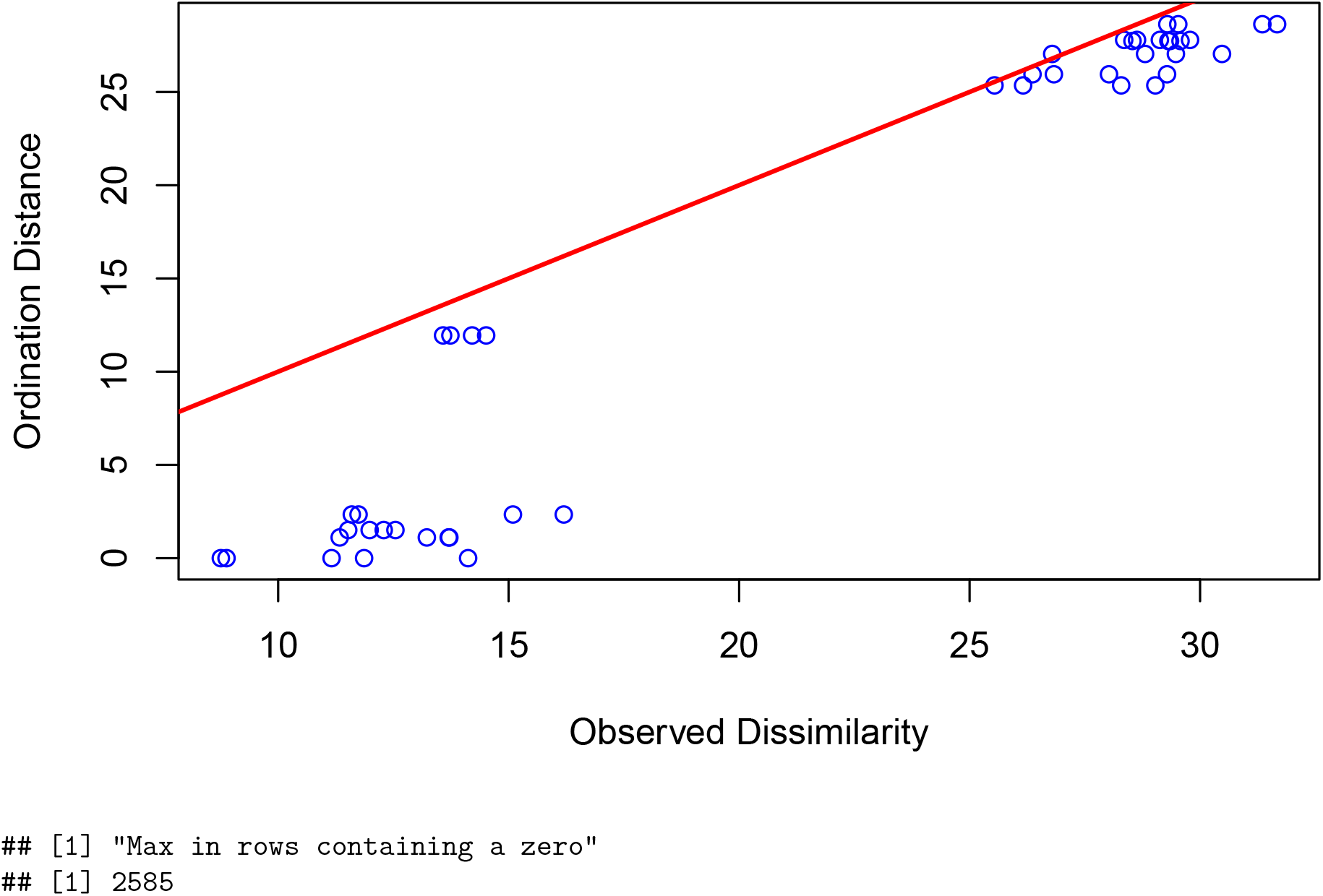
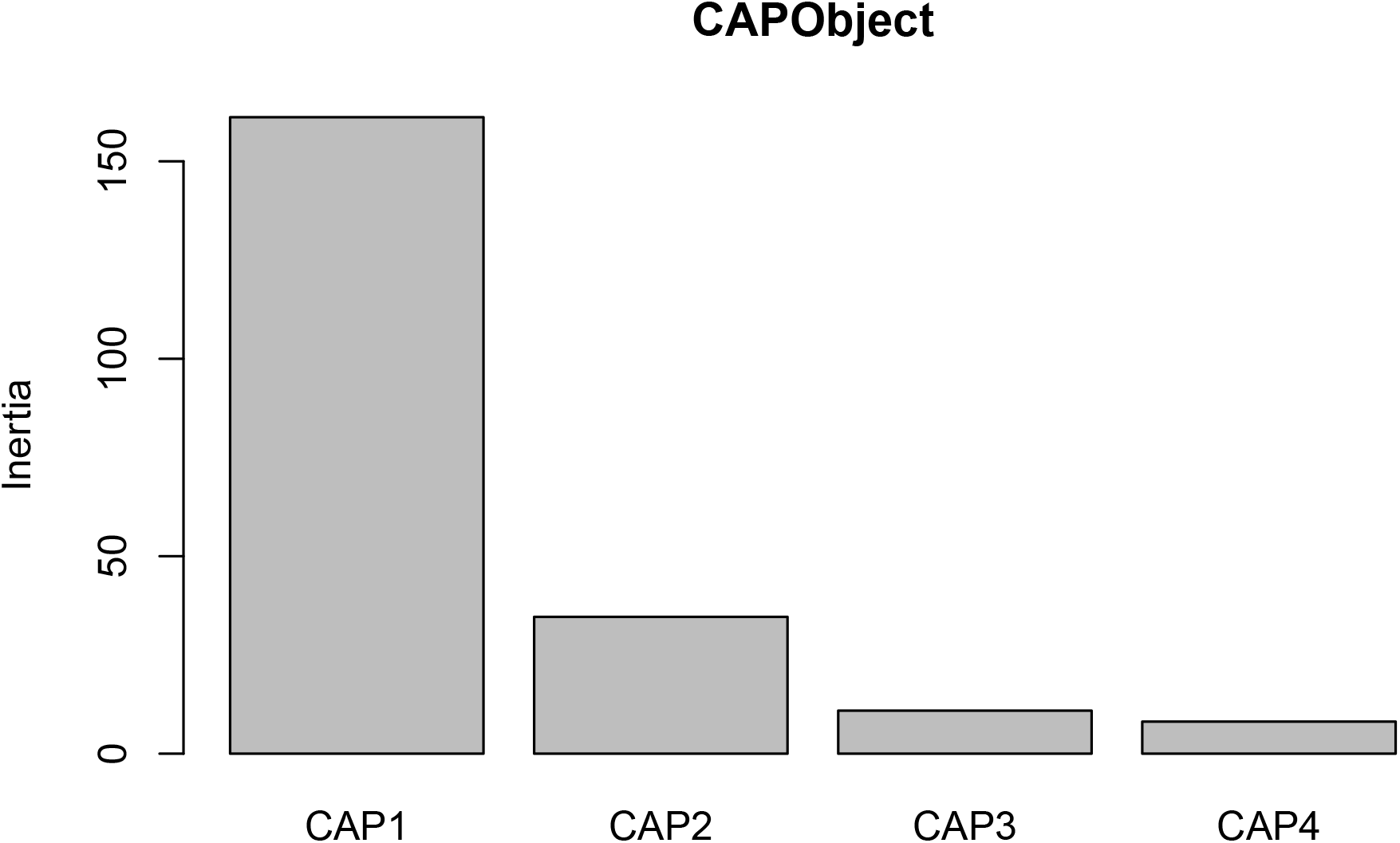
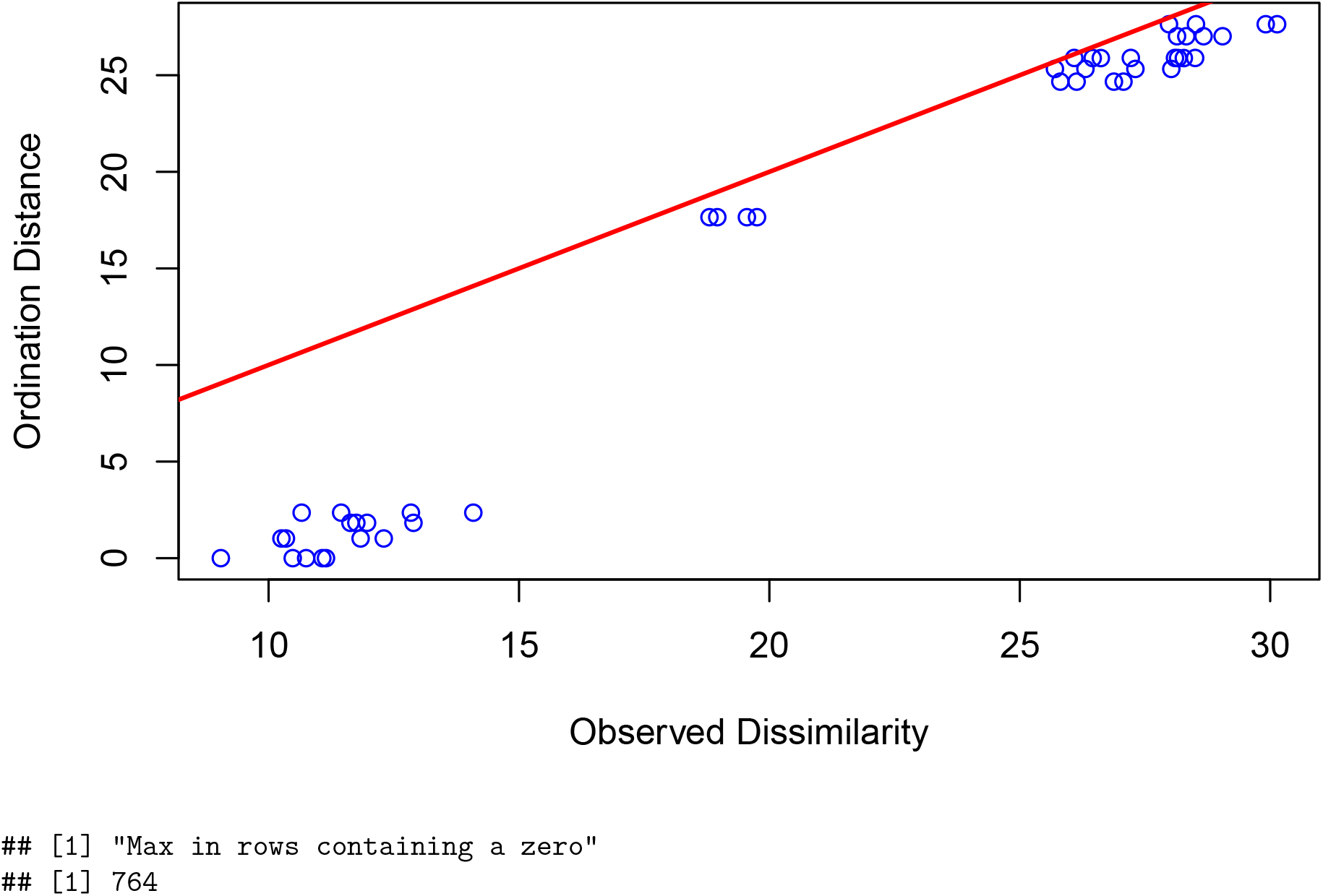
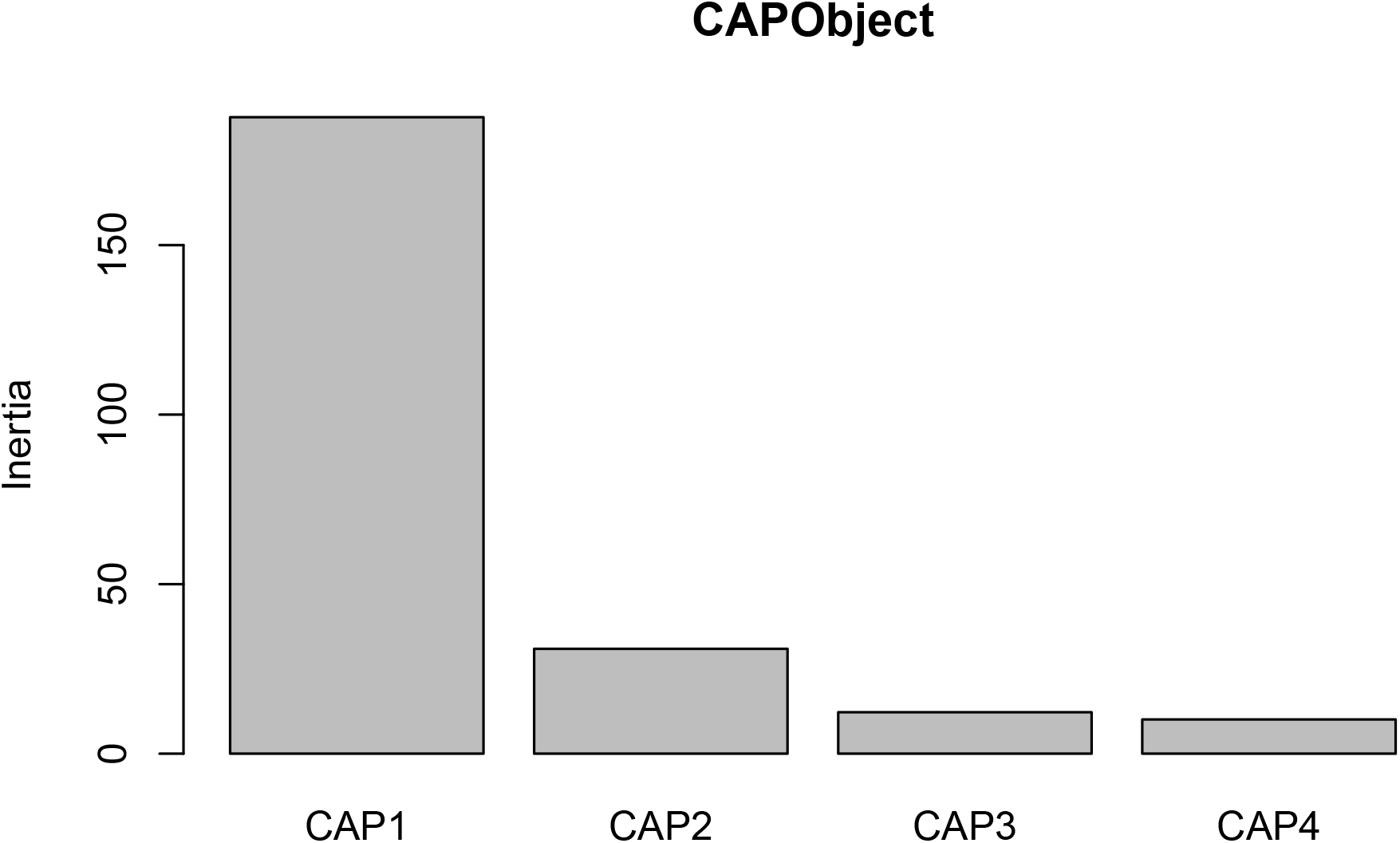
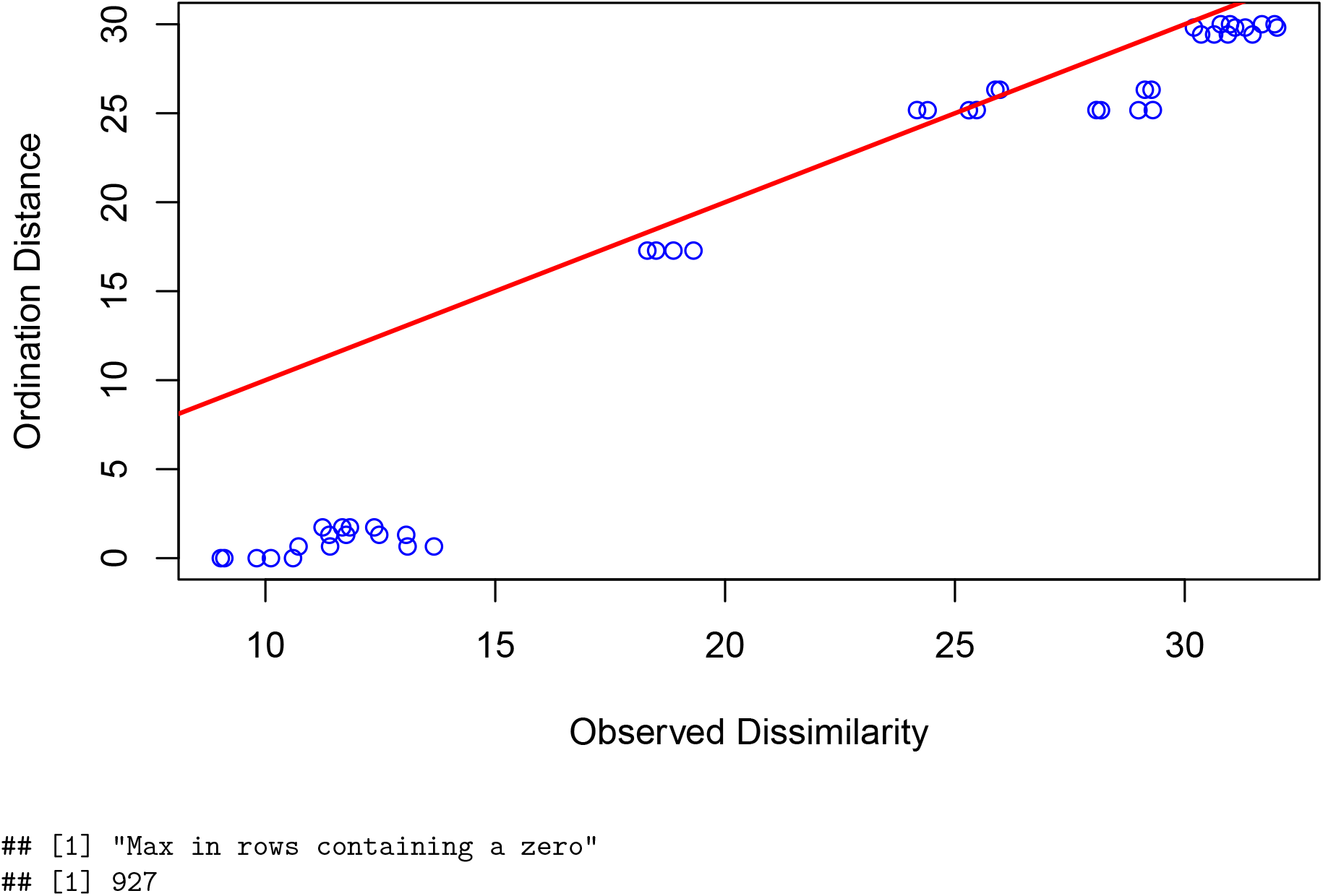
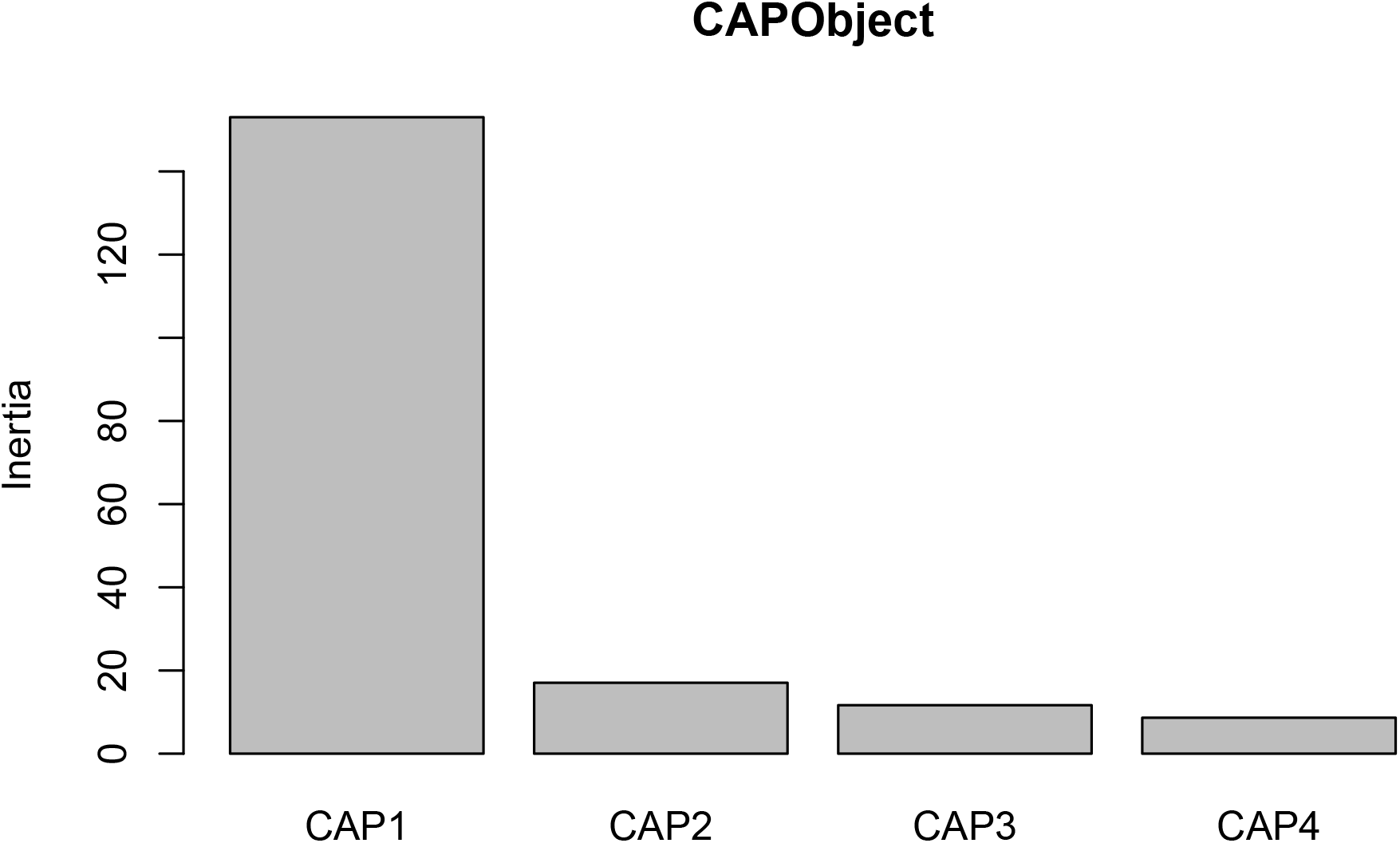
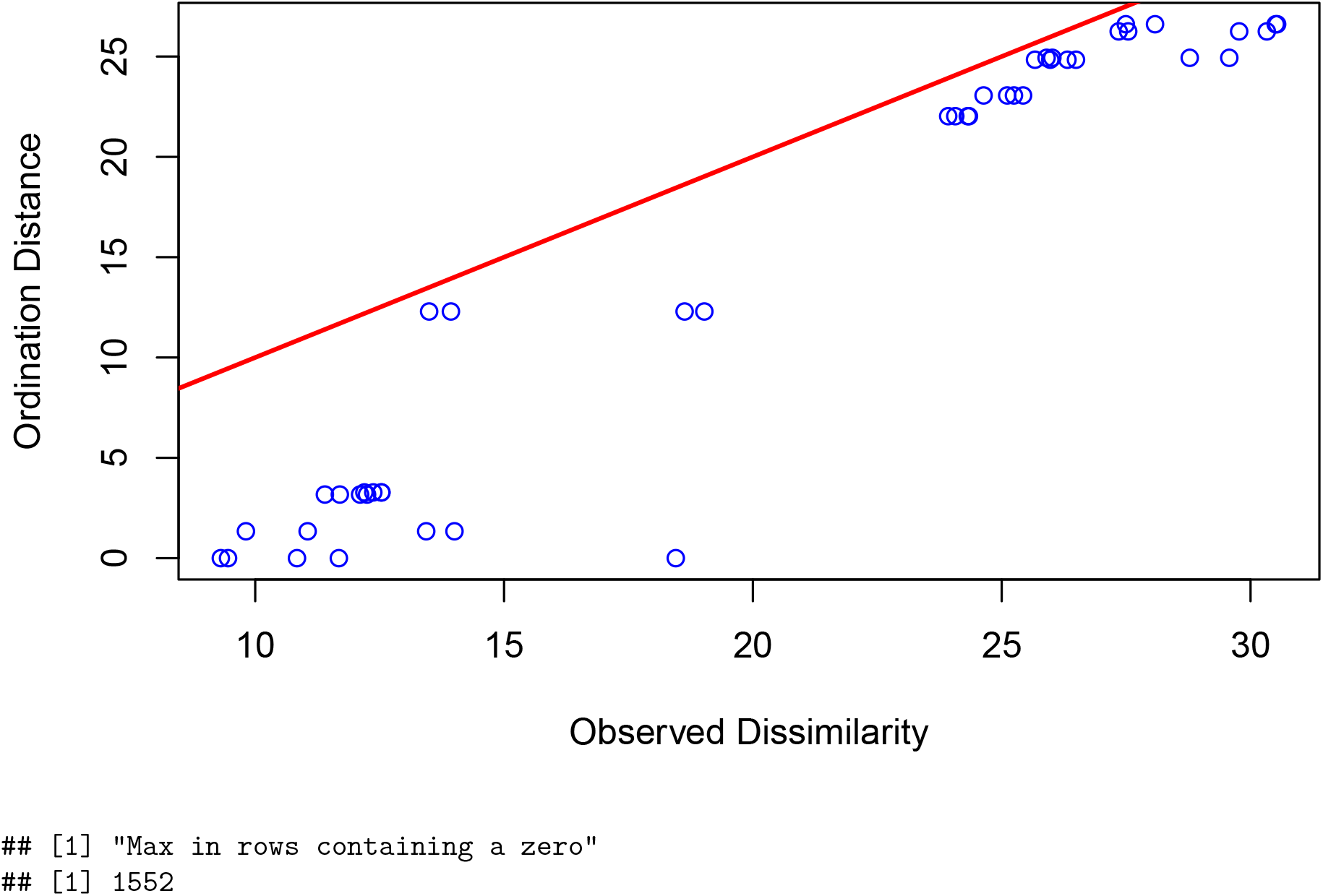
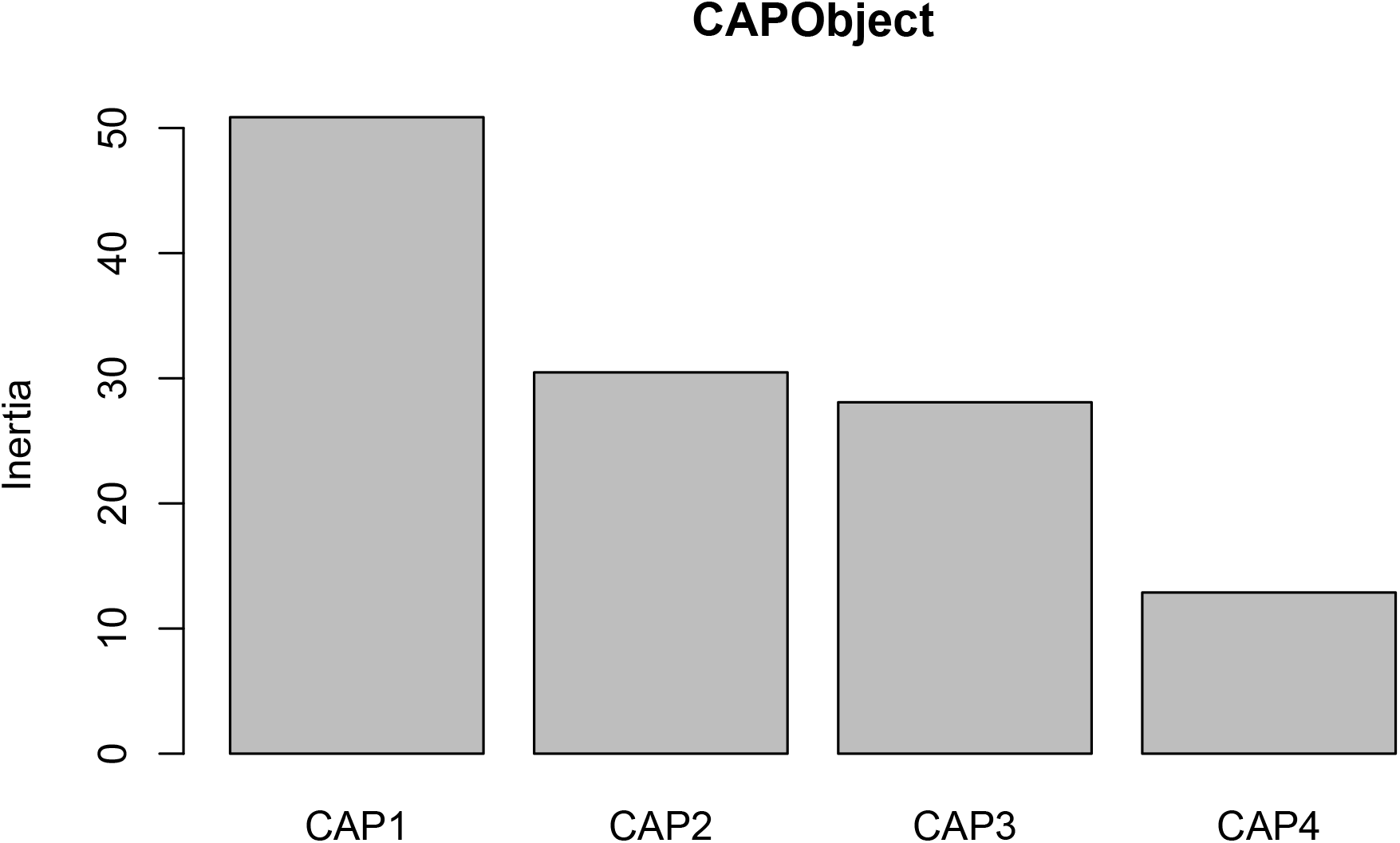
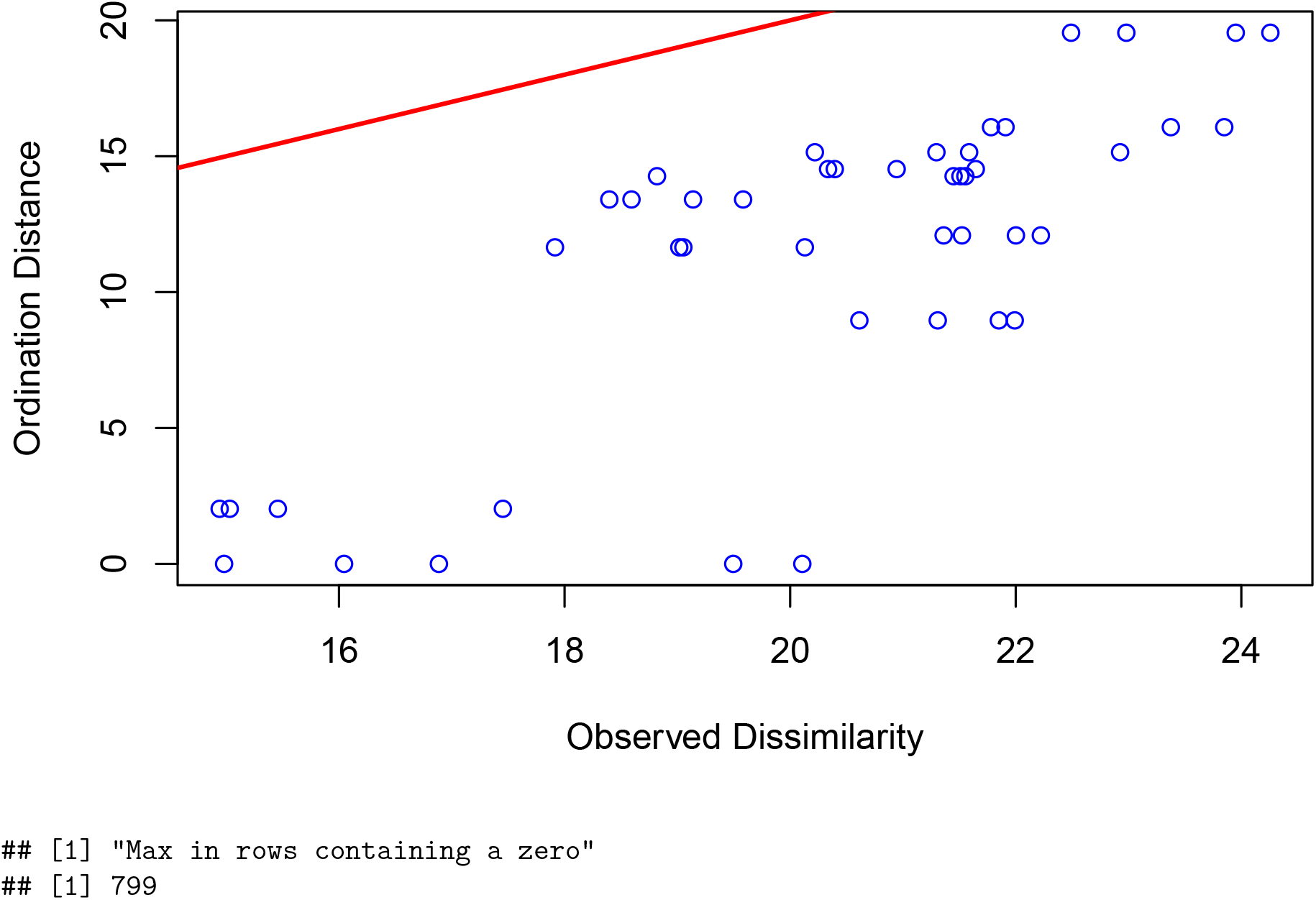
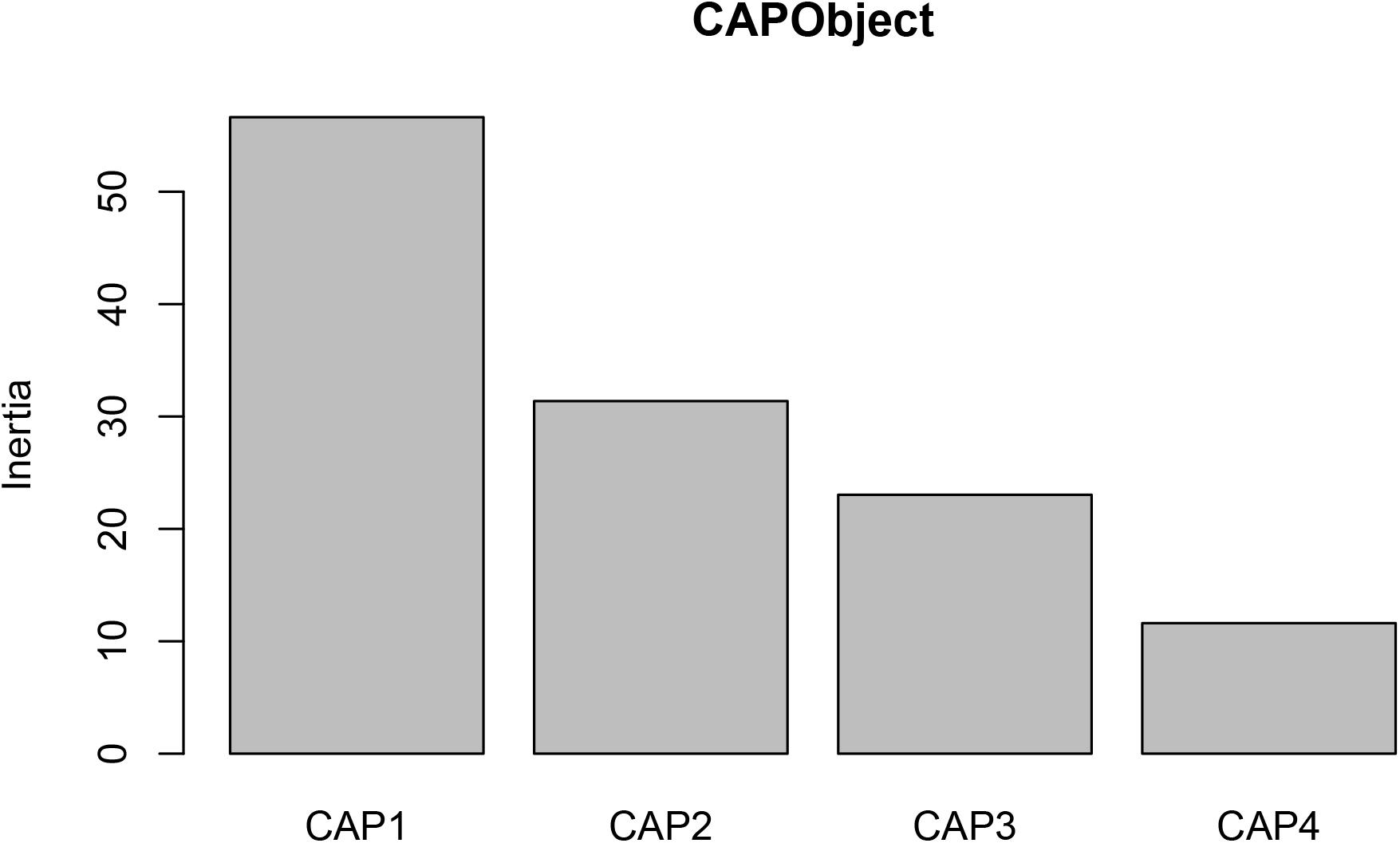
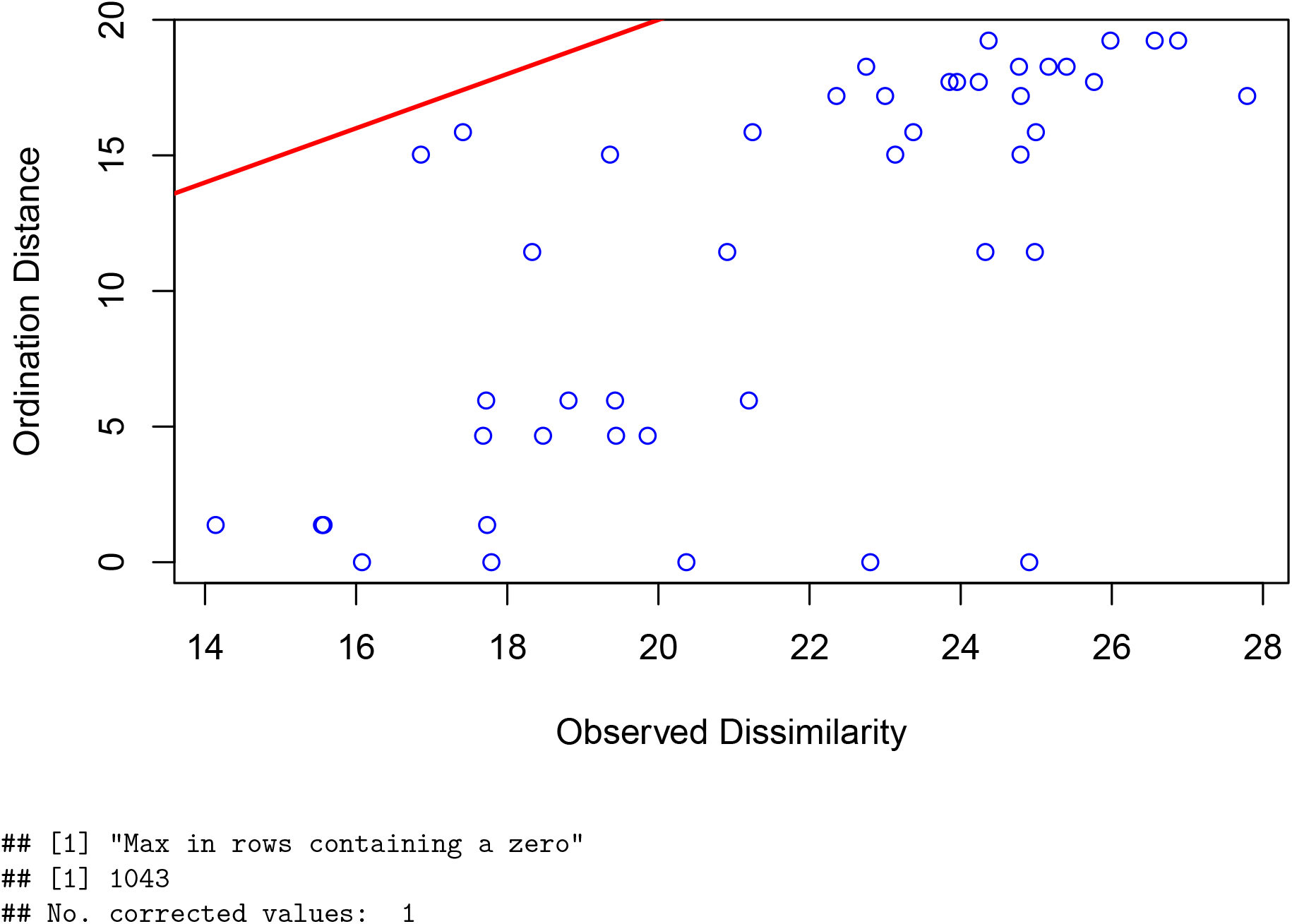
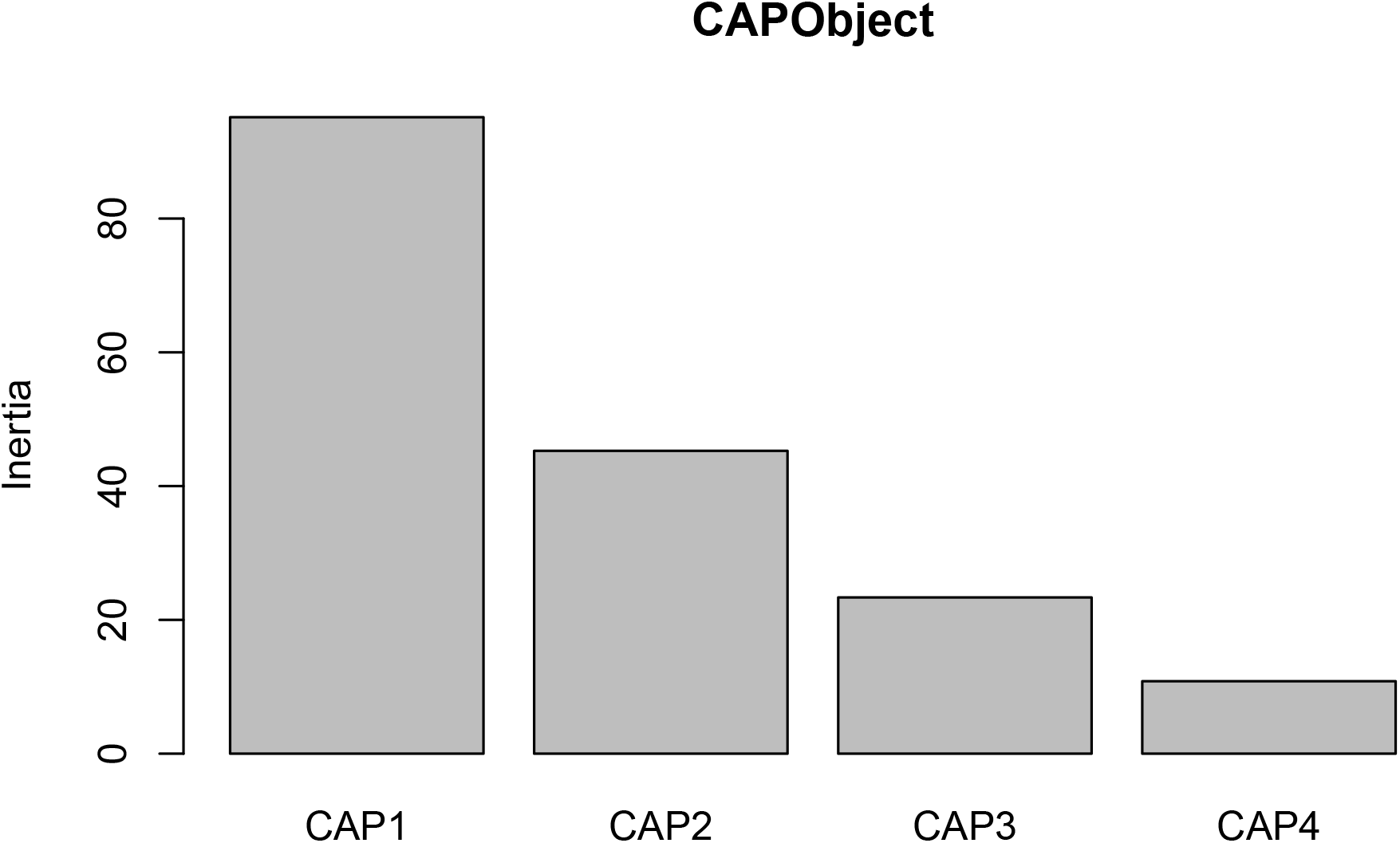
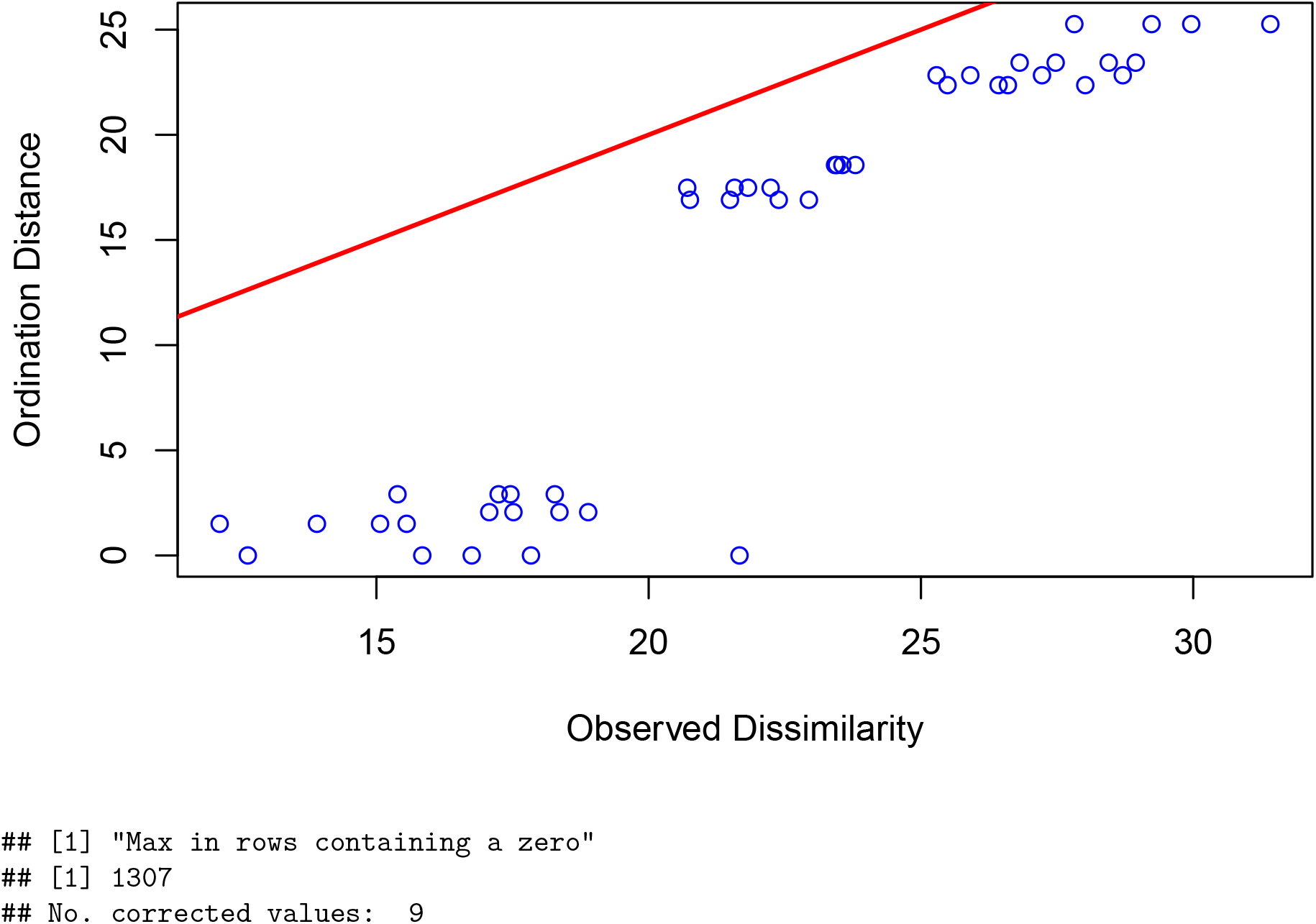
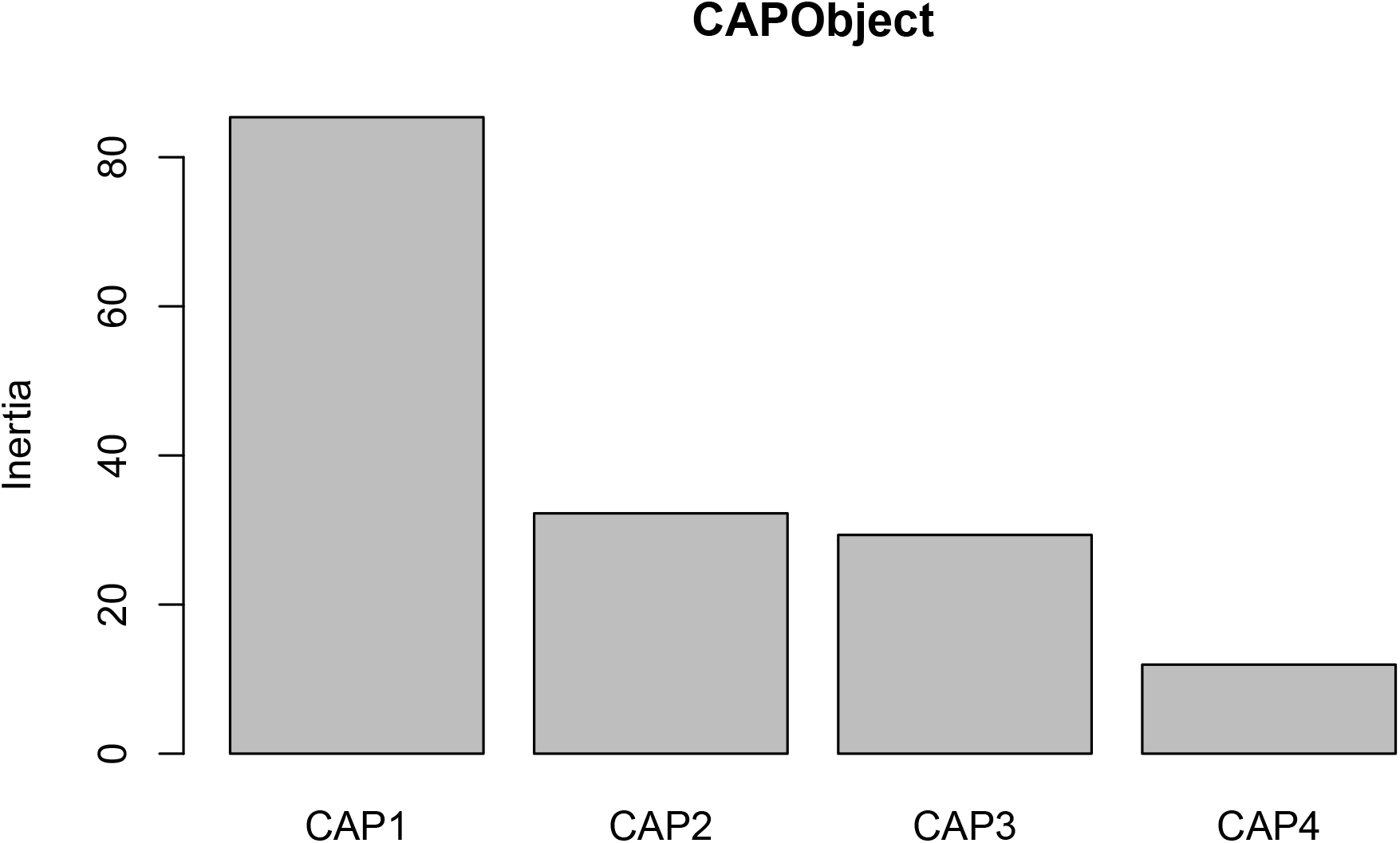
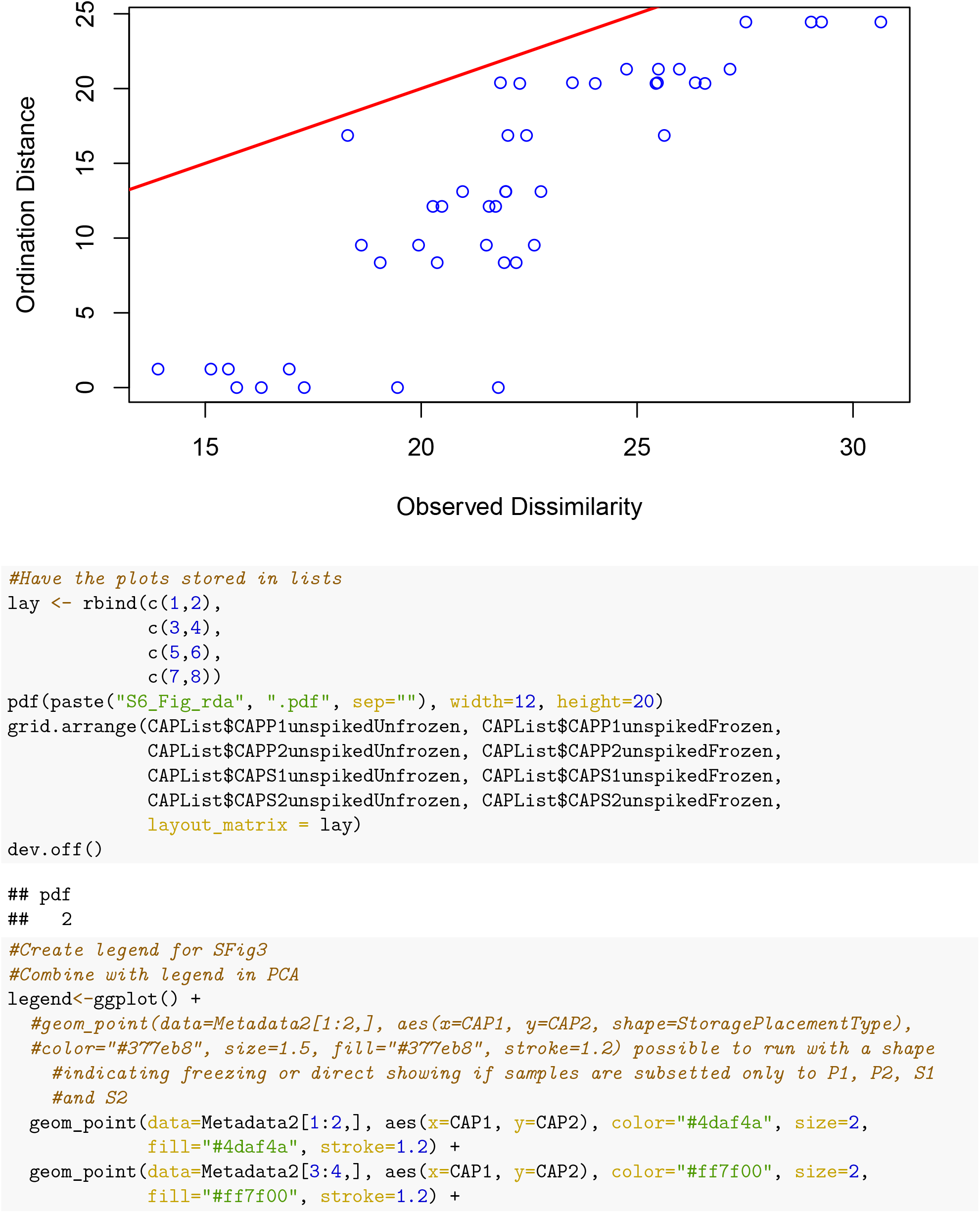
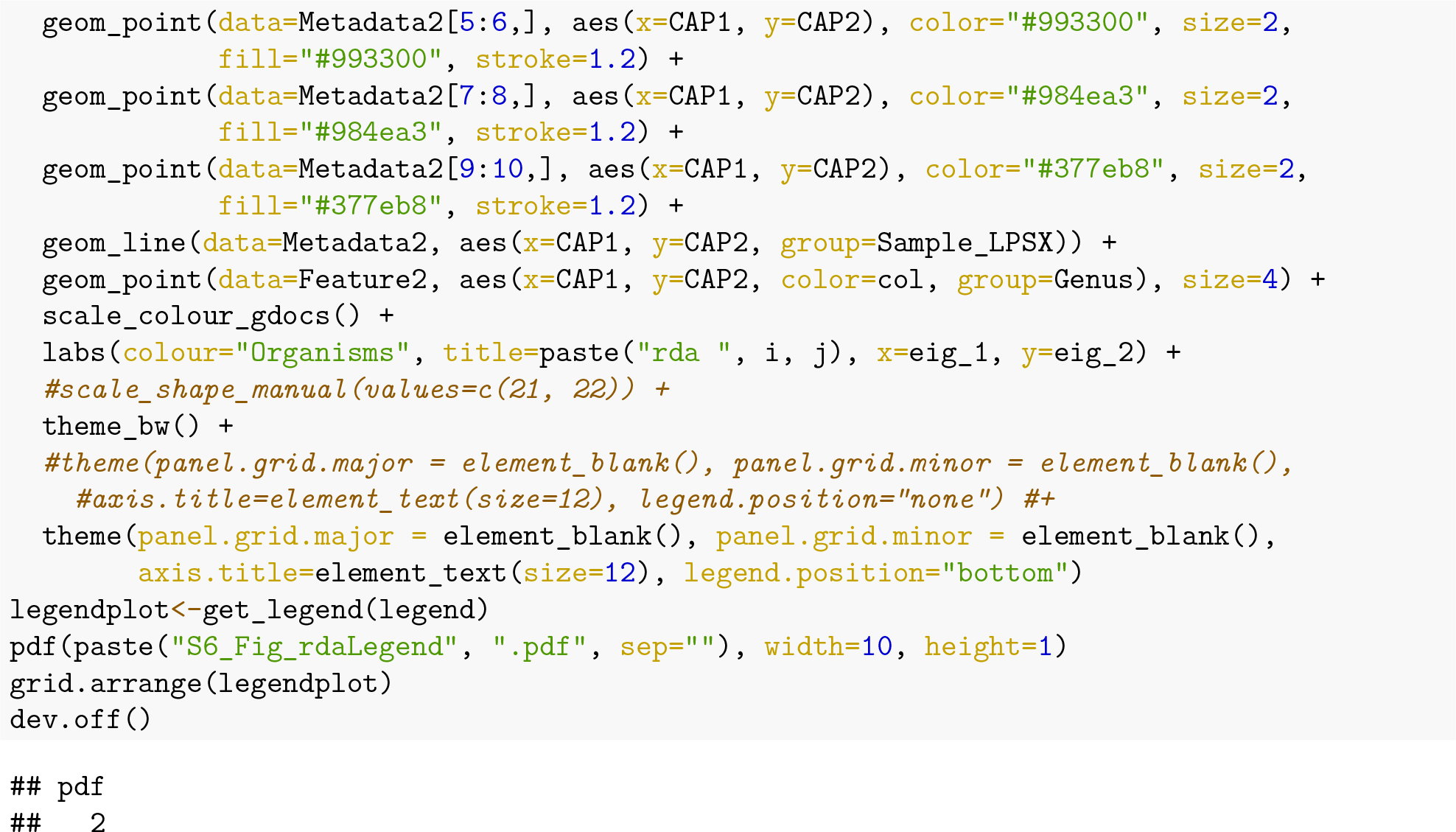

### Additional

#### Session information

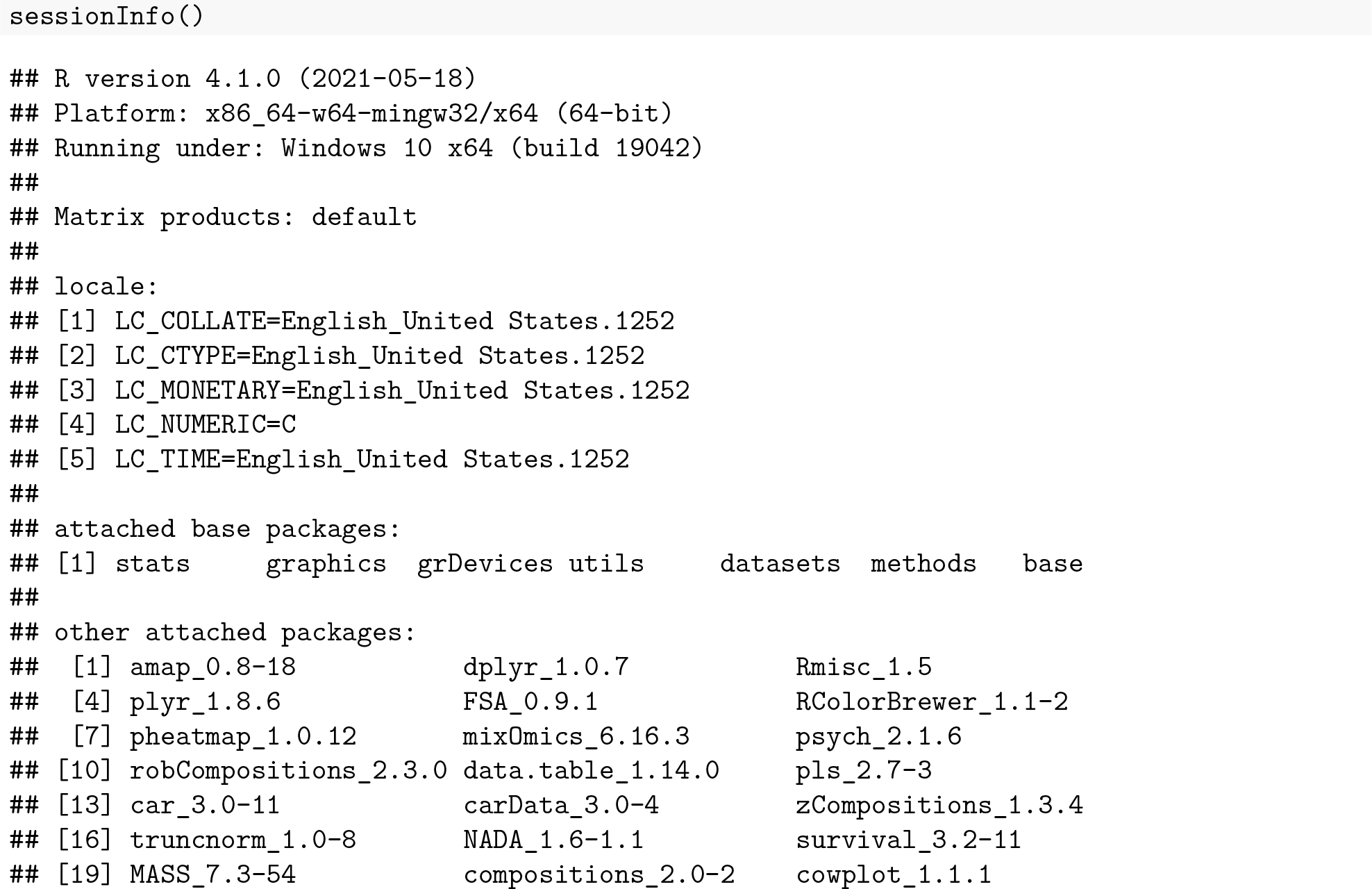
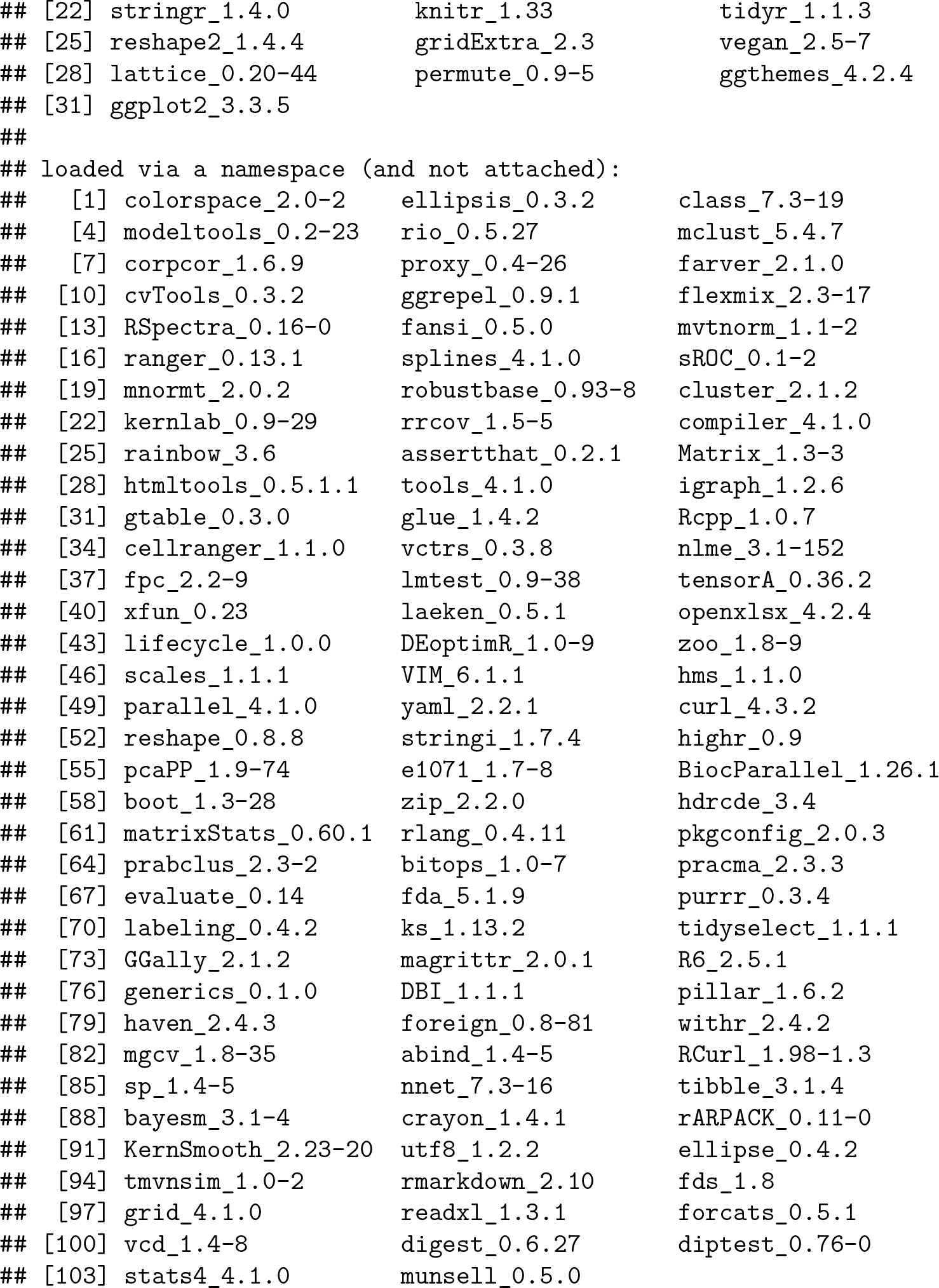

#### This document was processed on

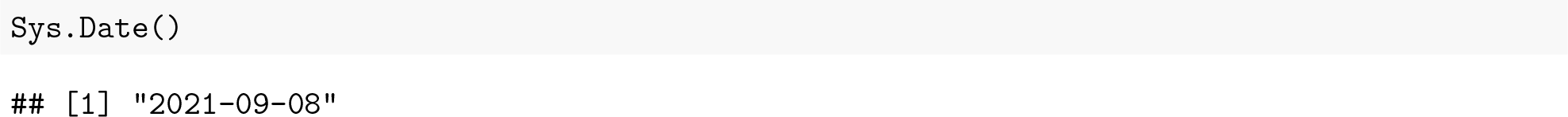

